# Cohort Profile: East London Genes & Health (ELGH), a community based population genomics and health study of British-Bangladeshi and British-Pakistani people

**DOI:** 10.1101/426163

**Authors:** Sarah Finer, Hilary C. Martin, Ahsan Khan, Karen A Hunt, Beverley MacLaughlin, Zaheer Ahmed, Richard Ashcroft, Ceri Durham, Daniel G MacArthur, Mark I McCarthy, John Robson, Bhavi Trivedi, Chris Griffiths, John Wright, Richard C Trembath, David A van Heel

## Abstract

- East London Genes & Health (ELGH) is a large scale, community genomics and health study (to date >34,000 volunteers; target 100,000 volunteers).
- ELGH was set up in 2015 to gain deeper understanding of health and disease, and underlying genetic influences, in British-Bangladeshi and British-Pakistani people living in east London.
- ELGH prioritises studies in areas important to, and identified by, the community it represents. Current priorities include cardiometabolic diseases and mental illness, these being of notably high prevalence and severity. However studies in any scientific area are possible, subject to community advisory group and ethical approval.
- ELGH combines health data science (using linked UK National Health Service (NHS) electronic health record data) with exome sequencing and SNP array genotyping to elucidate the genetic influence on health and disease, including the contribution from high rates of parental relatedness on rare genetic variation and homozygosity (autozygosity), in two understudied ethnic groups. Linkage to longitudinal health record data enables both retrospective and prospective analyses.
- Through Stage 2 studies, ELGH offers researchers the opportunity to undertake recall-by-genotype and/or recall-by-phenotype studies on volunteers. Sub-cohort, trial-within-cohort, and other study designs are possible.
- ELGH is a fully collaborative, open access resource, open to academic and life sciences industry scientific research partners.

## Why was the cohort set up?

East London Genes & Health (ELGH) is a community based, long-term study of health and disease in British-Bangladeshi and British-Pakistani people in east London. ELGH has a population-based design incorporating cutting-edge genomics with electronic health record (EHR) data linkage and targeted recall-by-genotype (RbG) studies. ELGH currently has >34,000 volunteers with funding to expand to 100,000 volunteers by 2023. ELGH is an open access data resource, and its research will impact a population at high need and redress the poor representation of non-White ethnic groups in existing population genomic cohorts^1^.

Almost a quarter of the world’s population, and 5% of the UK population, are of South Asian origin^2^. The risk of coronary heart disease is 3-4 times higher, and type 2 diabetes (T2D) 2-4 times higher in UK South Asians compared with Europeans^3,4^. East London incorporates one of the UK’s largest South Asian communities (29% of 1.95 million people), of which 70% are British-Bangladeshi and British-Pakistani, and its population live in high levels of deprivation (Tower Hamlets, Hackney, Barking and Dagenham are the 9^th^, 10^th^ and 11^th^ most deprived local authorities in England)^5^. Compared to White Europeans, South Asians living in east London have a two-fold greater risk of developing T2D^6^, nearly double the risk of non-alcoholic liver disease^7^, and over double the risk of multimorbidity^8^, with the onset of cardiovascular disease occurring 8 years earlier in men^8^. Determinants of poor cardiometabolic health start early in the life course, with higher rates of overweight and obese children in east London compared to the UK average^5^.

Recent genomic advances offer exciting potential to better understand the genetic causation of disease^9^, and to direct pharmacotherapy to rare loss-of-function gene variants^10^. Genetic variation relevant to British-Bangladeshi and British-Pakistani populations, such as autozygosity arising from parental relatedness, is under-researched with regards to potential effects on complex adult phenotypes at a population level^11,12^.

ELGH fosters authentic, inclusive, long-term engagement in its research, to deliver future health benefits to the population it represents. Community involvement in ELGH helps prioritise areas for research, including T2D, cardiovascular disease, dementia and mental health. ELGH undertakes a range of public engagement work, including collaboration with the award-winning Centre of the Cell^13^.

## Who is in the cohort?

ELGH (see ***Figure 1***) incorporates population-wide recruitment to Stage 1 studies, and targeted recruitment to Stage 2 recall-by-genotype (RbG) studies. Stage 3 and 4 studies are planned.

**Figure 1:**
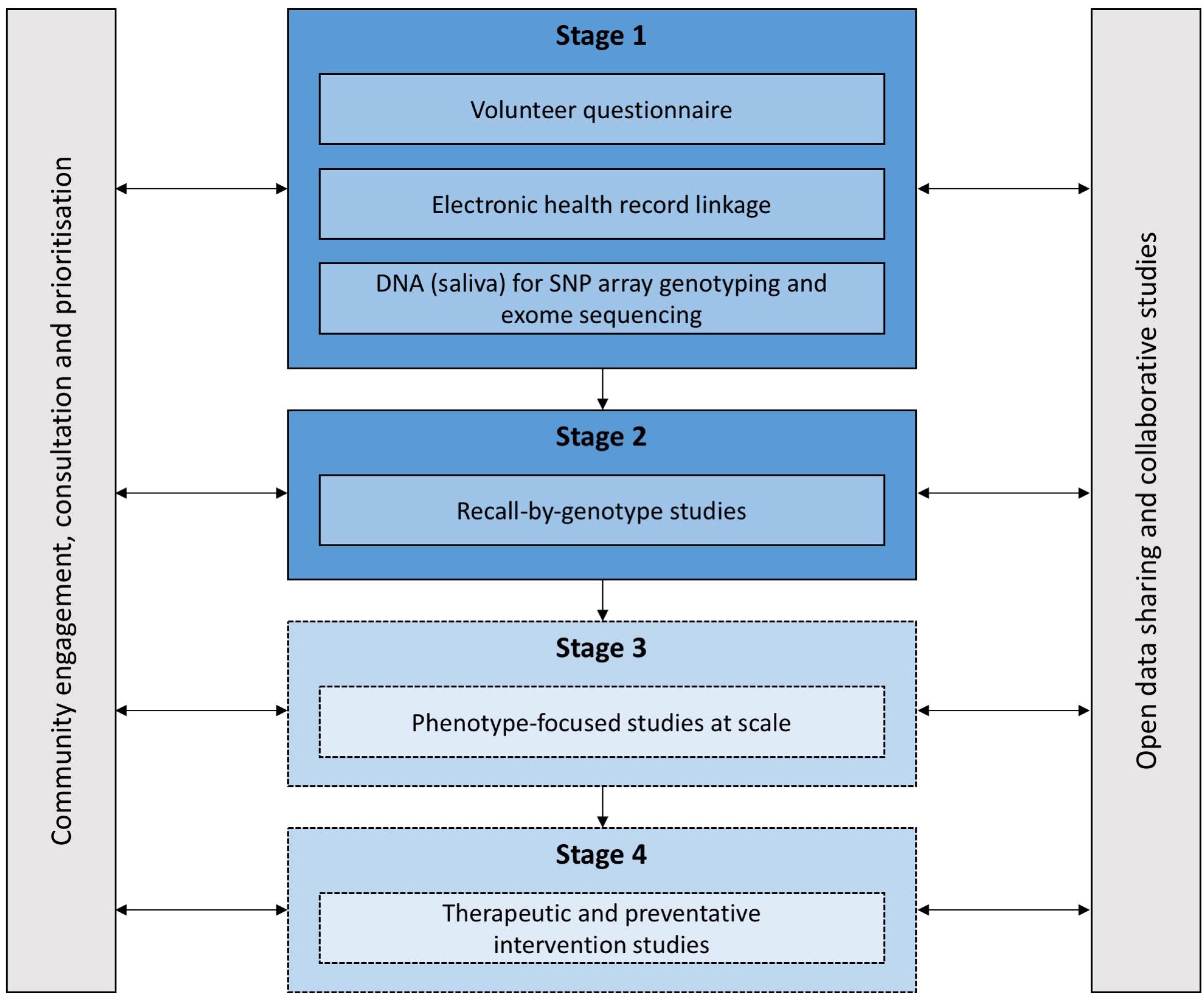
ELGH study design. Stage 1 and 2 studies have commenced. Stage 3 and 4 studies will commence in 2019.

During Stage 1, ELGH invites voluntary participation of all British-Bangladeshi and British-Pakistani individuals aged 16 and over, living in, working in, or within reach of, east London. Recruitment is largely undertaken by bilingual health researchers, and takes place in: (a) community settings, e.g. mosques, markets and libraries, supported by a third-sector partner organisation (Social Action for Health), and (b) healthcare settings, e.g. GP surgeries, outpatient clinics. Stage 1 volunteers complete a brief questionnaire, give consent to lifelong EHR linkage, and donate a saliva sample for DNA extraction and genetic tests. Between April 2015 and January 2019, ELGH has recruited 34,482 volunteers to Stage 1. At the most recent data linkage (November 2018), 97% of 31,646 had valid NHS numbers: 61% had linked primary care health record data available; 84% had linked secondary care data. By 2020, near-complete (>95%) linkage to primary care health records is expected with improved data connectivity, supported by Health Data Research UK. Recruitment into outer London regions, and a new study site in Bradford are planned for 2019/20, areas with similar ethnic populations and comparable health needs.

Summary data from the Stage 1 volunteer questionnaire and EHR data linkage are presented in ***Table 1***, including both baseline and longitudinal health data. Basic demographics of ELGH volunteers are compared to population-wide data in ***Figure 2*** and highlights that the convenience sampling approach in Stage 1 recruitment has achieved a sample broadly representative of the background population with regards age and sex, but which modestly favours recruitment of women over men in those <45 years. ELGH volunteers live in areas of high deprivation (97% in the most deprived 2 quintiles of the Index of Multiple Deprivation). Parental relatedness is reported by 19% of ELGH volunteers.

**Figure 2:**
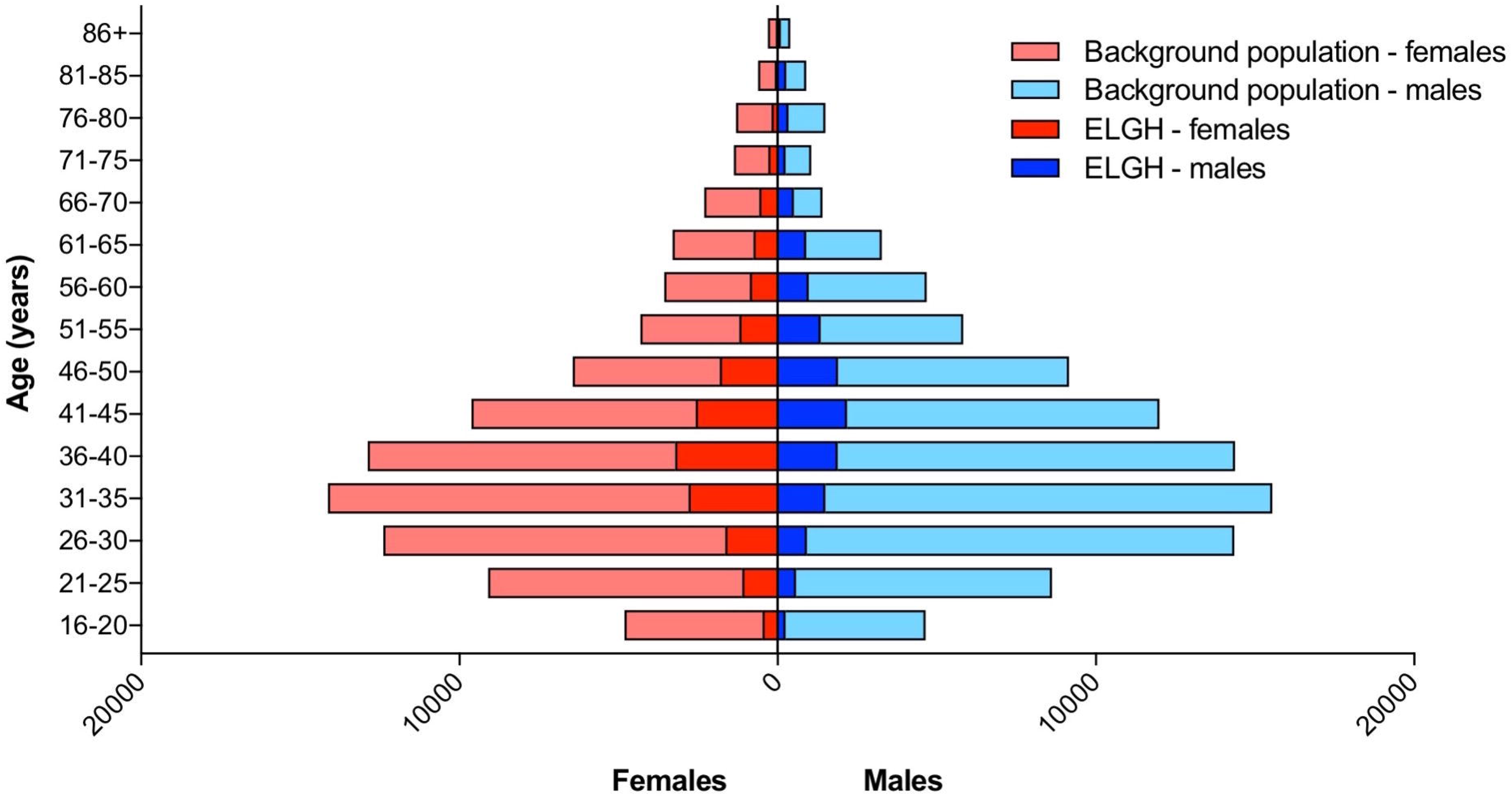
Population pyramid showing age and sex of ELGH volunteers (n=29,370) versus the total population of British-Bangladeshi and -Pakistani people (n=152,564) in East London (all NHS GP-registered adults residing in the London Boroughs of City and Hackney, Newham, Tower Hamlets, Waltham Forest), aged ≥ 16 years

**Table 1.** Baseline characteristics of ELGH volunteers from self-reported questionnaire and electronic health record data.

| Self-reported questionnaire data (n = 31646) |  |
| --- | --- |
| Year of birth n = 31634 | Median 1977 |
|  | Interquartile range 1967-1985; range 1915-2002 |
| Sex n = 31639 | Male n = 13928 (44%) |
|  | Female n = 17710 (56%) |
| Ethnicity n = 31508 | Bangladeshi or British Bangladeshi n = 21083 (67%) |
|  | Pakistani or British Pakistani n = 10319 (33%) |
| Parental relatedness | Yes = 5959 (18.8%) |
|  | No = 24601 (77.8%) |
|  | Don't know = 1013 (3.2%) |
|  | Not documented = 77 (0.2%) |
| Linked electronic health record data (n = 21514) |  |
| Index of Multiple Deprivation (2015) | Quintile 1 (most deprived), n = 12551 (58%) |
|  | Quintile 2, n = 8400 (39%) |
|  | Quintile 3, n = 411 (2%) |
|  | Quintile 4, n = 131 (0.6%) |
|  | Quintile 5 (least deprived), n = 11 (0%) |
| **Number of ELGH volunteers with common conditions:** |  |
| Type 2 diabetes | 4769 (22%) |
| Hypertension | 3956 (18%) |
| Ischaemic heart disease | 1048 (4.9%) |
| Dementia | 47 (0.2%) |
| Asthma | 2297 (10.7%) |
| COPD | 255 (1.2%) |
| **Number of ELGH volunteers with commonly recorded clinical data measured in the last 5 years:** |  |
| Body mass index | 18,654 (86.7%) |
|  | Mean = 27.16kg/m<sup>2</sup> |
|  | Standard deviation = 4.87kg/m<sup>2</sup> |
|  | Range = 14-69kg/m<sup>2</sup> |
| Smoking status | 18938 (88%) |
|  | Never smoked = 14353 (76%) |
|  | Ex-smoker = 2146 (11%) |
|  | Current smoker = 2276 (12%) |
|  | Uninformative coding = 194 (1%) |

ELGH operates under ethical approval, 14/LO/1240, from London South East NRES Committee of the Health Research Authority, dated 16 Sep 2014.

## How often have they been followed up?

ELGH contains real-world EHR data, its collection triggered by a broad range of clinical encounters including routine and emergency care. East London has an extensive track record of utilising routine clinical health care data (predominantly from primary care) in research studies^6,7,14^. Electronic performance dashboards are embedded in clinical practice, facilitating high quality and and equitable disease screening and clinical care^15, 16^. Primary care health records were digitised around 2000 and offer a rich source of data on clinical encounters since then, but also include pre-digitisation dates of diagnoses and summarised clinical events (e.g. type 2 diabetes, diagnosed in 1992). Health data linkage and extraction takes place 3-monthly and ELGH volunteers have consented for life long EHR access, facilitating longitudinal follow-up.

ELGH can invite volunteers to Stage 2 studies up to 4 times per year for more detailed study visits, e.g. recall by genotype (RbG) and/or phenotype, for clinical assessment and collection of biological samples, subject to ethics approval, volunteer acceptability and community advisory group approval. At September 2018, around 40 ELGH volunteers have participated in Stage 2 RbG studies.

## What has been measured?

Available data are summarised in Table 2:

- **Volunteer questionnaire** (*supplementary file 1*): This self-report questionnaire collects brief data including: name, date of birth, sex, ethnicity, contact details, diabetes status, parental relatedness, and overall assessment of general health and wellbeing. The questionnaire has been designed to facilitate high throughput recruitment and volunteer inclusivity where language and cultural differences exist, and to be used with or without researcher assistance. Completion of the volunteer questionnaire triggers health record data linkage via NHS number.
- **NHS primary care health record data linkage:** We first design data extraction in human-readable form (*Supplementary file 2*) and then code this in Structured Query Language (*Supplementary file 4*). Coded fields are extracted from EHR systems, curated to research phenotypes of interest, and developed both incrementally and on demand. Search terms are used, including READ2 diagnostic codes, prescribing data, laboratory test results and clinical measurements and processes. Data concordance was checked between volunteer questionnaires and their EHR, with >99% concordance for gender and year of birth. Almost all cases of data discordance were due to technical errors with questionnaire optical character recognition or user data completion, and were resolved with manual checking. Data outside clinically plausible ranges, or with clear data entry errors are removed. A detailed description of our data processing is in *Supplementary file 3*. Missing data exist, but at relatively low frequency in routinely collected and incentivised clinical measures, e.g. smoking status is recorded in the EHR of 88% of volunteers in the 5 years prior to the most recent data linkage. Repeated measures of routinely collected data, and cross-validation across information sources can mitigate the impact of missing data where it exists, as can statistical techniques, such as sensitivity analysis and multiple imputation^17^.
- **NHS secondary care health record data linkage:** linkage to Barts Health NHS Trust data provides secondary care data for all ELGH volunteers who have attended this hospital system (24,852 volunteers at the last linkage). Available data includes clinician-coded SNOMED-CT acute and chronic problem lists, laboratory and imaging results. OPCS-4 (Office for Population, Censuses and Surveys) and ICD-10 (World Health Organisation International Classification of Diseases and Related Health Problems) codes are available for every finished episode of care. For example, maternity data linkage within Barts Health identified 4,172 female ELGH volunteers with maternity records available for one or more pregnancies.
- **Planned linkage to health record datasets:** Linkage to other local hospitals (including those providing mental and community healthcare) is planned in 2019. ELGH wil also link to further datasets in 2019, including national NHS Hospital Episode Statistics (HES) and NHS Mortality Data^18, 19^ to include admissions and discharge, diagnosis and operation codes, maternity, psychiatric, critical care from 1997, and accident and emergency data, ICD-10 and OPCS-4 codes from 2008. NHS mortality data provides data on cause of death. Other planned data linkages to national registries include the National Cancer Registration and Analysis Service and National Cardiovascular Outcomes Research.
- **Genomics:** DNA is extracted from Oragene (DNA Genotek) saliva system and stored from all Stage 1 volunteers. To date, 20-40X depth exome sequencing has been performed (n=3,781) or is in progress (n=1,492) on volunteers reporting parental relatedness. By mid 2019 (funding secured) 50,000 samples from stage 1 volunteers will be genotyped on the Illumina Infinium Global Screening Array v3.0 (including 46,662 multi-disease variants)^20^. Array content includes rare disease-associated mutations (e.g. all pathogenic and likely pathogenic variants in ClinVar), pharmacogenetic associations and genome wide coverage for association studies, polygenic risk score, and Mendelian randomisation studies. In 2019/2020, if support is secured from an evolving Life Sciences Industry Consortium, or elsewhere, high-depth exome sequencing will be performed on up to 50,000 volunteer samples. The intention is for genotyping and high depth exome sequencing to be performed on up to 100,000 volunteer samples by 2023.
- **Samples for other-omics:** core study samples are taken from all volunteers recalled in stage 2 studies, including a blood cell pellet (for DNA, protein), plasma aliquots, and blood cell RNA preservation (Paxgene) for studies including methylation assays, transcriptomics, proteomics, lipidomics and metabolomics.

**Table 2:** Summary of all data types currently available in ELGH for Stage 1 volunteers, and planned for late 2018 onwards.

| Data source | Data fields | Volunteers | Data quality | Duration of data collection |
| --- | --- | --- | --- | --- |
| **Volunteer questionnaire** | Basic details: name, date of birth, ethnicity, address, GP, NHS number<br>Self-reported diabetes status<br>Self-reported parental relatedness<br>Self-assessment of overall health and wellbeing | All | High quality | Cross-sectional at study entry |
| **Electronic health records** | Primary care (GP) records (currently coded in READ2 or CTV3 clinical terminologies):<br>Sociodemographic data<br>Diagnoses<br>Prescribing data<br>Clinical measurements, e.g. height, weight, body mass index, blood pressure<br>Laboratory tests (e.g. blood tests)<br>Care processes, e.g. referrals<br>Quality and Outcomes Framework indicators: including care processes and outcomes for common diseases (e.g. diabetes, asthma, depression), public health concerns (smoking, obesity), and preventative measures (e.g. blood pressure checks)<br>NHS Health Checks: screening for diabetes, heart disease, kidney disease, stroke and dementia offered to 40-74 year olds | All volunteers where data linkage is possible (currently 61%, due to increase to >95% in 2019 with linkage to a wider GP practice network) | High quality | Real-world data with access to all available historic (retrospective) data and lifelong (prospective) data |
|  | Secondary care (hospital) records:<br>Diagnoses (ICD10) and procedures (OPCS-4)<br>Chronic problem listing (SNOMED)<br>Laboratory tests (e.g. blood tests, histopathology, microbiology)<br>Maternity records | All volunteers in contact with secondary (hospital) care | Moderate quality | Real-world data, includes retrospective data since 2012 and lifelong (prospective) data |
| **External health record data sets and registries** | Hospital Episode Statistics<br>Office for National Statistics mortality data<br>National Cancer Registration and Analysis Service, NCRAS<br>National Cardiovascular Outcomes Research: national audits | All volunteers (planned) | Moderate quality | Real-world data, includes retrospective data since 2003, and lifelong (prospective) data |
| **Genetic investigations** | Whole exome sequencing | All volunteers reporting parental relatedness (currently 19%) | High quality | Not applicable |
|  | Illumina Infinium Global Screening Array v3.0+MD | All volunteers up to 50,000 (to be undertaken in early 2019) | High quality | Not applicable |
| **Recall studies** | Bespoke clinical phenotyping and sample collection according to genotype of interest. Core samples taken on all for methylation assays, transcriptomics, lipidomics and metabolomics | 40 volunteers to date, with approval for all volunteers to be approached for recall up to 4 times per year | High quality | Dependent on protocol |

## What has it found: key findings and publications?

ELGH is a new resource that continues to grow in size and content and, to date, has been used for three main areas of work:

- **Characterisation of common phenotypes**

Using Type 2 diabetes (T2D) as an exemplar, we show the ability for detailed phenotypic characterisation of ELGH volunteers using EHRs **(*Table 3*)**. Of 21,514 volunteers in ELGH with available linked EHR data, 4,769 (22%) have a diagnosis of T2D in their primary care record. Basic sociodemographic data (age, gender, ethnicity) of volunteers was recorded in 100%, and smoking status had been obtained within 2 years of the most recent data linkage in 94%. In over 97% of volunteers with T2D, body mass index, markers of glucose control (HbA1c) and serum cholesterol were measured and available in the 2 years prior to ELGH participation. Hypertension, ischaemic heart disease and chronic kidney disease were observed in 47%, 15% and 11% of the 4,769, and erectile dysfunction was present in 26% of men. Retinal complications of T2D are recorded and graded, with 82% of volunteers having undergone screening within the last 2 years. Prescribing data show recent insulin prescriptions in 16%, and the use of single or multiple non-insulin agents, as well as use of cardiovascular drugs (e.g. lipid lowering therapy). These data show the potential to perform cross-sectional analyses in ELGH from EHR data.

**Table 3.**
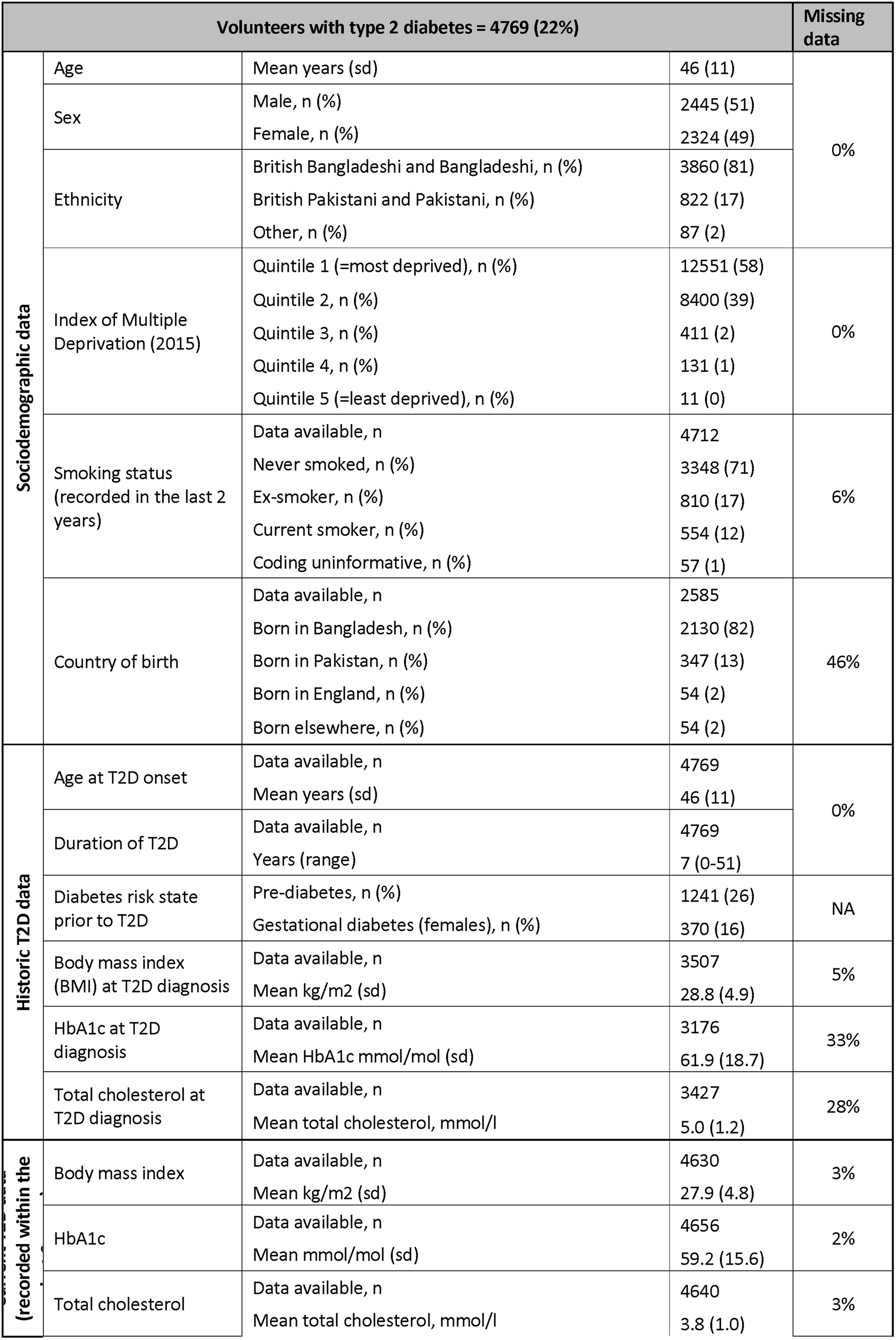
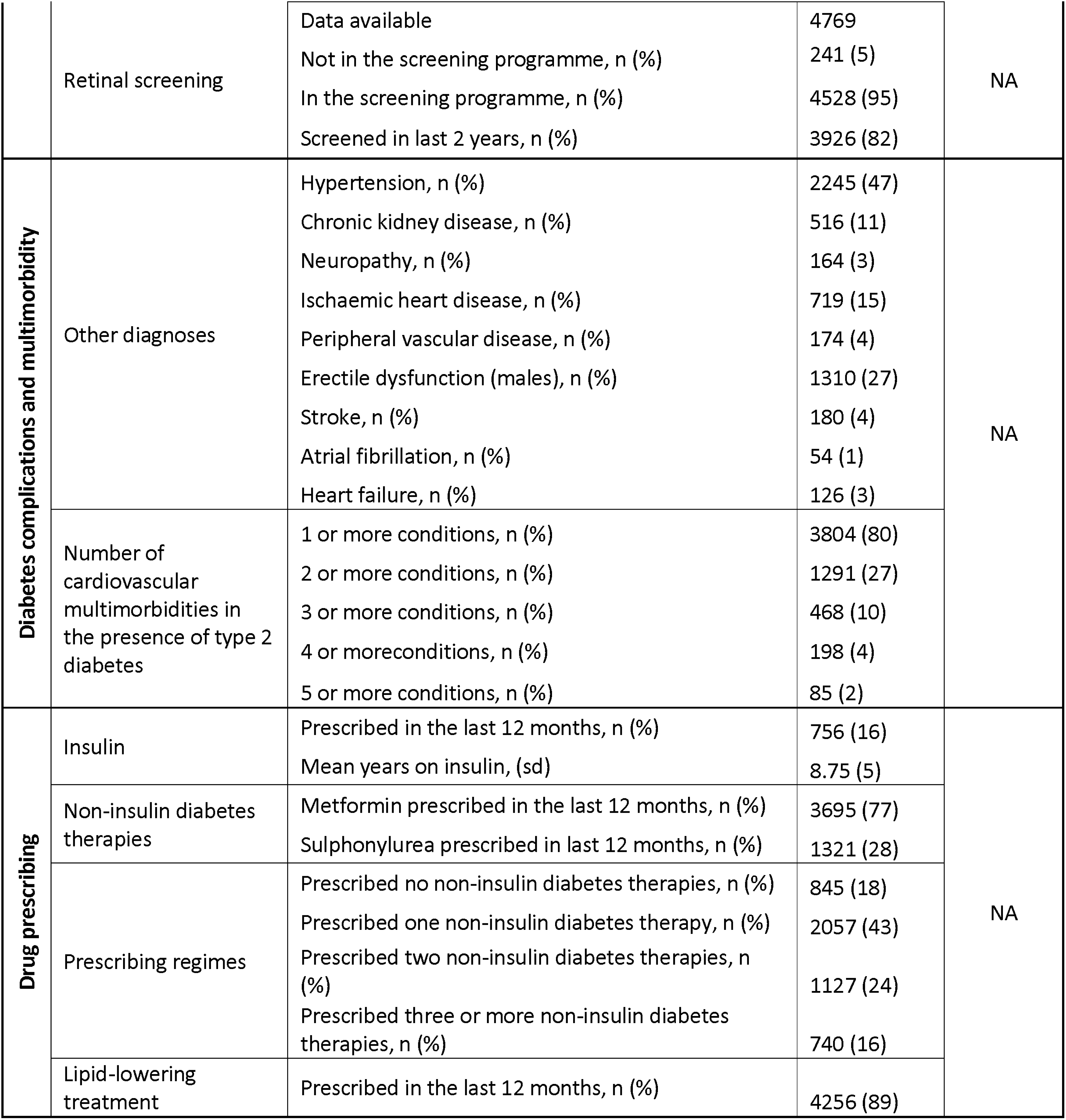
Example of a specific disease phenotype: characteristics of ELGH volunteers with type 2 diabetes. Data are presented in summary and descriptive formats as indicated. Missing data is estimated where available, e.g. for clinical care processes and measurements, but not diagnostic coding where the absence of a code is taken to indicate the absence of a diagnosis.

EHR data also gives the potential to study longitudinal phenotypic traits, retrospectively and prospectively. Median duration of T2D in ELGH volunteers was 7 years (range 0-51 years). For all volunteers with T2D, year of onset was recorded, and prescribing data and clinical measurements (including body mass index, HbA1c and cholesterol) at the time of diagnosis (+/− 6 months) were available for nearly two-thirds of volunteers. Prior to T2D onset, 26% (993) had had a diagnosis of pre-diabetes, and 16% (370) of women had had a diagnosis of gestational diabetes, allowing study of progression from at-risk to disease states.

Multimorbidity is an increasing problem in ageing populations with high rates of chronic long-term disease; in the ELGH population we identified that of the 4,769 ELGH volunteers with T2D, 80% had at at least 1, and 27% had 2 or more cardiovascular multimorbidities (***Table 3***).

- **Rare allele frequency gene variants occurring as homozygotes, including predicted loss of function knockouts.**

All ELGH volunteers self-reporting parental relatedness (19%) have been selected for exome sequencing. Genomic autozygosity (homozygous regions of the genome identical by descent from a recent common ancestor) means that rare allele frequency variants normally only seen as heterozygotes are enriched for homozygote genotypes. ELGH expands existing, smaller studies of autozygosity to investigate the health and population effects of such variants, with a focus on loss of function variants^11,21,22^. The accuracy of self-reported parental relatedness to actual autozygosity measured at the DNA level by exome sequencing (***Figure 3***) is a modest predictor of actual autozygosity, e.g. 8.2% of individuals who declare that their parents are not related in fact have >2.5% genomic autozygosity. For British-Bangladeshi volunteers, mean autozygosity is slightly lower than expected given the reported parental relationship (possibly due to confusion over the meaning of e.g. “first cousin” versus “second cousin”), whereas for British-Pakistani volunteers, mean autozygosity is slightly higher than expected (possibly due to historical parental relatedness).

- **Recall by genotype (and/or phenotype) studies**

**Figure 3.**
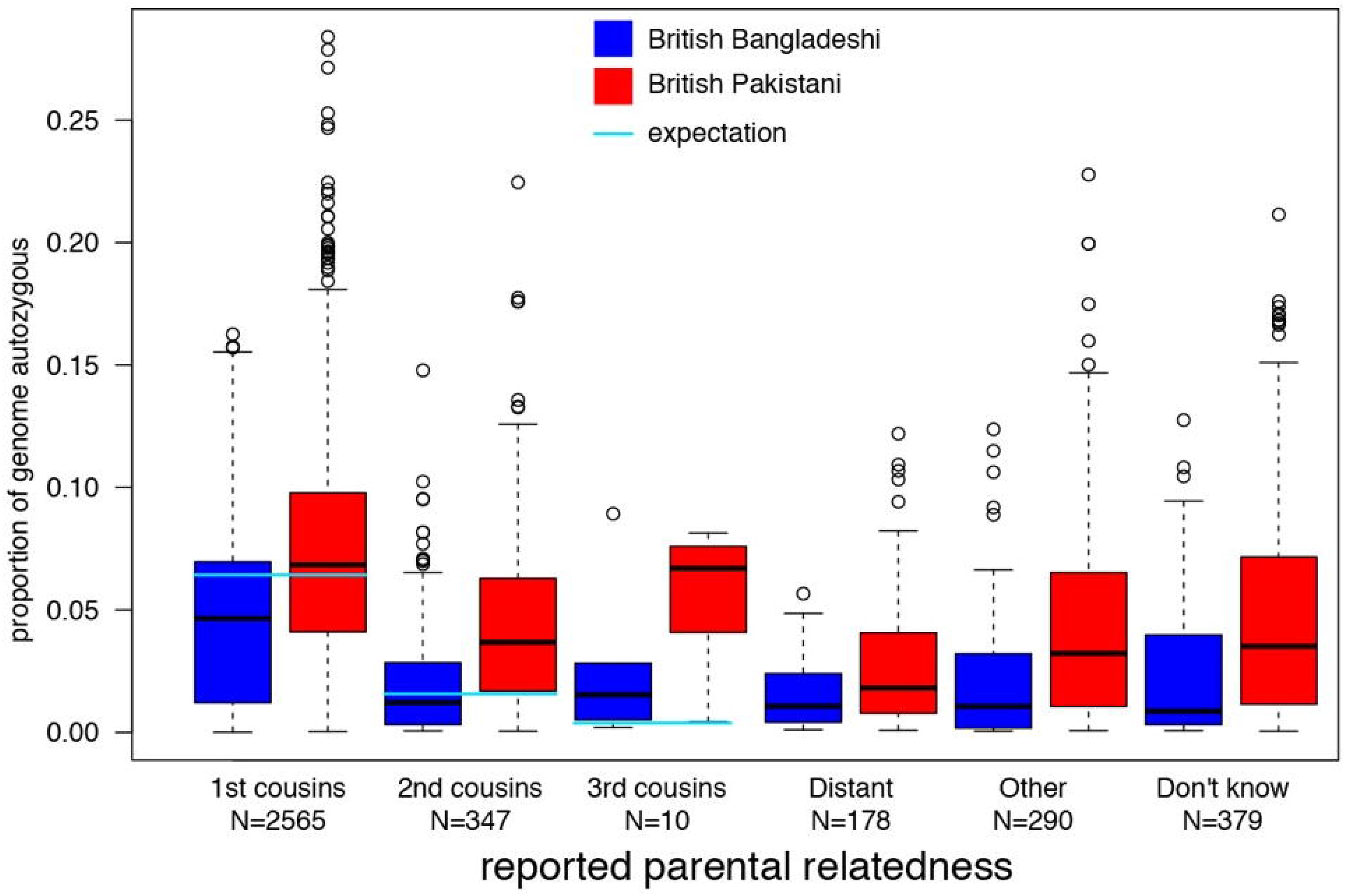
Distribution of levels of autozygosity as a fraction of the genome in ELGH volunteers, split according to self-reported parental relatedness and ethnicity (Tukey box plot showing median, lower and upper quartiles, quartiles +/− 1.5x interquartile range, and outliers).

RbG studies, applied to population cohorts with genomic data, are of increasing research interest^23^ and use the random allocation of alleles at conception (Mendelian Randomisation) to aid causal inference in population studies, reduce biases seen with observational studies, and develop functional studies. RbG studies can target specific single variants (or an allelic series for a gene) and polygenic variants (e.g. extremes of polygenic risk scores).

ELGH is undertaking RbG studies in Stage 2 using bespoke clinical phenotyping tailored to genotype or phenotype of interest. To date, three research consortia have undertaken ELGH RbG studies, one recalling volunteers with loss-of-function gene variants relevant to immune phenotypes, another phenotyping individuals with rare variants in genes implicated in T2D and obesity, and a third involving an industrial partnership to aid therapeutic development for a rare autosomal recessive metabolic disorder^24^. Successful recall completion rates to these RbG studies are between 30-40%.

## What are the main strengths and weaknesses?

ELGH has multiple strengths as a large, population-based study, and its novel, pragmatic design offers opportunities to combine genomic investigation with longitudinal and cross-sectional description of health and disease, and reflects a trend away from traditional epidemiological cohort design^25^. ELGH reaches a British-Bangladeshi and -Pakistani population with a high burden of disease, generalisable to a wider global population, and building on existing genetic studies that have been criticised for focusing on white European populations and have substantially under-recruited from minority ethnic groups^26^. High rates of autozygosity in ELGH volunteers lead to homozygous genotypes at variants with rare allele frequencies that will aid gene discovery, and RbG studies will generate novel translational impact^11,24^. Future studies on autozygosity will inform novel population level insights into the impact of genetic variation on health. The ability to invite all volunteers to Stage 2 studies offers the possibility to develop sub-cohorts and trials-within-cohorts in the future.

Our community-based recruitment approach offers broad reach into the target population however, to date, ELGH has modestly over-recruited British-Bangladeshi versus British-Pakistani volunteers. To support increased recruitment of British-Pakistani volunteers, recruitment is expanding into outer London boroughs and a new Bradford Genes & Health.

The use of real-world EHR data is both a strength and weakness of ELGH. Strengths include the ability to obtain high quality longitudinal data available on multiple diseases and disease risks via primary care in large numbers of volunteers in a feasible and cost-effective manner. Data linkage is not yet complete, but will improve in 2019 with improved infrastructure and linkage to national registries and databases. EHR data may be inferior to traditional observational studies in ascertaining some phenotypes, e.g. recent diseases of minor severity (which do not necessarily require healthcare access) or subclinical disease. Additionally, whilst outcomes can be studied relatively well, EHR data has limited opportunity to study certain exposures, e.g. health behaviours, diet or other environmental influences.

## Can I get hold of the data? Where can I find out more?

ELGH offers an open-access resource to international, academic and industrial researchers to drive high-impact, world-class science. Data access is managed at several levels, as follows:

- Level 1 - Fully open data. Summary data is distributed via our <u>website</u>, e.g. genotype counts and annotation of knockout variants from exome sequencing, and prevalence of phenotype and traits data.
- Level 2 - Genotype data (SNP chip genotyping, or high throughput sequencing) are (or will be) available under Data Access Agreements granted by the independent Wellcome Sanger Institute Data Access Committee. Individual sequencing (e.g. cram) and genotype files (e.g. vcf) are available within 6 months on the European Genome-phenome Archive^27^ (EGA).
- Level 3 - Individual-level phenotype data are held in an ISO27001 compliant Data Safe Haven environment under Data Access Agreement, currently hosted by the UK Secure e-Research Platform^28,29^. The data safe haven contains the latest genetic data linked to the questionnaire and health record phenotypes, and data export is tightly controlled. This “bring researchers to the data” model allows us to share regular data updates, maintain complex data linkages and avoid large file data transfers. This model provides robust reassurance to volunteers that their health data will be carefully looked after, with maximum security against data breaches.

External researchers can to apply to undertake research with ELGH via a formal application process, (details are available on the <u>website</u>), and most will be required to have their own research ethics approval to work with ELGH. Applications are assessed by both the Executive Board and Community Advisory Group, according to community prioritisation, acceptability and scientific merit.

## Acknowledgements

We acknowledge, with gratitude, the substantial contribution to ELGH from all of its volunteers who have provided samples and health record access, and have consented for recall. We thank the supporting recruitment teams and community organisations, including Social Action for Health. Invaluable support has also been received from the General Practitioners in the local health system who have made routinely collected health data available for research with support of the Queen Mary University of London Clinical Effectiveness Group. We are also grateful to the NHS National Institute for Health Research (NIHR) North-Thames Clinical Research Network (CRN), NHS Clinical Commissioning Groups (CCGs) (Tower Hamlets, City and Hackney, Newham, Waltham Forest, Barking and Dagenham), and Barts Health NHS Trust. ELGH looks forward to its expansion in 2019 in Bradford, Luton and Watford.

## Supplementary files

**Supplementary file 1:** Stage 1 volunteer questionnaire (.pdf file)

**Supplementary file 2:** Primary care electronic health record data extraction template (.xls file)

**Supplementary file 3:** Data processing protocol for primary care electronic health record data

**Supplementary file 4:** SQL code for primary care electronic health record data extraction

