## Supplementary File 3 for "Cohort Profile: East London Genes & Health (ELGH), a community based population genomics and health study of British-Bangladeshi and British-Pakistani people"

**Data processing protocol for primary care electronic health record data**

**Background**

Primary care electronic health record (EHR) data is linked via NHS number obtained via the volunteer questionnaire. Data linkage incorporates an NHS number validation step. Linked primary care electronic health record data are available for 68% (n = 21,514) of the total cohort (n = 31,646). Data are available for volunteers registered with a GP practice in the following London boroughs: Tower Hamlets, Newham, City and Hackney. At present, we do not have access to linked primary care EHR data for volunteers registered with GP practices in Waltham Forest, Luton, Barking, Havering, Redbridge and Essex, however data from practices in these boroughs will be available in future.

The primary care EHR dataset includes data items from the following categories:

- Demographic information
- Diagnoses
- Clinical observations
- Laboratory results
- Prescriptions

Data pertaining to all clinical terms (e.g. diagnoses and observations) is based on presence of relevant Read codes, version 2, (1) in the volunteer’s primary care EHR. All codes are extracted with the data at which it was entered and, where relevant, units of measurement. Primary care data extractions do not currently include free text noting of diagnoses in GP documentation.

Data are particularly rich in clinical contexts where routine data collection is standardised and incentivised, such as by the Quality and Outcomes Framework used in NHS primary care or supported by local quality improvement services which have been established over two decades.

Our most recent data linkage and download (R8) was undertaken on 12/11/18.

**Analysis summary:**

In the first instance, basic descriptive analyses of demographic information and disease counts were conducted, without data processing e.g. if a volunteer had a Read code for Type 1 diabetes as well as a Read Code for Type 2 diabetes, they were counted twice and contributed to the total number of both subgroups.

A more detailed descriptive analysis of a subgroup of volunteers with Type 2 diabetes was then conducted to describe the following:

- Demographic information
- Smoking status
- Retinal disease
- Multimorbidity
- Body mass index (at the time of diabetes diagnosis and latest, if taken in the past 2 years)
- HbA1c (at the time of diabetes diagnosis and latest, if taken in the past 2 years)
- Total Cholesterol (at the time of diabetes diagnosis and latest, if taken in the past 2 years)
- Prescribing for diabetes medications
- Prescribing for lipid lowering treatment and cardiovascular risk/disease

**Data processing:**

In the small number of instances where items of demographic data were not available in the primary care EHR dataset (date of birth and ethnicity), these data items were populated using data from the volunteer questionnaire by linking the unique volunteer questionnaire ID to their NHS number.

The following descriptive analyses were then performed using the methodology described below:

- **IMD 2015 Score**

Each volunteer’s Lower-layer Super Output Area code (available in the primary care EHR dataset) was matched to the corresponding IMD 2015 score as outlined by the Ministry of Housing, Communities and Local Government (2).

The proportion of volunteers with Type 2 diabetes (based on presence of a Read code for Type 2 diabetes in the primary care EHR) in each IMD 2015 quintile was then calculated.

- **Smoking status**

Where Read codes for two or more different smoking statuses were present in a volunteer’s primary care EHR and the dates of entry were the same, all smoking statuses for that volunteer were ignored (i.e. treated as if no smoking status was recorded).

Where Read codes for two or more different smoking statuses were present in a volunteer’s primary care EHR but dates of entry were different, only the Read code with the most recent date of entry was used for analysis.

Smoking status recorded in the past 5 years was summarised for all volunteers in the cohort, and smoking status recorded in the past 2 years was summarised for volunteers with Type 2 Diabetes (based on presence of a Read code for Type 2 diabetes in the primary care EHR).

- **Age at onset of Type 2 Diabetes**

The date of entry of a Read code for Type 2 diabetes was used as a proxy for the date of onset of Type 2 diabetes. The difference between the date of entry of a Read Code for Type 2 diabetes and the volunteer’s date of birth, was used to calculate the age at onset of Type 2 diabetes.

We did not exclude from the analysis volunteers for whom the Read code for Type 2 diabetes had the same date of entry as the date of registration at GP practice, even though it is likely that these reflect cases where the diagnosis of diabetes was already known at the time of registration and thus the date of onset of diabetes is not the date of entry of the Read code. For these volunteers, the implication is that the age at onset of diabetes is earlier than we have calculated using our methodology.

- **Duration of Type 2 diabetes**

The most recent primary care EHR dataset (R8) was extracted on 12/11/2018. The difference between the extraction date and the date of entry of a Read code for Type 2 diabetes was used to calculate the duration of Type 2 diabetes.

We did not exclude from the analysis volunteers for whom the Read code for Type 2 diabetes had the same date of entry as the date of registration at GP practice, even though it is likely that these reflect cases where the diagnosis of diabetes was already known at the time of registration and thus the date of onset of diabetes is not the date of entry of the Read code. For these volunteers, the implication is that the duration of disease is longer than we have calculated using our methodology.

- **Retinal disease**

In cases where a Read code for presence of retinal disease and a Read code for absence of retinal disease were both present in a volunteer’s primary care EHR and related to the same eye (i.e. left or right eye) and the dates of entry were the same, all retinal disease statuses for that eye were ignored (i.e. treated as if no Read code for retinal disease status was entered).

In cases where a Read code for presence of retinal disease and a Read code for absence of retinal disease were both present in a volunteer’s primary care EHR and related to the same eye but the dates of entry were different, only the Read code with the most recent date of entry was used for analysis.

- **Adult weight**

Weight data were available from the primary care EHR for the following time periods:

1. Earliest recorded weight
2. Weight recorded in a ± 6 month window around the time of diabetes diagnosis
3. Latest recorded weight

In cases where only one measurement of weight had been recorded for a volunteer, this value populated both the earliest recorded weight variable and the latest recorded weight variable.

Adult weight was defined as any measurement of weight taken at age ≥ 16 years. Any weight data taken at age < 16 years were not included in the analysis. A median weight (using all available weight variables if more than one weight measurement taken at age ≥ 16 years was present) was not calculated due to the variability per volunteer in the length of time that may have passed between each of the available weight variables, and due to the degree to which weight fluctuates over time.

Instead, we calculated time (in years) since the latest recorded weight for all volunteers and created two subgroups to perform further analyses:

1. Participants with a weight measurement in the past 5 years
2. Participants with a weight measurement in the past 2 years AND a Read code for Type 2 diabetes

All adult weights < 20kg were excluded from the analyses.

- **Adult height**

Height data were available from the primary care EHR for the following time periods:

1. Earliest recorded height
2. Height recorded in a ± 6 month window around the time of diabetes diagnosis
3. Latest recorded height

In cases where only one measurement of height was taken, this value populated both the earliest recorded height and latest recorded height variables.

Adult height was defined as any measurement of height taken at age ≥ 16 years. Height data taken at age < 16 years were not included in the analysis.  To inform later calculation of body mass index, and given that individuals typically have height measured less frequently than weight and that it varies less across the lifecourse, a median adult height (using all available height variables if more than one height measurement taken at age ≥ 16 years was present) was calculated for each volunteer, once the below data quality steps were performed.

1. In cases where age was ≥ 16 years and unit of measurement was absent (blank) and value was between 10 and 30 (inclusive), three assumptions were made:The unit was omitted in error
2. The intention was to enter the unit ‘metres’
3. A decimal place had been omitted in error

Based on these three assumptions, the unit of measurement of ‘metres’ was assigned to the value and the value was converted to the equivalent value in metres using the conversion calculation: [ x/10] where x represents the original value.

In cases where age was ≥ 16 years and unit of measurement was ‘centimetres’ and value was between 10 and 30 (inclusive), three assumptions were made:

1. The unit was entered in error
2. The intention was to enter the unit ‘metres’
3. A decimal place had been omitted in error

Based on these three assumptions, the unit of measurement of ‘centimetres’ was changed to ‘metres’ and the value was converted to the equivalent value in metres using the conversion calculation: [ x/10] where x represents the original value.

In cases where age was ≥ 16 years and unit of measurement was ‘centimetres’ and value was ≤ 3, the assumption was made that the unit of ‘centimetres’ was recorded in error and the unit of measurement was changed to ‘metres’, leaving the value intact.

In cases where age was ≥ 16 years and unit of measurement was ‘metres’ and value was ≥3, the assumption was made that the unit of ‘metres’ was recorded in error and the unit of measurement was changed to ‘centimetres’, leaving the value intact.

Once the above data quality steps were performed, all remaining values where age was ≥ 16 years and unit of measurement was ‘centimetres’ were converted to the equivalent value in metres using the conversion calculation: [x/100] where x represents the value in centimetres.

Any values ≤ 1 metre were then excluded from the analyses.

- **Body mass index (BMI)**

BMI data were available from the primary care EHR for the following time periods:

1. Earliest recorded BMI
2. BMI recorded in a ± 6 month window around the time of diabetes diagnosis
3. Latest recorded BMI

However, given that many weight measurements in the EHR are not paired to a height measurement taken at the same time, a new adult BMI value was calculated for each volunteer using latest adult weight (if weight taken in last 5 years) and median adult height (as above). For volunteers with Type 2 diabetes (based on presence of a Read code for Type 2 diabetes), a new value was calculated for ‘BMI in a ± 6 month window around the time of diabetes diagnosis’ using the value for weight recorded in a ± 6 month window around the time of diabetes diagnosis and median adult height (as above). For volunteers with Type 2 diabetes, a new value was also calculated for ‘BMI in past 2 years’ using latest adult weight (if weight taken in last 5 years) and median adult height (as above).

Any recalculated BMI values which were >75kg/m^2^ or <12kg/m^2^ were then excluded from the analysis.

- **HbA1c**

HbA1c data were available from the primary care EHR for the following time periods:

1. Earliest recorded HbA1c
2. HbA1c recorded in a ± 6 month window around the time of diabetes diagnosis
3. Latest recorded HbA1c

The majority of laboratory data enter the primary care EHR via a direct electronic link from the laboratory and data are therefore rarely entered manually in primary care. In the small number of instances where laboratory results are manually entered in primary care (e.g. a GP transcribing a result from a letter into the primary care EHR), there is potential for human error, therefore the following data quality steps were performed for each volunteer, in accordance with MASTERMIND study (3):

In cases where the HbA1c value was between 3.9 and 20 (inclusive) and the unit of measurement was either ‘mmol/mol’ or empty (blank), the following assumptions were made:

1. The data was manually entered
2. The unit ‘mmol/mol’ was entered in error
3. The intention had been to enter ‘percent’ as the unit of measurement

If these criteria were met, the unit was changed to percent, leaving the value intact.

In cases where the HbA1c value was between 20 and 195 (inclusive) and the unit of measurement was either ‘percent’ or empty (blank), the following assumptions were made:

1. The data was manually entered
2. The unit ‘percent’ was entered in error
3. The intention had been to enter ‘mmol/mol’ as the unit of measurement

If these criteria were met, the unit was changed to mmol/mol, leaving the value intact.

Once the above data quality steps were performed, all values where unit of measurement was ‘percent’ were then converted to the equivalent ‘mmol/mol’ value, using the conversion calculation: [(x – 2.152) / 0.09148] where x represents the HbA1c value in percent.

All values > 200 mmol/mol were then removed from the analysis.

- **Co-morbidities**

For volunteers with a Read code for Type 2 diabetes, the following diagnoses (available in the primary care EHR dataset) were included as co-morbidities:

1. Proliferative retinopathy (i.e. sight-threatening)
2. Maculopathy (i.e. sight-threatening)
3. Erectile dysfunction
4. Neuropathy
5. Peripheral vascular disease
6. Atrial fibrillation/flutter
7. Ischaemic heart disease
8. Heart failure
9. Stroke
10. CKD (stage 3 or worse)
11. Hypertension

The proportion of volunteers with a Read Code for Type 2 diabetes and a Read code for each of these diagnoses was calculated, as well as the proportion of volunteers with a Read Code for Type 2 diabetes and more than 1 co-morbidity.
