## Supplementary File 4 for "Cohort Profile: East London Genes & Health (ELGH), a community based population genomics and health study of British-Bangladeshi and British-Pakistani people"

**Supplementary file 4: SQL code for primary care electronic health record data extraction**

1. Demographics and Diagnoses

```
USE CEG

GO
 -- DROP all temp tables if exist
 BEGIN -- drop all temp tables if exist

	DECLARE @DropGlobal bit=0 --Default dont drop global temp table

	DECLARE @DROP_STATEMENT nvarchar(1000)
	DECLARE cursorDEL CURSOR FOR
	SELECT 'DROP TABLE '
	   + case
			   when name like '##%' then name
			   when name like '#%' then SUBSTRING(name, 1, CHARINDEX( '____', name)-1)
		end as DropSQL
	from tempdb..sysobjects
	WHERE name LIKE '#%'
	   AND OBJECT_ID('tempdb..' + name) IS NOT NULL
	   AND name not like case
						   when @DropGlobal=0 then '##%' --//Exclude global temp
						   else '#######%'    --//some fack expression so we can
											   --//select global temp for delete
						end
	OPEN cursorDEL
	FETCH NEXT FROM cursorDEL INTO @DROP_STATEMENT
	WHILE @@FETCH_STATUS = 0
	BEGIN
	EXEC (@DROP_STATEMENT)
	--print @DROP_STATEMENT
	FETCH NEXT FROM cursorDEL INTO @DROP_STATEMENT
	END
	CLOSE cursorDEL
	DEALLOCATE cursorDEL

END

GO

------------------  Demographics -----------------------------------------------------------------------

SELECT
	 [SK_ServiceProviderID]
	,[SK_PatientID]
	,[Gender]
	,[YearOfDeath]
	,[DateRegistered]
	,[DateRegisteredEnd]
	,[EthnicityCode]
	,[LSOACode]
	,(SELECT TOP 1 IMD.Score FROM FactEngine.[shared].[IMDScoreByLSOA] AS IMD WHERE IMD.LSOA = Dup.LSOACode AND IMD.ReferenceYear = 2010) AS [IMD 2010 Score]


INTO #Demographics

FROM
(

	SELECT
		 [SK_ServiceProviderID]
		,[SK_PatientID]
		,[Gender]
		,[DateRegistered]
		,[DateRegisteredEnd]
		,[YearOfDeath]
		,[EthnicityCode]
		,[LSOACode]
		,row_number() over(partition by [SK_PatientID] order by [DateRegistered] DESC) as rn

	FROM [08V].[PatientDemographics] P
	WHERE SK_PatientID IN(SELECT SK_PatientID FROM ceg.GenomicsData WHERE CCG = '08V')

)Dup

WHERE rn = 1

GO


------------------ Country of birth ----------------------------------------------------------------------------------------

SELECT
	 [SK_PatientID]
	,[ClinicalCode] CountryOfBirthCode
	,(SELECT TOP 1 Term FROM Dictionary.[dbo].[ReadCodes] R WHERE R.ReadCode COLLATE Latin1_General_CS_AS = Country.[ClinicalCode]) AS CountryOfBirthTerm

INTO  #CountryOfBirth

FROM
(
	SELECT

		 [SK_PatientID]
		,[ClinicalCode]
		,[EventDate]
		,row_number() over(partition by [SK_PatientID] order by [EventDate] DESC) as rn

	FROM  [08V].[GPEncounter]
	WHERE
		(
			[ClinicalCode] COLLATE Latin1_General_Bin LIKE '13[d-h]%'		-- COLLATE Latin1_General_Bin
			OR
			[ClinicalCode] COLLATE Latin1_General_Bin LIKE '13[j-k]%'		-- COLLATE Latin1_General_Bin
			OR
			[ClinicalCode] COLLATE Latin1_General_CS_AS = '13Zq'
		)
		AND
		SK_PatientID IN(SELECT SK_PatientID FROM ceg.GenomicsData WHERE CCG = '08V')

)as Country

WHERE rn = 1

GO

------------- Ethnicity-------------------------------------------------------------------------------------------------------------------


SELECT
	 [SK_PatientID]
	,[ClinicalCode] EthnicityCode
	,[EventDate] Ethnicity_DateRecorded

INTO  #Ethnicity

FROM
(
	SELECT
		 [SK_PatientID]
		,[ClinicalCode]
		,[EventDate]
		,row_number() over(partition by [SK_PatientID] order by [EventDate] DESC) as rn

	FROM  [08V].[GPEncounter]
	WHERE
		(
			[ClinicalCode] COLLATE Latin1_General_CS_AS LIKE '9S%'
			OR
			[ClinicalCode] COLLATE Latin1_General_CS_AS LIKE '9i%'
		)
		AND
		SK_PatientID IN(SELECT SK_PatientID FROM ceg.GenomicsData WHERE CCG = '08V')

)as Eth

WHERE rn = 1

GO

------- FAMILY HISTORY ------------------------------------------------------------------------------------------------
------------------  Family History of Diabetes ------------------------------------------------------------------------


SELECT
	 [SK_PatientID]
	,[ClinicalCode] FH_DiabetesCode
	,[EventDate] FH_DiabetesDate


INTO  #FH_Diabetes

FROM
(
	SELECT
		 [SK_PatientID]
		,[ClinicalCode]
		,[EventDate]
		,row_number() over(partition by [SK_PatientID] order by [EventDate] DESC) as rn

	FROM  [08V].[GPEncounter]
	WHERE
		[ClinicalCode] COLLATE Latin1_General_CS_AS = '1253'
		AND
		SK_PatientID IN(SELECT SK_PatientID FROM ceg.GenomicsData WHERE CCG = '08V')

)as FH_IHD

WHERE rn = 1

GO


------------------  Family History of Ischaemic heart disease (IHD) ------------------------------------------------------------------------


SELECT
	 [SK_PatientID]
	,[ClinicalCode] FH_IHDCode
	,[EventDate] FH_IHDDate


INTO  #FH_IHD

FROM
(
	SELECT
		 [SK_PatientID]
		,[ClinicalCode]
		,[EventDate]
		,row_number() over(partition by [SK_PatientID] order by [EventDate] DESC) as rn

	FROM  [08V].[GPEncounter]
	WHERE
		[ClinicalCode] COLLATE Latin1_General_CS_AS LIKE '12C2%'	--% added for Synonym
		AND
		SK_PatientID IN(SELECT SK_PatientID FROM ceg.GenomicsData WHERE CCG = '08V')

)as FH_IHD

WHERE rn = 1

GO

-- DIABETES DIAGNOSES --------------------------------------------------------------------------------------------------
------------------ Type 1 Diabetes QOF earliest ever -----------------------------------------------------------------------


SELECT
	 [SK_PatientID]
	,[ClinicalCode] DiabetesT1Code
	,[EventDate]	DiabetesT1DateRecorded

INTO  #DiabetesT1QoF

FROM
(
	SELECT
		*

	FROM
	(
		SELECT
			 [SK_PatientID]
			,[ClinicalCode]
			,[EventDate]
			,row_number() over(partition by [SK_PatientID] order by [EventDate] ASC) as rn

		FROM
			[08V].[GPEncounter]

		WHERE
			(
				[ClinicalCode] COLLATE Latin1_General_CS_AS LIKE 'C10E%'		--C10E% Type 1 diabetes mellitus
				OR
				[ClinicalCode] COLLATE Latin1_General_CS_AS LIKE 'C10ER%'	-- % for Synonym			--C10ER Latent autoimmune diabetes mellitus in adult
			)
			AND
			SK_PatientID IN(SELECT SK_PatientID FROM ceg.GenomicsData WHERE CCG = '08V')

	)as T1

	WHERE
		rn = 1

) AS DiabT1

WHERE
	SK_PatientID NOT IN	-- QOF definition for Diabetes excludes diabetes resolved codes if appearing after the latest diagnostic code
	(
		SELECT
			[SK_PatientID]

		FROM
		(
			SELECT
				 [SK_PatientID]
				,[ClinicalCode]
				,[EventDate]
				,row_number() over(partition by [SK_PatientID] order by [EventDate] DESC) as rn

			FROM
				[08V].[GPEncounter]

			WHERE
				(
					[ClinicalCode] COLLATE Latin1_General_CS_AS LIKE 'C10E%'		--C10E% Type 1 diabetes mellitus
					OR
					[ClinicalCode] COLLATE Latin1_General_CS_AS LIKE 'C10ER%'	-- % for Synonym 		--C10ER Latent autoimmune diabetes mellitus in adult
					OR
					[ClinicalCode] COLLATE Latin1_General_CS_AS IN('21263', '212H')  --Diabetes resolved
				)
				AND
				SK_PatientID IN(SELECT SK_PatientID FROM ceg.GenomicsData WHERE CCG = '08V')

		) AS  Res

		WHERE
			rn = 1	AND	[ClinicalCode] IN('21263', '212H')	--Diabetes resolved
	)

GO


------------------ Type 2 Diabetes QOF earliest ever -----------------------------------------------------------------------

/*
C10F% Type 2 diabetes mellitus (exclude C10F8 Reaven's syndrome),
C109J Insulin treated Type 2 diabetes mellitus,
*/


SELECT
	 [SK_PatientID]
	,[ClinicalCode] DiabetesT2Code
	,[EventDate]	DiabetesT2DateRecorded

INTO  #DiabetesT2QoF

FROM
(
	SELECT
		*

	FROM
	(
		SELECT
			 [SK_PatientID]
			,[ClinicalCode]
			,[EventDate]
			,row_number() over(partition by [SK_PatientID] order by [EventDate] ASC) as rn

		FROM
			[08V].[GPEncounter]

		WHERE
			(
				(
					[ClinicalCode] COLLATE Latin1_General_CS_AS LIKE 'C10F%' --C10F% Type 2 diabetes mellitus
					AND
					[ClinicalCode] COLLATE Latin1_General_CS_AS <> 'C10F8' --C10F8 Reaven's syndrome
				)
				OR
				[ClinicalCode] COLLATE Latin1_General_CS_AS LIKE 'C109J%' -- % for Synonym	 --C109J Insulin treated Type 2 diabetes mellitus
			)
			AND
			SK_PatientID IN(SELECT SK_PatientID FROM ceg.GenomicsData WHERE CCG = '08V')

	)as T2

	WHERE
		rn = 1

) AS DiabT2

WHERE
	SK_PatientID NOT IN	-- QOF definition for Diabetes excludes diabetes resolved codes if appearing after the latest diagnostic code
	(
		SELECT
			[SK_PatientID]

		FROM
		(
			SELECT
				 [SK_PatientID]
				,[ClinicalCode]
				,[EventDate]
				,row_number() over(partition by [SK_PatientID] order by [EventDate] DESC) as rn

			FROM
				[08V].[GPEncounter]

			WHERE
				(
					(
						[ClinicalCode] COLLATE Latin1_General_CS_AS LIKE 'C10F%'		--C10F% Type 2 diabetes mellitus
						AND
						[ClinicalCode] COLLATE Latin1_General_CS_AS <> 'C10F8'				--C10F8 Reaven's syndrome
					)
					OR
					[ClinicalCode] COLLATE Latin1_General_CS_AS LIKE 'C109J%' -- % for Synonym			--C109J Insulin treated Type 2 diabetes mellitus
					OR
					[ClinicalCode] COLLATE Latin1_General_CS_AS IN('21263', '212H')  --Diabetes resolved
				)
				AND
				SK_PatientID IN(SELECT SK_PatientID FROM ceg.GenomicsData WHERE CCG = '08V')

		) AS  Res

		WHERE
			rn = 1	AND	[ClinicalCode] IN('21263', '212H')	--Diabetes resolved
	)

GO

------------------ Secondary Diabetes earliest ever -----------------------------------------------------------------------
/*
C10G% Secondary pancreatic diabetes mellitus,
C10H% Diabetes mellitus induced by non-steroid drugs,
C10N% Secondary diabetes mellitus,

C10B Diabetes mellitus induced by steroids,
*/


SELECT
	 [SK_PatientID]
	,[ClinicalCode] SecondaryDiabetesCode
	,[EventDate] SecondaryDiabetesDateRecorded

INTO  #SecondaryDiabetesQoF

FROM
(
	SELECT
		*

	FROM
	(
		SELECT
			 [SK_PatientID]
			,[ClinicalCode]
			,[EventDate]
			,row_number() over(partition by [SK_PatientID] order by [EventDate] ASC) as rn

		FROM
			[08V].[GPEncounter]

		WHERE
			(
					[ClinicalCode] COLLATE Latin1_General_Bin LIKE 'C10[G-H]%'	 --C10G% Secondary pancreatic diabetes mellitus		C10H% Diabetes mellitus induced by non-steroid drugs
				OR	[ClinicalCode] COLLATE Latin1_General_CS_AS LIKE 'C10N%'	 --C10N% Secondary diabetes mellitus
				OR	[ClinicalCode] COLLATE Latin1_General_CS_AS = 'C10B'		 --Diabetes mellitus induced by steroids
			)
			AND
			SK_PatientID IN(SELECT SK_PatientID FROM ceg.GenomicsData WHERE CCG = '08V')

	)as SecdaryDiab

	WHERE
		rn = 1

) AS SecondaryDiabetes

WHERE
	SK_PatientID NOT IN	-- QOF definition for Diabetes excludes diabetes resolved codes if appearing after the latest diagnostic code
	(
		SELECT
			[SK_PatientID]

		FROM
		(
			SELECT
				 [SK_PatientID]
				,[ClinicalCode]
				,[EventDate]
				,row_number() over(partition by [SK_PatientID] order by [EventDate] DESC) as rn

			FROM
				[08V].[GPEncounter]

			WHERE
				(						-- COLLATE Latin1_General_Bin
						[ClinicalCode] COLLATE Latin1_General_CS_AS LIKE 'C10[G-H]%'	--C10G% Secondary pancreatic diabetes mellitus		C10H% Diabetes mellitus induced by non-steroid drugs
					OR	[ClinicalCode] COLLATE Latin1_General_CS_AS LIKE 'C10N%'		 --C10N% Secondary diabetes mellitus
					OR	[ClinicalCode] COLLATE Latin1_General_CS_AS = 'C10B'			 --Diabetes mellitus induced by steroids
					OR	[ClinicalCode] COLLATE Latin1_General_CS_AS IN('21263', '212H')  --Diabetes resolved
				)
				AND
				SK_PatientID IN(SELECT SK_PatientID FROM ceg.GenomicsData WHERE CCG = '08V')

		) AS  Res

		WHERE
			rn = 1	AND	[ClinicalCode] IN('21263', '212H')	--Diabetes resolved
	)

GO


------------------ Other types of diabetes earliest ever -----------------------------------------------------------------------
/*
C10A, C10C, C10D, C1A%, C10M%, C326%, A3A2, C135, PKyP, Q441, C10FS, C150%, C3500, C10N1, C370%, PKyF, C1zy4, PH3yA, PKy93, PKy1
*/


SELECT
	 [SK_PatientID]
	,[ClinicalCode] OtherTypesOfDiabetesCode
	,[EventDate] OtherTypesOfDiabetesDateRecorded

INTO  #OtherTypesOfDiabetes

FROM
(
	SELECT

		 [SK_PatientID]
		,[ClinicalCode]
		,[EventDate]
		,row_number() over(partition by [SK_PatientID] order by [EventDate] ASC) as rn

	FROM  [08V].[GPEncounter]
	WHERE
		(

				[ClinicalCode] COLLATE Latin1_General_CS_AS LIKE 'C10M%'	--C10M% Lipoatrophic diabetes mellitus
			OR	[ClinicalCode] COLLATE Latin1_General_CS_AS LIKE 'C150%'	--C150% Cushing's syndrome
			OR  [ClinicalCode] COLLATE Latin1_General_CS_AS LIKE 'C1A%'		--C1A% Insulin resistance
			OR	[ClinicalCode] COLLATE Latin1_General_CS_AS LIKE 'C326%'	--C10N% Lipodystrophy
			OR	[ClinicalCode] COLLATE Latin1_General_CS_AS LIKE 'C370%'	--C370%	Cystic fibrosis
			OR	[ClinicalCode] COLLATE Latin1_General_CS_AS IN
				(
					 'A3A2', 'A3A2-1'			-- Synonym	--Whipple's disease
					,'C10A', 'C10A-1'			-- Synonym	--Malnutrition-related diabetes mellitus
					,'C10C', 'C10C-1', 'C10C-2' -- Synonym		--Diabetes mellitus autosomal dominant
					,'C10D', 'C10D-1'			-- Synonym	--Diabetes mellitus autosomal dominant type 2
					,'C10FS'					--Maternally inherited diabetes mellitus
					,'C10N1'					--Cystic fibrosis related diabetes mellitus
					,'C135', 'C135-1', 'C135-2'	-- Synonym	--Diabetes insipidus
					,'C1zy4'					--Werner's syndrome
					,'C3500', 'C3500-1', 'C3500-2', 'C3500-3' -- Synonym	--Haemochromatosis
					,'PH3yA'					--Bloom syndrome
					,'PKy1', 'PKy1-1'			-- Synonym		--Laurence-Moon-Biedl syndrome
					,'PKy93'					--Prader - Willi syndrome
					,'PKyF'						--Alstrom syndrome
					,'PKyP', 'PKyP-1'			-- Synonym		--Diabetes insipidus, diabetes mellitus, optic atrophy and deafness
					,'Q441'						--Neonatal diabetes mellitus
				)
		)
		AND
		SK_PatientID IN(SELECT SK_PatientID FROM ceg.GenomicsData WHERE CCG = '08V')

)as OtherTypesOfDiab

WHERE rn = 1

GO


------------------ Pancreatic disease/surgery earliest ever -----------------------------------------------------------------------
/*
J670%, J671%, J6710, J6711, J67y6, 9b8G
*/


SELECT
	 [SK_PatientID]
	,[ClinicalCode] PancreaticDiseaseCode
	,[EventDate] PancreaticDiseaseDateRecorded

INTO  #PancreaticDisease

FROM
(
	SELECT

		 [SK_PatientID]
		,[ClinicalCode]
		,[EventDate]
		,row_number() over(partition by [SK_PatientID] order by [EventDate] ASC) as rn

	FROM  [08V].[GPEncounter]
	WHERE
		(
				[ClinicalCode] COLLATE Latin1_General_CS_AS LIKE 'J670%'	--Acute pancreatitis
			OR	[ClinicalCode] COLLATE Latin1_General_CS_AS LIKE 'J671%'	--Chronic pancreatitis
			OR	[ClinicalCode] COLLATE Latin1_General_CS_AS IN
				(
					 'J6710'		--Alcohol-induced chronic pancreatitis
					,'J6711'		--Gallstone chronic pancreatitis
					,'J67y6'		--Exocrine pancreatic insufficiency
					,'9b8G'			--Hepatobiliary and pancreatic surgery
				)
		)
		AND
		SK_PatientID IN(SELECT SK_PatientID FROM ceg.GenomicsData WHERE CCG = '08V')

)as PancreaticDis

WHERE rn = 1

GO


------------------ Gestational diabetes mellitus (Diabetes in pregnancy) earliest ever -----------------------------------------------------------------------

SELECT
	 [SK_PatientID]
	,[ClinicalCode] GestationalDiabetesCode
	,[EventDate]	GestationalDiabetesDateRecorded

INTO  #GestationalDiabetes

FROM
(
	SELECT

		 [SK_PatientID]
		,[ClinicalCode]
		,[EventDate]
		,row_number() over(partition by [SK_PatientID] order by [EventDate] ASC) as rn

	FROM  [08V].[GPEncounter]
	WHERE
		(
			[ClinicalCode] COLLATE Latin1_General_CS_AS LIKE 'L1808%'	--Synonym	--L1808 Diabetes mellitus arising in pregnancy
			OR
			[ClinicalCode] COLLATE Latin1_General_CS_AS LIKE 'L1809%'	--Synonym	--L1809 Gestational diabetes mellitus
		)
		AND
		SK_PatientID IN(SELECT SK_PatientID FROM ceg.GenomicsData WHERE CCG = '08V')

)as GestDiab

WHERE rn = 1

GO


------------------ Diabetes emergencies latest ever -----------------------------------------------------------------------
/*
C10EM, C109K
*/

SELECT
	 [SK_PatientID]
	,[ClinicalCode] EmergenciesDiabetesCode
	,[EventDate]	EmergenciesDiabetesDateRecorded

INTO  #EmergenciesDiabetes

FROM
(
	SELECT

		 [SK_PatientID]
		,[ClinicalCode]
		,[EventDate]
		,row_number() over(partition by [SK_PatientID] order by [EventDate] DESC) as rn

	FROM  [08V].[GPEncounter]
	WHERE
		(
			[ClinicalCode] COLLATE Latin1_General_CS_AS LIKE 'C10EM%'	--Synonym	--Type 1 diabetes mellitus with ketoacidosis
			OR
			[ClinicalCode] COLLATE Latin1_General_CS_AS LIKE 'C109K%'	--Synonym	--Hyperosmolar non-ketotic state in type 2 diabetes mellitus
		)
		AND
		SK_PatientID IN(SELECT SK_PatientID FROM ceg.GenomicsData WHERE CCG = '08V')

)as EmergenciesDia

WHERE rn = 1

GO


------------------ Bariatric surgery earliest ever -----------------------------------------------------------------------
/*
76110, 76111, 76112, 76113, 76114, 76115, 76116, 76130, 76131, 76132, 76133, 76134, 76135, 76136, 76140, 76141, 76160, 76166
*/


SELECT
	 [SK_PatientID]
	,[ClinicalCode] BariatricSurgeryCode
	,[EventDate] BariatricSurgeryDateRecorded

INTO  #BariatricSurgery

FROM
(
	SELECT

		 [SK_PatientID]
		,[ClinicalCode]
		,[EventDate]
		,row_number() over(partition by [SK_PatientID] order by [EventDate] ASC) as rn

	FROM  [08V].[GPEncounter]
	WHERE
		(
			[ClinicalCode] COLLATE Latin1_General_CS_AS LIKE '7611[0-6]%'
			OR
			[ClinicalCode] COLLATE Latin1_General_CS_AS LIKE '7613[0-6]%'
			OR
			[ClinicalCode] COLLATE Latin1_General_CS_AS LIKE '76160%'	-- Synonym		--Bypass of stomach by anastomosis of stomach to jejunum NEC
			OR
			[ClinicalCode] COLLATE Latin1_General_CS_AS IN
			(
				 '76140'				--Bypass of stomach by anastomosis of oesophagus to duodenum
				,'76141','76141-1'		--Bypass of stomach by anastomosis of stomach to duodenum
				,'76166'				--Laparoscopic gastric bypass
			)
		)
		AND
		SK_PatientID IN(SELECT SK_PatientID FROM ceg.GenomicsData WHERE CCG = '08V')

)as BariatricSurg

WHERE rn = 1

GO

-- DIABETES RISK SCORES ---------------------------------------------------------------------------------------------------------------------------------
------------------ Diabetes risk 1: Prediabetes code OR glucose intolerance codes -----------------------------------------------------------------------
---------------------- Earliest --------------------------------------------------------------------------------------------------------------


SELECT
	 [SK_PatientID]
	,[ClinicalCode] PrediabetesRisk1EarliestCode
	,[EventDate]	PrediabetesRisk1EarliestDateRecorded

INTO  #Risk1PrediabetesEarliest

FROM
(
	SELECT
		*

	FROM
	(
		SELECT
			 [SK_PatientID]
			,[ClinicalCode]
			,[EventDate]
			,row_number() over(partition by [SK_PatientID] order by [EventDate] ASC) as rn

		FROM
			[08V].[GPEncounter]

		WHERE
			(
				[ClinicalCode] COLLATE Latin1_General_CS_AS IN('C11y2', 'C11y3', 'C11y5')
				OR
				[ClinicalCode] COLLATE Latin1_General_CS_AS LIKE 'R10E%'	-- For synonyms terms
				OR
				[ClinicalCode] COLLATE Latin1_General_CS_AS LIKE 'R10D0%'
				OR
				[ClinicalCode] COLLATE Latin1_General_CS_AS LIKE 'R102%'
			)
			AND
			SK_PatientID IN(SELECT SK_PatientID FROM ceg.GenomicsData WHERE CCG = '08V')

	)as R1PrediabEarliest

	WHERE
		rn = 1

) AS Risk1PrediabetesEarliest

WHERE
	SK_PatientID NOT IN	-- Exclude if there is a more recent diabetes diagnostic codes
	(
		SELECT
			[SK_PatientID]

		FROM
		(
			SELECT
				 [SK_PatientID]
				,[ClinicalCode]
				,[EventDate]
				,row_number() over(partition by [SK_PatientID] order by [EventDate] DESC) as rn

			FROM
				[08V].[GPEncounter]

			WHERE
				(
					[ClinicalCode] COLLATE Latin1_General_CS_AS IN('C11y2', 'C11y3', 'C11y5')
					OR
					[ClinicalCode] COLLATE Latin1_General_CS_AS LIKE 'R10E%'	-- For synonyms terms
					OR
					[ClinicalCode] COLLATE Latin1_General_CS_AS LIKE 'R10D0%'
					OR
					[ClinicalCode] COLLATE Latin1_General_CS_AS LIKE 'R102%'
					OR
					[ClinicalCode] COLLATE Latin1_General_CS_AS IN('21263', '212H')  --Diabetes resolved
				)
				AND
				SK_PatientID IN(SELECT SK_PatientID FROM ceg.GenomicsData WHERE CCG = '08V')

		) AS  Res

		WHERE
			rn = 1	AND	[ClinicalCode] IN('21263', '212H')	--Diabetes resolved
	)

GO


---------------------- Latest --------------------------------------------------------------------------------------------------------------


SELECT
	 [SK_PatientID]
	,[ClinicalCode] PrediabetesRisk1LatestCode
	,[EventDate] PrediabetesRisk1LatestDateRecorded

INTO  #Risk1PrediabetesLatest

FROM
(
	SELECT

		 [SK_PatientID]
		,[ClinicalCode]
		,[EventDate]
		,row_number() over(partition by [SK_PatientID] order by [EventDate] DESC) as rn

	FROM  [08V].[GPEncounter]
	WHERE
		(
			[ClinicalCode] COLLATE Latin1_General_CS_AS IN('C11y2', 'C11y3', 'C11y5')
			OR
			[ClinicalCode] COLLATE Latin1_General_CS_AS LIKE 'R10E%'	-- For synonyms terms
			OR
			[ClinicalCode] COLLATE Latin1_General_CS_AS LIKE 'R10D0%'
			OR
			[ClinicalCode] COLLATE Latin1_General_CS_AS LIKE 'R102%'
			OR
			[ClinicalCode] COLLATE Latin1_General_CS_AS IN('21263', '212H')  --Diabetes resolved
		)
		AND
		SK_PatientID IN(SELECT SK_PatientID FROM ceg.GenomicsData WHERE CCG = '08V')

)as Risk1PrediabetesLatest

WHERE rn = 1 AND [ClinicalCode] NOT IN('21263', '212H')	---- Exclude if there is a more recent diabetes diagnostic codes

GO


------------------ Diabetes risk 2:  At risk of diabetes mellitus ------------------------------------------------------------------------------------------------
---------------------- Earliest --------------------------------------------------------------------------------------------------------------


SELECT
	 [SK_PatientID]
	,[ClinicalCode] Risk2MellitusEarliestCode
	,[EventDate]	Risk2MellitusEarliestDateRecorded

INTO  #Risk2MellitusEarliest

FROM
(
	SELECT
		*

	FROM
	(
		SELECT
			 [SK_PatientID]
			,[ClinicalCode]
			,[EventDate]
			,row_number() over(partition by [SK_PatientID] order by [EventDate] ASC) as rn

		FROM
			[08V].[GPEncounter]

		WHERE
			[ClinicalCode] COLLATE Latin1_General_CS_AS IN('14O8', '14O80')
			AND
			SK_PatientID IN(SELECT SK_PatientID FROM ceg.GenomicsData WHERE CCG = '08V')

	)as R2MellitusEarliest

	WHERE
		rn = 1

) AS Risk2MellitusEarliest

WHERE
	SK_PatientID NOT IN	-- Exclude if there is a more recent diabetes diagnostic codes
	(
		SELECT
			[SK_PatientID]

		FROM
		(
			SELECT
				 [SK_PatientID]
				,[ClinicalCode]
				,[EventDate]
				,row_number() over(partition by [SK_PatientID] order by [EventDate] DESC) as rn

			FROM
				[08V].[GPEncounter]

			WHERE
				(
					[ClinicalCode] COLLATE Latin1_General_CS_AS IN('14O8', '14O80')
					OR
					[ClinicalCode] COLLATE Latin1_General_CS_AS IN('21263', '212H')  --Diabetes resolved
				)
				AND
				SK_PatientID IN(SELECT SK_PatientID FROM ceg.GenomicsData WHERE CCG = '08V')

		) AS  Res

		WHERE
			rn = 1	AND	[ClinicalCode] IN('21263', '212H')	--Diabetes resolved
	)

GO


---------------------- Latest --------------------------------------------------------------------------------------------------------------


SELECT
	 [SK_PatientID]
	,[ClinicalCode] Risk2MellitusLatestCode
	,[EventDate] Risk2MellitusLatestDateRecorded

INTO  #Risk2MellitusLatest

FROM
(
	SELECT

		 [SK_PatientID]
		,[ClinicalCode]
		,[EventDate]
		,row_number() over(partition by [SK_PatientID] order by [EventDate] DESC) as rn

	FROM  [08V].[GPEncounter]
	WHERE
		(
			[ClinicalCode] COLLATE Latin1_General_CS_AS IN('14O8', '14O80')
			OR
			[ClinicalCode] COLLATE Latin1_General_CS_AS IN('21263', '212H')  --Diabetes resolved
		)
		AND
		SK_PatientID IN(SELECT SK_PatientID FROM ceg.GenomicsData WHERE CCG = '08V')

)as Risk2MellitusLatest

WHERE rn = 1 AND [ClinicalCode] NOT IN('21263', '212H')	-- Exclude if there is a more recent diabetes diagnostic codes

GO


------------------ Diabetes risk 3:  HbA1c --------------------------------------------------------------------------------------------------------------------
---------------------- Earliest --------------------------------------------------------------------------------------------------------------

SELECT
	 [SK_PatientID]
	,[ClinicalCode] Risk3EarliestHbA1cCode
	,[EventDate]	Risk3EarliestHbA1cDateRecorded
	,[Value]		Risk3EarliestHbA1cValue
	,[Units]		Risk3EarliestHbA1cUnits

INTO  #Risk3EarliestHbA1c

FROM
(
	SELECT
		*

	FROM
	(
		SELECT
			 [SK_PatientID]
			,[ClinicalCode]
			,[EventDate]
			,[Value]
			,[Units]
			,row_number() over(partition by [SK_PatientID] order by [EventDate] ASC) as rn

		FROM
			[08V].[GPEncounter]

		WHERE
			(
				([ClinicalCode] COLLATE Latin1_General_CS_AS = '42W4' AND [Value] BETWEEN 6 AND 6.4)	--[Units] = '%'
				OR
				([ClinicalCode] COLLATE Latin1_General_CS_AS = '42W5' AND [Value] BETWEEN 42 AND 47.99) --[Units] = 'mmol/mol'
			)
			AND
			SK_PatientID IN(SELECT SK_PatientID FROM ceg.GenomicsData WHERE CCG = '08V')

	)as R3HbA1c

	WHERE
		rn = 1

) AS Risk3EarliestHbA1c

WHERE
	SK_PatientID NOT IN	-- Exclude if there is a more recent diabetes diagnostic codes
	(
		SELECT
			[SK_PatientID]

		FROM
		(
			SELECT
				 [SK_PatientID]
				,[ClinicalCode]
				,[EventDate]
				,row_number() over(partition by [SK_PatientID] order by [EventDate] DESC) as rn

			FROM
				[08V].[GPEncounter]

			WHERE
				(
					([ClinicalCode] COLLATE Latin1_General_CS_AS = '42W4' AND [Value] BETWEEN 6 AND 6.4)	--[Units] = '%'
					OR
					([ClinicalCode] COLLATE Latin1_General_CS_AS = '42W5' AND [Value] BETWEEN 42 AND 47.99) --[Units] = 'mmol/mol'
					OR
					[ClinicalCode] COLLATE Latin1_General_CS_AS IN('21263', '212H')  --Diabetes resolved
				)
				AND
				SK_PatientID IN(SELECT SK_PatientID FROM ceg.GenomicsData WHERE CCG = '08V')

		) AS  Res

		WHERE
			rn = 1	AND	[ClinicalCode] IN('21263', '212H')	-- Exclude if there is a more recent diabetes diagnostic codes
	)

GO


---------------------- Latest --------------------------------------------------------------------------------------------------------------

SELECT
	 [SK_PatientID]
	,[ClinicalCode] Risk3LatestHbA1cCode
	,[EventDate]	Risk3LatestHbA1cDateRecorded
	,[Value]		Risk3LatestHbA1cValue
	,[Units]		Risk3LatestHbA1cUnits

INTO  #Risk3LatestHbA1c

FROM
(
	SELECT
		 [SK_PatientID]
		,[ClinicalCode]
		,[EventDate]
		,[Value]
		,[Units]
		,row_number() over(partition by [SK_PatientID] order by [EventDate] DESC) as rn

	FROM
		[08V].[GPEncounter]

	WHERE
		(
			([ClinicalCode] COLLATE Latin1_General_CS_AS = '42W4' AND [Value] BETWEEN 6 AND 6.4)	--[Units] = '%'
			OR
			([ClinicalCode] COLLATE Latin1_General_CS_AS = '42W5' AND [Value] BETWEEN 42 AND 47.99) --[Units] = 'mmol/mol'
			OR
			[ClinicalCode] COLLATE Latin1_General_CS_AS IN('21263', '212H')  --Diabetes resolved
		)
		AND
		SK_PatientID IN(SELECT SK_PatientID FROM ceg.GenomicsData WHERE CCG = '08V')

) AS Risk3LatestHbA1c

WHERE rn = 1 AND [ClinicalCode] NOT IN('21263', '212H')	-- Exclude if there is a more recent diabetes diagnostic codes

GO

------------------ Diabetes risk 4: Qdiabetes score -----------------------------------------------------------------------------------------------------------------
---------------------- Earliest --------------------------------------------------------------------------------------------------------------


SELECT
	 [SK_PatientID]
	,[ClinicalCode] Risk4QdiabetesScoreEarliestCode
	,[EventDate]	Risk4QdiabetesScoreEarliestDateRecorded
	,[Value]		QdiabetesEarliestScore
	,[Units]		QdiabetesEarliestUnits

INTO  #Risk4QdiabetesScoreEarliest

FROM
(
	SELECT
		*

	FROM
	(
		SELECT
			 [SK_PatientID]
			,[ClinicalCode]
			,[EventDate]
			,[Value]
			,[Units]
			,row_number() over(partition by [SK_PatientID] order by [EventDate] ASC) as rn

		FROM
			[08V].[GPEncounter]

		WHERE
			[ClinicalCode] COLLATE Latin1_General_CS_AS = '38Gj'
			AND
			SK_PatientID IN(SELECT SK_PatientID FROM ceg.GenomicsData WHERE CCG = '08V')

	)as Risk4QdiaScoreEarliest

	WHERE
		rn = 1

) AS Risk4QdiabetesScoreEarliest

WHERE
	SK_PatientID NOT IN	-- Exclude if there is a more recent diabetes diagnostic codes
	(
		SELECT
			[SK_PatientID]

		FROM
		(
			SELECT
				 [SK_PatientID]
				,[ClinicalCode]
				,[EventDate]
				,row_number() over(partition by [SK_PatientID] order by [EventDate] DESC) as rn

			FROM
				[08V].[GPEncounter]

			WHERE
				(
					[ClinicalCode] COLLATE Latin1_General_CS_AS = '38Gj'
					OR
					[ClinicalCode] COLLATE Latin1_General_CS_AS IN('21263', '212H')  --Diabetes resolved
				)
				AND
				SK_PatientID IN(SELECT SK_PatientID FROM ceg.GenomicsData WHERE CCG = '08V')

		) AS  Res

		WHERE
			rn = 1	AND	[ClinicalCode] IN('21263', '212H')	-- Exclude if there is a more recent diabetes diagnostic codes
	)

GO

---------------------- Latest --------------------------------------------------------------------------------------------------------------


SELECT
	 [SK_PatientID]
	,[ClinicalCode] Risk4QdiabetesScoreLatestCode
	,[EventDate] Risk4QdiabetesScoreLatestDateRecorded
	,[Value] QdiabetesLatestScore
	,[Units] QdiabetesLatestUnits

INTO  #Risk4QdiabetesScoreLatest

FROM
(
	SELECT

		 [SK_PatientID]
		,[ClinicalCode]
		,[EventDate]
		,[Value]
		,[Units]
		,row_number() over(partition by [SK_PatientID] order by [EventDate] DESC) as rn

	FROM  [08V].[GPEncounter]
	WHERE
		(
			[ClinicalCode] COLLATE Latin1_General_CS_AS = '38Gj'
			OR
			[ClinicalCode] COLLATE Latin1_General_CS_AS IN('21263', '212H')  --Diabetes resolved
		)
		AND
		SK_PatientID IN(SELECT SK_PatientID FROM ceg.GenomicsData WHERE CCG = '08V')

)as Risk4QdiabetesScoreLatest

WHERE rn = 1 AND [ClinicalCode] NOT IN('21263', '212H')	-- Exclude if there is a more recent diabetes diagnostic codes

GO


------------------ CVD Qrisk (10 year risk score ) ≥20% --------------------------------------------------------------------------------------
---------------------- Earliest --------------------------------------------------------------------------------------------------------------


SELECT
	 [SK_PatientID]
	,[ClinicalCode] CVDQriskScoreEarliestCode
	,[EventDate] CVDQriskScoreEarliestDateRecorded
	,[Value] CVDQriskScoreEarliestScore
	,[Units] CVDQriskScoreEarliestUnits

INTO  #CVDQriskScoreEarliest

FROM
(
	SELECT

		 [SK_PatientID]
		,[ClinicalCode]
		,[EventDate]
		,[Value]
		,[Units]
		,row_number() over(partition by [SK_PatientID] order by [EventDate] ASC) as rn

	FROM  [08V].[GPEncounter]
	WHERE
		(
			[ClinicalCode] COLLATE Latin1_General_CS_AS IN ('38DP', '38DF',  'EMISNQQR1', 'EMISNQQR2')
			AND
			Value >= 20
		)
		AND
		SK_PatientID IN(SELECT SK_PatientID FROM ceg.GenomicsData WHERE CCG = '08V')

)as CVDQriskScoreEarliest

WHERE rn = 1

GO

---------------------- Latest --------------------------------------------------------------------------------------------------------------


SELECT
	 [SK_PatientID]
	,[ClinicalCode] CVDQriskScoreLatestCode
	,[EventDate] CVDQriskScoreLatestDateRecorded
	,[Value] CVDQriskScoreLatestScore
	,[Units] CVDQriskScoreLatestUnits

INTO  #CVDQriskScoreLatest

FROM
(
	SELECT

		 [SK_PatientID]
		,[ClinicalCode]
		,[EventDate]
		,[Value]
		,[Units]
		,row_number() over(partition by [SK_PatientID] order by [EventDate] DESC) as rn

	FROM  [08V].[GPEncounter]
	WHERE
		(
			[ClinicalCode] COLLATE Latin1_General_CS_AS IN ('38DP', '38DF',  'EMISNQQR1', 'EMISNQQR2')
			AND
			Value >= 20
		)
		AND
		SK_PatientID IN(SELECT SK_PatientID FROM ceg.GenomicsData WHERE CCG = '08V')

)as CVDQriskScoreLatest

WHERE rn = 1

GO


-- DIABETES COMPLICATIONS ---------------------------------------------------------------------------------------------------------------------------------
------------------ Diabetic Retinopathy screening  --------------------------------------------------------------------------------------------------
/*
2BB, 68A8, 9N1v, 9N2f, 8I6F, 8I3X
*/


SELECT
	 [SK_PatientID]
	,[ClinicalCode] DiabeticRetinopathyScreeningLatestCode
	,[EventDate] DiabeticRetinopathyScreeningLatestDateRecorded


INTO  #DiabeticRetinopathyScreening

FROM
(
	SELECT

		 [SK_PatientID]
		,[ClinicalCode]
		,[EventDate]
		,row_number() over(partition by [SK_PatientID] order by [EventDate] DESC) as rn

	FROM  [08V].[GPEncounter]
	WHERE
		(
			--68A8, 9N1v, 9N2f, 8I6F, 8I3X							-- Synonym
			[ClinicalCode] COLLATE Latin1_General_CS_AS IN ('2BB', '2BB-1', '68A8', '9N1v', '9N2f', '8I6F', '8I3X')
		)
		AND
		SK_PatientID IN(SELECT SK_PatientID FROM ceg.GenomicsData WHERE CCG = '08V')

)as DiabeticRetinopathyScreening

WHERE rn = 1

GO


------------------ LEFT EYE Diabetic Retinopathy/Maculopathy Diagnosis --------------------------------------------------------------------------------------------------
---------------------- Earliest --------------------------------------------------------------------------------------------------------------
/*
2BBQ, 2BBS, 2BBV , EMISQLE13, 2BBX, 2BBl
*/

SELECT
	 [SK_PatientID]
	,[ClinicalCode] RetinopathyLeftEarliestCode
	,[EventDate] RetinopathyLeftEarliestDateRecorded


INTO  #RetinopathyLeftEarliest

FROM
(
	SELECT

		 [SK_PatientID]
		,[ClinicalCode]
		,[EventDate]
		,row_number() over(partition by [SK_PatientID] order by [EventDate] ASC) as rn

	FROM  [08V].[GPEncounter]
	WHERE
		(
			[ClinicalCode] COLLATE Latin1_General_CS_AS IN ('2BBQ', '2BBS', '2BBV', 'EMISQLE13', '2BBX', '2BBl')
		)
		AND
		SK_PatientID IN(SELECT SK_PatientID FROM ceg.GenomicsData WHERE CCG = '08V')

)AS RetinopathyLeftEarliest

WHERE rn = 1

GO

---------------------- Latest --------------------------------------------------------------------------------------------------------------


SELECT
	 [SK_PatientID]
	,[ClinicalCode] RetinopathyLeftLatestCode
	,[EventDate] RetinopathyLeftLatestDateRecorded


INTO  #RetinopathyLeftLatest

FROM
(
	SELECT

		 [SK_PatientID]
		,[ClinicalCode]
		,[EventDate]
		,row_number() over(partition by [SK_PatientID] order by [EventDate] DESC) as rn

	FROM  [08V].[GPEncounter]
	WHERE
		(
			[ClinicalCode] COLLATE Latin1_General_CS_AS IN ('2BBQ', '2BBS', '2BBV', 'EMISQLE13', '2BBX', '2BBl')
		)
		AND
		SK_PatientID IN(SELECT SK_PatientID FROM ceg.GenomicsData WHERE CCG = '08V')

) AS RetinopathyLeftLatest

WHERE rn = 1

GO


------------------ LEFT EYE NO Diabetic Retinopathy/Maculopathye --------------------------------------------------------------------------------------------------
---------------------- Earliest --------------------------------------------------------------------------------------------------------------
/*
2BBK, 2BBj
*/


SELECT
	 [SK_PatientID]
	,[ClinicalCode] NoRetinopathyLeftEarliestCode
	,[EventDate] NoRetinopathyLeftEarliestDateRecorded


INTO  #NoRetinopathyLeftEarliest

FROM
(
	SELECT

		 [SK_PatientID]
		,[ClinicalCode]
		,[EventDate]
		,row_number() over(partition by [SK_PatientID] order by [EventDate] ASC) as rn

	FROM  [08V].[GPEncounter]
	WHERE
		(
			[ClinicalCode] COLLATE Latin1_General_CS_AS IN ('2BBK', '2BBj')
		)
		AND
		SK_PatientID IN(SELECT SK_PatientID FROM ceg.GenomicsData WHERE CCG = '08V')

) AS NoRetinopathyLeftEarliest

WHERE rn = 1

GO

---------------------- Latest --------------------------------------------------------------------------------------------------------------


SELECT
	 [SK_PatientID]
	,[ClinicalCode] NoRetinopathyLeftLatestCode
	,[EventDate] NoRetinopathyLeftLatestDateRecorded


INTO  #NoRetinopathyLeftLatest

FROM
(
	SELECT

		 [SK_PatientID]
		,[ClinicalCode]
		,[EventDate]
		,row_number() over(partition by [SK_PatientID] order by [EventDate] DESC) as rn

	FROM  [08V].[GPEncounter]
	WHERE
		(
			[ClinicalCode] COLLATE Latin1_General_CS_AS IN ('2BBK', '2BBj')
		)
		AND
		SK_PatientID IN(SELECT SK_PatientID FROM ceg.GenomicsData WHERE CCG = '08V')

) AS NoRetinopathyLeftLatest

WHERE rn = 1

GO


------------------ RIGHT EYE Diabetic Retinopathy/Maculopathy  Diagnosis --------------------------------------------------------------------------------------------------
---------------------- Earliest --------------------------------------------------------------------------------------------------------------
/*
2BBP, 2BBR, 2BBT, EMISQRI13, 2BBW, 2BBk
*/

SELECT
	 [SK_PatientID]
	,[ClinicalCode] RetinopathyRightEyeEarliestCode
	,[EventDate] RetinopathyRightEyeEarliestDateRecorded


INTO  #RetinopathyRightEyeEarliest

FROM
(
	SELECT

		 [SK_PatientID]
		,[ClinicalCode]
		,[EventDate]
		,row_number() over(partition by [SK_PatientID] order by [EventDate] ASC) as rn

	FROM  [08V].[GPEncounter]
	WHERE
		(
			[ClinicalCode] COLLATE Latin1_General_CS_AS IN ('2BBP', '2BBR', '2BBT', 'EMISQRI13', '2BBW', '2BBk')
		)
		AND
		SK_PatientID IN(SELECT SK_PatientID FROM ceg.GenomicsData WHERE CCG = '08V')

) AS RetinopathyRightEyeEarliest

WHERE rn = 1

GO

---------------------- Latest --------------------------------------------------------------------------------------------------------------


SELECT
	 [SK_PatientID]
	,[ClinicalCode] RetinopathyRightEyeLatestCode
	,[EventDate] RetinopathyRightEyeLatestDateRecorded


INTO  #RetinopathyRightEyeLatest

FROM
(
	SELECT

		 [SK_PatientID]
		,[ClinicalCode]
		,[EventDate]
		,row_number() over(partition by [SK_PatientID] order by [EventDate] DESC) as rn

	FROM  [08V].[GPEncounter]
	WHERE
		(
			[ClinicalCode] COLLATE Latin1_General_CS_AS IN ('2BBP', '2BBR', '2BBT', 'EMISQRI13', '2BBW', '2BBk')
		)
		AND
		SK_PatientID IN(SELECT SK_PatientID FROM ceg.GenomicsData WHERE CCG = '08V')

) AS RetinopathyRightEyeLatest

WHERE rn = 1

GO


------------------ RIGHT EYE NO Diabetic Retinopathy/Maculopathy --------------------------------------------------------------------------------------------------
---------------------- Earliest --------------------------------------------------------------------------------------------------------------
/*
22BJ, 2BBi
*/

SELECT
	 [SK_PatientID]
	,[ClinicalCode] NoRetinopathyRightEarliestCode
	,[EventDate] NoRetinopathyRightEarliestDateRecorded


INTO  #NoRetinopathyRightEarliest

FROM
(
	SELECT

		 [SK_PatientID]
		,[ClinicalCode]
		,[EventDate]
		,row_number() over(partition by [SK_PatientID] order by [EventDate] ASC) as rn

	FROM  [08V].[GPEncounter]
	WHERE
		(
			[ClinicalCode] COLLATE Latin1_General_CS_AS IN ('2BBJ', '2BBi')	--	 (removed 22BJ)
		)
		AND
		SK_PatientID IN(SELECT SK_PatientID FROM ceg.GenomicsData WHERE CCG = '08V')

) AS NoRetinopathyRightEarliest

WHERE rn = 1

GO

---------------------- Latest --------------------------------------------------------------------------------------------------------------

SELECT
	 [SK_PatientID]
	,[ClinicalCode] NoRetinopathyRightLatestCode
	,[EventDate] NoRetinopathyRightLatestDateRecorded


INTO  #NoRetinopathyRightLatest

FROM
(
	SELECT

		 [SK_PatientID]
		,[ClinicalCode]
		,[EventDate]
		,row_number() over(partition by [SK_PatientID] order by [EventDate] DESC) as rn

	FROM  [08V].[GPEncounter]
	WHERE
		(
			[ClinicalCode] COLLATE Latin1_General_CS_AS IN ('2BBJ', '2BBi')	--	(removed 22BJ)
		)
		AND
		SK_PatientID IN(SELECT SK_PatientID FROM ceg.GenomicsData WHERE CCG = '08V')

) AS NoRetinopathyRightLatest

WHERE rn = 1

GO

------------------ Complains of erectile dysfunction earliest ever -----------------------------------------------------------------------
/*
1D1B, 1ABJ
*/

SELECT
	 [SK_PatientID]
	,[ClinicalCode] ComplainsEdCode
	,[EventDate] ComplainsEdDateRecorded

INTO  #ComplainsEdEarliest

FROM
(
	SELECT

		 [SK_PatientID]
		,[ClinicalCode]
		,[EventDate]
		,row_number() over(partition by [SK_PatientID] order by [EventDate] ASC) as rn

	FROM  [08V].[GPEncounter]
	WHERE
		(
			[ClinicalCode] COLLATE Latin1_General_CS_AS IN
			(
				'1598', '1598-1',	--Synonym
				'1ABB',
				'1ABC',
				'1D1B',
				'7C25E',
				'E2273', 'E2273-1'	--Synonym
			)  -- Removed '1ABJ'
		)
		AND
		SK_PatientID IN(SELECT SK_PatientID FROM ceg.GenomicsData WHERE CCG = '08V')

)as ComplainsEdEarliest

WHERE rn = 1

GO

-----------Peripheral Vascular Disease -----------------------------------------------------------------------------------
/*
G7310, G732, G7320, G7321, G7322, G7323, G7324, G733, G734, G735, G73y, G73y0, G73y1, G73z, G73z0, G73zz
*/


SELECT
	 [SK_PatientID]
	,[ClinicalCode] PVDCode
	,[EventDate]	PVDDateRecorded

INTO  #PVD

FROM
(
	SELECT
		 [SK_PatientID]
		,[ClinicalCode]
		,[EventDate]
		,row_number() over(partition by [SK_PatientID] order by [EventDate] ASC) as rn

	FROM  [08V].[GPEncounter]
	WHERE
		(
			[ClinicalCode] COLLATE Latin1_General_CS_AS IN ('G7310', 'G732', 'G7320', 'G7321', 'G7322', 'G7323', 'G7324', 'G733', 'G734', 'G735', 'G735-1',
															'G73y', 'G73y0', 'G73y1', 'G73z', 'G73z0', 'G73z0-1', 'G73z0-2', 'G73zz')
		)
		AND
		SK_PatientID IN(SELECT SK_PatientID FROM ceg.GenomicsData WHERE CCG = '08V')

)as PVD

WHERE rn = 1

GO


-----------Peripheral Arterial Disease [QOF]-----------------------------------------------------------------------------------


SELECT
	 [SK_PatientID]
	,[ClinicalCode] PADQoFCode
	,[EventDate] PADDateRecorded

INTO  #PAD_QoF

FROM
(
	SELECT
		 [SK_PatientID]
		,[ClinicalCode]
		,[EventDate]
		,row_number() over(partition by [SK_PatientID] order by [EventDate] ASC) as rn

	FROM  [08V].[GPEncounter]
	WHERE
		(
			(
				[ClinicalCode] COLLATE Latin1_General_CS_AS LIKE 'G73z%'
				AND
				[ClinicalCode] COLLATE Latin1_General_CS_AS <> 'G73z1'
			)
			OR
			[ClinicalCode] COLLATE Latin1_General_CS_AS LIKE 'G73%'	-- Synonym
			OR
			[ClinicalCode] COLLATE Latin1_General_CS_AS LIKE 'G74%'	-- Synonym
			OR
			[ClinicalCode] COLLATE Latin1_General_CS_AS IN ('G73y', 'Gyu74')
		)
		AND
		SK_PatientID IN(SELECT SK_PatientID FROM ceg.GenomicsData WHERE CCG = '08V')

)as pad

WHERE rn = 1

GO


-----------Abdominal Aortic Aneurysm-----------------------------------------------------------------------------------


SELECT
	 [SK_PatientID]
	,[ClinicalCode] AAAQoFCode
	,[EventDate] AAADateRecorded

INTO  #AAA_QoF

FROM
(
	SELECT
		 [SK_PatientID]
		,[ClinicalCode]
		,[EventDate]
		,row_number() over(partition by [SK_PatientID] order by [EventDate] ASC) as rn

	FROM  [08V].[GPEncounter]
	WHERE
		[ClinicalCode] COLLATE Latin1_General_CS_AS LIKE 'G71%'
		AND
		SK_PatientID IN(SELECT SK_PatientID FROM ceg.GenomicsData WHERE CCG = '08V')

)as aaa

WHERE rn = 1

GO


-----------Diabetic foot ulcer left-----------------------------------------------------------------------------------


SELECT
	 [SK_PatientID]
	,[ClinicalCode] FootUlcerLeftCode
	,[EventDate] FootUlcerLeftDateRecorded

INTO  #FootUlcerLeft

FROM
(
	SELECT
		 [SK_PatientID]
		,[ClinicalCode]
		,[EventDate]
		,row_number() over(partition by [SK_PatientID] order by [EventDate] ASC) as rn

	FROM  [08V].[GPEncounter]
	WHERE
		[ClinicalCode] COLLATE Latin1_General_CS_AS IN ('2G5L', '2G55')
		AND
		SK_PatientID IN(SELECT SK_PatientID FROM ceg.GenomicsData WHERE CCG = '08V')

)as FootUlcerLeft

WHERE rn = 1

GO

-----------Diabetic foot ulcer right-----------------------------------------------------------------------------------


SELECT
	 [SK_PatientID]
	,[ClinicalCode] FootUlcerRightCode
	,[EventDate] FootUlcerRightDateRecorded

INTO  #FootUlcerRight

FROM
(
	SELECT
		 [SK_PatientID]
		,[ClinicalCode]
		,[EventDate]
		,row_number() over(partition by [SK_PatientID] order by [EventDate] ASC) as rn

	FROM  [08V].[GPEncounter]
	WHERE
		[ClinicalCode] COLLATE Latin1_General_CS_AS IN ('2G5H', '2G54')
		AND
		SK_PatientID IN(SELECT SK_PatientID FROM ceg.GenomicsData WHERE CCG = '08V')

)as FootUlcerRight

WHERE rn = 1

GO


-----------Diabetic neuropathy-----------------------------------------------------------------------------------

SELECT
	 [SK_PatientID]
	,[ClinicalCode] NeuropathyCode
	,[EventDate] NeuropathyDateRecorded

INTO  #Neuropathy

FROM
(
	SELECT
		 [SK_PatientID]
		,[ClinicalCode]
		,[EventDate]
		,row_number() over(partition by [SK_PatientID] order by [EventDate] ASC) as rn

	FROM  [08V].[GPEncounter]
	WHERE
		(
			[ClinicalCode] COLLATE Latin1_General_CS_AS LIKE 'C106%'	--Synonym
			OR
			[ClinicalCode] COLLATE Latin1_General_CS_AS IN ('F367', 'F3y0')
		)
		AND
		SK_PatientID IN(SELECT SK_PatientID FROM ceg.GenomicsData WHERE CCG = '08V')

)as Neuropathy

WHERE rn = 1

GO

-----------Ischaemic Heart Disease\CHD  QOF-----------------------------------------------------------------------------------

SELECT
	 [SK_PatientID]
	,[ClinicalCode] IHD_CHDQoFCode
	,[EventDate] IHD_DateRecorded

INTO  #IHD_CHD_QoF

FROM
(
	SELECT
		 [SK_PatientID]
		,[ClinicalCode]
		,[EventDate]
		,row_number() over(partition by [SK_PatientID] order by [EventDate] ASC) as rn

	FROM  [08V].[GPEncounter]
	WHERE
		(
			   ([ClinicalCode] COLLATE Latin1_General_CS_AS LIKE 'G30%' AND [ClinicalCode] COLLATE Latin1_General_CS_AS <> 'G30A')
			OR ([ClinicalCode] COLLATE Latin1_General_CS_AS LIKE 'G31%' AND [ClinicalCode] COLLATE Latin1_General_CS_AS <> 'G310')
			OR ([ClinicalCode] COLLATE Latin1_General_CS_AS LIKE 'G33%' AND [ClinicalCode] COLLATE Latin1_General_CS_AS NOT IN('G331', 'G332'))
			OR ([ClinicalCode] COLLATE Latin1_General_CS_AS LIKE 'G34%' AND [ClinicalCode] COLLATE Latin1_General_CS_AS <> 'G341')
			OR [ClinicalCode] COLLATE Latin1_General_CS_AS LIKE 'G35%'
			OR [ClinicalCode] COLLATE Latin1_General_CS_AS LIKE 'G38%'
			OR [ClinicalCode] COLLATE Latin1_General_CS_AS LIKE 'G39%'
			OR [ClinicalCode] COLLATE Latin1_General_CS_AS LIKE 'G3y%'
			OR ([ClinicalCode] COLLATE Latin1_General_CS_AS LIKE 'Gyu3%' AND [ClinicalCode] COLLATE Latin1_General_CS_AS <> 'Gyu31')
			OR [ClinicalCode] COLLATE Latin1_General_CS_AS IN ('G3', 'G3-1', 'G3-2', 'G3-3')	--Synonym
			OR [ClinicalCode] COLLATE Latin1_General_CS_AS IN ('G32', 'G32-1')	--Synonym
			OR [ClinicalCode] COLLATE Latin1_General_CS_AS = 'G3z'
		)
		AND
		SK_PatientID IN(SELECT SK_PatientID FROM ceg.GenomicsData WHERE CCG = '08V')


)as ast

WHERE rn = 1

GO

----------- Atrial Fibrillation QOF (QOF definition for atrial fibrillation excludes atrial fibrillation resolved codes if appearing after the latest diagnostic code) ------------------

SELECT
	 [SK_PatientID]
	,[ClinicalCode] AFQoFCode
	,[EventDate]	AF_DateRecorded

INTO  #AF_QoF

FROM
(
	SELECT
		*

	FROM
	(
		SELECT
			 [SK_PatientID]
			,[ClinicalCode]
			,[EventDate]
			,row_number() over(partition by [SK_PatientID] order by [EventDate] ASC) as rn

		FROM
			[08V].[GPEncounter]

		WHERE
			[ClinicalCode] COLLATE Latin1_General_CS_AS LIKE 'G573%'
			AND
			SK_PatientID IN(SELECT SK_PatientID FROM ceg.GenomicsData WHERE CCG = '08V')

	)as af

	WHERE
		rn = 1

) AS Afib

WHERE
	SK_PatientID NOT IN	-- QOF definition for atrial fibrillation excludes atrial fibrillation resolved codes if appearing after the latest diagnostic code
	(
		SELECT
			[SK_PatientID]

		FROM
		(
			SELECT
				 [SK_PatientID]
				,[ClinicalCode]
				,[EventDate]
				,row_number() over(partition by [SK_PatientID] order by [EventDate] DESC) as rn

			FROM
				[08V].[GPEncounter]

			WHERE
				(
					[ClinicalCode] COLLATE Latin1_General_CS_AS LIKE 'G573%'
					OR
					[ClinicalCode] COLLATE Latin1_General_CS_AS = '212R' --AF resolved
				)
				AND
				SK_PatientID IN(SELECT SK_PatientID FROM ceg.GenomicsData WHERE CCG = '08V')

		) AS  Res

		WHERE
			rn = 1	AND	[ClinicalCode] = '212R' --AF resolved
	)

GO

-----------Heart failure QOF-----------------------------------------------------------------------------------

SELECT
	 [SK_PatientID]
	,[ClinicalCode] HFQoFCode
	,[EventDate] HFDateRecorded

INTO  #HF_QoF

FROM
(
	SELECT
		 [SK_PatientID]
		,[ClinicalCode]
		,[EventDate]
		,row_number() over(partition by [SK_PatientID] order by [EventDate] ASC) as rn

	FROM  [08V].[GPEncounter]
	WHERE
		(
			[ClinicalCode] COLLATE Latin1_General_CS_AS LIKE 'G58%'
			OR
			[ClinicalCode] COLLATE Latin1_General_CS_AS IN ('G1yz1', '662f', '662g', '662h', '662i')
		)
		AND
		SK_PatientID IN(SELECT SK_PatientID FROM ceg.GenomicsData WHERE CCG = '08V')

)as hf

WHERE rn = 1

GO

-----------Stroke/TIA [QOF]-----------------------------------------------------------------------------------

SELECT
	 [SK_PatientID]
	,[ClinicalCode] StrokeQoFCode
	,[EventDate]	StrokeDateRecorded

INTO  #Stroke_TIA_QoF

FROM
(
	SELECT
		 [SK_PatientID]
		,[ClinicalCode]
		,[EventDate]
		,row_number() over(partition by [SK_PatientID] order by [EventDate] ASC) as rn

	FROM  [08V].[GPEncounter]
	WHERE
		(
			([ClinicalCode] COLLATE Latin1_General_CS_AS LIKE 'G61%' AND [ClinicalCode] COLLATE Latin1_General_CS_AS <> 'G617')
			OR
			[ClinicalCode] COLLATE Latin1_General_CS_AS LIKE 'G64%'
			OR
			([ClinicalCode] COLLATE Latin1_General_CS_AS LIKE 'G65%' AND [ClinicalCode] COLLATE Latin1_General_CS_AS <> 'G655')
			OR
			([ClinicalCode] COLLATE Latin1_General_CS_AS LIKE 'G66%' AND [ClinicalCode] COLLATE Latin1_General_CS_AS <> 'G669')
			OR
			[ClinicalCode] COLLATE Latin1_General_CS_AS IN ('G63y0', 'G63y1', 'G6760', 'G6W', 'G6X', 'Gyu62', 'Gyu63', 'Gyu64', 'Gyu65','Gyu66', 'Gyu6F', 'Gyu6G', 'ZV12D', 'Fyu55')
		)
		AND
		SK_PatientID IN(SELECT SK_PatientID FROM ceg.GenomicsData WHERE CCG = '08V')

)as stroke

WHERE rn = 1

GO

----------- Hypertension QOF (QOF definition for Hypertension excludes Hypertension resolved codes if appearing after the latest diagnostic code) ------------------

SELECT
	 [SK_PatientID]
	,[ClinicalCode] HypertensionQoFCode
	,[EventDate]	HypertensionDateRecorded

INTO  #HypertensionQoF

FROM
(
	SELECT
		*

	FROM
	(
		SELECT
			 [SK_PatientID]
			,[ClinicalCode]
			,[EventDate]
			,row_number() over(partition by [SK_PatientID] order by [EventDate] ASC) as rn

		FROM
			[08V].[GPEncounter]

		WHERE
			(
					[ClinicalCode] COLLATE Latin1_General_CS_AS LIKE 'G20%'
				OR	[ClinicalCode] COLLATE Latin1_General_CS_AS LIKE 'G24%'
				OR	[ClinicalCode] COLLATE Latin1_General_CS_AS LIKE 'G25%'
				OR	[ClinicalCode] COLLATE Latin1_General_CS_AS IN ('G2', 'G2-1',		--Synonym
																	'G26', 'G26-1',		--Synonym
																	'G28', 'G2y', 'G2z', 'Gyu2', 'Gyu20')
			)
			AND
			SK_PatientID IN(SELECT SK_PatientID FROM ceg.GenomicsData WHERE CCG = '08V')

	)as hyper

	WHERE
		rn = 1

) AS Hypertension

WHERE
	SK_PatientID NOT IN	-- QOF definition for Hypertension excludes Hypertension resolved codes if appearing after the latest diagnostic code
	(
		SELECT
			[SK_PatientID]

		FROM
		(
			SELECT
				 [SK_PatientID]
				,[ClinicalCode]
				,[EventDate]
				,row_number() over(partition by [SK_PatientID] order by [EventDate] DESC) as rn

			FROM
				[08V].[GPEncounter]

			WHERE
				(
						[ClinicalCode] COLLATE Latin1_General_CS_AS LIKE 'G20%'
					OR	[ClinicalCode] COLLATE Latin1_General_CS_AS LIKE 'G24%'
					OR	[ClinicalCode] COLLATE Latin1_General_CS_AS LIKE 'G25%'
					OR	[ClinicalCode] COLLATE Latin1_General_CS_AS IN ('G2', 'G2-1',		--Synonym
																		'G26', 'G26-1',		--Synonym
																		'G28', 'G2y', 'G2z', 'Gyu2', 'Gyu20')
					OR	[ClinicalCode] COLLATE Latin1_General_CS_AS IN ('21261', '212K') -- Hypertension resolved
				)
				AND
				SK_PatientID IN(SELECT SK_PatientID FROM ceg.GenomicsData WHERE CCG = '08V')

		) AS  Res

		WHERE
			rn = 1	AND	[ClinicalCode] IN ('21261', '212K') -- Hypertension resolved
	)

GO

-----------Chronic kidney disease [QOF]-----------------------------------------------------------------------------------

SELECT
	 [SK_PatientID]
	,[ClinicalCode] CKDQoFCode
	,[EventDate]	CKDDateRecorded

INTO  #CKD_QoF

FROM
(
	SELECT
		 [SK_PatientID]
		,[ClinicalCode]
		,[EventDate]
		,row_number() over(partition by [SK_PatientID] order by [EventDate] ASC) as rn

	FROM  [08V].[GPEncounter]
	WHERE

		[ClinicalCode] COLLATE Latin1_General_CS_AS
		IN
		(
			'1Z12','1Z13', '1Z14', '1Z15', '1Z16',
			'1Z1B', '1Z1B-1', -- Synonym
			'1Z1C', '1Z1C-1', -- Synonym
			'1Z1D', '1Z1D-1', -- Synonym
			'1Z1E', '1Z1E-1', -- Synonym
			'1Z1F', '1Z1F-1', -- Synonym
			'1Z1G', '1Z1G-1', -- Synonym
			'1Z1H', '1Z1H-1', -- Synonym
			'1Z1J', '1Z1J-1', -- Synonym
			'1Z1K',	'1Z1K-1', -- Synonym
			'1Z1L', '1Z1L-1', -- Synonym
			'1Z1T', '1Z1V', '1Z1W', '1Z1X', '1Z1Y','1Z1Z', '1Z1a', '1Z1b', '1Z1c', '1Z1d', '1Z1e', '1Z1f','K053', 'K054', 'K055'
		)
		AND
		SK_PatientID IN(SELECT SK_PatientID FROM ceg.GenomicsData WHERE CCG = '08V')

)as ckd

WHERE rn = 1

GO

-- RELATED METABOLIC CONDITIONS ---------------------------------------------------------------------------------------------------------------------------------
-----------Polycystic ovarian syndrome --------------------------------------------------------------------------------

SELECT
	 [SK_PatientID]
	,[ClinicalCode] PCSCode
	,[EventDate] PCSDateRecorded

INTO  #PCS

FROM
(
	SELECT
		 [SK_PatientID]
		,[ClinicalCode]
		,[EventDate]
		,row_number() over(partition by [SK_PatientID] order by [EventDate] ASC) as rn

	FROM  [08V].[GPEncounter]
	WHERE
		[ClinicalCode] COLLATE Latin1_General_CS_AS = 'C165'
		AND
		SK_PatientID IN(SELECT SK_PatientID FROM ceg.GenomicsData WHERE CCG = '08V')

)as PCS

WHERE rn = 1

GO

-----------Obesity-related breathing disorders --------------------------------------------------------------------------------

SELECT
	 [SK_PatientID]
	,[ClinicalCode] ObesityDisorderCode
	,[EventDate] ObesityDisorderDateRecorded

INTO  #ObesityDisorder

FROM
(
	SELECT
		 [SK_PatientID]
		,[ClinicalCode]
		,[EventDate]
		,row_number() over(partition by [SK_PatientID] order by [EventDate] ASC) as rn

	FROM  [08V].[GPEncounter]
	WHERE
		[ClinicalCode] COLLATE Latin1_General_CS_AS IN ('Fy03', 'Fy03-1','C3802', 'C38y0', 'C38y0-1')	--Synonym ends with -1
		AND
		SK_PatientID IN(SELECT SK_PatientID FROM ceg.GenomicsData WHERE CCG = '08V')

)as PCS

WHERE rn = 1

GO

-----------Fatty liver ------------------------------------------------------------------------------------------------------

SELECT
	 [SK_PatientID]
	,[ClinicalCode] NAFLDCode
	,[EventDate] NAFLDDateRecorded

INTO  #NAFLD

FROM
(
	SELECT
		 [SK_PatientID]
		,[ClinicalCode]
		,[EventDate]
		,row_number() over(partition by [SK_PatientID] order by [EventDate] ASC) as rn

	FROM  [08V].[GPEncounter]
	WHERE
		[ClinicalCode] COLLATE Latin1_General_CS_AS IN ('J61y', 'J615', 'J615-1')
		AND
		SK_PatientID IN(SELECT SK_PatientID FROM ceg.GenomicsData WHERE CCG = '08V')

)as NAFLD

WHERE rn = 1

GO


-----------Familial hypercholesterolaemia --------------------------------------------------------------------------------


SELECT
	 [SK_PatientID]
	,[ClinicalCode] HypercholesterolCode
	,[EventDate] HypercholesterolDateRecorded

INTO  #Hypercholesterol

FROM
(
	SELECT
		 [SK_PatientID]
		,[ClinicalCode]
		,[EventDate]
		,row_number() over(partition by [SK_PatientID] order by [EventDate] ASC) as rn

	FROM  [08V].[GPEncounter]
	WHERE
		[ClinicalCode] COLLATE Latin1_General_CS_AS IN ('C3200', 'C3201', 'C3204', 'C3205', 'C3203', 'C3220')
		AND
		SK_PatientID IN(SELECT SK_PatientID FROM ceg.GenomicsData WHERE CCG = '08V')

)as Hypercholesterol

WHERE rn = 1

GO


------------------ Acanthosis earliest ever -----------------------------------------------------------------------
/*
PH3y5, M212
*/

SELECT
	 [SK_PatientID]
	,[ClinicalCode] AcanthosisCode
	,[EventDate] AcanthosisDateRecorded

INTO  #Acanthosis

FROM
(
	SELECT

		 [SK_PatientID]
		,[ClinicalCode]
		,[EventDate]
		,row_number() over(partition by [SK_PatientID] order by [EventDate] ASC) as rn

	FROM  [08V].[GPEncounter]
	WHERE
		(
			[ClinicalCode] COLLATE Latin1_General_CS_AS IN ('PH3y5', 'M212')		--Acanthosis nigricans, congenital, Acquired acanthosis nigricans
		)
		AND
		SK_PatientID IN(SELECT SK_PatientID FROM ceg.GenomicsData WHERE CCG = '08V')

)as Acanthosis

WHERE rn = 1

GO


-- RELATED AUTOIMMUNE & ENDOCRINE ------------------------------------------------------------------------------------------------
-----------Thyroid conditions --------------------------------------------------------------------------------


SELECT
	 [SK_PatientID]
	,[ClinicalCode] ThyroidCode
	,[EventDate]	ThyroidDateRecorded

INTO  #Thyroid

FROM
(
	SELECT
		 [SK_PatientID]
		,[ClinicalCode]
		,[EventDate]
		,row_number() over(partition by [SK_PatientID] order by [EventDate] ASC) as rn

	FROM  [08V].[GPEncounter]
	WHERE
		(		-- C02%, C03%, C04%, 1JM%, 1431, 1432, F3814, Q4337, C0A5
				[ClinicalCode] COLLATE Latin1_General_CS_AS LIKE 'C02%'
			OR	[ClinicalCode] COLLATE Latin1_General_CS_AS LIKE 'C03%'
			OR	[ClinicalCode] COLLATE Latin1_General_CS_AS LIKE 'C04%'
			OR	[ClinicalCode] COLLATE Latin1_General_CS_AS LIKE '1JM%'
			OR	[ClinicalCode] COLLATE Latin1_General_CS_AS IN ('1431', '1431-1', '1432', 'F3814', 'Q4337', 'C0A5')
		)
		AND
		SK_PatientID IN(SELECT SK_PatientID FROM ceg.GenomicsData WHERE CCG = '08V')

)as Thyroid

WHERE rn = 1

GO

-----------Autoimmune and endocrine conditions (non-thyroid) --------------------------------------------------------

SELECT
	 [SK_PatientID]
	,[ClinicalCode] AutoimmuneCode
	,[EventDate]	AutoimmuneDateRecorded

INTO  #Autoimmune

FROM
(
	SELECT
		 [SK_PatientID]
		,[ClinicalCode]
		,[EventDate]
		,row_number() over(partition by [SK_PatientID] order by [EventDate] ASC) as rn

	FROM  [08V].[GPEncounter]
	WHERE
		(
				[ClinicalCode] COLLATE Latin1_General_CS_AS LIKE 'J690%'
			OR	[ClinicalCode] COLLATE Latin1_General_CS_AS IN ('C1541', 'C1546',
																'C154z', 'C154z-1', 'C154z-2',			--Synonym
																'C180', 'C180-1', 						--Synonym
																'C181', 'C181-1', 'C181-2',				--Synonym
																'C182', 'C18y', 'C18z',
																'D010', 'D010-1', 'D010-2', 'D010-3',	--Synonym
																'M2951')
		)
		AND
		SK_PatientID IN(SELECT SK_PatientID FROM ceg.GenomicsData WHERE CCG = '08V')

)as Autoimmune

WHERE rn = 1

GO

-- Miscellaneous ---------------------------------------------------------------------------------------------------------------------------------
-----------Frailty index -----------------------------------------------------------------------------------------------------


SELECT
	 [SK_PatientID]
	,[ClinicalCode] FrailtyindexCode
	,[EventDate]	FrailtyindexDateRecorded

INTO  #Frailtyindex

FROM
(
	SELECT
		 [SK_PatientID]
		,[ClinicalCode]
		,[EventDate]
		,row_number() over(partition by [SK_PatientID] order by [EventDate] ASC) as rn

	FROM  [08V].[GPEncounter]
	WHERE
		(	--38Ql, 2Jd0, 2Jd1, 2Jd2
			[ClinicalCode] COLLATE Latin1_General_CS_AS = '38Ql'	-- Frailty index
			OR
			[ClinicalCode] COLLATE Latin1_General_CS_AS = '2Jd0'	-- Mild frailty
			OR
			[ClinicalCode] COLLATE Latin1_General_CS_AS = '2Jd1'	-- moderate frailty
			OR
			[ClinicalCode] COLLATE Latin1_General_CS_AS = '2Jd2'	-- Severe frailty
		)
		AND
		SK_PatientID IN(SELECT SK_PatientID FROM ceg.GenomicsData WHERE CCG = '08V')

)as Frailtyindex

WHERE rn = 1

GO

-----------NHS Health Check (NHS HC) --------------------------------------------------------------------------------


SELECT
	 [SK_PatientID]
	,[ClinicalCode] NHS_HC_Code
	,[EventDate]	NHS_HC_DateRecorded

INTO  #NHS_HC

FROM
(
	SELECT
		 [SK_PatientID]
		,[ClinicalCode]
		,[EventDate]
		,row_number() over(partition by [SK_PatientID] order by [EventDate] DESC) as rn

	FROM  [08V].[GPEncounter]
	WHERE
		[ClinicalCode] COLLATE Latin1_General_CS_AS LIKE '8BAg%'
		AND
		SK_PatientID IN(SELECT SK_PatientID FROM ceg.GenomicsData WHERE CCG = '08V')

)as NHSns

WHERE rn = 1

GO


--------------------------------------------------------------------------------------------------------------------------
---------------------------------------- Joining temp tables ------------------------------------------------------------------
--------------------------------------------------------------------------------------------------------------------------


SELECT

	(Select CommissionerName From Dictionary.[dbo].[Commissioner] Where CommissionerCode = '08V') AS Locality
	,GD.EncryptedNHSNumber

	--Demographics
	,GD.SK_PatientID
	,P.Gender
	,FactEngine.[shared].[fGetAge](GD.[DateOfBirth], '2018-10-01') AS [Age 2018-10-01]
	,GD.[DateOfBirth]
	,DatePart(year, GD.[DateOfBirth]) AS [Year of Birth]

	,[YearOfDeath]
	,[DateRegistered]
	,[DateRegisteredEnd]

	,CASE
		WHEN Ethnicity.EthnicityCode IS NOT NULL THEN Ethnicity.EthnicityCode COLLATE DATABASE_DEFAULT
		ELSE P.[EthnicityCode] COLLATE DATABASE_DEFAULT
	 END AS "Ethnicity Code"


	,P.[IMD 2010 Score]
	,P.LSOACode

	,CountryOfBirth.CountryOfBirthCode
	,CountryOfBirth.CountryOfBirthTerm

	--FAMILY HISTORY
	,FH_Diabetes.FH_DiabetesCode
	,FH_Diabetes.FH_DiabetesDate

	,FH_IHD.FH_IHDCode
	,FH_IHD.FH_IHDDate

	--DIABETES DIAGNOSES
	,T1.DiabetesT1Code
	,T1.DiabetesT1DateRecorded

	,T2.DiabetesT2Code
	,T2.DiabetesT2DateRecorded

	,SecondaryDiabetesQoF.SecondaryDiabetesCode
	,SecondaryDiabetesQoF.SecondaryDiabetesDateRecorded

	,OtherTypesOfDiabetes.OtherTypesOfDiabetesCode
	,OtherTypesOfDiabetes.OtherTypesOfDiabetesDateRecorded

	,PancreaticDisease.PancreaticDiseaseCode
	,PancreaticDisease.PancreaticDiseaseDateRecorded

	,GestationalDiabetes.GestationalDiabetesCode
	,GestationalDiabetes.GestationalDiabetesDateRecorded

	,EmergenciesDiabetes.EmergenciesDiabetesCode
	,EmergenciesDiabetes.EmergenciesDiabetesDateRecorded

	,BariatricSurgery.BariatricSurgeryCode
	,BariatricSurgery.BariatricSurgeryDateRecorded

	-- DIABETES RISK SCORES
	,Risk1PrediabetesEarliest.PrediabetesRisk1EarliestCode
	,Risk1PrediabetesEarliest.PrediabetesRisk1EarliestDateRecorded

	,Risk1PrediabetesLatest.PrediabetesRisk1LatestCode
	,Risk1PrediabetesLatest.PrediabetesRisk1LatestDateRecorded

	,Risk2MellitusEarliest.Risk2MellitusEarliestCode
	,Risk2MellitusEarliest.Risk2MellitusEarliestDateRecorded

	,Risk2MellitusLatest.Risk2MellitusLatestCode
	,Risk2MellitusLatest.Risk2MellitusLatestDateRecorded

	,Risk3EarliestHbA1c.Risk3EarliestHbA1cCode
	,Risk3EarliestHbA1c.Risk3EarliestHbA1cDateRecorded
	,Risk3EarliestHbA1c.Risk3EarliestHbA1cValue
	,Risk3EarliestHbA1c.Risk3EarliestHbA1cUnits

	,Risk3LatestHbA1c.Risk3LatestHbA1cCode
	,Risk3LatestHbA1c.Risk3LatestHbA1cDateRecorded
	,Risk3LatestHbA1c.Risk3LatestHbA1cValue
	,Risk3LatestHbA1c.Risk3LatestHbA1cUnits

	,Risk4QdiabetesScoreEarliest.Risk4QdiabetesScoreEarliestCode
	,Risk4QdiabetesScoreEarliest.Risk4QdiabetesScoreEarliestDateRecorded
	,Risk4QdiabetesScoreEarliest.QdiabetesEarliestScore
	,Risk4QdiabetesScoreEarliest.QdiabetesEarliestUnits

	,Risk4QdiabetesScoreLatest.Risk4QdiabetesScoreLatestCode
	,Risk4QdiabetesScoreLatest.Risk4QdiabetesScoreLatestDateRecorded
	,Risk4QdiabetesScoreLatest.QdiabetesLatestScore
	,Risk4QdiabetesScoreLatest.QdiabetesLatestUnits

	,CVDQriskScoreEarliest.CVDQriskScoreEarliestCode
	,CVDQriskScoreEarliest.CVDQriskScoreEarliestDateRecorded
	,CVDQriskScoreEarliest.CVDQriskScoreEarliestScore
	,CVDQriskScoreEarliest.CVDQriskScoreEarliestUnits

	,CVDQriskScoreLatest.CVDQriskScoreLatestCode
	,CVDQriskScoreLatest.CVDQriskScoreLatestDateRecorded
	,CVDQriskScoreLatest.CVDQriskScoreLatestScore
	,CVDQriskScoreLatest.CVDQriskScoreLatestUnits

	-- DIABETES COMPLICATIONS

	,DiabeticRetinopathyScreening.DiabeticRetinopathyScreeningLatestCode
	,DiabeticRetinopathyScreening.DiabeticRetinopathyScreeningLatestDateRecorded

	,RetinopathyLeftEarliest.RetinopathyLeftEarliestCode
	,RetinopathyLeftEarliest.RetinopathyLeftEarliestDateRecorded

	,RetinopathyLeftLatest.RetinopathyLeftLatestCode
	,RetinopathyLeftLatest.RetinopathyLeftLatestDateRecorded

	,NoRetinopathyLeftEarliest.NoRetinopathyLeftEarliestCode
	,NoRetinopathyLeftEarliest.NoRetinopathyLeftEarliestDateRecorded

	,NoRetinopathyLeftLatest.NoRetinopathyLeftLatestCode
	,NoRetinopathyLeftLatest.NoRetinopathyLeftLatestDateRecorded


	,RetinopathyRightEyeEarliest.RetinopathyRightEyeEarliestCode
	,RetinopathyRightEyeEarliest.RetinopathyRightEyeEarliestDateRecorded

	,RetinopathyRightEyeLatest.RetinopathyRightEyeLatestCode
	,RetinopathyRightEyeLatest.RetinopathyRightEyeLatestDateRecorded

	,NoRetinopathyRightEarliest.NoRetinopathyRightEarliestCode
	,NoRetinopathyRightEarliest.NoRetinopathyRightEarliestDateRecorded

	,NoRetinopathyRightLatest.NoRetinopathyRightLatestCode
	,NoRetinopathyRightLatest.NoRetinopathyRightLatestDateRecorded


	,ComplainsEdEarliest.ComplainsEdCode
	,ComplainsEdEarliest.ComplainsEdDateRecorded

	,PVD.PVDCode
	,PVD.PVDDateRecorded

	,PAD_QoF.PADQoFCode
	,PAD_QoF.PADDateRecorded

	,AAA_QoF.AAAQoFCode
	,AAA_QoF.AAAQoFCode

	,FootUlcerLeft.FootUlcerLeftCode
	,FootUlcerLeft.FootUlcerLeftDateRecorded

	,FootUlcerRight.FootUlcerRightCode
	,FootUlcerRight.FootUlcerRightDateRecorded

	,Neuropathy.NeuropathyCode
	,Neuropathy.NeuropathyDateRecorded

	,IHD_CHD_QoF.IHD_CHDQoFCode
	,IHD_CHD_QoF.IHD_DateRecorded

	,AF_QoF.AFQoFCode
	,AF_QoF.AF_DateRecorded

	,HF_QoF.HFQoFCode
	,HF_QoF.HFDateRecorded

	,Stroke_TIA_QoF.StrokeQoFCode
	,Stroke_TIA_QoF.StrokeDateRecorded

	,HypertensionQoF.HypertensionQoFCode
	,HypertensionQoF.HypertensionDateRecorded

	,CKD_QoF.CKDQoFCode
	,CKD_QoF.CKDDateRecorded


	--RELATED METABOLIC CONDITIONS
	,PCS.PCSCode
	,PCS.PCSDateRecorded

	,ObesityDisorder.ObesityDisorderCode
	,ObesityDisorder.ObesityDisorderDateRecorded

	,NAFLD.NAFLDCode
	,NAFLD.NAFLDDateRecorded

	,Hypercholesterol.HypercholesterolCode
	,Hypercholesterol.HypercholesterolDateRecorded

	,Acanthosis.AcanthosisCode
	,Acanthosis.AcanthosisDateRecorded

	-- RELATED AUTOIMMUNE & ENDOCRINE
	,Thyroid.ThyroidCode
	,Thyroid.ThyroidDateRecorded

	,Autoimmune.AutoimmuneCode
	,Autoimmune.AutoimmuneDateRecorded

	--Miscellaneous
	,Frailtyindex.FrailtyindexCode
	,Frailtyindex.FrailtyindexDateRecorded

	,NHS_HC.NHS_HC_Code
	,NHS_HC.NHS_HC_DateRecorded

FROM
	ceg.GenomicsData GD

	INNER JOIN #Demographics P ON P.SK_PatientID = GD.SK_PatientID
	LEFT JOIN #Ethnicity as Ethnicity ON Ethnicity.SK_PatientID = P.SK_PatientID
	LEFT JOIN #CountryOfBirth CountryOfBirth ON CountryOfBirth.SK_PatientID = P.SK_PatientID

	LEFT JOIN  #FH_Diabetes as FH_Diabetes ON FH_Diabetes.SK_PatientID = P.SK_PatientID
	LEFT JOIN  #FH_IHD as FH_IHD ON FH_IHD.SK_PatientID = P.SK_PatientID

	LEFT JOIN  #DiabetesT1QoF as T1 ON T1.SK_PatientID = P.SK_PatientID
	LEFT JOIN  #DiabetesT2QoF as T2 ON T2.SK_PatientID = P.SK_PatientID
	LEFT JOIN  #SecondaryDiabetesQoF as SecondaryDiabetesQoF ON SecondaryDiabetesQoF.SK_PatientID = P.SK_PatientID
	LEFT JOIN  #OtherTypesOfDiabetes as OtherTypesOfDiabetes ON OtherTypesOfDiabetes.SK_PatientID = P.SK_PatientID
	LEFT JOIN  #PancreaticDisease AS PancreaticDisease ON PancreaticDisease.SK_PatientID = P.SK_PatientID
	LEFT JOIN  #GestationalDiabetes as GestationalDiabetes ON GestationalDiabetes.SK_PatientID = P.SK_PatientID
	LEFT JOIN  #EmergenciesDiabetes AS EmergenciesDiabetes ON EmergenciesDiabetes.SK_PatientID = P.SK_PatientID
	LEFT JOIN  #BariatricSurgery AS BariatricSurgery ON BariatricSurgery.SK_PatientID = P.SK_PatientID

	LEFT JOIN  #Risk1PrediabetesEarliest  AS Risk1PrediabetesEarliest ON Risk1PrediabetesEarliest.SK_PatientID = P.SK_PatientID
	LEFT JOIN  #Risk1PrediabetesLatest AS Risk1PrediabetesLatest ON Risk1PrediabetesLatest.SK_PatientID = P.SK_PatientID
	LEFT JOIN  #Risk2MellitusEarliest AS Risk2MellitusEarliest ON Risk2MellitusEarliest.SK_PatientID = P.SK_PatientID
	LEFT JOIN  #Risk2MellitusLatest AS Risk2MellitusLatest ON Risk2MellitusLatest.SK_PatientID = P.SK_PatientID

	LEFT JOIN  #Risk3EarliestHbA1c AS Risk3EarliestHbA1c ON Risk3EarliestHbA1c.SK_PatientID = P.SK_PatientID
	LEFT JOIN  #Risk3LatestHbA1c AS Risk3LatestHbA1c ON Risk3LatestHbA1c.SK_PatientID = P.SK_PatientID

	LEFT JOIN  #Risk4QdiabetesScoreEarliest AS Risk4QdiabetesScoreEarliest ON Risk4QdiabetesScoreEarliest.SK_PatientID = P.SK_PatientID
	LEFT JOIN  #Risk4QdiabetesScoreLatest AS Risk4QdiabetesScoreLatest ON Risk4QdiabetesScoreLatest.SK_PatientID = P.SK_PatientID
	LEFT JOIN  #CVDQriskScoreEarliest AS CVDQriskScoreEarliest ON CVDQriskScoreEarliest.SK_PatientID = P.SK_PatientID
	LEFT JOIN  #CVDQriskScoreLatest AS CVDQriskScoreLatest ON CVDQriskScoreLatest.SK_PatientID = P.SK_PatientID

	-- DIABETES COMPLICATIONS
	LEFT JOIN  #DiabeticRetinopathyScreening AS DiabeticRetinopathyScreening ON DiabeticRetinopathyScreening.SK_PatientID = P.SK_PatientID

	LEFT JOIN  #RetinopathyLeftEarliest AS RetinopathyLeftEarliest ON RetinopathyLeftEarliest.SK_PatientID = P.SK_PatientID
	LEFT JOIN  #RetinopathyLeftLatest AS RetinopathyLeftLatest ON RetinopathyLeftLatest.SK_PatientID = P.SK_PatientID

	LEFT JOIN  #NoRetinopathyLeftEarliest AS NoRetinopathyLeftEarliest ON NoRetinopathyLeftEarliest.SK_PatientID = P.SK_PatientID
	LEFT JOIN  #NoRetinopathyLeftLatest AS NoRetinopathyLeftLatest ON NoRetinopathyLeftLatest.SK_PatientID = P.SK_PatientID

	LEFT JOIN  #RetinopathyRightEyeEarliest AS RetinopathyRightEyeEarliest ON RetinopathyRightEyeEarliest.SK_PatientID = P.SK_PatientID
	LEFT JOIN  #RetinopathyRightEyeLatest AS RetinopathyRightEyeLatest ON RetinopathyRightEyeLatest.SK_PatientID = P.SK_PatientID

	LEFT JOIN  #NoRetinopathyRightEarliest AS NoRetinopathyRightEarliest ON NoRetinopathyRightEarliest.SK_PatientID = P.SK_PatientID
	LEFT JOIN  #NoRetinopathyRightLatest AS NoRetinopathyRightLatest ON NoRetinopathyRightLatest.SK_PatientID = P.SK_PatientID

	LEFT JOIN  #ComplainsEdEarliest AS ComplainsEdEarliest ON ComplainsEdEarliest.SK_PatientID = P.SK_PatientID

	LEFT JOIN  #PVD AS PVD ON PVD.SK_PatientID = P.SK_PatientID
	LEFT JOIN  #PAD_QoF AS PAD_QoF ON PAD_QoF.SK_PatientID = P.SK_PatientID
	LEFT JOIN  #AAA_QoF AS AAA_QoF ON AAA_QoF.SK_PatientID = P.SK_PatientID

	LEFT JOIN  #FootUlcerLeft AS FootUlcerLeft ON FootUlcerLeft.SK_PatientID = P.SK_PatientID
	LEFT JOIN  #FootUlcerRight AS FootUlcerRight ON FootUlcerRight.SK_PatientID = P.SK_PatientID

	LEFT JOIN  #Neuropathy AS Neuropathy ON Neuropathy.SK_PatientID = P.SK_PatientID
	LEFT JOIN  #IHD_CHD_QoF AS IHD_CHD_QoF ON IHD_CHD_QoF.SK_PatientID = P.SK_PatientID
	LEFT JOIN  #AF_QoF AS AF_QoF ON AF_QoF.SK_PatientID = P.SK_PatientID
	LEFT JOIN  #HF_QoF AS HF_QoF ON HF_QoF.SK_PatientID = P.SK_PatientID
	LEFT JOIN  #Stroke_TIA_QoF AS Stroke_TIA_QoF ON Stroke_TIA_QoF.SK_PatientID = P.SK_PatientID
	LEFT JOIN  #HypertensionQoF AS HypertensionQoF ON HypertensionQoF.SK_PatientID = P.SK_PatientID
	LEFT JOIN  #CKD_QoF AS CKD_QoF ON CKD_QoF.SK_PatientID = P.SK_PatientID


	--RELATED METABOLIC CONDITIONS
	LEFT JOIN  #PCS AS PCS ON PCS.SK_PatientID = P.SK_PatientID
	LEFT JOIN  #ObesityDisorder AS ObesityDisorder ON ObesityDisorder.SK_PatientID = P.SK_PatientID
	LEFT JOIN  #NAFLD AS NAFLD ON NAFLD.SK_PatientID = P.SK_PatientID
	LEFT JOIN  #Hypercholesterol AS Hypercholesterol ON Hypercholesterol.SK_PatientID = P.SK_PatientID
	LEFT JOIN  #Acanthosis AS Acanthosis ON Acanthosis.SK_PatientID = P.SK_PatientID

	-- RELATED AUTOIMMUNE & ENDOCRINE
	LEFT JOIN  #Thyroid AS Thyroid ON Thyroid.SK_PatientID = P.SK_PatientID
	LEFT JOIN  #Autoimmune AS Autoimmune ON Autoimmune.SK_PatientID = P.SK_PatientID

	--Miscellaneous
	LEFT JOIN  #Frailtyindex AS Frailtyindex ON Frailtyindex.SK_PatientID = P.SK_PatientID
	LEFT JOIN  #NHS_HC AS NHS_HC ON NHS_HC.SK_PatientID = P.SK_PatientID

GO

/*

BEGIN -- Drop all temp tables used

	DECLARE @DropGlobal bit=0 --Default dont drop global temp table

	DECLARE @DROP_STATEMENT nvarchar(1000)
	DECLARE cursorDEL CURSOR FOR
	SELECT 'DROP TABLE '
	   + case
			   when name like '##%' then name
			   when name like '#%' then SUBSTRING(name, 1, CHARINDEX( '____', name)-1)
		end as DropSQL
	from tempdb..sysobjects
	WHERE name LIKE '#%'
	   AND OBJECT_ID('tempdb..' + name) IS NOT NULL
	   AND name not like case
						   when @DropGlobal=0 then '##%' --//Exclude global temp
						   else '#######%'    --//some fack expression so we can
											   --//select global temp for delete
						end
	OPEN cursorDEL
	FETCH NEXT FROM cursorDEL INTO @DROP_STATEMENT
	WHILE @@FETCH_STATUS = 0
	BEGIN
	EXEC (@DROP_STATEMENT)
	print @DROP_STATEMENT
	FETCH NEXT FROM cursorDEL INTO @DROP_STATEMENT
	END
	CLOSE cursorDEL
	DEALLOCATE cursorDEL

END

GO
*/
```

2. Other diagnoses

```
USE CEG

-- drop all temp tables if exist
IF OBJECT_ID('tempdb..#AsthmaQoF', 'U') IS NOT NULL DROP TABLE #AsthmaQoF
IF OBJECT_ID('tempdb..#COPD_QoF', 'U') IS NOT NULL DROP TABLE #COPD_QoF
IF OBJECT_ID('tempdb..#IPF', 'U') IS NOT NULL DROP TABLE #IPF
IF OBJECT_ID('tempdb..#AMD', 'U') IS NOT NULL DROP TABLE #AMD
IF OBJECT_ID('tempdb..#Glaucoma', 'U') IS NOT NULL DROP TABLE #Glaucoma
IF OBJECT_ID('tempdb..#RA', 'U') IS NOT NULL DROP TABLE #RA
IF OBJECT_ID('tempdb..#Lupus', 'U') IS NOT NULL DROP TABLE #Lupus
IF OBJECT_ID('tempdb..#Crohns', 'U') IS NOT NULL DROP TABLE #Crohns
IF OBJECT_ID('tempdb..#UC', 'U') IS NOT NULL DROP TABLE #UC
IF OBJECT_ID('tempdb..#Eczema', 'U') IS NOT NULL DROP TABLE #Eczema
IF OBJECT_ID('tempdb..#PCD', 'U') IS NOT NULL DROP TABLE #PCD

-- EXTRA DIAGNOSES -----------------------------------------------------------------------------------------------------------------------------------
----------- Asthma QOF v37 (QOF definition for asthma excludes asthma resolved codes if appearing after the latest diagnostic code) ------------------

SELECT
	 [SK_PatientID]
	,[ClinicalCode] AsthmaQoFCode
	,[EventDate]	AsthmaDateRecorded

INTO  #AsthmaQoF

FROM
(
	SELECT
		*

	FROM
	(
		SELECT
			 [SK_PatientID]
			,[ClinicalCode]
			,[EventDate]
			,row_number() over(partition by [SK_PatientID] order by [EventDate] ASC) as rn

		FROM
			[08V].[GPEncounter]

		WHERE
			(
				([ClinicalCode] COLLATE Latin1_General_CS_AS LIKE 'H33%' AND [ClinicalCode] COLLATE Latin1_General_CS_AS <> 'H333')
				OR													 -- Synonym
				[ClinicalCode] COLLATE Latin1_General_CS_AS IN('H3120', 'H3120-1', 'H3B', '173A')
			)
			AND
			SK_PatientID IN(SELECT SK_PatientID FROM ceg.GenomicsData WHERE CCG = '08V')

	)as ast

	WHERE
		rn = 1

) AS AsthmaQoF

WHERE
	SK_PatientID NOT IN	-- QOF definition for asthma excludes asthma resolved codes if appearing after the latest diagnostic code
	(
		SELECT
			[SK_PatientID]

		FROM
		(
			SELECT
				 [SK_PatientID]
				,[ClinicalCode]
				,[EventDate]
				,row_number() over(partition by [SK_PatientID] order by [EventDate] DESC) as rn

			FROM
				[08V].[GPEncounter]

			WHERE
				(
					(
						([ClinicalCode] COLLATE Latin1_General_CS_AS LIKE 'H33%' AND [ClinicalCode] COLLATE Latin1_General_CS_AS <> 'H333')
						OR														-- Synonym
						[ClinicalCode] COLLATE Latin1_General_CS_AS IN('H3120', 'H3120-1', 'H3B', '173A')
					)
					OR
					[ClinicalCode] COLLATE Latin1_General_CS_AS IN('21262','212G')  --Asthma resolved
				)
				AND
				SK_PatientID IN(SELECT SK_PatientID FROM ceg.GenomicsData WHERE CCG = '08V')

		) AS  Res

		WHERE
			rn = 1	AND	[ClinicalCode] IN('21262','212G')  --Asthma resolved
	)

GO

-----------COPD QOF v37 (QOF definition for COPD excludes COPD resolved codes if appearing after the latest diagnostic code)--------

SELECT
	 [SK_PatientID]
	,[ClinicalCode] COPDQoFCode
	,[EventDate]	COPD_DateRecorded

INTO  #COPD_QoF

FROM
(
	SELECT
		*

	FROM
	(
		SELECT
			 [SK_PatientID]
			,[ClinicalCode]
			,[EventDate]
			,row_number() over(partition by [SK_PatientID] order by [EventDate] ASC) as rn

		FROM
			[08V].[GPEncounter]

		WHERE
			(
				   ([ClinicalCode] COLLATE Latin1_General_CS_AS LIKE 'H31%' AND [ClinicalCode] COLLATE Latin1_General_CS_AS  NOT IN('H3101', 'H31y0', 'H3122'))
				OR [ClinicalCode] COLLATE Latin1_General_CS_AS LIKE 'H32%'
				OR [ClinicalCode] COLLATE Latin1_General_CS_AS IN
				   (
					 'H3', 'H3-1'	-- Synonym
					,'H36', 'H37', 'H38', 'H39', 'H3A', 'H3B', 'H3y', 'H3y-1', 'H3z', 'H3z-1'	-- Synonym
					,'H4640', 'H4641', 'H5832', 'Hyu30', 'Hyu31'
				   )
			)
			AND
			SK_PatientID IN(SELECT SK_PatientID FROM ceg.GenomicsData WHERE CCG = '08V')

	)as copd

	WHERE
		rn = 1

) AS COPD_QoF

WHERE
	SK_PatientID NOT IN	-- QOF definition for COPD excludes COPD resolved codes if appearing after the latest diagnostic code
	(
		SELECT
			[SK_PatientID]

		FROM
		(
			SELECT
				 [SK_PatientID]
				,[ClinicalCode]
				,[EventDate]
				,row_number() over(partition by [SK_PatientID] order by [EventDate] DESC) as rn

			FROM
				[08V].[GPEncounter]

			WHERE
				(
					   ([ClinicalCode] COLLATE Latin1_General_CS_AS LIKE 'H31%' AND [ClinicalCode] COLLATE Latin1_General_CS_AS  NOT IN('H3101', 'H31y0', 'H3122'))
					OR [ClinicalCode] COLLATE Latin1_General_CS_AS LIKE 'H32%'
					OR [ClinicalCode] COLLATE Latin1_General_CS_AS IN
					   (
						 'H3', 'H3-1'	-- Synonym
						,'H36', 'H37', 'H38', 'H39', 'H3A', 'H3B', 'H3y', 'H3y-1', 'H3z', 'H3z-1'	-- Synonym
						,'H4640', 'H4641', 'H5832', 'Hyu30', 'Hyu31'
					   )
					OR
					[ClinicalCode] COLLATE Latin1_General_CS_AS = '2126F' -- COPD resolved
				)
				AND
				SK_PatientID IN(SELECT SK_PatientID FROM ceg.GenomicsData WHERE CCG = '08V')

		) AS  Res

		WHERE
			rn = 1	AND	[ClinicalCode] = '2126F' -- COPD resolved
	)

GO


-----------Idiopathic Pulmonary Fibrosis -----------------------------------------------------------------------------------

SELECT
	 [SK_PatientID]
	,[ClinicalCode] IPFCode
	,[EventDate]	IPF_DateRecorded

INTO  #IPF

FROM
(
	SELECT
		 [SK_PatientID]
		,[ClinicalCode]
		,[EventDate]
		,row_number() over(partition by [SK_PatientID] order by [EventDate] ASC) as rn

	FROM  [08V].[GPEncounter]
	WHERE		-- 	H5631, H5632
		(
			[ClinicalCode] COLLATE Latin1_General_CS_AS IN ('H5631', 'H5632')
		)
		AND
		SK_PatientID IN(SELECT SK_PatientID FROM ceg.GenomicsData WHERE CCG = '08V')

)as IPF

WHERE rn = 1

GO


-----------Age related macular degeneration -----------------------------------------------------------------------------------

SELECT
	 [SK_PatientID]
	,[ClinicalCode] AMDCode
	,[EventDate]	AMDDateRecorded

INTO  #AMD

FROM
(
	SELECT
		 [SK_PatientID]
		,[ClinicalCode]
		,[EventDate]
		,row_number() over(partition by [SK_PatientID] order by [EventDate] ASC) as rn

	FROM  [08V].[GPEncounter]
	WHERE
		(
			[ClinicalCode] COLLATE Latin1_General_CS_AS IN ('F4250', 'F4251', 'F4252', 'F4252-1')
		)
		AND
		SK_PatientID IN(SELECT SK_PatientID FROM ceg.GenomicsData WHERE CCG = '08V')

) AS AMD

WHERE rn = 1

GO


-----------Glaucoma -----------------------------------------------------------------------------------

SELECT
	 [SK_PatientID]
	,[ClinicalCode] GlaucomaCode
	,[EventDate] GlaucomaDateRecorded

INTO  #Glaucoma

FROM
(
	SELECT
		 [SK_PatientID]
		,[ClinicalCode]
		,[EventDate]
		,row_number() over(partition by [SK_PatientID] order by [EventDate] ASC) as rn

	FROM  [08V].[GPEncounter]
	WHERE
		(
			[ClinicalCode] COLLATE Latin1_General_CS_AS LIKE 'F45%'
		)
		AND
		SK_PatientID IN(SELECT SK_PatientID FROM ceg.GenomicsData WHERE CCG = '08V')

) AS Glaucoma

WHERE rn = 1

GO

-----------Rheumatoid arthritis (QOF v37)-----------------------------------------------------------------------------------

SELECT
	 [SK_PatientID]
	,[ClinicalCode] RACode
	,[EventDate] RADateRecorded

INTO  #RA

FROM
(
	SELECT
		 [SK_PatientID]
		,[ClinicalCode]
		,[EventDate]
		,row_number() over(partition by [SK_PatientID] order by [EventDate] ASC) as rn

	FROM  [08V].[GPEncounter]
	WHERE
		(
				[ClinicalCode] COLLATE Latin1_General_CS_AS LIKE 'N040%'
			OR	([ClinicalCode] COLLATE Latin1_General_CS_AS LIKE 'N042%' AND [ClinicalCode] COLLATE Latin1_General_CS_AS <> 'N0420')
			OR	[ClinicalCode] COLLATE Latin1_General_CS_AS IN ('N041', 'N047', 'N04X',
																'N04y0', 'N04y0-1', 'N04y0-2',
																'N04y2', 'Nyu11', 'Nyu12', 'Nyu1G', 'Nyu10', 'G5yA', 'G5y8')
		)
		AND
		SK_PatientID IN(SELECT SK_PatientID FROM ceg.GenomicsData WHERE CCG = '08V')

)as RA

WHERE rn = 1

GO


-----------Systemic lupus erythematosus (SLE) -----------------------------------------------------------------------------------

SELECT
	 [SK_PatientID]
	,[ClinicalCode] LupusCode
	,[EventDate] LupusDateRecorded

INTO  #Lupus

FROM
(
	SELECT
		 [SK_PatientID]
		,[ClinicalCode]
		,[EventDate]
		,row_number() over(partition by [SK_PatientID] order by [EventDate] ASC) as rn

	FROM  [08V].[GPEncounter]
	WHERE
		(
			[ClinicalCode] COLLATE Latin1_General_CS_AS LIKE 'N000%'
		)
		AND
		SK_PatientID IN(SELECT SK_PatientID FROM ceg.GenomicsData WHERE CCG = '08V')

) AS Lupus

WHERE rn = 1

GO


-----------Crohn's Disease -----------------------------------------------------------------------------------

SELECT
	 [SK_PatientID]
	,[ClinicalCode] CrohnsCode
	,[EventDate] CrohnsDateRecorded

INTO  #Crohns

FROM
(
	SELECT
		 [SK_PatientID]
		,[ClinicalCode]
		,[EventDate]
		,row_number() over(partition by [SK_PatientID] order by [EventDate] ASC) as rn

	FROM  [08V].[GPEncounter]
	WHERE
		(
			[ClinicalCode] COLLATE Latin1_General_CS_AS LIKE 'J40%'
		)
		AND
		SK_PatientID IN(SELECT SK_PatientID FROM ceg.GenomicsData WHERE CCG = '08V')

) AS Crohns

WHERE rn = 1

GO


-----------Ulcerative colitis and/or proctitis -----------------------------------------------------------------------------------

SELECT
	 [SK_PatientID]
	,[ClinicalCode] UC_Code
	,[EventDate] UCDateRecorded

INTO  #UC

FROM
(
	SELECT
		 [SK_PatientID]
		,[ClinicalCode]
		,[EventDate]
		,row_number() over(partition by [SK_PatientID] order by [EventDate] ASC) as rn

	FROM  [08V].[GPEncounter]
	WHERE
		(
			[ClinicalCode] COLLATE Latin1_General_CS_AS LIKE 'J41%'
		)
		AND
		SK_PatientID IN(SELECT SK_PatientID FROM ceg.GenomicsData WHERE CCG = '08V')

) AS UC

WHERE rn = 1

GO

-----------Eczema -----------------------------------------------------------------------------------------------------


SELECT
	 [SK_PatientID]
	,[ClinicalCode] EczemaCode
	,[EventDate] EczemaDateRecorded

INTO  #Eczema

FROM
(
	SELECT
		 [SK_PatientID]
		,[ClinicalCode]
		,[EventDate]
		,row_number() over(partition by [SK_PatientID] order by [EventDate] ASC) as rn

	FROM  [08V].[GPEncounter]
	WHERE
		[ClinicalCode] COLLATE Latin1_General_CS_AS LIKE 'M11%'
		AND
		SK_PatientID IN(SELECT SK_PatientID FROM ceg.GenomicsData WHERE CCG = '08V')

)as Ecz

WHERE rn = 1

GO

-----------Primary ciliary dyskinesia (PCD) --------------------------------------------------------------------------------


SELECT
	 [SK_PatientID]
	,[ClinicalCode] PCDQoFCode
	,[EventDate] PCDDateRecorded

INTO  #PCD

FROM
(
	SELECT
		 [SK_PatientID]
		,[ClinicalCode]
		,[EventDate]
		,row_number() over(partition by [SK_PatientID] order by [EventDate] DESC) as rn

	FROM  [08V].[GPEncounter]
	WHERE
		[ClinicalCode] COLLATE Latin1_General_CS_AS = 'Pyu9D'
		AND
		SK_PatientID IN(SELECT SK_PatientID FROM ceg.GenomicsData WHERE CCG = '08V')

)as pcd

WHERE rn = 1

GO


--------------------------------------------------------------------------------------------------------------------------
---------------------------------------- Joining Tables ------------------------------------------------------------------
--------------------------------------------------------------------------------------------------------------------------


SELECT

	(Select CommissionerName From Dictionary.[dbo].[Commissioner] Where CommissionerCode = '08V') AS Locality
	,P.EncryptedNHSNumber

	,P.SK_PatientID

	--EXTRA DIAGNOSES
	,AsthmaQoF.AsthmaQoFCode
	,AsthmaQoF.AsthmaDateRecorded

	,COPD_QoF.COPDQoFCode
	,COPD_QoF.COPD_DateRecorded

	,IPF.IPFCode
	,IPF.IPF_DateRecorded

	,AMD.AMDCode
	,AMD.AMDDateRecorded

	,Glaucoma.GlaucomaCode
	,Glaucoma.GlaucomaDateRecorded

	,RA.RACode
	,RA.RADateRecorded

	,Lupus.LupusCode
	,Lupus.LupusDateRecorded

	,Crohns.CrohnsCode
	,Crohns.CrohnsDateRecorded

	,UC.UC_Code
	,UC.UCDateRecorded

	,Eczema.EczemaCode
	,Eczema.EczemaDateRecorded

	,PCD.PCDQoFCode
	,PCD.PCDDateRecorded

FROM
	ceg.GenomicsData AS P

	--  EXTRA DIAGNOSES
	LEFT JOIN #AsthmaQoF AS AsthmaQoF ON AsthmaQoF.SK_PatientID = P.SK_PatientID
	LEFT JOIN #COPD_QoF AS COPD_QoF ON COPD_QoF.SK_PatientID = P.SK_PatientID
	LEFT JOIN #IPF AS IPF ON IPF.SK_PatientID = P.SK_PatientID
	LEFT JOIN #AMD AS AMD ON AMD.SK_PatientID = P.SK_PatientID
	LEFT JOIN #Glaucoma AS Glaucoma ON Glaucoma.SK_PatientID = P.SK_PatientID
	LEFT JOIN #RA AS RA ON RA.SK_PatientID = P.SK_PatientID
	LEFT JOIN #Lupus AS Lupus ON Lupus.SK_PatientID = P.SK_PatientID
	LEFT JOIN #Crohns AS Crohns ON Crohns.SK_PatientID = P.SK_PatientID
	LEFT JOIN #UC AS UC ON UC.SK_PatientID = P.SK_PatientID
	LEFT JOIN  #Eczema AS Eczema ON Eczema.SK_PatientID = P.SK_PatientID
	LEFT JOIN  #PCD AS PCD ON PCD.SK_PatientID = P.SK_PatientID

WHERE
	P.CCG = '08V'

GO


-- drop all temp tables used
DROP TABLE #AsthmaQoF
DROP TABLE #COPD_QoF
DROP TABLE #IPF
DROP TABLE #AMD
DROP TABLE #Glaucoma
DROP TABLE #RA
DROP TABLE #Lupus
DROP TABLE #Crohns
DROP TABLE #UC
DROP TABLE #Eczema
DROP TABLE #PCD

GO
```

3. Neuro and mental health

```
USE CEG

-- drop all temp tables if exist
BEGIN -- drop all temp tables if exist

	DECLARE @DropGlobal bit=0 --Default dont drop global temp table

	DECLARE @DROP_STATEMENT nvarchar(1000)
	DECLARE cursorDEL CURSOR FOR
	SELECT 'DROP TABLE '
	   + case
			   when name like '##%' then name
			   when name like '#%' then SUBSTRING(name, 1, CHARINDEX( '____', name)-1)
		end as DropSQL
	from tempdb..sysobjects
	WHERE name LIKE '#%'
	   AND OBJECT_ID('tempdb..' + name) IS NOT NULL
	   AND name not like case
						   when @DropGlobal=0 then '##%' --//Exclude global temp
						   else '#######%'    --//some fack expression so we can
											   --//select global temp for delete
						end
	OPEN cursorDEL
	FETCH NEXT FROM cursorDEL INTO @DROP_STATEMENT
	WHILE @@FETCH_STATUS = 0
	BEGIN
	EXEC (@DROP_STATEMENT)
	--print @DROP_STATEMENT
	FETCH NEXT FROM cursorDEL INTO @DROP_STATEMENT
	END
	CLOSE cursorDEL
	DEALLOCATE cursorDEL

END

GO


-- NEURO & MENTAL HEALTH -----------------------------------------------------------------------------------------------------------------------------------
-----------Amyotrophic lateral sclerosis (motor neurone disease) -----------------------------------------------------------------------------------

SELECT
	 [SK_PatientID]
	,[ClinicalCode] MNDCode
	,[EventDate] MND_DateRecorded

INTO  #MND

FROM
(
	SELECT
		 [SK_PatientID]
		,[ClinicalCode]
		,[EventDate]
		,row_number() over(partition by [SK_PatientID] order by [EventDate] ASC) as rn

	FROM  [08V].[GPEncounter]
	WHERE
		(
			[ClinicalCode] COLLATE Latin1_General_CS_AS LIKE 'F152%'
		)
		AND
		SK_PatientID IN(SELECT SK_PatientID FROM ceg.GenomicsData WHERE CCG = '08V')

) AS MND

WHERE rn = 1

GO


-----------Multiple sclerosis -----------------------------------------------------------------------------------

SELECT
	 [SK_PatientID]
	,[ClinicalCode] MSCode
	,[EventDate] MSDateRecorded

INTO  #MS

FROM
(
	SELECT
		 [SK_PatientID]
		,[ClinicalCode]
		,[EventDate]
		,row_number() over(partition by [SK_PatientID] order by [EventDate] ASC) as rn

	FROM  [08V].[GPEncounter]
	WHERE
		(
			[ClinicalCode] COLLATE Latin1_General_CS_AS LIKE 'F20%'
		)
		AND
		SK_PatientID IN(SELECT SK_PatientID FROM ceg.GenomicsData WHERE CCG = '08V')

) AS MS

WHERE rn = 1

GO


-----------Parkinson's Disease -----------------------------------------------------------------------------------


SELECT
	 [SK_PatientID]
	,[ClinicalCode] ParkinsonsCode
	,[EventDate] ParkinsonsDateRecorded

INTO  #Parkinsons

FROM
(
	SELECT
		 [SK_PatientID]
		,[ClinicalCode]
		,[EventDate]
		,row_number() over(partition by [SK_PatientID] order by [EventDate] ASC) as rn

	FROM  [08V].[GPEncounter]

	WHERE		--F12%, F1303, F11x9,147F
		(
			[ClinicalCode] COLLATE Latin1_General_CS_AS LIKE 'F12%'
			OR
			[ClinicalCode] COLLATE Latin1_General_CS_AS IN('F1303', 'F11x9', '147F')
		)
		AND
		SK_PatientID IN(SELECT SK_PatientID FROM ceg.GenomicsData WHERE CCG = '08V')

) AS Parkinsons

WHERE rn = 1

GO


-----------Atypical Parkinson's -----------------------------------------------------------------------------------


SELECT
	 [SK_PatientID]
	,[ClinicalCode] AtypicalPD_Code
	,[EventDate] AtypicalPD_DateRecorded

INTO #AtypicalPD

FROM
(
	SELECT
		 [SK_PatientID]
		,[ClinicalCode]
		,[EventDate]
		,row_number() over(partition by [SK_PatientID] order by [EventDate] ASC) as rn

	FROM  [08V].[GPEncounter]
	WHERE
		(
			[ClinicalCode] COLLATE Latin1_General_CS_AS IN('F24y0', 'F24y0-1', 'F24y0-2') -- cover Synonym
			OR
			[ClinicalCode] COLLATE Latin1_General_CS_AS IN('F24y2', 'F11y2', 'F174')
		)
		AND
		SK_PatientID IN(SELECT SK_PatientID FROM ceg.GenomicsData WHERE CCG = '08V')

) AS AtypicalPD

WHERE rn = 1

GO


-----------Vascular Dementia -----------------------------------------------------------------------------------


SELECT
	 [SK_PatientID]
	,[ClinicalCode] VasDemCode
	,[EventDate] VasDemDateRecorded

INTO  #VasDem

FROM
(
	SELECT
		 [SK_PatientID]
		,[ClinicalCode]
		,[EventDate]
		,row_number() over(partition by [SK_PatientID] order by [EventDate] ASC) as rn

	FROM  [08V].[GPEncounter]
	WHERE
		(
			[ClinicalCode] COLLATE Latin1_General_CS_AS LIKE 'Eu01%'
		)
		AND
		SK_PatientID IN(SELECT SK_PatientID FROM ceg.GenomicsData WHERE CCG = '08V')

) AS VasDem

WHERE rn = 1

GO


-----------Alzheimer's Disease -----------------------------------------------------------------------------------

SELECT
	 [SK_PatientID]
	,[ClinicalCode] AlzheimersCode
	,[EventDate] AlzheimersDateRecorded

INTO  #Alzheimers

FROM
(
	SELECT
		 [SK_PatientID]
		,[ClinicalCode]
		,[EventDate]
		,row_number() over(partition by [SK_PatientID] order by [EventDate] ASC) as rn

	FROM  [08V].[GPEncounter]
	WHERE
		(
			[ClinicalCode] COLLATE Latin1_General_CS_AS LIKE 'F110%'
		)
		AND
		SK_PatientID IN(SELECT SK_PatientID FROM ceg.GenomicsData WHERE CCG = '08V')

) AS Alzheimers

WHERE rn = 1

GO


-----------Dementia (QOF v37)-----------------------------------------------------------------------------------

SELECT
	 [SK_PatientID]
	,[ClinicalCode] DementiaCode
	,[EventDate] DementiaDateRecorded

INTO  #DementiaQoF

FROM
(
	SELECT
		 [SK_PatientID]
		,[ClinicalCode]
		,[EventDate]
		,row_number() over(partition by [SK_PatientID] order by [EventDate] ASC) as rn

	FROM  [08V].[GPEncounter]
	WHERE
		(
				[ClinicalCode] COLLATE Latin1_General_CS_AS LIKE 'A411%'
			OR	[ClinicalCode] COLLATE Latin1_General_CS_AS LIKE 'E00%'
			OR	[ClinicalCode] COLLATE Latin1_General_CS_AS LIKE 'Eu00%'
			OR	[ClinicalCode] COLLATE Latin1_General_CS_AS LIKE 'Eu01%'
			OR	[ClinicalCode] COLLATE Latin1_General_CS_AS LIKE 'Eu02%'
			OR	[ClinicalCode] COLLATE Latin1_General_CS_AS LIKE 'E012%'
			OR	[ClinicalCode] COLLATE Latin1_General_CS_AS IN ('E02y1', 'E041', 'Eu041', 'F110', 'F111', 'F112', 'F116', 'F118', 'F21y2', 'A410', 'Eu107', 'Eu107-1', 'Eu107-2', 'F11x7')
		)
		AND
		SK_PatientID IN(SELECT SK_PatientID FROM ceg.GenomicsData WHERE CCG = '08V')

)as Dementia

WHERE rn = 1

GO

-----------Cognitive decline --------------------------------------------------------------------------------------

SELECT
	 [SK_PatientID]
	,[ClinicalCode] CognitiveCode
	,[EventDate]	CognitiveDateRecorded

INTO  #Cognitive

FROM
(
	SELECT
		 [SK_PatientID]
		,[ClinicalCode]
		,[EventDate]
		,row_number() over(partition by [SK_PatientID] order by [EventDate] ASC) as rn

	FROM  [08V].[GPEncounter]
	WHERE
		(		-- 28E%, 3AE1 , 3AE2 , Eu057, Ryu5 , Ryu51, Eu05%
				[ClinicalCode] COLLATE Latin1_General_CS_AS LIKE '28E%'
			OR	[ClinicalCode] COLLATE Latin1_General_CS_AS LIKE 'Eu05%'
			OR	[ClinicalCode] COLLATE Latin1_General_CS_AS IN ('3AE1', '3AE2', 'Eu057', 'Ryu5', 'Ryu51')
		)
		AND
		SK_PatientID IN(SELECT SK_PatientID FROM ceg.GenomicsData WHERE CCG = '08V')

)as Cognitive

WHERE rn = 1

GO

-----------Memory disorders --------------------------------------------------------------------------------------

SELECT
	 [SK_PatientID]
	,[ClinicalCode] MemoryCode
	,[EventDate]	MemoryDateRecorded

INTO  #Memory

FROM
(
	SELECT
		 [SK_PatientID]
		,[ClinicalCode]
		,[EventDate]
		,row_number() over(partition by [SK_PatientID] order by [EventDate] ASC) as rn

	FROM  [08V].[GPEncounter]
	WHERE
		[ClinicalCode] COLLATE Latin1_General_CS_AS IN
		(
			 '1B1A', '1B1A-1', '1B1A-2', '1B1A-3'		-- Synonym (-)
			,'1B1Y', '1B1a', '1S21','28G'
			,'3A10', '3A20', '3A30', '3A40', '3A50', '3A60', '3A70', '3A80', '3A91', '3AA1'
			,'8BIk', '8HTY','9Nk1'
			,'E2A10', 'E2A11'
			,'R00z0', 'R00z0-1'
		)
		AND
		SK_PatientID IN(SELECT SK_PatientID FROM ceg.GenomicsData WHERE CCG = '08V')

)as Memory

WHERE rn = 1

GO

-----------Brain tumours -----------------------------------------------------------------------------------

SELECT
	 [SK_PatientID]
	,[ClinicalCode] BrainTumourCode
	,[EventDate]	BrainTumourDateRecorded

INTO  #BrainTumour

FROM
(
	SELECT
		 [SK_PatientID]
		,[ClinicalCode]
		,[EventDate]
		,row_number() over(partition by [SK_PatientID] order by [EventDate] ASC) as rn

	FROM  [08V].[GPEncounter]
	WHERE
		(		--B7F0, B51%, 2B2%, B583%
				[ClinicalCode] COLLATE Latin1_General_CS_AS LIKE '2B2%'
			OR	[ClinicalCode] COLLATE Latin1_General_CS_AS LIKE 'B51%'
			OR	[ClinicalCode] COLLATE Latin1_General_CS_AS LIKE 'B583%'
			OR	[ClinicalCode] COLLATE Latin1_General_CS_AS IN ('B7F0', 'B7F0-1')
		)
		AND
		SK_PatientID IN(SELECT SK_PatientID FROM ceg.GenomicsData WHERE CCG = '08V')

) AS BrainTumour

WHERE rn = 1

GO

-----------Brain damage childhood -----------------------------------------------------------------------------------

SELECT
	 [SK_PatientID]
	,[ClinicalCode] BrainDamageCode
	,[EventDate]	BrainDamageDateRecorded

INTO  #BrainDamage

FROM
(
	SELECT
		 [SK_PatientID]
		,[ClinicalCode]
		,[EventDate]
		,row_number() over(partition by [SK_PatientID] order by [EventDate] ASC) as rn

	FROM  [08V].[GPEncounter]
	WHERE
		(
			[ClinicalCode] COLLATE Latin1_General_CS_AS IN
			(
				'Q200z', 'Q200z-1',								-- Synonym (-)
				'129', '129-1', '129-2',						-- Synonym (-)
				'Q2004', 'Q2004-1',								-- Synonym (-)
				'F23', 'F23-1', 'F23-2', 'F23-3', 'F23-4',		-- Synonym (-)
				'PJ0', 'PJ0-1', 'PJ0-2', 'PJ0-3',				-- Synonym (-)
				'P22', 'Q200', '13Z4E', 'Eu81', 'PJ'
			)


		)
		AND
		SK_PatientID IN(SELECT SK_PatientID FROM ceg.GenomicsData WHERE CCG = '08V')

) AS BrainDamage

WHERE rn = 1

GO


-----------Epilepsy  -----------------------------------------------------------------------------------

SELECT
	 [SK_PatientID]
	,[ClinicalCode] EpilepsyCode
	,[EventDate]	EpilepsyDateRecorded

INTO  #Epilepsy

FROM
(
	SELECT
		 [SK_PatientID]
		,[ClinicalCode]
		,[EventDate]
		,row_number() over(partition by [SK_PatientID] order by [EventDate] ASC) as rn

	FROM  [08V].[GPEncounter]
	WHERE
		(
			[ClinicalCode] COLLATE Latin1_General_CS_AS LIKE 'F250%'
			OR
			[ClinicalCode] COLLATE Latin1_General_CS_AS LIKE 'F251%'
			OR
			[ClinicalCode] COLLATE Latin1_General_CS_AS LIKE 'F254%'
			OR
			[ClinicalCode] COLLATE Latin1_General_CS_AS LIKE 'F255%'
			OR
			[ClinicalCode] COLLATE Latin1_General_CS_AS LIKE 'F25y%'
			OR
			[ClinicalCode] COLLATE Latin1_General_CS_AS IN('F253', 'F253-1')	-- Synonym (-)
			OR
			[ClinicalCode] COLLATE Latin1_General_CS_AS IN('F25z', 'F25z-1')	-- Synonym (-)
			OR
			[ClinicalCode] COLLATE Latin1_General_CS_AS IN('F1321', 'F1321-1')	-- Synonym (-)
			OR
			[ClinicalCode] COLLATE Latin1_General_CS_AS IN ('F25', 'F252', 'F257', 'F25B', 'F25C', 'F25D', 'F25E', 'F25F', 'F25X', 'SC200')
		)
		AND
		SK_PatientID IN(SELECT SK_PatientID FROM ceg.GenomicsData WHERE CCG = '08V')

)as Epilepsy

WHERE rn = 1

GO

----------Schizophrenic disorders (QOF v37) --------------------------------------------------

SELECT
	 [SK_PatientID]
	,[ClinicalCode] Schizophrenic_QoFCode
	,[EventDate]	Schizophrenic_QoFDateRecorded

INTO  #SchizophrenicQof

FROM
(
	SELECT
		 [SK_PatientID]
		,[ClinicalCode]
		,[EventDate]
		,row_number() over(partition by [SK_PatientID] order by [EventDate] ASC) as rn

	FROM  [08V].[GPEncounter]

	WHERE
		(	--ELGH v8:   E10%,  E12% , E13%, E2122, Eu2%
			   [ClinicalCode] COLLATE Latin1_General_CS_AS LIKE 'E10%'
			OR [ClinicalCode] COLLATE Latin1_General_CS_AS LIKE 'E12%'
			OR [ClinicalCode] COLLATE Latin1_General_CS_AS LIKE 'E13%'
			OR [ClinicalCode] COLLATE Latin1_General_CS_AS LIKE 'Eu2%'
			OR [ClinicalCode] COLLATE Latin1_General_CS_AS = 'E2122'
		)
		AND
		SK_PatientID IN(SELECT SK_PatientID FROM ceg.GenomicsData WHERE CCG = '08V')

)as Schizophrenic

WHERE rn = 1

GO


----------Bipolar, manic disorders (QOF v37) --------------------------------------------------

SELECT
	 [SK_PatientID]
	,[ClinicalCode] Bipolar_QoFCode
	,[EventDate] Bipolar_QoFDateRecorded

INTO  #Bipolar_QoF

FROM
(
	SELECT
		 [SK_PatientID]
		,[ClinicalCode]
		,[EventDate]
		,row_number() over(partition by [SK_PatientID] order by [EventDate] ASC) as rn

	FROM  [08V].[GPEncounter]

	WHERE
		(	--ELGH v8:   E110%, E111%, E1124, E1134, E114%, E115%, E116%, E117%, E11y%, E11z, E11z0, E11zz, Eu30% , Eu31% , Eu323, Eu328, Eu333, Eu329, Eu32A
			   [ClinicalCode] COLLATE Latin1_General_CS_AS LIKE 'E110%'
			OR [ClinicalCode] COLLATE Latin1_General_CS_AS LIKE 'E111%'
			OR [ClinicalCode] COLLATE Latin1_General_CS_AS LIKE 'E114%'
			OR [ClinicalCode] COLLATE Latin1_General_CS_AS LIKE 'E115%'
			OR [ClinicalCode] COLLATE Latin1_General_CS_AS LIKE 'E116%'
			OR [ClinicalCode] COLLATE Latin1_General_CS_AS LIKE 'E117%'
			OR [ClinicalCode] COLLATE Latin1_General_CS_AS LIKE 'E11y%'
			OR [ClinicalCode] COLLATE Latin1_General_CS_AS LIKE 'Eu30%'
			OR [ClinicalCode] COLLATE Latin1_General_CS_AS LIKE 'Eu31%'
			OR [ClinicalCode] COLLATE Latin1_General_CS_AS LIKE 'Eu333%'	-- % to cover Synonym
			OR
			   [ClinicalCode] COLLATE Latin1_General_CS_AS IN ('Eu32', 'Eu32-1', 'Eu32-2', 'Eu32-3')	-- Synonym (-)
			OR
			   [ClinicalCode] COLLATE Latin1_General_CS_AS IN ('Eu323', 'Eu323-1', 'Eu323-2', 'Eu323-3', 'Eu323-4')	-- Synonym (-)
			OR
			   [ClinicalCode] COLLATE Latin1_General_CS_AS IN ('E1124', 'E1134', 'E11z', 'E11z0', 'E11zz', 'Eu328', 'Eu329')
		)
		AND
		SK_PatientID IN(SELECT SK_PatientID FROM ceg.GenomicsData WHERE CCG = '08V')

)as Bipolar

WHERE rn = 1

GO


------------------Depression (QOF) ------------------------------------------------------------------------

SELECT
	 [SK_PatientID]
	,[ClinicalCode] DepressionQoFCode
	,[EventDate] DepressionQoFDateRecorded

INTO #DepressionQoF

FROM
(
	SELECT
		 [SK_PatientID]
		,[ClinicalCode]
		,[EventDate]
		,row_number() over(partition by [SK_PatientID] order by [EventDate] ASC) as rn

	FROM  [08V].[GPEncounter]
	WHERE
			--ELGH v8:   E0013, E0021, E112%, E113%, E118, E11y2, E11z2, E130, E135, E2003, E291, E2B, E2B1, Eu204, Eu251, Eu32%, Eu33%, Eu341, Eu412, Eu530, Eu530-1, Eu530-2
		(
			[ClinicalCode] COLLATE Latin1_General_CS_AS LIKE 'E112%'
			OR
			[ClinicalCode] COLLATE Latin1_General_CS_AS LIKE 'E113%'
			OR
			[ClinicalCode] COLLATE Latin1_General_CS_AS LIKE 'Eu32%'
			OR
			[ClinicalCode] COLLATE Latin1_General_CS_AS LIKE 'Eu33%'
			OR
			[ClinicalCode] COLLATE Latin1_General_CS_AS IN('E130', 'E130-1')
			OR
			[ClinicalCode] COLLATE Latin1_General_CS_AS IN('Eu251', 'Eu251-1', 'Eu251-2')
			OR
			[ClinicalCode] COLLATE Latin1_General_CS_AS IN('Eu341', 'Eu341-2', 'Eu341-3', 'Eu341-4', 'Eu341-4')
			OR
			[ClinicalCode] COLLATE Latin1_General_CS_AS IN('Eu412', 'Eu412-1')
			OR
			[ClinicalCode] COLLATE Latin1_General_CS_AS IN('Eu530', 'Eu530-1', 'Eu530-2')
			OR
			[ClinicalCode] COLLATE Latin1_General_CS_AS IN('E0013', 'E0021', 'E118', 'E11y2', 'E11z2', 'E135', 'E2003', 'E291', 'E2B', 'E2B1', 'Eu204')
		)
		-- AND
		-- [ClinicalCode] COLLATE Latin1_General_CS_AS NOT IN ('Eu32A', 'Eu32B', 'Eu329')
		-- [ClinicalCode] COLLATE Latin1_General_CS_AS = '212S'	--Depression Resolved
		AND
		SK_PatientID IN(SELECT SK_PatientID FROM ceg.GenomicsData WHERE CCG = '08V')

)as DepressionQof

WHERE rn = 1

GO

-- ANXIETY DISORDERS ------------------------------------------------------------------------------------------------
----------- Anxiety -------------------------------------------------------------------------------------------------

SELECT
	 [SK_PatientID]
	,[ClinicalCode] AnxietyCode
	,[EventDate]	AnxietyDateRecorded

INTO  #Anxiety

FROM
(
	SELECT
		*

	FROM
	(
		SELECT
			 [SK_PatientID]
			,[ClinicalCode]
			,[EventDate]
			,row_number() over(partition by [SK_PatientID] order by [EventDate] ASC) as rn

		FROM  [08V].[GPEncounter]
		WHERE
			(
				[ClinicalCode] COLLATE Latin1_General_CS_AS LIKE 'E200%'
				OR
				[ClinicalCode] COLLATE Latin1_General_CS_AS LIKE 'Eu40%'
				OR
				[ClinicalCode] COLLATE Latin1_General_CS_AS LIKE 'EU41%'
				OR
				[ClinicalCode] COLLATE Latin1_General_CS_AS LIKE 'Eu054%'
				OR
				[ClinicalCode] COLLATE Latin1_General_CS_AS IN('1466', 'Eu341', 'Eu341-1', 'Eu341-2', 'Eu341-3', 'Eu341-4')	 -- Synonym(-)
			)
			AND
			SK_PatientID IN(SELECT SK_PatientID FROM ceg.GenomicsData WHERE CCG = '08V')

	)as Anxty

	WHERE rn = 1

)as Anxiety

WHERE
	SK_PatientID NOT IN	-- Excludes anxiety resolved codes if appearing after the latest anxiety diagnostic code
	(
		SELECT
			[SK_PatientID]

		FROM
		(
			SELECT
				 [SK_PatientID]
				,[ClinicalCode]
				,[EventDate]
				,row_number() over(partition by [SK_PatientID] order by [EventDate] DESC) as rn

			FROM
				[08V].[GPEncounter]

			WHERE
				(
					[ClinicalCode] COLLATE Latin1_General_CS_AS LIKE 'E200%'
					OR
					[ClinicalCode] COLLATE Latin1_General_CS_AS LIKE 'Eu40%'
					OR
					[ClinicalCode] COLLATE Latin1_General_CS_AS LIKE 'EU41%'
					OR
					[ClinicalCode] COLLATE Latin1_General_CS_AS LIKE 'Eu054%'
					OR
					[ClinicalCode] COLLATE Latin1_General_CS_AS IN('1466', 'Eu341', 'Eu341-1', 'Eu341-2', 'Eu341-3', 'Eu341-4')	 -- Synonym(-)
					OR
					[ClinicalCode] COLLATE Latin1_General_CS_AS = '2126J'  --2126J	Anxiety resolved
				)
				AND
				SK_PatientID IN(SELECT SK_PatientID FROM ceg.GenomicsData WHERE CCG = '08V')

		) AS  Res

		WHERE
			rn = 1	AND	[ClinicalCode] = '2126J'	--2126J	Anxiety resolved
	)

GO

----------- PTSD -------------------------------------------------------------------------------------------------

SELECT
	 [SK_PatientID]
	,[ClinicalCode] PTSDCode
	,[EventDate]	PTSDDateRecorded

INTO  #PTSD

FROM
(
	SELECT
		 [SK_PatientID]
		,[ClinicalCode]
		,[EventDate]
		,row_number() over(partition by [SK_PatientID] order by [EventDate] ASC) as rn

	FROM  [08V].[GPEncounter]
	WHERE
		[ClinicalCode] COLLATE Latin1_General_CS_AS LIKE 'Eu431%'
		AND
		SK_PatientID IN(SELECT SK_PatientID FROM ceg.GenomicsData WHERE CCG = '08V')

)as PTSD

WHERE rn = 1

GO


----------- OCD -------------------------------------------------------------------------------------------------

SELECT
	 [SK_PatientID]
	,[ClinicalCode] OCDCode
	,[EventDate]	OCDDateRecorded

INTO  #OCD

FROM
(
	SELECT
		 [SK_PatientID]
		,[ClinicalCode]
		,[EventDate]
		,row_number() over(partition by [SK_PatientID] order by [EventDate] ASC) as rn

	FROM  [08V].[GPEncounter]
	WHERE
		(
			[ClinicalCode] COLLATE Latin1_General_CS_AS LIKE 'Eu605%'      -- % to cover Synonym
			OR
			[ClinicalCode] COLLATE Latin1_General_CS_AS LIKE 'Eu42%'
		)
		AND
		SK_PatientID IN(SELECT SK_PatientID FROM ceg.GenomicsData WHERE CCG = '08V')

)as OCD

WHERE rn = 1

GO

----------- Panic disorder/panic attack  -------------------------------------------------------------------------------------

SELECT
	 [SK_PatientID]
	,[ClinicalCode] PanicDisorderCode
	,[EventDate]	PanicDisorderDateRecorded

INTO  #PanicDisorder

FROM
(
	SELECT
		 [SK_PatientID]
		,[ClinicalCode]
		,[EventDate]
		,row_number() over(partition by [SK_PatientID] order by [EventDate] ASC) as rn

	FROM  [08V].[GPEncounter]
	WHERE
		(
			[ClinicalCode] COLLATE Latin1_General_CS_AS LIKE 'E2001%'      -- % to cover Synonym
			OR
			[ClinicalCode] COLLATE Latin1_General_CS_AS LIKE 'E202%'
		)
		AND
		SK_PatientID IN(SELECT SK_PatientID FROM ceg.GenomicsData WHERE CCG = '08V')

)as PanicDisorder

WHERE rn = 1

GO


----------- Social anxiety disorder (childhood)  -------------------------------------------------------------------------------------------------

SELECT
	 [SK_PatientID]
	,[ClinicalCode] SocialAnxietyCode
	,[EventDate]	SocialAnxietyDateRecorded

INTO  #SocialAnxiety

FROM
(
	SELECT
		 [SK_PatientID]
		,[ClinicalCode]
		,[EventDate]
		,row_number() over(partition by [SK_PatientID] order by [EventDate] ASC) as rn

	FROM  [08V].[GPEncounter]
	WHERE
		[ClinicalCode] COLLATE Latin1_General_CS_AS LIKE 'Eu932%'      -- % to cover Synonym
		AND
		SK_PatientID IN(SELECT SK_PatientID FROM ceg.GenomicsData WHERE CCG = '08V')

)as SocialAnxiety

WHERE rn = 1

GO

----------- Phobic disorders -------------------------------------------------------------------------------------------------

SELECT
	 [SK_PatientID]
	,[ClinicalCode] PhobiasCode
	,[EventDate]	PhobiasDateRecorded

INTO  #Phobias

FROM
(
	SELECT
		 [SK_PatientID]
		,[ClinicalCode]
		,[EventDate]
		,row_number() over(partition by [SK_PatientID] order by [EventDate] ASC) as rn

	FROM  [08V].[GPEncounter]
	WHERE
		(
			[ClinicalCode] COLLATE Latin1_General_CS_AS = 'Eu931'
			OR
			[ClinicalCode] COLLATE Latin1_General_CS_AS LIKE 'E202%'
			OR
			[ClinicalCode] COLLATE Latin1_General_CS_AS LIKE 'Eu40%'
		)
		AND
		SK_PatientID IN(SELECT SK_PatientID FROM ceg.GenomicsData WHERE CCG = '08V')

)as Phobias

WHERE rn = 1

GO

----------- Seasonal Affective Disorder -------------------------------------------------------------------------------------------------

SELECT
	 [SK_PatientID]
	,[ClinicalCode] SAD_Code
	,[EventDate]	SAD_DateRecorded

INTO  #SAD

FROM
(
	SELECT
		 [SK_PatientID]
		,[ClinicalCode]
		,[EventDate]
		,row_number() over(partition by [SK_PatientID] order by [EventDate] ASC) as rn

	FROM  [08V].[GPEncounter]
	WHERE
		(
			[ClinicalCode] COLLATE Latin1_General_CS_AS = 'E118'
			OR
			[ClinicalCode] COLLATE Latin1_General_CS_AS LIKE 'Eu33%'
		)
		AND
		SK_PatientID IN(SELECT SK_PatientID FROM ceg.GenomicsData WHERE CCG = '08V')

)as SAD

WHERE rn = 1

GO


----------- Dissociative disorder/conversion disorder -------------------------------------------------------------------------------------------------

SELECT
	 [SK_PatientID]
	,[ClinicalCode] ConversionCode
	,[EventDate]	ConversionDateRecorded

INTO  #Conversion

FROM
(
	SELECT
		 [SK_PatientID]
		,[ClinicalCode]
		,[EventDate]
		,row_number() over(partition by [SK_PatientID] order by [EventDate] ASC) as rn

	FROM  [08V].[GPEncounter]
	WHERE
		(

			[ClinicalCode] COLLATE Latin1_General_CS_AS LIKE 'E2016%'      -- % to cover Synonym
			OR
			[ClinicalCode] COLLATE Latin1_General_CS_AS LIKE 'Eu44%'
			OR
			[ClinicalCode] COLLATE Latin1_General_CS_AS = 'Eu055'
		)
		AND
		SK_PatientID IN(SELECT SK_PatientID FROM ceg.GenomicsData WHERE CCG = '08V')

)as Conversion

WHERE rn = 1

GO

----------- Adjustment disorder (includes acute stress reaction) ------------------------------------------------------------------------------------------

SELECT
	 [SK_PatientID]
	,[ClinicalCode] AdjustmentCode
	,[EventDate]	AdjustmentDateRecorded

INTO  #Adjustment

FROM
(
	SELECT
		 [SK_PatientID]
		,[ClinicalCode]
		,[EventDate]
		,row_number() over(partition by [SK_PatientID] order by [EventDate] ASC) as rn

	FROM  [08V].[GPEncounter]
	WHERE
		(
			[ClinicalCode] COLLATE Latin1_General_CS_AS LIKE 'E29%'
			OR
			[ClinicalCode] COLLATE Latin1_General_CS_AS LIKE 'Eu43%'
		)
		AND
		SK_PatientID IN(SELECT SK_PatientID FROM ceg.GenomicsData WHERE CCG = '08V')

)as Adjustment

WHERE rn = 1

GO

----------- Somatoform disorder/hypochondria -------------------------------------------------------------------------------------------------

SELECT
	 [SK_PatientID]
	,[ClinicalCode] SomatoformCode
	,[EventDate]	SomatoformDateRecorded

INTO  #Somatoform

FROM
(
	SELECT
		 [SK_PatientID]
		,[ClinicalCode]
		,[EventDate]
		,row_number() over(partition by [SK_PatientID] order by [EventDate] ASC) as rn

	FROM  [08V].[GPEncounter]
	WHERE
		[ClinicalCode] COLLATE Latin1_General_CS_AS LIKE 'Eu45%'
		AND
		SK_PatientID IN(SELECT SK_PatientID FROM ceg.GenomicsData WHERE CCG = '08V')

)as Somatoform

WHERE rn = 1

GO

----------- Eating disorder -------------------------------------------------------------------------------------------------

SELECT
	 [SK_PatientID]
	,[ClinicalCode] EatingDisorderCode
	,[EventDate]	EatingDisorderDateRecorded

INTO  #EatingDisorder

FROM
(
	SELECT
		 [SK_PatientID]
		,[ClinicalCode]
		,[EventDate]
		,row_number() over(partition by [SK_PatientID] order by [EventDate] ASC) as rn

	FROM  [08V].[GPEncounter]
	WHERE
		(	-- Eu50%, Eu275%, 1JZ
			[ClinicalCode] COLLATE Latin1_General_CS_AS = '1JZ'
			OR
			[ClinicalCode] COLLATE Latin1_General_CS_AS LIKE 'Eu275%'
			OR
			[ClinicalCode] COLLATE Latin1_General_CS_AS LIKE 'Eu50%'
		)
		AND
		SK_PatientID IN(SELECT SK_PatientID FROM ceg.GenomicsData WHERE CCG = '08V')

)as EatingDisorder

WHERE rn = 1

GO

----------- Alcohol/substance/psychoactive/opiate use/abuse/dependence --------------------------------------------------------

SELECT
	 [SK_PatientID]
	,[ClinicalCode] DependenceAndAbuseCode
	,[EventDate]	DependenceAndAbuseDateRecorded

INTO  #DependenceAndAbuse

FROM
(
	SELECT
		*

	FROM
	(
		SELECT
			 [SK_PatientID]
			,[ClinicalCode]
			,[EventDate]
			,row_number() over(partition by [SK_PatientID] order by [EventDate] ASC) as rn

		FROM  [08V].[GPEncounter]
		WHERE
			(	-- E23%, E24%, 13cM%, 1T%, Eu55, ZV114, 13cH, 146F.,  13c1, Eu112, Eu102, Eu182, Eu142, Eu122, Eu152, Eu162, Eu132

				[ClinicalCode] COLLATE Latin1_General_CS_AS LIKE 'E23%'
				OR
				[ClinicalCode] COLLATE Latin1_General_CS_AS LIKE 'E24%'
				OR
				[ClinicalCode] COLLATE Latin1_General_CS_AS LIKE '13cM%'
				OR
				[ClinicalCode] COLLATE Latin1_General_CS_AS LIKE '1T%'
				OR
				[ClinicalCode] COLLATE Latin1_General_CS_AS LIKE 'Eu55%'       -- % to cover Synonym
				OR
				[ClinicalCode] COLLATE Latin1_General_CS_AS LIKE 'Eu112%'      -- % to cover Synonym
				OR
				[ClinicalCode] COLLATE Latin1_General_CS_AS LIKE 'Eu102%'      -- % to cover Synonym
				OR
				[ClinicalCode] COLLATE Latin1_General_CS_AS LIKE 'Eu182%'      -- % to cover Synonym
				OR
				[ClinicalCode] COLLATE Latin1_General_CS_AS LIKE 'Eu142%'      -- % to cover Synonym
				OR
				[ClinicalCode] COLLATE Latin1_General_CS_AS LIKE 'Eu122%'      -- % to cover Synonym
				OR
				[ClinicalCode] COLLATE Latin1_General_CS_AS LIKE 'Eu152%'      -- % to cover Synonym
				OR
				[ClinicalCode] COLLATE Latin1_General_CS_AS LIKE 'Eu162%'      -- % to cover Synonym
				OR
				[ClinicalCode] COLLATE Latin1_General_CS_AS LIKE 'Eu132%'      -- % to cover Synonym
				OR
				[ClinicalCode] COLLATE Latin1_General_CS_AS IN('ZV114', '13cH', '146F', '13c1')
			)
			AND
			SK_PatientID IN(SELECT SK_PatientID FROM ceg.GenomicsData WHERE CCG = '08V')

	)as DependenceAbuse

	WHERE rn = 1

)as DependenceAndAbuse

WHERE
	SK_PatientID NOT IN	-- Excludes Alcohol dependence resolved codes if appearing after the latest acohol dependence diagnostic code
	(
		SELECT
			[SK_PatientID]

		FROM
		(
			SELECT
				 [SK_PatientID]
				,[ClinicalCode]
				,[EventDate]
				,row_number() over(partition by [SK_PatientID] order by [EventDate] DESC) as rn

			FROM
				[08V].[GPEncounter]

			WHERE
				(
					[ClinicalCode] COLLATE Latin1_General_CS_AS LIKE 'E23%'
					OR
					[ClinicalCode] COLLATE Latin1_General_CS_AS LIKE 'E24%'
					OR
					[ClinicalCode] COLLATE Latin1_General_CS_AS LIKE '13cM%'
					OR
					[ClinicalCode] COLLATE Latin1_General_CS_AS LIKE '1T%'
					OR
					[ClinicalCode] COLLATE Latin1_General_CS_AS LIKE 'Eu55%'       -- % to cover Synonym
					OR
					[ClinicalCode] COLLATE Latin1_General_CS_AS LIKE 'Eu112%'      -- % to cover Synonym
					OR
					[ClinicalCode] COLLATE Latin1_General_CS_AS LIKE 'Eu102%'      -- % to cover Synonym
					OR
					[ClinicalCode] COLLATE Latin1_General_CS_AS LIKE 'Eu182%'      -- % to cover Synonym
					OR
					[ClinicalCode] COLLATE Latin1_General_CS_AS LIKE 'Eu142%'      -- % to cover Synonym
					OR
					[ClinicalCode] COLLATE Latin1_General_CS_AS LIKE 'Eu122%'      -- % to cover Synonym
					OR
					[ClinicalCode] COLLATE Latin1_General_CS_AS LIKE 'Eu152%'      -- % to cover Synonym
					OR
					[ClinicalCode] COLLATE Latin1_General_CS_AS LIKE 'Eu162%'      -- % to cover Synonym
					OR
					[ClinicalCode] COLLATE Latin1_General_CS_AS LIKE 'Eu132%'      -- % to cover Synonym
					OR
					[ClinicalCode] COLLATE Latin1_General_CS_AS IN('ZV114', '13cH', '146F', '13c1')
					OR
					[ClinicalCode] COLLATE Latin1_General_CS_AS = '2126C'  --2126C	Alcohol dependence resolved
				)
				AND
				SK_PatientID IN(SELECT SK_PatientID FROM ceg.GenomicsData WHERE CCG = '08V')

		) AS  Res

		WHERE
			rn = 1	AND	[ClinicalCode] = '2126C'	--2126C	Alcohol dependence resolved
	)

GO

----------- ADHD/ADD -------------------------------------------------------------------------------------------------

SELECT
	 [SK_PatientID]
	,[ClinicalCode] ADHD_ADDCode
	,[EventDate]	ADHD_ADDDateRecorded

INTO  #ADHD_ADD

FROM
(
	SELECT
		 [SK_PatientID]
		,[ClinicalCode]
		,[EventDate]
		,row_number() over(partition by [SK_PatientID] order by [EventDate] ASC) as rn

	FROM  [08V].[GPEncounter]
	WHERE
		(
			[ClinicalCode] COLLATE Latin1_General_CS_AS IN('Eu9y7', 'Eu900', 'Eu900-1') -- Synonym (-)
			OR
			[ClinicalCode] COLLATE Latin1_General_CS_AS LIKE 'E2E0%'
		)
		AND
		SK_PatientID IN(SELECT SK_PatientID FROM ceg.GenomicsData WHERE CCG = '08V')

)as ADHD_ADD

WHERE rn = 1

GO

----------- Personality disorder -------------------------------------------------------------------------------------------------

SELECT
	 [SK_PatientID]
	,[ClinicalCode] PersonalityDisorderCode
	,[EventDate]	PersonalityDisorderDateRecorded

INTO  #PersonalityDisorder

FROM
(
	SELECT
		 [SK_PatientID]
		,[ClinicalCode]
		,[EventDate]
		,row_number() over(partition by [SK_PatientID] order by [EventDate] ASC) as rn

	FROM  [08V].[GPEncounter]
	WHERE
		[ClinicalCode] COLLATE Latin1_General_CS_AS LIKE 'E21%'
		AND
		SK_PatientID IN(SELECT SK_PatientID FROM ceg.GenomicsData WHERE CCG = '08V')

)as PersonalityDisorder

WHERE rn = 1

GO

----------- Habit and impulse disorders -------------------------------------------------------------------------------------------------

SELECT
	 [SK_PatientID]
	,[ClinicalCode] HabitCode
	,[EventDate]	HabitDateRecorded

INTO  #Habit

FROM
(
	SELECT
		 [SK_PatientID]
		,[ClinicalCode]
		,[EventDate]
		,row_number() over(partition by [SK_PatientID] order by [EventDate] ASC) as rn

	FROM  [08V].[GPEncounter]
	WHERE
		[ClinicalCode] COLLATE Latin1_General_CS_AS LIKE 'Eu63%'
		AND
		SK_PatientID IN(SELECT SK_PatientID FROM ceg.GenomicsData WHERE CCG = '08V')

)as Habit

WHERE rn = 1

GO

------------------Learning Disabilities -----------------------------------------------------------------------------

SELECT
	 [SK_PatientID]
	,[ClinicalCode] LearningDisabilitiesCode
	,[EventDate] LearningDisabilitiesDateRecorded

INTO #LearningDisabilities

FROM
(
	SELECT
		 [SK_PatientID]
		,[ClinicalCode]
		,[EventDate]
		,row_number() over(partition by [SK_PatientID] order by [EventDate] ASC) as rn

	FROM  [08V].[GPEncounter]
	WHERE
			--ELGH v8:   E3%, Eu7%, Eu814, Eu815, Eu816, Eu817, Eu81z, Eu818, 918e
		(
			[ClinicalCode] COLLATE Latin1_General_CS_AS LIKE 'E3%'
			OR
			[ClinicalCode] COLLATE Latin1_General_CS_AS LIKE 'Eu7%'
			OR
			[ClinicalCode] COLLATE Latin1_General_CS_AS LIKE 'Eu81z%'      -- % to cover Synonym
			OR
			[ClinicalCode] COLLATE Latin1_General_CS_AS IN('Eu814', 'Eu815', 'Eu816', 'Eu817', 'Eu818', '918e')
		)
		AND
		SK_PatientID IN(SELECT SK_PatientID FROM ceg.GenomicsData WHERE CCG = '08V')

)as LD

WHERE rn = 1

GO

----------- Autism/Aspergers -------------------------------------------------------------------------------------------------

SELECT
	 [SK_PatientID]
	,[ClinicalCode] AutismCode
	,[EventDate]	AutismDateRecorded

INTO  #Autism

FROM
(
	SELECT
		 [SK_PatientID]
		,[ClinicalCode]
		,[EventDate]
		,row_number() over(partition by [SK_PatientID] order by [EventDate] ASC) as rn

	FROM  [08V].[GPEncounter]
	WHERE
		(
			[ClinicalCode] COLLATE Latin1_General_CS_AS LIKE 'E140%'
			OR
			[ClinicalCode] COLLATE Latin1_General_CS_AS LIKE 'Eu840%'      -- % to cover Synonym
			OR
			[ClinicalCode] COLLATE Latin1_General_CS_AS LIKE 'Eu841%'      -- % to cover Synonym
		)
		AND
		SK_PatientID IN(SELECT SK_PatientID FROM ceg.GenomicsData WHERE CCG = '08V')

)as Autism

WHERE rn = 1

GO

----------- Speech/language disorder -------------------------------------------------------------------------------------------------

SELECT
	 [SK_PatientID]
	,[ClinicalCode] SpeechLanguageDisorderCode
	,[EventDate]	SpeechLanguageDisorderDateRecorded

INTO  #SpeechLanguageDisorder

FROM
(
	SELECT
		 [SK_PatientID]
		,[ClinicalCode]
		,[EventDate]
		,row_number() over(partition by [SK_PatientID] order by [EventDate] ASC) as rn

	FROM  [08V].[GPEncounter]
	WHERE
		(	-- E2F3%, Eu80%
			[ClinicalCode] COLLATE Latin1_General_CS_AS LIKE 'E2F3%'
			OR
			[ClinicalCode] COLLATE Latin1_General_CS_AS LIKE 'Eu80%'
		)
		AND
		SK_PatientID IN(SELECT SK_PatientID FROM ceg.GenomicsData WHERE CCG = '08V')

)as SpeechLanguageDisorder

WHERE rn = 1

GO

----------- Conduct disorder -------------------------------------------------------------------------------------------------

SELECT
	 [SK_PatientID]
	,[ClinicalCode] ConductDisorderCode
	,[EventDate]	ConductDisorderDateRecorded

INTO  #ConductDisorder

FROM
(
	SELECT
		 [SK_PatientID]
		,[ClinicalCode]
		,[EventDate]
		,row_number() over(partition by [SK_PatientID] order by [EventDate] ASC) as rn

	FROM  [08V].[GPEncounter]
	WHERE
		[ClinicalCode] COLLATE Latin1_General_CS_AS LIKE 'Eu91%'
		AND
		SK_PatientID IN(SELECT SK_PatientID FROM ceg.GenomicsData WHERE CCG = '08V')

)as ConductDisorder

WHERE rn = 1

GO

----------- Tourette's syndrome/tic disorder -------------------------------------------------------------------------------------------------

SELECT
	 [SK_PatientID]
	,[ClinicalCode] TouretteSyndromeDisorderCode
	,[EventDate]	TouretteSyndromeDisorderDateRecorded

INTO  #TouretteSyndromeDisorder

FROM
(
	SELECT
		 [SK_PatientID]
		,[ClinicalCode]
		,[EventDate]
		,row_number() over(partition by [SK_PatientID] order by [EventDate] ASC) as rn

	FROM  [08V].[GPEncounter]
	WHERE
		[ClinicalCode] COLLATE Latin1_General_CS_AS = 'E2723'
		AND
		SK_PatientID IN(SELECT SK_PatientID FROM ceg.GenomicsData WHERE CCG = '08V')

)as TouretteSyndromeDisorder

WHERE rn = 1

GO

----------- Gender identity disorders -------------------------------------------------------------------------------------------------

SELECT
	 [SK_PatientID]
	,[ClinicalCode] GenderIdentityDisordersCode
	,[EventDate]	GenderIdentityDisordersDateRecorded

INTO  #GenderIdentityDisorders

FROM
(
	SELECT
		 [SK_PatientID]
		,[ClinicalCode]
		,[EventDate]
		,row_number() over(partition by [SK_PatientID] order by [EventDate] ASC) as rn

	FROM  [08V].[GPEncounter]
	WHERE
		[ClinicalCode] COLLATE Latin1_General_CS_AS LIKE 'Eu64%'
		AND
		SK_PatientID IN(SELECT SK_PatientID FROM ceg.GenomicsData WHERE CCG = '08V')

)as GenderIdentityDisorders

WHERE rn = 1


GO


--------------------------------------------------------------------------------------------------------------------------
---------------------------------------- Joining Tables ------------------------------------------------------------------
--------------------------------------------------------------------------------------------------------------------------


SELECT

	(Select CommissionerName From Dictionary.[dbo].[Commissioner] Where CommissionerCode = '08V') AS Locality
	,P.EncryptedNHSNumber

	,P.SK_PatientID

	--NEURO & MENTAL HEALTH
	,MND.MNDCode
	,MND.MND_DateRecorded

	,MS.MSCode
	,MS.MSDateRecorded

	,Parkinsons.ParkinsonsCode
	,Parkinsons.ParkinsonsDateRecorded

	,AtypicalPD.AtypicalPD_Code
	,AtypicalPD.AtypicalPD_DateRecorded

	,VasDem.VasDemCode
	,VasDem.VasDemDateRecorded

	,Alzheimers.AlzheimersCode
	,Alzheimers.AlzheimersDateRecorded

	,DementiaQoF.DementiaCode
	,DementiaQoF.DementiaDateRecorded

	,Cognitive.CognitiveCode
	,Cognitive.CognitiveDateRecorded

	,Memory.MemoryCode
	,Memory.MemoryDateRecorded

	,BrainTumour.BrainTumourCode
	,BrainTumour.BrainTumourDateRecorded

	,BrainDamage.BrainDamageCode
	,BrainDamage.BrainDamageDateRecorded

	,Epilepsy.EpilepsyCode
	,Epilepsy.EpilepsyDateRecorded

	,SchizophrenicQof.Schizophrenic_QoFCode
	,SchizophrenicQof.Schizophrenic_QoFDateRecorded

	,Bipolar_QoF.Bipolar_QoFCode
	,Bipolar_QoF.Bipolar_QoFDateRecorded

	,DepressionQoF.DepressionQoFCode
	,DepressionQoF.DepressionQoFDateRecorded

	-- ANXIETY DISORDERS
	,Anxiety.AnxietyCode
	,Anxiety.AnxietyDateRecorded

	,PTSD.PTSDCode
	,PTSD.PTSDDateRecorded

	,OCD.OCDCode
	,OCD.OCDDateRecorded

	,PanicDisorder.PanicDisorderCode
	,PanicDisorder.PanicDisorderDateRecorded

	,SocialAnxiety.SocialAnxietyCode
	,SocialAnxiety.SocialAnxietyDateRecorded

	,Phobias.PhobiasCode
	,Phobias.PhobiasDateRecorded

	,SAD.SAD_Code
	,SAD.SAD_DateRecorded

	,Conversion.ConversionCode
	,Conversion.ConversionDateRecorded

	,Adjustment.AdjustmentCode
	,Adjustment.AdjustmentDateRecorded

	,Somatoform.SomatoformCode
	,Somatoform.SomatoformDateRecorded

	,EatingDisorder.EatingDisorderCode
	,EatingDisorder.EatingDisorderDateRecorded

	,DependenceAndAbuse.DependenceAndAbuseCode
	,DependenceAndAbuse.DependenceAndAbuseDateRecorded

	,ADHD_ADD.ADHD_ADDCode
	,ADHD_ADD.ADHD_ADDDateRecorded

	,PersonalityDisorder.PersonalityDisorderCode
	,PersonalityDisorder.PersonalityDisorderDateRecorded

	,Habit.HabitCode
	,Habit.HabitDateRecorded

	,LearningDisabilities.LearningDisabilitiesCode
	,LearningDisabilities.LearningDisabilitiesDateRecorded


	,Autism.AutismCode
	,Autism.AutismDateRecorded

	,SpeechLanguageDisorder.SpeechLanguageDisorderCode
	,SpeechLanguageDisorder.SpeechLanguageDisorderDateRecorded

	,ConductDisorder.ConductDisorderCode
	,ConductDisorder.ConductDisorderDateRecorded

	,TouretteSyndromeDisorder.TouretteSyndromeDisorderCode
	,TouretteSyndromeDisorder.TouretteSyndromeDisorderDateRecorded

	,GenderIdentityDisorders.GenderIdentityDisordersCode
	,GenderIdentityDisorders.GenderIdentityDisordersDateRecorded

FROM
	ceg.GenomicsData AS P

	--  EXTRA DIAGNOSES
	LEFT JOIN #MND AS MND ON MND.SK_PatientID = P.SK_PatientID
	LEFT JOIN #MS AS MS ON MS.SK_PatientID = P.SK_PatientID
	LEFT JOIN #Parkinsons AS Parkinsons ON Parkinsons.SK_PatientID = P.SK_PatientID
	LEFT JOIN #AtypicalPD AS AtypicalPD ON AtypicalPD.SK_PatientID = P.SK_PatientID
	LEFT JOIN #VasDem AS VasDem ON VasDem.SK_PatientID = P.SK_PatientID
	LEFT JOIN #Alzheimers AS Alzheimers ON Alzheimers.SK_PatientID = P.SK_PatientID
	LEFT JOIN #DementiaQoF AS DementiaQoF ON DementiaQoF.SK_PatientID = P.SK_PatientID
	LEFT JOIN #Cognitive AS Cognitive ON Cognitive.SK_PatientID = P.SK_PatientID
	LEFT JOIN #Memory AS Memory ON Memory.SK_PatientID = P.SK_PatientID
	LEFT JOIN #BrainTumour AS BrainTumour ON BrainTumour.SK_PatientID = P.SK_PatientID
	LEFT JOIN #BrainDamage AS BrainDamage ON BrainDamage.SK_PatientID = P.SK_PatientID
	LEFT JOIN #Epilepsy AS Epilepsy ON Epilepsy.SK_PatientID = P.SK_PatientID
	LEFT JOIN #SchizophrenicQof AS SchizophrenicQof ON SchizophrenicQof.SK_PatientID = P.SK_PatientID
	LEFT JOIN #Bipolar_QoF AS Bipolar_QoF ON Bipolar_QoF.SK_PatientID = P.SK_PatientID

	LEFT JOIN #DepressionQoF AS DepressionQoF ON DepressionQoF.SK_PatientID = P.SK_PatientID

	-- ANXIETY DISORDERS
	LEFT JOIN #Anxiety	AS Anxiety  ON Anxiety.SK_PatientID = P.SK_PatientID
	LEFT JOIN #PTSD	AS PTSD ON PTSD.SK_PatientID = P.SK_PatientID
	LEFT JOIN #OCD	AS OCD ON OCD.SK_PatientID = P.SK_PatientID
	LEFT JOIN #PanicDisorder AS PanicDisorder ON PanicDisorder.SK_PatientID = P.SK_PatientID
	LEFT JOIN #SocialAnxiety AS SocialAnxiety ON SocialAnxiety.SK_PatientID = P.SK_PatientID
	LEFT JOIN #Phobias AS Phobias ON Phobias.SK_PatientID = P.SK_PatientID
	LEFT JOIN #SAD AS SAD ON SAD.SK_PatientID = P.SK_PatientID
	LEFT JOIN #Conversion	AS Conversion  ON Conversion.SK_PatientID = P.SK_PatientID
	LEFT JOIN #Adjustment AS Adjustment ON Adjustment.SK_PatientID = P.SK_PatientID
	LEFT JOIN #Somatoform AS Somatoform ON Somatoform.SK_PatientID = P.SK_PatientID
	LEFT JOIN #EatingDisorder AS EatingDisorder ON EatingDisorder.SK_PatientID = P.SK_PatientID
	LEFT JOIN #DependenceAndAbuse	AS DependenceAndAbuse ON DependenceAndAbuse.SK_PatientID = P.SK_PatientID
	LEFT JOIN #ADHD_ADD	AS ADHD_ADD ON ADHD_ADD.SK_PatientID = P.SK_PatientID
	LEFT JOIN #PersonalityDisorder AS PersonalityDisorder ON PersonalityDisorder.SK_PatientID = P.SK_PatientID
	LEFT JOIN #Habit AS Habit ON Habit.SK_PatientID = P.SK_PatientID
	LEFT JOIN #LearningDisabilities	AS LearningDisabilities ON LearningDisabilities.SK_PatientID = P.SK_PatientID
	LEFT JOIN #Autism AS Autism ON Autism.SK_PatientID = P.SK_PatientID
	LEFT JOIN #SpeechLanguageDisorder AS SpeechLanguageDisorder ON SpeechLanguageDisorder.SK_PatientID = P.SK_PatientID
	LEFT JOIN #ConductDisorder AS ConductDisorder ON ConductDisorder.SK_PatientID = P.SK_PatientID
	LEFT JOIN #TouretteSyndromeDisorder	AS TouretteSyndromeDisorder ON TouretteSyndromeDisorder.SK_PatientID = P.SK_PatientID
	LEFT JOIN #GenderIdentityDisorders AS GenderIdentityDisorders ON GenderIdentityDisorders.SK_PatientID = P.SK_PatientID

WHERE
	P.CCG = '08V'

GO


-- drop all temp tables used
BEGIN -- drop all temp tables used

	DECLARE @DropGlobal bit=0 --Default dont drop global temp table

	DECLARE @DROP_STATEMENT nvarchar(1000)
	DECLARE cursorDEL CURSOR FOR
	SELECT 'DROP TABLE '
	   + case
			   when name like '##%' then name
			   when name like '#%' then SUBSTRING(name, 1, CHARINDEX( '____', name)-1)
		end as DropSQL
	from tempdb..sysobjects
	WHERE name LIKE '#%'
	   AND OBJECT_ID('tempdb..' + name) IS NOT NULL
	   AND name not like case
						   when @DropGlobal=0 then '##%' --//Exclude global temp
						   else '#######%'    --//some fack expression so we can
											   --//select global temp for delete
						end
	OPEN cursorDEL
	FETCH NEXT FROM cursorDEL INTO @DROP_STATEMENT
	WHILE @@FETCH_STATUS = 0
	BEGIN
	EXEC (@DROP_STATEMENT)
	--print @DROP_STATEMENT
	FETCH NEXT FROM cursorDEL INTO @DROP_STATEMENT
	END
	CLOSE cursorDEL
	DEALLOCATE cursorDEL

END

GO
```

4. Clinic data

```
USE CEG

-- drop all temp tables if exist
BEGIN -- drop all temp tables if exist

	DECLARE @DropGlobal bit=0 --Default dont drop global temp table

	DECLARE @DROP_STATEMENT nvarchar(1000)
	DECLARE cursorDEL CURSOR FOR
	SELECT 'DROP TABLE '
	   + case
			   when name like '##%' then name
			   when name like '#%' then SUBSTRING(name, 1, CHARINDEX( '____', name)-1)
		end as DropSQL
	from tempdb..sysobjects
	WHERE name LIKE '#%'
	   AND OBJECT_ID('tempdb..' + name) IS NOT NULL
	   AND name not like case
						   when @DropGlobal=0 then '##%' --//Exclude global temp
						   else '#######%'    --//some fack expression so we can
											   --//select global temp for delete
						end

	OPEN cursorDEL
	FETCH NEXT FROM cursorDEL INTO @DROP_STATEMENT
	WHILE @@FETCH_STATUS = 0
	BEGIN
	EXEC (@DROP_STATEMENT)
	--print @DROP_STATEMENT
	FETCH NEXT FROM cursorDEL INTO @DROP_STATEMENT
	END
	CLOSE cursorDEL
	DEALLOCATE cursorDEL

END

GO

-- REPORT 2 CLINIC DATA --------------------------------------------------------------------------------------------------


-- DIABETES DIAGNOSES --------------------------------------------------------------------------------------------------
------------------ Type 1 Diabetes QOF earliest ever -----------------------------------------------------------------------


SELECT
	 [SK_PatientID]
	,[ClinicalCode] DiabetesT1Code
	,[EventDate]	DiabetesT1DateRecorded

INTO  #DiabetesT1QoF

FROM
(
	SELECT
		*

	FROM
	(
		SELECT
			 [SK_PatientID]
			,[ClinicalCode]
			,[EventDate]
			,row_number() over(partition by [SK_PatientID] order by [EventDate] ASC) as rn

		FROM
			[08V].[GPEncounter]

		WHERE
			(
				[ClinicalCode] COLLATE Latin1_General_CS_AS LIKE 'C10E%'	--C10E% Type 1 diabetes mellitus
				OR
				[ClinicalCode] COLLATE Latin1_General_CS_AS LIKE 'C10ER%'	-- % for Synonym		--C10ER Latent autoimmune diabetes mellitus in adult
			)
			AND
			SK_PatientID IN(SELECT SK_PatientID FROM ceg.GenomicsData WHERE CCG = '08V')

	)as T1

	WHERE
		rn = 1

) AS DiabT1

WHERE
	SK_PatientID NOT IN	-- QOF definition for Diabetes excludes diabetes resolved codes if appearing after the latest diagnostic code
	(
		SELECT
			[SK_PatientID]

		FROM
		(
			SELECT
				 [SK_PatientID]
				,[ClinicalCode]
				,[EventDate]
				,row_number() over(partition by [SK_PatientID] order by [EventDate] DESC) as rn

			FROM
				[08V].[GPEncounter]

			WHERE
				(
					[ClinicalCode] COLLATE Latin1_General_CS_AS LIKE 'C10E%'	--C10E% Type 1 diabetes mellitus
					OR
					[ClinicalCode] COLLATE Latin1_General_CS_AS LIKE 'C10ER%'	-- % for Synonym		--C10ER Latent autoimmune diabetes mellitus in adult
					OR
					[ClinicalCode] COLLATE Latin1_General_CS_AS IN('21263', '212H')  --Diabetes resolved
				)
				AND
				SK_PatientID IN(SELECT SK_PatientID FROM ceg.GenomicsData WHERE CCG = '08V')

		) AS  Res

		WHERE
			rn = 1	AND	[ClinicalCode] IN('21263', '212H')	--Diabetes resolved
	)

GO


------------------ Type 2 Diabetes QOF earliest ever -----------------------------------------------------------------------

/*
C10F% Type 2 diabetes mellitus (exclude C10F8 Reaven's syndrome),
C109J Insulin treated Type 2 diabetes mellitus,
*/


SELECT
	 [SK_PatientID]
	,[ClinicalCode] DiabetesT2Code
	,[EventDate]	DiabetesT2DateRecorded

INTO  #DiabetesT2QoF

FROM
(
	SELECT
		*

	FROM
	(
		SELECT
			 [SK_PatientID]
			,[ClinicalCode]
			,[EventDate]
			,row_number() over(partition by [SK_PatientID] order by [EventDate] ASC) as rn

		FROM
			[08V].[GPEncounter]

		WHERE
			(
				(
					[ClinicalCode] COLLATE Latin1_General_CS_AS LIKE 'C10F%' --C10F% Type 2 diabetes mellitus
					AND
					[ClinicalCode] COLLATE Latin1_General_CS_AS <> 'C10F8' --C10F8 Reaven's syndrome
				)
				OR
				[ClinicalCode] COLLATE Latin1_General_CS_AS  LIKE 'C109J%' -- % for Synonym --C109J Insulin treated Type 2 diabetes mellitus
			)
			AND
			SK_PatientID IN(SELECT SK_PatientID FROM ceg.GenomicsData WHERE CCG = '08V')

	)as T2

	WHERE
		rn = 1

) AS DiabT2

WHERE
	SK_PatientID NOT IN	-- QOF definition for Diabetes excludes diabetes resolved codes if appearing after the latest diagnostic code
	(
		SELECT
			[SK_PatientID]

		FROM
		(
			SELECT
				 [SK_PatientID]
				,[ClinicalCode]
				,[EventDate]
				,row_number() over(partition by [SK_PatientID] order by [EventDate] DESC) as rn

			FROM
				[08V].[GPEncounter]

			WHERE
				(
					(
						[ClinicalCode] COLLATE Latin1_General_CS_AS LIKE 'C10F%'		--C10F% Type 2 diabetes mellitus
						AND
						[ClinicalCode] COLLATE Latin1_General_CS_AS <> 'C10F8'				--C10F8 Reaven's syndrome
					)
					OR
					[ClinicalCode] COLLATE Latin1_General_CS_AS  LIKE 'C109J%' -- % for Synonym			--C109J Insulin treated Type 2 diabetes mellitus
					OR
					[ClinicalCode] COLLATE Latin1_General_CS_AS IN('21263', '212H')  --Diabetes resolved
				)
				AND
				SK_PatientID IN(SELECT SK_PatientID FROM ceg.GenomicsData WHERE CCG = '08V')

		) AS  Res

		WHERE
			rn = 1	AND	[ClinicalCode] IN('21263', '212H')	--Diabetes resolved
	)

GO


-- CLINIC DATA  ---------------------------------------------------------------------------------------------------------
------------------ Current smoker (QOF v37) ------------------------------------------------------------------------

SELECT
	 [SK_PatientID]
	,[ClinicalCode] CurrentSmokerQoFCode
	,[EventDate]	CurrentSmokerQoFDateRecorded

INTO  #CurrentSmokerQoF

FROM
(
	SELECT
		 [SK_PatientID]
		,[ClinicalCode]
		,[EventDate]
		,row_number() over(partition by E.[SK_PatientID] order by [EventDate] DESC) as rn

	FROM  [08V].[GPEncounter] E
	WHERE
		(	-- QOF v37: 1372.-1376. , 137C.-137D. , 137G.-137H. , 137J. , 137M. , 137P.-137R. , 137V. , 137X.-137f. , 137h. , 137m. , 137o. , 137..

			[ClinicalCode] COLLATE Latin1_General_CS_AS  IN ('137', '137-1')	--Synonym(-)
			OR
			[ClinicalCode] COLLATE Latin1_General_CS_AS LIKE '137[2-6]%'		-- % to cover Synonym
			OR
			[ClinicalCode] COLLATE Latin1_General_CS_AS IN('137C', '137D', '137G', '137H', '137J', '137M', '137P', '137P-1', '137Q', '137Q-1', '137R', '137V', '137X', '137Y', '137Z')	--Synonym(-)
			OR
			[ClinicalCode] COLLATE Latin1_General_CS_AS IN('137a', '137b', '137c', '137d', '137e', '137f', '137h', '137m', '137o')
		)
		AND
		SK_PatientID IN(SELECT G.SK_PatientID FROM ceg.GenomicsData AS G WHERE CCG = '08V')

)as CurrentSmokerQoF

WHERE rn = 1

GO

------------------ Ex-smoker (QOF v37) ------------------------------------------------------------------------

SELECT
	 [SK_PatientID]
	,[ClinicalCode] ExSmokerQoFCode
	,[EventDate]	ExSmokerQoFDateRecorded

INTO  #ExSmokerQoF

FROM
(
	SELECT
		 [SK_PatientID]
		,[ClinicalCode]
		,[EventDate]
		,row_number() over(partition by E.[SK_PatientID] order by [EventDate] DESC) as rn

	FROM  [08V].[GPEncounter] E
	WHERE
		(	-- QOF v37: 1377.-137B. , 137F. , 137K. , 137N.-137O. , 137S.-137T. , 137j. , 137l.

			[ClinicalCode] COLLATE Latin1_General_CS_AS  IN ('1377', '1378', '1379')
			OR
			[ClinicalCode] COLLATE Latin1_General_CS_AS IN('137A', '137B', '137F', '137K', '137N', '137O', '137S', '137T')
			OR
			[ClinicalCode] COLLATE Latin1_General_CS_AS IN('137j', '137l')
		)
		AND
		SK_PatientID IN(SELECT G.SK_PatientID FROM ceg.GenomicsData AS G WHERE CCG = '08V')

)as ExSmokerQoF

WHERE rn = 1

GO

------------------ Never smoked (QOF v37) ------------------------------------------------------------------------

SELECT
	 [SK_PatientID]
	,[ClinicalCode] NeverSmokedQoFCode
	,[EventDate]	NeverSmokedQoFDateRecorded

INTO  #NeverSmokedQoF

FROM
(
	SELECT
		 [SK_PatientID]
		,[ClinicalCode]
		,[EventDate]
		,row_number() over(partition by E.[SK_PatientID] order by [EventDate] DESC) as rn

	FROM  [08V].[GPEncounter] E
	WHERE
		-- QOF v37: 1371
		[ClinicalCode] COLLATE Latin1_General_CS_AS IN ('1371', '1371-1')
		AND
		SK_PatientID IN(SELECT G.SK_PatientID FROM ceg.GenomicsData AS G WHERE CCG = '08V')

)as NeverSmokedQoF

WHERE rn = 1

GO


------------------  On examination weight (O/E weight) --------------------------------------------------------
---------------------- Weight Earliest ----------------------------------------------------------------------------


SELECT
	 [SK_PatientID]
	,[ClinicalCode] WeightEarliestCode
	,[EventDate] WeightEarliestDate
	,[Value] WeightEarliestScore
	,[Units] WeightEarliestUnits
	,[AgeAtEvent] WeightEarliestAgeAtEvent

INTO  #WeightEarliest

FROM
(
	SELECT
		 [SK_PatientID]
		,[ClinicalCode]
		,[EventDate]
		,[Value]
		,[Units]
		,[AgeAtEvent]
		,row_number() over(partition by [SK_PatientID] order by [EventDate] ASC) as rn

	FROM  [08V].[GPEncounter]
	WHERE
		[ClinicalCode] COLLATE Latin1_General_CS_AS = '22A'
		AND
		[Value] <> 0
		AND
		SK_PatientID IN(SELECT SK_PatientID FROM ceg.GenomicsData WHERE CCG = '08V')

)as WeightEarliest

WHERE rn = 1

GO

---------------------- Weight +/- 6 month window around  T1 or T2 diagnosis -------------------------------------


SELECT
	 [SK_PatientID]
	,[ClinicalCode] Weight6MonthsWindowCode
	,[EventDate] Weight6MonthsWindowDate
	,[Value] Weight6MonthsWindowScore
	,[Units] Weight6MonthsWindowUnits
	,[AgeAtEvent] Weight6MonthsWindowAgeAtEvent

INTO  #Weight6MonthsWindow

FROM
(
	SELECT
		 E.[SK_PatientID]
		,[ClinicalCode]
		,[EventDate]
		,[Value]
		,[Units]
		,[AgeAtEvent]
		,row_number() over(partition by E.[SK_PatientID] order by [EventDate] ASC) as rn

	FROM  [08V].[GPEncounter] E
	LEFT JOIN #DiabetesT1QoF AS T1 ON T1.SK_PatientID = E.SK_PatientID
	LEFT JOIN #DiabetesT2QoF AS T2 ON T2.SK_PatientID = E.SK_PatientID

	WHERE
		[ClinicalCode] COLLATE Latin1_General_CS_AS = '22A'
		AND
		[Value] <> 0
		AND
		(
			T1.SK_PatientID IS NOT NULL
			OR
			T2.SK_PatientID IS NOT NULL
		)
		AND
		(
			[EventDate] BETWEEN DATEADD(month, -6, T1.DiabetesT1DateRecorded) AND DATEADD(month, 6, T1.DiabetesT1DateRecorded)
			OR
			[EventDate] BETWEEN DATEADD(month, -6, T2.DiabetesT2DateRecorded) AND DATEADD(month, 6, T2.DiabetesT2DateRecorded)
		)
		AND
		E.SK_PatientID IN(SELECT SK_PatientID FROM ceg.GenomicsData WHERE CCG = '08V')

) AS Weight6MonthsWindow

WHERE rn = 1

GO


---------------------- Weight Latest ever----------------------------------------------------------------------------


SELECT
	 [SK_PatientID]
	,[ClinicalCode] WeightLatestCode
	,[EventDate] WeightLatestDate
	,[Value] WeightLatestScore
	,[Units] WeightLatestUnits
	,[AgeAtEvent] WeightLatestAgeAtEvent

INTO  #WeightLatest

FROM
(
	SELECT
		 [SK_PatientID]
		,[ClinicalCode]
		,[EventDate]
		,[Value]
		,[Units]
		,[AgeAtEvent]
		,row_number() over(partition by [SK_PatientID] order by [EventDate] DESC) as rn

	FROM  [08V].[GPEncounter]
	WHERE
		[ClinicalCode] COLLATE Latin1_General_CS_AS = '22A'
		AND
		[Value] <> 0
		AND
		SK_PatientID IN(SELECT SK_PatientID FROM ceg.GenomicsData WHERE CCG = '08V')

)as WeightLatest

WHERE rn = 1

GO


------------------  On examination height (O/E height) --------------------------------------------------------
----------------------Height Earliest ----------------------------------------------------------------------------

SELECT
	 [SK_PatientID]
	,[ClinicalCode] HeightEarliestCode
	,[EventDate] HeightEarliestDate
	,[Value] HeightEarliestScore
	,[Units] HeightEarliestUnits
	,[AgeAtEvent] HeightEarliestAgeAtEvent

INTO  #HeightEarliest

FROM
(
	SELECT
		 [SK_PatientID]
		,[ClinicalCode]
		,[EventDate]
		,[Value]
		,[Units]
		,[AgeAtEvent]
		,row_number() over(partition by [SK_PatientID] order by [EventDate] ASC) as rn

	FROM  [08V].[GPEncounter]
	WHERE
		[ClinicalCode] COLLATE Latin1_General_CS_AS = '229'
		AND
		[Value] <> 0
		AND
		SK_PatientID IN(SELECT SK_PatientID FROM ceg.GenomicsData WHERE CCG = '08V')

)as HeightEarliest

WHERE rn = 1

GO

---------------------- Height +/- 6 month window around  T1 or T2 diagnosis -------------------------------------

SELECT
	 [SK_PatientID]
	,[ClinicalCode] Height6MonthsWindowCode
	,[EventDate] Height6MonthsWindowDate
	,[Value] Height6MonthsWindowScore
	,[Units] Height6MonthsWindowUnits
	,[AgeAtEvent] Height6MonthsWindowAgeAtEvent

INTO  #Height6MonthsWindow

FROM
(
	SELECT
		 E.[SK_PatientID]
		,[ClinicalCode]
		,[EventDate]
		,[Value]
		,[Units]
		,[AgeAtEvent]
		,row_number() over(partition by E.[SK_PatientID] order by [EventDate] ASC) as rn

	FROM  [08V].[GPEncounter] E
	LEFT JOIN #DiabetesT1QoF AS T1 ON T1.SK_PatientID = E.SK_PatientID
	LEFT JOIN #DiabetesT2QoF AS T2 ON T2.SK_PatientID = E.SK_PatientID

	WHERE
		[ClinicalCode] COLLATE Latin1_General_CS_AS = '229'
		AND
		[Value] <> 0
		AND
		(
			T1.SK_PatientID IS NOT NULL
			OR
			T2.SK_PatientID IS NOT NULL
		)
		AND
		(
			[EventDate] BETWEEN DATEADD(month, -6, T1.DiabetesT1DateRecorded) AND DATEADD(month, 6, T1.DiabetesT1DateRecorded)
			OR
			[EventDate] BETWEEN DATEADD(month, -6, T2.DiabetesT2DateRecorded) AND DATEADD(month, 6, T2.DiabetesT2DateRecorded)
		)
		AND
		E.SK_PatientID IN(SELECT SK_PatientID FROM ceg.GenomicsData WHERE CCG = '08V')

) AS Height6MonthsWindow

WHERE rn = 1

GO


---------------------- Height Latest ----------------------------------------------------------------------------

SELECT
	 [SK_PatientID]
	,[ClinicalCode] HeightLatestCode
	,[EventDate] HeightLatestDate
	,[Value] HeightLatestScore
	,[Units] HeightLatestUnits
	,[AgeAtEvent] HeightLatestAgeAtEvent

INTO  #HeightLatest

FROM
(
	SELECT
		 [SK_PatientID]
		,[ClinicalCode]
		,[EventDate]
		,[Value]
		,[Units]
		,[AgeAtEvent]
		,row_number() over(partition by [SK_PatientID] order by [EventDate] DESC) as rn

	FROM  [08V].[GPEncounter]
	WHERE
		[ClinicalCode] COLLATE Latin1_General_CS_AS = '229'
		AND
		[Value] <> 0
		AND
		SK_PatientID IN(SELECT SK_PatientID FROM ceg.GenomicsData WHERE CCG = '08V')

)as HeightLatest

WHERE rn = 1

GO


------------------  BMI  ---------------------------------------------------------------------------------------
----------------------BMI Earliest ----------------------------------------------------------------------------


SELECT
	 [SK_PatientID]
	,[ClinicalCode] BMIEarliestCode
	,[EventDate] BMIEarliestDate
	,[Value] BMIEarliestScore
	,[Units] BMIEarliestUnits
	,[AgeAtEvent] BMIEarliestAgeAtEvent

INTO  #BMIEarliest

FROM
(
	SELECT
		 [SK_PatientID]
		,[ClinicalCode]
		,[EventDate]
		,[Value]
		,[Units]
		,[AgeAtEvent]
		,row_number() over(partition by [SK_PatientID] order by [EventDate] ASC) as rn

	FROM  [08V].[GPEncounter]
	WHERE
		[ClinicalCode] COLLATE Latin1_General_CS_AS = '22K'
		AND
		[Value] <> 0
		AND
		SK_PatientID IN(SELECT SK_PatientID FROM ceg.GenomicsData WHERE CCG = '08V')

)as BMIEarliest

WHERE rn = 1

GO


---------------------- BMI +/- 6 month window around  T1 or T2 diagnosis -------------------------------------


SELECT
	 [SK_PatientID]
	,[ClinicalCode] BMI6MonthsWindowCode
	,[EventDate] BMI6MonthsWindowDate
	,[Value] BMI6MonthsWindowScore
	,[Units] BMI6MonthsWindowUnits
	,[AgeAtEvent] BMI6MonthsWindowAgeAtEvent

INTO  #BMI6MonthsWindow

FROM
(
	SELECT
		 E.[SK_PatientID]
		,[ClinicalCode]
		,[EventDate]
		,[Value]
		,[Units]
		,[AgeAtEvent]
		,row_number() over(partition by E.[SK_PatientID] order by [EventDate] ASC) as rn

	FROM  [08V].[GPEncounter] E
	LEFT JOIN #DiabetesT1QoF AS T1 ON T1.SK_PatientID = E.SK_PatientID
	LEFT JOIN #DiabetesT2QoF AS T2 ON T2.SK_PatientID = E.SK_PatientID

	WHERE
		[ClinicalCode] COLLATE Latin1_General_CS_AS = '22K'
		AND
		[Value] <> 0
		AND
		(
			T1.SK_PatientID IS NOT NULL
			OR
			T2.SK_PatientID IS NOT NULL
		)
		AND
		(
			[EventDate] BETWEEN DATEADD(month, -6, T1.DiabetesT1DateRecorded) AND DATEADD(month, 6, T1.DiabetesT1DateRecorded)
			OR
			[EventDate] BETWEEN DATEADD(month, -6, T2.DiabetesT2DateRecorded) AND DATEADD(month, 6, T2.DiabetesT2DateRecorded)
		)
		AND
		E.SK_PatientID IN(SELECT SK_PatientID FROM ceg.GenomicsData WHERE CCG = '08V')

) AS BMI6MonthsWindow

WHERE rn = 1

GO


---------------------- BMI Latest ----------------------------------------------------------------------------


SELECT
	 [SK_PatientID]
	,[ClinicalCode] BMILatestCode
	,[EventDate] BMILatestDate
	,[Value] BMILatestScore
	,[Units] BMILatestUnits
	,[AgeAtEvent] BMILatestAgeAtEvent

INTO  #BMILatest

FROM
(
	SELECT
		 [SK_PatientID]
		,[ClinicalCode]
		,[EventDate]
		,[Value]
		,[Units]
		,[AgeAtEvent]
		,row_number() over(partition by [SK_PatientID] order by [EventDate] DESC) as rn

	FROM  [08V].[GPEncounter]
	WHERE
		[ClinicalCode] COLLATE Latin1_General_CS_AS = '22K'
		AND
		[Value] <> 0
		AND
		SK_PatientID IN(SELECT SK_PatientID FROM ceg.GenomicsData WHERE CCG = '08V')

)as BMILatest

WHERE rn = 1

GO

------------------ HbA1c ----------------------------------------------------------------------------------------------
----------------------HbA1c Earliest ----------------------------------------------------------------------------------

SELECT
	 [SK_PatientID]
	,[ClinicalCode] HbA1cEarliestCode
	,[EventDate] HbA1cEarliestDateRecorded
	,[Value] HbA1cEarliestValue
	,[Units] HbA1cEarliestUnits

INTO  #HbA1cEarliest

FROM
(
	SELECT

		 [SK_PatientID]
		,[ClinicalCode]
		,[EventDate]
		,[Value]
		,[Units]
		,row_number() over(partition by [SK_PatientID] order by [EventDate] ASC) as rn

	FROM  [08V].[GPEncounter]
	WHERE

		[ClinicalCode] COLLATE Latin1_General_CS_AS IN('42W4', '42W5')
		AND
		[Value] <> 0
		AND
		SK_PatientID IN(SELECT SK_PatientID FROM ceg.GenomicsData WHERE CCG = '08V')

)as HbA1cEarliest

WHERE rn = 1 --AND HbA1cEarliest.VALUE > 0

GO


---------------------- HbA1c +/- 6 month window around  T1 or T2 diagnosis -------------------------------------

SELECT
	 [SK_PatientID]
	,[ClinicalCode] HbA1c6MonthsWindowCode
	,[EventDate] HbA1c6MonthsWindowDate
	,[Value] HbA1c6MonthsWindowScore
	,[Units] HbA1c6MonthsWindowUnits

INTO  #HbA1c6MonthsWindow

FROM
(
	SELECT
		 E.[SK_PatientID]
		,[ClinicalCode]
		,[EventDate]
		,[Value]
		,[Units]
		,row_number() over(partition by E.[SK_PatientID] order by [EventDate] ASC) as rn

	FROM  [08V].[GPEncounter] E
	LEFT JOIN #DiabetesT1QoF AS T1 ON T1.SK_PatientID = E.SK_PatientID
	LEFT JOIN #DiabetesT2QoF AS T2 ON T2.SK_PatientID = E.SK_PatientID

	WHERE
		[ClinicalCode] COLLATE Latin1_General_CS_AS IN('42W4', '42W5')
		AND
		[Value] <> 0
		AND
		(
			T1.SK_PatientID IS NOT NULL
			OR
			T2.SK_PatientID IS NOT NULL
		)
		AND
		(
			[EventDate] BETWEEN DATEADD(month, -6, T1.DiabetesT1DateRecorded) AND DATEADD(month, 6, T1.DiabetesT1DateRecorded)
			OR
			[EventDate] BETWEEN DATEADD(month, -6, T2.DiabetesT2DateRecorded) AND DATEADD(month, 6, T2.DiabetesT2DateRecorded)
		)
		AND
		E.SK_PatientID IN(SELECT SK_PatientID FROM ceg.GenomicsData WHERE CCG = '08V')

) AS HbA1c6MonthsWindow

WHERE rn = 1

GO

---------------------- HbA1c Latest ----------------------------------------------------------------------------

SELECT
	 [SK_PatientID]
	,[ClinicalCode] HbA1cLatestCode
	,[EventDate] HbA1cLatestDateRecorded
	,[Value] HbA1cLatestValue
	,[Units] HbA1cLatestUnits

INTO  #HbA1cLatest

FROM
(
	SELECT
		 [SK_PatientID]
		,[ClinicalCode]
		,[EventDate]
		,[Value]
		,[Units]
		,row_number() over(partition by [SK_PatientID] order by [EventDate] DESC) as rn

	FROM  [08V].[GPEncounter]
	WHERE

		[ClinicalCode] COLLATE Latin1_General_CS_AS IN('42W4', '42W5')
		AND
		[Value] <> 0
		AND
		SK_PatientID IN(SELECT SK_PatientID FROM ceg.GenomicsData WHERE CCG = '08V')

)as HbA1cLatest


WHERE rn = 1

GO


------------------Cholesterol ----------------------------------------------------------------------------------------------
---------------------- Cholesterol Earliest ----------------------------------------------------------------------------

SELECT
	 [SK_PatientID]
	,[ClinicalCode] CholesterolEarliestCode
	,[EventDate] CholesterolEarliestDateRecorded
	,[Value] CholesterolEarliestValue
	,[Units] CholesterolEarliestUnits

INTO  #CholesterolEarliest

FROM
(
	SELECT

		 [SK_PatientID]
		,[ClinicalCode]
		,[EventDate]
		,[Value]
		,[Units]
		,row_number() over(partition by [SK_PatientID] order by [EventDate] ASC) as rn

	FROM  [08V].[GPEncounter]
	WHERE

		[ClinicalCode] COLLATE Latin1_General_CS_AS = '44P'
		AND
		Value <> 0
		AND
		SK_PatientID IN(SELECT SK_PatientID FROM ceg.GenomicsData WHERE CCG = '08V')

)as CholesterolEarliest

WHERE rn = 1

GO


---------------------- Cholesterol +/- 6 month window around  T1 or T2 diagnosis -------------------------------------

SELECT
	 [SK_PatientID]
	,[ClinicalCode] Cholesterol6MonthsWindowCode
	,[EventDate] Cholesterol6MonthsWindowDate
	,[Value] Cholesterol6MonthsWindowScore
	,[Units] Cholesterol6MonthsWindowUnits

INTO  #Cholesterol6MonthsWindow

FROM
(
	SELECT
		 E.[SK_PatientID]
		,[ClinicalCode]
		,[EventDate]
		,[Value]
		,[Units]
		,row_number() over(partition by E.[SK_PatientID] order by [EventDate] ASC) as rn

	FROM  [08V].[GPEncounter] E
	LEFT JOIN #DiabetesT1QoF AS T1 ON T1.SK_PatientID = E.SK_PatientID
	LEFT JOIN #DiabetesT2QoF AS T2 ON T2.SK_PatientID = E.SK_PatientID

	WHERE
		[ClinicalCode] COLLATE Latin1_General_CS_AS = '44P'
		AND
		[Value] <> 0
		AND
		(
			T1.SK_PatientID IS NOT NULL
			OR
			T2.SK_PatientID IS NOT NULL
		)
		AND
		(
			[EventDate] BETWEEN DATEADD(month, -6, T1.DiabetesT1DateRecorded) AND DATEADD(month, 6, T1.DiabetesT1DateRecorded)
			OR
			[EventDate] BETWEEN DATEADD(month, -6, T2.DiabetesT2DateRecorded) AND DATEADD(month, 6, T2.DiabetesT2DateRecorded)
		)
		AND
		E.SK_PatientID IN(SELECT SK_PatientID FROM ceg.GenomicsData WHERE CCG = '08V')

) AS Cholesterol6MonthsWindow

WHERE rn = 1

GO


---------------------- Cholesterol Latest ----------------------------------------------------------------------------

SELECT
	 [SK_PatientID]
	,[ClinicalCode] CholesterolLatestCode
	,[EventDate] CholesterolLatestDateRecorded
	,[Value] CholesterolLatestValue
	,[Units] CholesterolLatestUnits

INTO  #CholesterolLatest

FROM
(
	SELECT
		 [SK_PatientID]
		,[ClinicalCode]
		,[EventDate]
		,[Value]
		,[Units]
		,row_number() over(partition by [SK_PatientID] order by [EventDate] DESC) as rn

	FROM  [08V].[GPEncounter]
	WHERE

		[ClinicalCode] COLLATE Latin1_General_CS_AS = '44P'
		AND
		[Value] <> 0
		AND
		SK_PatientID IN(SELECT SK_PatientID FROM ceg.GenomicsData WHERE CCG = '08V')

)as CholesterolLatest


WHERE rn = 1

GO


------------------HDL cholesterol ----------------------------------------------------------------------------------------------
---------------------- HDL cholesterol Earliest ----------------------------------------------------------------------------

SELECT
	 [SK_PatientID]
	,[ClinicalCode] HDL_CholesterolEarliestCode
	,[EventDate] HDL_CholesterolEarliestDateRecorded
	,[Value] HDL_CholesterolEarliestValue
	,[Units] HDL_CholesterolEarliestUnits

INTO  #HDL_CholesterolEarliest

FROM
(
	SELECT

		 [SK_PatientID]
		,[ClinicalCode]
		,[EventDate]
		,[Value]
		,[Units]
		,row_number() over(partition by [SK_PatientID] order by [EventDate] ASC) as rn

	FROM  [08V].[GPEncounter]
	WHERE

		[ClinicalCode] COLLATE Latin1_General_CS_AS = '44P5'
		AND
		[Value] <> 0
		AND
		SK_PatientID IN(SELECT SK_PatientID FROM ceg.GenomicsData WHERE CCG = '08V')

)as HDL_CholesterolEarliest

WHERE rn = 1

GO


---------------------- HDL cholesterol +/- 6 month window around  T1 or T2 diagnosis -------------------------------------

SELECT
	 [SK_PatientID]
	,[ClinicalCode] HDL_Cholesterol6MonthsWindowCode
	,[EventDate] HDL_Cholesterol6MonthsWindowDate
	,[Value] HDL_Cholesterol6MonthsWindowScore
	,[Units] HDL_Cholesterol6MonthsWindowUnits

INTO  #HDL_Cholesterol6MonthsWindow

FROM
(
	SELECT
		 E.[SK_PatientID]
		,[ClinicalCode]
		,[EventDate]
		,[Value]
		,[Units]
		,row_number() over(partition by E.[SK_PatientID] order by [EventDate] ASC) as rn

	FROM  [08V].[GPEncounter] E
	LEFT JOIN #DiabetesT1QoF AS T1 ON T1.SK_PatientID = E.SK_PatientID
	LEFT JOIN #DiabetesT2QoF AS T2 ON T2.SK_PatientID = E.SK_PatientID

	WHERE
		[ClinicalCode] COLLATE Latin1_General_CS_AS = '44P5'
		AND
		[Value] <> 0
		AND
		(
			T1.SK_PatientID IS NOT NULL
			OR
			T2.SK_PatientID IS NOT NULL
		)
		AND
		(
			[EventDate] BETWEEN DATEADD(month, -6, T1.DiabetesT1DateRecorded) AND DATEADD(month, 6, T1.DiabetesT1DateRecorded)
			OR
			[EventDate] BETWEEN DATEADD(month, -6, T2.DiabetesT2DateRecorded) AND DATEADD(month, 6, T2.DiabetesT2DateRecorded)
		)
		AND
		E.SK_PatientID IN(SELECT SK_PatientID FROM ceg.GenomicsData WHERE CCG = '08V')

) AS HDL_Cholesterol6MonthsWindow

WHERE rn = 1

GO


---------------------- HDL cholesterol Latest ----------------------------------------------------------------------------

SELECT
	 [SK_PatientID]
	,[ClinicalCode] HDL_CholesterolLatestCode
	,[EventDate] HDL_CholesterolLatestDateRecorded
	,[Value] HDL_CholesterolLatestValue
	,[Units] HDL_CholesterolLatestUnits

INTO  #HDL_CholesterolLatest

FROM
(
	SELECT
		 [SK_PatientID]
		,[ClinicalCode]
		,[EventDate]
		,[Value]
		,[Units]
		,row_number() over(partition by [SK_PatientID] order by [EventDate] DESC) as rn

	FROM  [08V].[GPEncounter]
	WHERE

		[ClinicalCode] COLLATE Latin1_General_CS_AS = '44P5'
		AND
		[Value] <> 0
		AND
		SK_PatientID IN(SELECT SK_PatientID FROM ceg.GenomicsData WHERE CCG = '08V')

)as HDL_CholesterolLatest


WHERE rn = 1

GO


------------------Serum LDL cholesterol (LDL-C) ----------------------------------------------------------------------------------
---------------------- Serum LDL Cholesterol Earliest ----------------------------------------------------------------------------

SELECT
	 [SK_PatientID]
	,[ClinicalCode] SerumLDL_CholesterolEarliestCode
	,[EventDate] SerumLDL_CholesterolEarliestDateRecorded
	,[Value] SerumLDL_CholesterolEarliestValue
	,[Units] SerumLDL_CholesterolEarliestUnits

INTO  #SerumLDL_CholesterolEarliest

FROM
(
	SELECT

		 [SK_PatientID]
		,[ClinicalCode]
		,[EventDate]
		,[Value]
		,[Units]
		,row_number() over(partition by [SK_PatientID] order by [EventDate] ASC) as rn

	FROM  [08V].[GPEncounter]
	WHERE

		[ClinicalCode] COLLATE Latin1_General_CS_AS = '44P6'
		AND
		[Value] <> 0
		AND
		SK_PatientID IN(SELECT SK_PatientID FROM ceg.GenomicsData WHERE CCG = '08V')

)as SerumLDL_CholesterolEarliest

WHERE rn = 1

GO


---------------------- Serum LDL Cholesterol +/- 6 month window around  T1 or T2 diagnosis -------------------------------------

SELECT
	 [SK_PatientID]
	,[ClinicalCode] SerumLDL_Cholesterol6MonthsWindowCode
	,[EventDate] SerumLDL_Cholesterol6MonthsWindowDate
	,[Value] SerumLDL_Cholesterol6MonthsWindowScore
	,[Units] SerumLDL_Cholesterol6MonthsWindowUnits

INTO  #SerumLDL_Cholesterol6MonthsWindow

FROM
(
	SELECT
		 E.[SK_PatientID]
		,[ClinicalCode]
		,[EventDate]
		,[Value]
		,[Units]
		,row_number() over(partition by E.[SK_PatientID] order by [EventDate] ASC) as rn

	FROM  [08V].[GPEncounter] E
	LEFT JOIN #DiabetesT1QoF AS T1 ON T1.SK_PatientID = E.SK_PatientID
	LEFT JOIN #DiabetesT2QoF AS T2 ON T2.SK_PatientID = E.SK_PatientID

	WHERE
		[ClinicalCode] COLLATE Latin1_General_CS_AS = '44P6'
		AND
		[Value] <> 0
		AND
		(
			T1.SK_PatientID IS NOT NULL
			OR
			T2.SK_PatientID IS NOT NULL
		)
		AND
		(
			[EventDate] BETWEEN DATEADD(month, -6, T1.DiabetesT1DateRecorded) AND DATEADD(month, 6, T1.DiabetesT1DateRecorded)
			OR
			[EventDate] BETWEEN DATEADD(month, -6, T2.DiabetesT2DateRecorded) AND DATEADD(month, 6, T2.DiabetesT2DateRecorded)
		)
		AND
		E.SK_PatientID IN(SELECT SK_PatientID FROM ceg.GenomicsData WHERE CCG = '08V')

) AS SerumLDL_Cholesterol6MonthsWindow

WHERE rn = 1

GO


---------------------- Serum LDL Cholesterol Latest ----------------------------------------------------------------------------

SELECT
	 [SK_PatientID]
	,[ClinicalCode] SerumLDL_CholesterolLatestCode
	,[EventDate] SerumLDL_CholesterolLatestDateRecorded
	,[Value] SerumLDL_CholesterolLatestValue
	,[Units] SerumLDL_CholesterolLatestUnits

INTO  #SerumLDL_CholesterolLatest

FROM
(
	SELECT
		 [SK_PatientID]
		,[ClinicalCode]
		,[EventDate]
		,[Value]
		,[Units]
		,row_number() over(partition by [SK_PatientID] order by [EventDate] DESC) as rn

	FROM  [08V].[GPEncounter]
	WHERE

		[ClinicalCode] COLLATE Latin1_General_CS_AS = '44P6'
		AND
		[Value] <> 0
		AND
		SK_PatientID IN(SELECT SK_PatientID FROM ceg.GenomicsData WHERE CCG = '08V')

)as SerumLDL_CholesterolLatest


WHERE rn = 1

GO


------------------ Serum triglycerides ----------------------------------------------------------------------------------------------
---------------------- Serum triglycerides Earliest ----------------------------------------------------------------------------

SELECT
	 [SK_PatientID]
	,[ClinicalCode] SerumTriglyceridesEarliestCode
	,[EventDate] SerumTriglyceridesEarliestDateRecorded
	,[Value] SerumTriglyceridesEarliestValue
	,[Units] SerumTriglyceridesEarliestUnits

INTO  #SerumTriglyceridesEarliest

FROM
(
	SELECT

		 [SK_PatientID]
		,[ClinicalCode]
		,[EventDate]
		,[Value]
		,[Units]
		,row_number() over(partition by [SK_PatientID] order by [EventDate] ASC) as rn

	FROM  [08V].[GPEncounter]
	WHERE

		[ClinicalCode] COLLATE Latin1_General_CS_AS LIKE '44Q%'
		AND
		[Value] <> 0
		AND
		SK_PatientID IN(SELECT SK_PatientID FROM ceg.GenomicsData WHERE CCG = '08V')

)as SerumTriglyEarliest

WHERE rn = 1

GO

---------------------- Serum triglycerides +/- 6 month window around  T1 or T2 diagnosis -------------------------------------

SELECT
	 [SK_PatientID]
	,[ClinicalCode] SerumTriglycerides6MonthsWindowCode
	,[EventDate] SerumTriglycerides6MonthsWindowDate
	,[Value] SerumTriglycerides6MonthsWindowScore
	,[Units] SerumTriglycerides6MonthsWindowUnits

INTO  #SerumTriglycerides6MonthsWindow

FROM
(
	SELECT
		 E.[SK_PatientID]
		,[ClinicalCode]
		,[EventDate]
		,[Value]
		,[Units]
		,row_number() over(partition by E.[SK_PatientID] order by [EventDate] ASC) as rn

	FROM  [08V].[GPEncounter] E
	LEFT JOIN #DiabetesT1QoF AS T1 ON T1.SK_PatientID = E.SK_PatientID
	LEFT JOIN #DiabetesT2QoF AS T2 ON T2.SK_PatientID = E.SK_PatientID

	WHERE
		[ClinicalCode] COLLATE Latin1_General_CS_AS LIKE '44Q%'
		AND
		[Value] <> 0
		AND
		(
			T1.SK_PatientID IS NOT NULL
			OR
			T2.SK_PatientID IS NOT NULL
		)
		AND
		(
			[EventDate] BETWEEN DATEADD(month, -6, T1.DiabetesT1DateRecorded) AND DATEADD(month, 6, T1.DiabetesT1DateRecorded)
			OR
			[EventDate] BETWEEN DATEADD(month, -6, T2.DiabetesT2DateRecorded) AND DATEADD(month, 6, T2.DiabetesT2DateRecorded)
		)
		AND
		E.SK_PatientID IN(SELECT SK_PatientID FROM ceg.GenomicsData WHERE CCG = '08V')

) AS SerumTriglycerides6MonthsWindow

WHERE rn = 1

GO


---------------------- Serum triglycerides Latest ----------------------------------------------------------------------------

SELECT
	 [SK_PatientID]
	,[ClinicalCode] SerumTriglyceridesLatestCode
	,[EventDate] SerumTriglyceridesLatestDateRecorded
	,[Value] SerumTriglyceridesLatestValue
	,[Units] SerumTriglyceridesLatestUnits

INTO  #SerumTriglyceridesLatest

FROM
(
	SELECT
		 [SK_PatientID]
		,[ClinicalCode]
		,[EventDate]
		,[Value]
		,[Units]
		,row_number() over(partition by [SK_PatientID] order by [EventDate] DESC) as rn

	FROM  [08V].[GPEncounter]
	WHERE

		[ClinicalCode] COLLATE Latin1_General_CS_AS LIKE '44Q%'
		AND
		[Value] <> 0
		AND
		SK_PatientID IN(SELECT SK_PatientID FROM ceg.GenomicsData WHERE CCG = '08V')

)as SerumTriglyLatest


WHERE rn = 1

GO


------------------Urine protein/creatinine ratio earliest ever ----------------------------------------------------------------------------------------------

SELECT
	 [SK_PatientID]
	,[ClinicalCode] UrineProteinEarliestCode
	,[EventDate] UrineProteinEarliestDateRecorded
	,[Value] UrineProteinEarliestValue
	,[Units] UrineProteinEarliestUnits

INTO  #UrineProteinEarliest

FROM
(
	SELECT

		 [SK_PatientID]
		,[ClinicalCode]
		,[EventDate]
		,[Value]
		,[Units]
		,row_number() over(partition by [SK_PatientID] order by [EventDate] ASC) as rn

	FROM  [08V].[GPEncounter]
	WHERE

		[ClinicalCode] COLLATE Latin1_General_CS_AS = '44lD'
		AND
		[Value] <> 0
		AND
		SK_PatientID IN(SELECT SK_PatientID FROM ceg.GenomicsData WHERE CCG = '08V')

)as UrineProteinEarliest

WHERE rn = 1

GO


------------------Urine protein/creatinine ratio latest ever -----------------------------------------------------------------------------------------------------

SELECT
	 [SK_PatientID]
	,[ClinicalCode] UrineProteinLatestCode
	,[EventDate] UrineProteinLatestDateRecorded
	,[Value] UrineProteinLatestValue
	,[Units] UrineProteinLatestUnits

INTO  #UrineProteinLatest

FROM
(
	SELECT
		 [SK_PatientID]
		,[ClinicalCode]
		,[EventDate]
		,[Value]
		,[Units]
		,row_number() over(partition by [SK_PatientID] order by [EventDate] DESC) as rn

	FROM  [08V].[GPEncounter]
	WHERE

		[ClinicalCode] COLLATE Latin1_General_CS_AS = '44lD'
		AND
		[Value] <> 0
		AND
		SK_PatientID IN(SELECT SK_PatientID FROM ceg.GenomicsData WHERE CCG = '08V')

)as UrineProteinLatest

WHERE rn = 1

GO


------------------Urine albumin/creatinine ratio earliest ever ----------------------------------------------------------------------------------------------

SELECT
	 [SK_PatientID]
	,[ClinicalCode] CreatininerRatioEarliestCode
	,[EventDate] CreatininerRatioEarliestDateRecorded
	,[Value] CreatininerRatioEarliestValue
	,[Units] CreatininerRatioEarliestUnits

INTO  #CreatininerRatioEarliest

FROM
(
	SELECT

		 [SK_PatientID]
		,[ClinicalCode]
		,[EventDate]
		,[Value]
		,[Units]
		,row_number() over(partition by [SK_PatientID] order by [EventDate] ASC) as rn

	FROM  [08V].[GPEncounter]
	WHERE

		[ClinicalCode] COLLATE Latin1_General_CS_AS = '46TC'
		AND
		[Value] <> 0
		AND
		SK_PatientID IN(SELECT SK_PatientID FROM ceg.GenomicsData WHERE CCG = '08V')

)as CreatininerRatioEarliest

WHERE rn = 1

GO


------------------Urine albumin/creatinine ratio latest ever -----------------------------------------------------------------------------------------------------

SELECT
	 [SK_PatientID]
	,[ClinicalCode] CreatininerRatioLatestCode
	,[EventDate] CreatininerRatioLatestDateRecorded
	,[Value] CreatininerRatioLatestValue
	,[Units] CreatininerRatioLatestUnits

INTO  #CreatininerRatioLatest

FROM
(
	SELECT
		 [SK_PatientID]
		,[ClinicalCode]
		,[EventDate]
		,[Value]
		,[Units]
		,row_number() over(partition by [SK_PatientID] order by [EventDate] DESC) as rn

	FROM  [08V].[GPEncounter]
	WHERE

		[ClinicalCode] COLLATE Latin1_General_CS_AS = '46TC'
		AND
		[Value] <> 0
		AND
		SK_PatientID IN(SELECT SK_PatientID FROM ceg.GenomicsData WHERE CCG = '08V')

)as CreatininerRatioLatest

WHERE rn = 1

GO


------------------Urine Albumin earliest ever ----------------------------------------------------------------------------------------------

SELECT
	 [SK_PatientID]
	,[ClinicalCode] UrineAlbuminEarliestCode
	,[EventDate] UrineAlbuminEarliestDateRecorded
	,[Value] UrineAlbuminEarliestValue
	,[Units] UrineAlbuminEarliestUnits

INTO  #UrineAlbuminEarliest

FROM
(
	SELECT

		 [SK_PatientID]
		,[ClinicalCode]
		,[EventDate]
		,[Value]
		,[Units]
		,row_number() over(partition by [SK_PatientID] order by [EventDate] ASC) as rn

	FROM  [08V].[GPEncounter]
	WHERE

		[ClinicalCode] COLLATE Latin1_General_CS_AS = '46N4'
		AND
		[Value] <> 0
		AND
		SK_PatientID IN(SELECT SK_PatientID FROM ceg.GenomicsData WHERE CCG = '08V')

)as UrineAlbuminEarliest

WHERE rn = 1

GO


------------------Urine Albumin latest ever -----------------------------------------------------------------------------------------------------

SELECT
	 [SK_PatientID]
	,[ClinicalCode] UrineAlbuminLatestCode
	,[EventDate] UrineAlbuminLatestDateRecorded
	,[Value] UrineAlbuminLatestValue
	,[Units] UrineAlbuminLatestUnits

INTO  #UrineAlbuminLatest

FROM
(
	SELECT
		 [SK_PatientID]
		,[ClinicalCode]
		,[EventDate]
		,[Value]
		,[Units]
		,row_number() over(partition by [SK_PatientID] order by [EventDate] DESC) as rn

	FROM  [08V].[GPEncounter]
	WHERE

		[ClinicalCode] COLLATE Latin1_General_CS_AS = '46N4'
		AND
		[Value] <> 0
		AND
		SK_PatientID IN(SELECT SK_PatientID FROM ceg.GenomicsData WHERE CCG = '08V')

)as UrineAlbuminLatest

WHERE rn = 1

GO


------------------Systolic Blood Pressure earliest ever ----------------------------------------------------------------------------------------------

SELECT
	 [SK_PatientID]
	,[ClinicalCode] SysBPEarliestCode
	,[EventDate] SysBPEarliestDateRecorded
	,[Value] SysBPEarliestValue
	,[Units] SysBPEarliestUnits

INTO  #SysBPEarliest

FROM
(
	SELECT

		 [SK_PatientID]
		,[ClinicalCode]
		,[EventDate]
		,[Value]
		,[Units]
		,row_number() over(partition by [SK_PatientID] order by [EventDate] ASC) as rn

	FROM  [08V].[GPEncounter]
	WHERE

		[ClinicalCode] COLLATE Latin1_General_CS_AS = '2469'
		AND
		[Value] <> 0
		AND
		SK_PatientID IN(SELECT SK_PatientID FROM ceg.GenomicsData WHERE CCG = '08V')

)as SysBPEarliest

WHERE rn = 1

GO


------------------Systolic Blood Pressure latest ever -----------------------------------------------------------------------------------------------------


SELECT
	 [SK_PatientID]
	,[ClinicalCode] SysBPLatestCode
	,[EventDate] SysBPLatestDateRecorded
	,[Value] SysBPLatestValue
	,[Units] SysBPLatestUnits

INTO  #SysBPLatest

FROM
(
	SELECT
		 [SK_PatientID]
		,[ClinicalCode]
		,[EventDate]
		,[Value]
		,[Units]
		,row_number() over(partition by [SK_PatientID] order by [EventDate] DESC) as rn

	FROM  [08V].[GPEncounter]
	WHERE

		[ClinicalCode] COLLATE Latin1_General_CS_AS = '2469'
		AND
		[Value] <> 0
		AND
		SK_PatientID IN(SELECT SK_PatientID FROM ceg.GenomicsData WHERE CCG = '08V')

)as SysBPLatest


WHERE rn = 1

GO


------------------Diastolic Blood Pressure earliest ever ----------------------------------------------------------------------------------------------

SELECT
	 [SK_PatientID]
	,[ClinicalCode] DiaBPEarliestCode
	,[EventDate] DiaBPEarliestDateRecorded
	,[Value] DiaBPEarliestValue
	,[Units] DiaBPEarliestUnits

INTO  #DiaBPEarliest

FROM
(
	SELECT

		 [SK_PatientID]
		,[ClinicalCode]
		,[EventDate]
		,[Value]
		,[Units]
		,row_number() over(partition by [SK_PatientID] order by [EventDate] ASC) as rn

	FROM  [08V].[GPEncounter]
	WHERE

		[ClinicalCode] COLLATE Latin1_General_CS_AS = '246A'
		AND
		[Value] <> 0
		AND
		SK_PatientID IN(SELECT SK_PatientID FROM ceg.GenomicsData WHERE CCG = '08V')

)as DiaBPEarliest

WHERE rn = 1

GO


------------------Diastolic Blood Pressure latest ever -----------------------------------------------------------------------------------------------------

SELECT
	 [SK_PatientID]
	,[ClinicalCode] DiaBPLatestCode
	,[EventDate] DiaBPLatestDateRecorded
	,[Value] DiaBPLatestValue
	,[Units] DiaBPLatestUnits

INTO  #DiaBPLatest

FROM
(
	SELECT
		 [SK_PatientID]
		,[ClinicalCode]
		,[EventDate]
		,[Value]
		,[Units]
		,row_number() over(partition by [SK_PatientID] order by [EventDate] DESC) as rn

	FROM  [08V].[GPEncounter]
	WHERE

		[ClinicalCode] COLLATE Latin1_General_CS_AS = '246A'
		AND
		[Value] <> 0
		AND
		SK_PatientID IN(SELECT SK_PatientID FROM ceg.GenomicsData WHERE CCG = '08V')

)as DiaBPLatest


WHERE rn = 1

GO


------------------Serum Creatinine earliest ever ----------------------------------------------------------------------------------------------

SELECT
	 [SK_PatientID]
	,[ClinicalCode] CreatinineEarliestCode
	,[EventDate] CreatinineEarliestDateRecorded
	,[Value] CreatinineEarliestValue
	,[Units] CreatinineEarliestUnits

INTO  #CreatinineEarliest

FROM
(
	SELECT

		 [SK_PatientID]
		,[ClinicalCode]
		,[EventDate]
		,[Value]
		,[Units]
		,row_number() over(partition by [SK_PatientID] order by [EventDate] ASC) as rn

	FROM  [08V].[GPEncounter]
	WHERE

		[ClinicalCode] COLLATE Latin1_General_CS_AS = '44J3'
		AND
		[Value] <> 0
		AND
		SK_PatientID IN(SELECT SK_PatientID FROM ceg.GenomicsData WHERE CCG = '08V')

)as CreatinineEarliest

WHERE rn = 1

GO


------------------Serum Creatinine latest ever -----------------------------------------------------------------------------------------------------

SELECT
	 [SK_PatientID]
	,[ClinicalCode] CreatinineLatestCode
	,[EventDate] CreatinineLatestDateRecorded
	,[Value] CreatinineLatestValue
	,[Units] CreatinineLatestUnits

INTO  #CreatinineLatest

FROM
(
	SELECT

		 [SK_PatientID]
		,[ClinicalCode]
		,[EventDate]
		,[Value]
		,[Units]
		,row_number() over(partition by [SK_PatientID] order by [EventDate] Desc) as rn

	FROM  [08V].[GPEncounter]
	WHERE

		[ClinicalCode] COLLATE Latin1_General_CS_AS = '44J3'
		AND
		[Value] <> 0
		AND
		SK_PatientID IN(SELECT SK_PatientID FROM ceg.GenomicsData WHERE CCG = '08V')

)as CreatinineLatest

WHERE rn = 1

GO


------------------ eGFR earliest ever ----------------------------------------------------------------------------------------------

SELECT
	 [SK_PatientID]
	,[ClinicalCode] eGFR_EarliestCode
	,[EventDate] eGFR_EarliestDateRecorded
	,[Value] eGFR_EarliestValue
	,[Units] eGFR_EarliestUnits

INTO  #eGFR_Earliest

FROM
(
	SELECT

		 [SK_PatientID]
		,[ClinicalCode]
		,[EventDate]
		,[Value]
		,[Units]
		,row_number() over(partition by [SK_PatientID] order by [EventDate] ASC) as rn

	FROM  [08V].[GPEncounter]
	WHERE

		[ClinicalCode] COLLATE Latin1_General_CS_AS = '451E'
		AND
		[Value] <> 0
		AND
		SK_PatientID IN(SELECT SK_PatientID FROM ceg.GenomicsData WHERE CCG = '08V')

)as eGFR_Earliest

WHERE rn = 1

GO


------------------ eGFR latest ever -----------------------------------------------------------------------------------------------------

SELECT
	 [SK_PatientID]
	,[ClinicalCode] eGFR_LatestCode
	,[EventDate] eGFR_LatestDateRecorded
	,[Value] eGFR_LatestValue
	,[Units] eGFR_LatestUnits

INTO  #eGFR_Latest

FROM
(
	SELECT

		 [SK_PatientID]
		,[ClinicalCode]
		,[EventDate]
		,[Value]
		,[Units]
		,row_number() over(partition by [SK_PatientID] order by [EventDate] DESC) as rn

	FROM  [08V].[GPEncounter]
	WHERE

		[ClinicalCode] COLLATE Latin1_General_CS_AS = '451E'
		AND
		[Value] <> 0
		AND
		SK_PatientID IN(SELECT SK_PatientID FROM ceg.GenomicsData WHERE CCG = '08V')

)as eGFR_Latest

WHERE rn = 1

GO


------------------Glutamic acid decarboxylase antibody level (GAD)  latest ever -------------------------------------------------------------

SELECT
	 [SK_PatientID]
	,[ClinicalCode] GAD_Code
	,[EventDate] GAD_DateRecorded
	,[Value] GAD_Value
	,[Units] GAD_Units

INTO  #GAD

FROM
(
	SELECT

		 [SK_PatientID]
		,[ClinicalCode]
		,[EventDate]
		,[Value]
		,[Units]
		,row_number() over(partition by [SK_PatientID] order by [EventDate] DESC) as rn

	FROM  [08V].[GPEncounter]
	WHERE

		[ClinicalCode] COLLATE Latin1_General_CS_AS = '43m8'
		AND
		[Value] <> 0
		AND
		SK_PatientID IN(SELECT SK_PatientID FROM ceg.GenomicsData WHERE CCG = '08V')

)as GAD

WHERE rn = 1

GO


------------------ Islet cell antibody level latest ever --------------------------------------------------------------------------

SELECT
	 [SK_PatientID]
	,[ClinicalCode] IsletCellCode
	,[EventDate] IsletCellDateRecorded
	,[Value] IsletCellValue
	,[Units] IsletCellUnits

INTO  #IsletCell

FROM
(
	SELECT

		 [SK_PatientID]
		,[ClinicalCode]
		,[EventDate]
		,[Value]
		,[Units]
		,row_number() over(partition by [SK_PatientID] order by [EventDate] DESC) as rn

	FROM  [08V].[GPEncounter]
	WHERE

		[ClinicalCode] COLLATE Latin1_General_CS_AS = '43a3'
		AND
		[Value] <> 0
		AND
		SK_PatientID IN(SELECT SK_PatientID FROM ceg.GenomicsData WHERE CCG = '08V')

)as IsletCell

WHERE rn = 1

GO


------------------ Insulinoma-associated antigen-2 antibody level (IA2) latest ever --------------------------------------------------------

SELECT
	 [SK_PatientID]
	,[ClinicalCode] IA2_Code
	,[EventDate] IA2_DateRecorded
	,[Value] IA2_Value
	,[Units] IA2_Units

INTO  #IA2

FROM
(
	SELECT

		 [SK_PatientID]
		,[ClinicalCode]
		,[EventDate]
		,[Value]
		,[Units]
		,row_number() over(partition by [SK_PatientID] order by [EventDate] DESC) as rn

	FROM  [08V].[GPEncounter]
	WHERE

		[ClinicalCode] COLLATE Latin1_General_CS_AS = '43aw'
		AND
		[Value] <> 0
		AND
		SK_PatientID IN(SELECT SK_PatientID FROM ceg.GenomicsData WHERE CCG = '08V')

)as IA2

WHERE rn = 1

GO


------------------ Coeliac Autoantibody Profile latest ever --------------------------------------------------------------------------

SELECT
	 [SK_PatientID]
	,[ClinicalCode] CoeliacAutoantibodyCode
	,[EventDate] CoeliacAutoantibodyDateRecorded

INTO  #CoeliacAutoantibody

FROM
(
	SELECT

		 [SK_PatientID]
		,[ClinicalCode]
		,[EventDate]
		,row_number() over(partition by [SK_PatientID] order by [EventDate] DESC) as rn

	FROM  [08V].[GPEncounter]
	WHERE

		[ClinicalCode] COLLATE Latin1_General_CS_AS IN ('68W3' , '68W4')
		AND
		SK_PatientID IN(SELECT SK_PatientID FROM ceg.GenomicsData WHERE CCG = '08V')

)as CoeliacAutoantibody

WHERE rn = 1

GO


------------------ Anti-thyroid peroxidase latest ever --------------------------------------------------------------------------

SELECT
	 [SK_PatientID]
	,[ClinicalCode] PeroxidaseCode
	,[EventDate] PeroxidaseDateRecorded
	,[Value] PeroxidaseValue
	,[Units] PeroxidaseUnits

INTO  #Peroxidase

FROM
(
	SELECT

		 [SK_PatientID]
		,[ClinicalCode]
		,[EventDate]
		,[Value]
		,[Units]
		,row_number() over(partition by [SK_PatientID] order by [EventDate] DESC) as rn

	FROM  [08V].[GPEncounter]
	WHERE

		[ClinicalCode] COLLATE Latin1_General_CS_AS LIKE '43Gd%'
		AND
		[Value] <> 0
		AND
		SK_PatientID IN(SELECT SK_PatientID FROM ceg.GenomicsData WHERE CCG = '08V')

)as Peroxidase

WHERE rn = 1

GO


------------------ C - peptide latest ever --------------------------------------------------------------------------

SELECT
	 [SK_PatientID]
	,[ClinicalCode] CpeptideCode
	,[EventDate] CpeptideDateRecorded
	,[Value] CpeptideValue
	,[Units] CpeptideUnits

INTO  #Cpeptide

FROM
(
	SELECT

		 [SK_PatientID]
		,[ClinicalCode]
		,[EventDate]
		,[Value]
		,[Units]
		,row_number() over(partition by [SK_PatientID] order by [EventDate] DESC) as rn

	FROM  [08V].[GPEncounter]
	WHERE

		[ClinicalCode] COLLATE Latin1_General_CS_AS LIKE '44Za%'
		AND
		[Value] <> 0
		AND
		SK_PatientID IN(SELECT SK_PatientID FROM ceg.GenomicsData WHERE CCG = '08V')

)as Cpeptide

WHERE rn = 1

GO


------------------ Serum insulin latest ever --------------------------------------------------------------------------

SELECT
	 [SK_PatientID]
	,[ClinicalCode] SerumInsulinCode
	,[EventDate] SerumInsulinDateRecorded
	,[Value] SerumInsulinValue
	,[Units] SerumInsulinUnits

INTO  #SerumInsulin

FROM
(
	SELECT

		 [SK_PatientID]
		,[ClinicalCode]
		,[EventDate]
		,[Value]
		,[Units]
		,row_number() over(partition by [SK_PatientID] order by [EventDate] DESC) as rn

	FROM  [08V].[GPEncounter]
	WHERE

		[ClinicalCode] COLLATE Latin1_General_CS_AS = '4493'
		AND
		[Value] <> 0
		AND
		SK_PatientID IN(SELECT SK_PatientID FROM ceg.GenomicsData WHERE CCG = '08V')

)as SerumInsulin

WHERE rn = 1

GO


------------------  Serum (ALT) alanine aminotransferase level latest ever --------------------------------------------------------------------------

SELECT
	 [SK_PatientID]
	,[ClinicalCode] SerumALT_Code
	,[EventDate]	SerumALT_DateRecorded
	,[Value]		SerumALT_Value
	,[Units]		SerumALT_Units

INTO  #SerumALT

FROM
(
	SELECT
		 [SK_PatientID]
		,[ClinicalCode]
		,[EventDate]
		,[Value]
		,[Units]
		,row_number() over(partition by [SK_PatientID] order by [EventDate] DESC) as rn

	FROM  [08V].[GPEncounter]
	WHERE

		[ClinicalCode] COLLATE Latin1_General_CS_AS = '44GB'
		AND
		[Value] <> 0
		AND
		SK_PatientID IN(SELECT SK_PatientID FROM ceg.GenomicsData WHERE CCG = '08V')

)as SerumALT

WHERE rn = 1

GO


------------------ AST serum level latest ever --------------------------------------------------------------------------

SELECT
	 [SK_PatientID]
	,[ClinicalCode] AST_SerumCode
	,[EventDate]	AST_SerumDateRecorded
	,[Value]		AST_SerumValue
	,[Units]		AST_SerumUnits

INTO  #AST_Serum

FROM
(
	SELECT
		 [SK_PatientID]
		,[ClinicalCode]
		,[EventDate]
		,[Value]
		,[Units]
		,row_number() over(partition by [SK_PatientID] order by [EventDate] DESC) as rn

	FROM  [08V].[GPEncounter]
	WHERE

		[ClinicalCode] COLLATE Latin1_General_CS_AS IN('44HB', '44HB-1')
		AND
		[Value] <> 0
		AND
		SK_PatientID IN(SELECT SK_PatientID FROM ceg.GenomicsData WHERE CCG = '08V')

)as AST_Serum

WHERE rn = 1

GO

------------------ Thyroid function test latest ever --------------------------------------------------------------------------

SELECT
	 [SK_PatientID]
	,[ClinicalCode] TSH_T4Code
	,[EventDate] TSH_T4DateRecorded
	,[Value]	 TSH_T4Value
	,[Units]	 TSH_T4Units

INTO  #TSH_T4

FROM
(
	SELECT
		 [SK_PatientID]
		,[ClinicalCode]
		,[EventDate]
		,[Value]
		,[Units]
		,row_number() over(partition by [SK_PatientID] order by [EventDate] DESC) as rn

	FROM  [08V].[GPEncounter]
	WHERE
		(
			[ClinicalCode] COLLATE Latin1_General_CS_AS LIKE '442A%'
			OR
			[ClinicalCode] COLLATE Latin1_General_CS_AS LIKE '4426%'
		)
		AND
		[Value] <> 0
		AND
		SK_PatientID IN(SELECT SK_PatientID FROM ceg.GenomicsData WHERE CCG = '08V')

)as TSH_T4

WHERE rn = 1

GO


--------------------------------------------------------------------------------------------------------------------------
---------------------------------------- Joining Tables ------------------------------------------------------------------
--------------------------------------------------------------------------------------------------------------------------


SELECT

	(Select CommissionerName From Dictionary.[dbo].[Commissioner] Where CommissionerCode = '08V') AS Locality
	,P.EncryptedNHSNumber

	,P.SK_PatientID


	-- CLINIC DATA
	,CurrentSmokerQoF.CurrentSmokerQoFCode
	,CurrentSmokerQoF.CurrentSmokerQoFDateRecorded

	,ExSmokerQoF.ExSmokerQoFCode
	,ExSmokerQoF.ExSmokerQoFDateRecorded

	,NeverSmokedQoF.NeverSmokedQoFCode
	,NeverSmokedQoF.NeverSmokedQoFDateRecorded

	,WeightEarliest.WeightEarliestCode
	,WeightEarliest.WeightEarliestDate
	,WeightEarliest.WeightEarliestScore
	,WeightEarliest.WeightEarliestUnits
	,WeightEarliest.WeightEarliestAgeAtEvent

	,Weight6MonthsWindow.Weight6MonthsWindowCode
	,Weight6MonthsWindow.Weight6MonthsWindowDate
	,Weight6MonthsWindow.Weight6MonthsWindowScore
	,Weight6MonthsWindow.Weight6MonthsWindowUnits
	,Weight6MonthsWindow.Weight6MonthsWindowAgeAtEvent

	,WeightLatest.WeightLatestCode
	,WeightLatest.WeightLatestDate
	,WeightLatest.WeightLatestScore
	,WeightLatest.WeightLatestUnits
	,WeightLatest.WeightLatestAgeAtEvent

	--,WeightHighestEver.WeightHighestEverCode
	--,WeightHighestEver.WeightHighestEverDate
	--,WeightHighestEver.WeightHighestEverScore
	--,WeightHighestEver.WeightHighestEverUnits
	--,WeightHighestEver.WeightHighestEverAgeAtEvent

	,HeightEarliest.HeightEarliestCode
	,HeightEarliest.HeightEarliestDate
	,HeightEarliest.HeightEarliestScore
	,HeightEarliest.HeightEarliestUnits
	,HeightEarliest.HeightEarliestAgeAtEvent

	,Height6MonthsWindow.Height6MonthsWindowCode
	,Height6MonthsWindow.Height6MonthsWindowDate
	,Height6MonthsWindow.Height6MonthsWindowScore
	,Height6MonthsWindow.Height6MonthsWindowUnits
	,Height6MonthsWindow.Height6MonthsWindowAgeAtEvent

	,HeightLatest.HeightLatestCode
	,HeightLatest.HeightLatestDate
	,HeightLatest.HeightLatestScore
	,HeightLatest.HeightLatestUnits
	,HeightLatest.HeightLatestAgeAtEvent

	,BMIEarliest.BMIEarliestCode
	,BMIEarliest.BMIEarliestDate
	,BMIEarliest.BMIEarliestScore
	,BMIEarliest.BMIEarliestUnits
	,BMIEarliest.BMIEarliestAgeAtEvent

	,BMI6MonthsWindow.BMI6MonthsWindowCode
	,BMI6MonthsWindow.BMI6MonthsWindowDate
	,BMI6MonthsWindow.BMI6MonthsWindowScore
	,BMI6MonthsWindow.BMI6MonthsWindowUnits
	,BMI6MonthsWindow.BMI6MonthsWindowAgeAtEvent

	,BMILatest.BMILatestCode
	,BMILatest.BMILatestDate
	,BMILatest.BMILatestScore
	,BMILatest.BMILatestUnits
	,BMILatest.BMILatestAgeAtEvent

	--,BMIHighestEver.BMI_HighestEverCode
	--,BMIHighestEver.BMI_HighestEverDate
	--,BMIHighestEver.BMI_HighestEverScore
	--,BMIHighestEver.BMI_HighestEverUnits
	--,BMIHighestEver.BMI_HighestEverAgeAtEvent

	,HbA1cEarliest.HbA1cEarliestCode
	,HbA1cEarliest.HbA1cEarliestDateRecorded
	,HbA1cEarliest.HbA1cEarliestValue
	,HbA1cEarliest.HbA1cEarliestUnits

	,HbA1c6MonthsWindow.HbA1c6MonthsWindowCode
	,HbA1c6MonthsWindow.HbA1c6MonthsWindowDate
	,HbA1c6MonthsWindow.HbA1c6MonthsWindowScore
	,HbA1c6MonthsWindow.HbA1c6MonthsWindowUnits

	,HbA1cLatest.HbA1cLatestCode
	,HbA1cLatest.HbA1cLatestDateRecorded
	,HbA1cLatest.HbA1cLatestValue
	,HbA1cLatest.HbA1cLatestUnits

	--,HbA1cHighestEver.HbA1cHighestEverCode
	--,HbA1cHighestEver.HbA1cHighestEverDateRecorded
	--,HbA1cHighestEver.HbA1cHighestEverValue
	--,HbA1cHighestEver.HbA1cHighestEverUnits

	,CholesterolEarliest.CholesterolEarliestCode
	,CholesterolEarliest.CholesterolEarliestDateRecorded
	,CholesterolEarliest.CholesterolEarliestValue
	,CholesterolEarliest.CholesterolEarliestUnits

	,Cholesterol6MonthsWindow.Cholesterol6MonthsWindowCode
	,Cholesterol6MonthsWindow.Cholesterol6MonthsWindowDate
	,Cholesterol6MonthsWindow.Cholesterol6MonthsWindowScore
	,Cholesterol6MonthsWindow.Cholesterol6MonthsWindowUnits

	,CholesterolLatest.CholesterolLatestCode
	,CholesterolLatest.CholesterolLatestDateRecorded
	,CholesterolLatest.CholesterolLatestValue
	,CholesterolLatest.CholesterolLatestUnits

	--,CholesterolHighestEver.CholesterolHighestEverCode
	--,CholesterolHighestEver.CholesterolHighestEverDateRecorded
	--,CholesterolHighestEver.CholesterolHighestEverValue
	--,CholesterolHighestEver.CholesterolHighestEverUnits

	,HDL_CholesterolEarliest.HDL_CholesterolEarliestCode
	,HDL_CholesterolEarliest.HDL_CholesterolEarliestDateRecorded
	,HDL_CholesterolEarliest.HDL_CholesterolEarliestValue
	,HDL_CholesterolEarliest.HDL_CholesterolEarliestUnits

	,HDL_Cholesterol6MonthsWindow.HDL_Cholesterol6MonthsWindowCode
	,HDL_Cholesterol6MonthsWindow.HDL_Cholesterol6MonthsWindowDate
	,HDL_Cholesterol6MonthsWindow.HDL_Cholesterol6MonthsWindowScore
	,HDL_Cholesterol6MonthsWindow.HDL_Cholesterol6MonthsWindowUnits

	,HDL_CholesterolLatest.HDL_CholesterolLatestCode
	,HDL_CholesterolLatest.HDL_CholesterolLatestDateRecorded
	,HDL_CholesterolLatest.HDL_CholesterolLatestValue
	,HDL_CholesterolLatest.HDL_CholesterolLatestUnits

	--,HDL_CholesterolHighestEver.HDL_CholesterolHighestEverCode
	--,HDL_CholesterolHighestEver.HDL_CholesterolHighestEverDateRecorded
	--,HDL_CholesterolHighestEver.HDL_CholesterolHighestEverValue
	--,HDL_CholesterolHighestEver.HDL_CholesterolHighestEverUnits

	,SerumLDL_CholesterolEarliest.SerumLDL_CholesterolEarliestCode
	,SerumLDL_CholesterolEarliest.SerumLDL_CholesterolEarliestDateRecorded
	,SerumLDL_CholesterolEarliest.SerumLDL_CholesterolEarliestValue
	,SerumLDL_CholesterolEarliest.SerumLDL_CholesterolEarliestUnits

	,SerumLDL_Cholesterol6MonthsWindow.SerumLDL_Cholesterol6MonthsWindowCode
	,SerumLDL_Cholesterol6MonthsWindow.SerumLDL_Cholesterol6MonthsWindowDate
	,SerumLDL_Cholesterol6MonthsWindow.SerumLDL_Cholesterol6MonthsWindowScore
	,SerumLDL_Cholesterol6MonthsWindow.SerumLDL_Cholesterol6MonthsWindowUnits

	,SerumLDL_CholesterolLatest.SerumLDL_CholesterolLatestCode
	,SerumLDL_CholesterolLatest.SerumLDL_CholesterolLatestDateRecorded
	,SerumLDL_CholesterolLatest.SerumLDL_CholesterolLatestValue
	,SerumLDL_CholesterolLatest.SerumLDL_CholesterolLatestUnits

	--,SerumLDL_CholesterolHighestEver.SerumLDL_CholesterolHighestEverCode
	--,SerumLDL_CholesterolHighestEver.SerumLDL_CholesterolHighestEverDateRecorded
	--,SerumLDL_CholesterolHighestEver.SerumLDL_CholesterolHighestEverValue
	--,SerumLDL_CholesterolHighestEver.SerumLDL_CholesterolHighestEverUnits

	,SerumTriglyceridesEarliest.SerumTriglyceridesEarliestCode
	,SerumTriglyceridesEarliest.SerumTriglyceridesEarliestDateRecorded
	,SerumTriglyceridesEarliest.SerumTriglyceridesEarliestValue
	,SerumTriglyceridesEarliest.SerumTriglyceridesEarliestUnits

	,SerumTriglycerides6MonthsWindow.SerumTriglycerides6MonthsWindowCode
	,SerumTriglycerides6MonthsWindow.SerumTriglycerides6MonthsWindowDate
	,SerumTriglycerides6MonthsWindow.SerumTriglycerides6MonthsWindowScore
	,SerumTriglycerides6MonthsWindow.SerumTriglycerides6MonthsWindowUnits

	,SerumTriglyceridesLatest.SerumTriglyceridesLatestCode
	,SerumTriglyceridesLatest.SerumTriglyceridesLatestDateRecorded
	,SerumTriglyceridesLatest.SerumTriglyceridesLatestValue
	,SerumTriglyceridesLatest.SerumTriglyceridesLatestUnits

	--,SerumTriglyceridesHighestEver.SerumTriglyceridesHighestEverCode
	--,SerumTriglyceridesHighestEver.SerumTriglyceridesHighestEverDateRecorded
	--,SerumTriglyceridesHighestEver.SerumTriglyceridesHighestEverValue
	--,SerumTriglyceridesHighestEver.SerumTriglyceridesHighestEverUnits

	,UrineProteinEarliest.UrineProteinEarliestCode
	,UrineProteinEarliest.UrineProteinEarliestDateRecorded
	,UrineProteinEarliest.UrineProteinEarliestValue
	,UrineProteinEarliest.UrineProteinEarliestUnits

	,UrineProteinLatest.UrineProteinLatestCode
	,UrineProteinLatest.UrineProteinLatestDateRecorded
	,UrineProteinLatest.UrineProteinLatestValue
	,UrineProteinLatest.UrineProteinLatestUnits

	,CreatininerRatioEarliest.CreatininerRatioEarliestCode
	,CreatininerRatioEarliest.CreatininerRatioEarliestDateRecorded
	,CreatininerRatioEarliest.CreatininerRatioEarliestValue
	,CreatininerRatioEarliest.CreatininerRatioEarliestUnits

	,CreatininerRatioLatest.CreatininerRatioLatestCode
	,CreatininerRatioLatest.CreatininerRatioLatestDateRecorded
	,CreatininerRatioLatest.CreatininerRatioLatestValue
	,CreatininerRatioLatest.CreatininerRatioLatestUnits

	,UrineAlbuminEarliest.UrineAlbuminEarliestCode
	,UrineAlbuminEarliest.UrineAlbuminEarliestDateRecorded
	,UrineAlbuminEarliest.UrineAlbuminEarliestValue
	,UrineAlbuminEarliest.UrineAlbuminEarliestUnits

	,UrineAlbuminLatest.UrineAlbuminLatestCode
	,UrineAlbuminLatest.UrineAlbuminLatestDateRecorded
	,UrineAlbuminLatest.UrineAlbuminLatestValue
	,UrineAlbuminLatest.UrineAlbuminLatestUnits

	,SysBPEarliest.SysBPEarliestCode
	,SysBPEarliest.SysBPEarliestDateRecorded
	,SysBPEarliest.SysBPEarliestValue
	,SysBPEarliest.SysBPEarliestUnits

	,SysBPLatest.SysBPLatestCode
	,SysBPLatest.SysBPLatestDateRecorded
	,SysBPLatest.SysBPLatestValue
	,SysBPLatest.SysBPLatestUnits

	--,SysBPHighestEver.SysBPHighestEverCode
	--,SysBPHighestEver.SysBPHighestEverDateRecorded
	--,SysBPHighestEver.SysBPHighestEverValue
	--,SysBPHighestEver.SysBPHighestEverUnits

	,DiaBPEarliest.DiaBPEarliestCode
	,DiaBPEarliest.DiaBPEarliestDateRecorded
	,DiaBPEarliest.DiaBPEarliestValue
	,DiaBPEarliest.DiaBPEarliestUnits

	,DiaBPLatest.DiaBPLatestCode
	,DiaBPLatest.DiaBPLatestDateRecorded
	,DiaBPLatest.DiaBPLatestValue
	,DiaBPLatest.DiaBPLatestUnits

	--,DiaBPHighestEver.DiaBPHighestEverCode
	--,DiaBPHighestEver.DiaBPHighestEverDateRecorded
	--,DiaBPHighestEver.DiaBPHighestEverValue
	--,DiaBPHighestEver.DiaBPHighestEverUnits

	,CreatinineEarliest.CreatinineEarliestCode
	,CreatinineEarliest.CreatinineEarliestDateRecorded
	,CreatinineEarliest.CreatinineEarliestValue
	,CreatinineEarliest.CreatinineEarliestUnits

	,CreatinineLatest.CreatinineLatestCode
	,CreatinineLatest.CreatinineLatestDateRecorded
	,CreatinineLatest.CreatinineLatestValue
	,CreatinineLatest.CreatinineLatestUnits

	,eGFR_Earliest.eGFR_EarliestCode
	,eGFR_Earliest.eGFR_EarliestDateRecorded
	,eGFR_Earliest.eGFR_EarliestValue
	,eGFR_Earliest.eGFR_EarliestUnits

	,eGFR_Latest.eGFR_LatestCode
	,eGFR_Latest.eGFR_LatestDateRecorded
	,eGFR_Latest.eGFR_LatestValue
	,eGFR_Latest.eGFR_LatestUnits

	,GAD.GAD_Code
	,GAD.GAD_DateRecorded
	,GAD.GAD_Value
	,GAD.GAD_Units

	,IsletCell.IsletCellCode
	,IsletCell.IsletCellDateRecorded
	,IsletCell.IsletCellValue
	,IsletCell.IsletCellUnits

	,IA2.IA2_Code
	,IA2.IA2_DateRecorded
	,IA2.IA2_Value
	,IA2.IA2_Units

	,CoeliacAutoantibody.CoeliacAutoantibodyCode
	,CoeliacAutoantibody.CoeliacAutoantibodyDateRecorded

	,Peroxidase.PeroxidaseCode
	,Peroxidase.PeroxidaseDateRecorded
	,Peroxidase.PeroxidaseValue
	,Peroxidase.PeroxidaseUnits

	,Cpeptide.CpeptideCode
	,Cpeptide.CpeptideDateRecorded
	,Cpeptide.CpeptideValue
	,Cpeptide.CpeptideUnits

	,SerumInsulin.SerumInsulinCode
	,SerumInsulin.SerumInsulinDateRecorded
	,SerumInsulin.SerumInsulinValue
	,SerumInsulin.SerumInsulinUnits

	,SerumALT.SerumALT_Code
	,SerumALT.SerumALT_DateRecorded
	,SerumALT.SerumALT_Value
	,SerumALT.SerumALT_Units

	,AST_Serum.AST_SerumCode
	,AST_Serum.AST_SerumDateRecorded
	,AST_Serum.AST_SerumValue
	,AST_Serum.AST_SerumUnits

	,TSH_T4.TSH_T4Code
	,TSH_T4.TSH_T4DateRecorded
	,TSH_T4.TSH_T4Value
	,TSH_T4.TSH_T4Units

FROM
	CEG.ceg.GenomicsData P

	-- CLINIC DATA

	--LEFT JOIN  #Smoking AS Smoking ON Smoking.SK_PatientID = P.SK_PatientID

	LEFT JOIN  #CurrentSmokerQoF AS CurrentSmokerQoF ON CurrentSmokerQoF.SK_PatientID = P.SK_PatientID
	LEFT JOIN  #ExSmokerQoF AS ExSmokerQoF ON ExSmokerQoF.SK_PatientID = P.SK_PatientID
	LEFT JOIN  #NeverSmokedQoF AS NeverSmokedQoF ON NeverSmokedQoF.SK_PatientID = P.SK_PatientID

	LEFT JOIN  #WeightEarliest AS WeightEarliest ON WeightEarliest.SK_PatientID = P.SK_PatientID
	LEFT JOIN  #Weight6MonthsWindow AS Weight6MonthsWindow ON Weight6MonthsWindow.SK_PatientID = P.SK_PatientID
	LEFT JOIN  #WeightLatest AS WeightLatest ON WeightLatest.SK_PatientID = P.SK_PatientID
	--LEFT JOIN  #WeightHighestEver AS WeightHighestEver ON WeightHighestEver.SK_PatientID = P.SK_PatientID

	LEFT JOIN  #HeightEarliest AS HeightEarliest ON HeightEarliest.SK_PatientID = P.SK_PatientID
	LEFT JOIN  #Height6MonthsWindow AS Height6MonthsWindow ON Height6MonthsWindow.SK_PatientID = P.SK_PatientID
	LEFT JOIN  #HeightLatest AS HeightLatest ON HeightLatest.SK_PatientID = P.SK_PatientID

	LEFT JOIN  #BMIEarliest AS BMIEarliest ON BMIEarliest.SK_PatientID = P.SK_PatientID
	LEFT JOIN  #BMI6MonthsWindow AS BMI6MonthsWindow ON BMI6MonthsWindow.SK_PatientID = P.SK_PatientID
	LEFT JOIN  #BMILatest AS BMILatest ON BMILatest.SK_PatientID = P.SK_PatientID
	--LEFT JOIN  #BMIHighestEver AS BMIHighestEver ON BMIHighestEver.SK_PatientID = P.SK_PatientID

	LEFT JOIN  #HbA1cEarliest AS HbA1cEarliest ON HbA1cEarliest.SK_PatientID = P.SK_PatientID
	LEFT JOIN  #HbA1c6MonthsWindow AS HbA1c6MonthsWindow ON HbA1c6MonthsWindow.SK_PatientID = P.SK_PatientID
	LEFT JOIN  #HbA1cLatest AS HbA1cLatest ON HbA1cLatest.SK_PatientID = P.SK_PatientID
	--LEFT JOIN  #HbA1cHighestEver AS HbA1cHighestEver ON HbA1cHighestEver.SK_PatientID = P.SK_PatientID

	LEFT JOIN  #CholesterolEarliest AS CholesterolEarliest ON CholesterolEarliest.SK_PatientID = P.SK_PatientID
	LEFT JOIN  #Cholesterol6MonthsWindow AS Cholesterol6MonthsWindow ON Cholesterol6MonthsWindow.SK_PatientID = P.SK_PatientID
	LEFT JOIN  #CholesterolLatest AS CholesterolLatest ON CholesterolLatest.SK_PatientID = P.SK_PatientID
	--LEFT JOIN  #CholesterolHighestEver AS CholesterolHighestEver ON CholesterolHighestEver.SK_PatientID = P.SK_PatientID

	LEFT JOIN  #HDL_CholesterolEarliest AS HDL_CholesterolEarliest ON HDL_CholesterolEarliest.SK_PatientID = P.SK_PatientID
	LEFT JOIN  #HDL_Cholesterol6MonthsWindow AS HDL_Cholesterol6MonthsWindow ON HDL_Cholesterol6MonthsWindow.SK_PatientID = P.SK_PatientID
	LEFT JOIN  #HDL_CholesterolLatest AS HDL_CholesterolLatest ON HDL_CholesterolLatest.SK_PatientID = P.SK_PatientID
	--LEFT JOIN  #HDL_CholesterolHighestEver AS HDL_CholesterolHighestEver ON HDL_CholesterolHighestEver.SK_PatientID = P.SK_PatientID

	LEFT JOIN  #SerumLDL_CholesterolEarliest AS SerumLDL_CholesterolEarliest ON SerumLDL_CholesterolEarliest.SK_PatientID = P.SK_PatientID
	LEFT JOIN  #SerumLDL_Cholesterol6MonthsWindow AS SerumLDL_Cholesterol6MonthsWindow ON SerumLDL_Cholesterol6MonthsWindow.SK_PatientID = P.SK_PatientID
	LEFT JOIN  #SerumLDL_CholesterolLatest AS SerumLDL_CholesterolLatest ON SerumLDL_CholesterolLatest.SK_PatientID = P.SK_PatientID
	--LEFT JOIN  #SerumLDL_CholesterolHighestEver AS SerumLDL_CholesterolHighestEver ON SerumLDL_CholesterolHighestEver.SK_PatientID = P.SK_PatientID

	LEFT JOIN  #SerumTriglyceridesEarliest AS SerumTriglyceridesEarliest ON SerumTriglyceridesEarliest.SK_PatientID = P.SK_PatientID
	LEFT JOIN  #SerumTriglycerides6MonthsWindow AS SerumTriglycerides6MonthsWindow ON SerumTriglycerides6MonthsWindow.SK_PatientID = P.SK_PatientID
	LEFT JOIN  #SerumTriglyceridesLatest AS SerumTriglyceridesLatest ON SerumTriglyceridesLatest.SK_PatientID = P.SK_PatientID
	--LEFT JOIN  #SerumTriglyceridesHighestEver AS SerumTriglyceridesHighestEver ON SerumTriglyceridesHighestEver.SK_PatientID = P.SK_PatientID

	LEFT JOIN  #UrineProteinEarliest AS UrineProteinEarliest ON UrineProteinEarliest.SK_PatientID = P.SK_PatientID
	LEFT JOIN  #UrineProteinLatest AS UrineProteinLatest ON UrineProteinLatest.SK_PatientID = P.SK_PatientID

	LEFT JOIN  #CreatininerRatioEarliest AS CreatininerRatioEarliest ON CreatininerRatioEarliest.SK_PatientID = P.SK_PatientID
	LEFT JOIN  #CreatininerRatioLatest AS CreatininerRatioLatest ON CreatininerRatioLatest.SK_PatientID = P.SK_PatientID

	LEFT JOIN  #UrineAlbuminEarliest AS UrineAlbuminEarliest ON UrineAlbuminEarliest.SK_PatientID = P.SK_PatientID
	LEFT JOIN  #UrineAlbuminLatest AS UrineAlbuminLatest ON UrineAlbuminLatest.SK_PatientID = P.SK_PatientID

	LEFT JOIN  #SysBPEarliest AS SysBPEarliest ON SysBPEarliest.SK_PatientID = P.SK_PatientID
	LEFT JOIN  #SysBPLatest AS SysBPLatest ON SysBPLatest.SK_PatientID = P.SK_PatientID
	--LEFT JOIN  #SysBPHighestEver AS SysBPHighestEver ON SysBPHighestEver.SK_PatientID = P.SK_PatientID

	LEFT JOIN  #DiaBPEarliest AS DiaBPEarliest ON DiaBPEarliest.SK_PatientID = P.SK_PatientID
	LEFT JOIN  #DiaBPLatest AS DiaBPLatest ON DiaBPLatest.SK_PatientID = P.SK_PatientID
	--LEFT JOIN  #DiaBPHighestEver AS DiaBPHighestEver ON DiaBPHighestEver.SK_PatientID = P.SK_PatientID

	LEFT JOIN  #CreatinineEarliest AS CreatinineEarliest ON CreatinineEarliest.SK_PatientID = P.SK_PatientID
	LEFT JOIN  #CreatinineLatest AS CreatinineLatest ON CreatinineLatest.SK_PatientID = P.SK_PatientID

	LEFT JOIN  #eGFR_Earliest AS eGFR_Earliest ON eGFR_Earliest.SK_PatientID = P.SK_PatientID
	LEFT JOIN  #eGFR_Latest AS eGFR_Latest ON eGFR_Latest.SK_PatientID = P.SK_PatientID

	LEFT JOIN  #GAD AS GAD ON GAD.SK_PatientID = P.SK_PatientID
	LEFT JOIN  #IsletCell AS IsletCell ON IsletCell.SK_PatientID = P.SK_PatientID
	LEFT JOIN  #IA2 AS IA2 ON IA2.SK_PatientID = P.SK_PatientID
	LEFT JOIN  #CoeliacAutoantibody AS CoeliacAutoantibody ON CoeliacAutoantibody.SK_PatientID = P.SK_PatientID
	LEFT JOIN  #Peroxidase AS Peroxidase ON Peroxidase.SK_PatientID = P.SK_PatientID
	LEFT JOIN  #Cpeptide AS Cpeptide ON Cpeptide.SK_PatientID = P.SK_PatientID
	LEFT JOIN  #SerumInsulin AS SerumInsulin ON SerumInsulin.SK_PatientID = P.SK_PatientID

	LEFT JOIN  #SerumALT AS SerumALT ON SerumALT.SK_PatientID = P.SK_PatientID
	LEFT JOIN  #AST_Serum AS AST_Serum ON AST_Serum.SK_PatientID = P.SK_PatientID

	LEFT JOIN  #TSH_T4 AS TSH_T4 ON TSH_T4.SK_PatientID = P.SK_PatientID

WHERE
	P.CCG = '08V'

GO

/*
--drop all used temp tables
BEGIN -- drop all used temp tables

	DECLARE @DropGlobal bit=0 --Default dont drop global temp table

	DECLARE @DROP_STATEMENT nvarchar(1000)
	DECLARE cursorDEL CURSOR FOR
	SELECT 'DROP TABLE '
	   + case
			   when name like '##%' then name
			   when name like '#%' then SUBSTRING(name, 1, CHARINDEX( '____', name)-1)
		end as DropSQL
	from tempdb..sysobjects
	WHERE name LIKE '#%'
	   AND OBJECT_ID('tempdb..' + name) IS NOT NULL
	   AND name not like case
						   when @DropGlobal=0 then '##%' --//Exclude global temp
						   else '#######%'    --//some fack expression so we can
											   --//select global temp for delete
						end
	OPEN cursorDEL
	FETCH NEXT FROM cursorDEL INTO @DROP_STATEMENT
	WHILE @@FETCH_STATUS = 0
	BEGIN
	EXEC (@DROP_STATEMENT)
	--print @DROP_STATEMENT
	FETCH NEXT FROM cursorDEL INTO @DROP_STATEMENT
	END
	CLOSE cursorDEL
	DEALLOCATE cursorDEL

END

GO
*/
```

5. Prescribing

```
USE CEG

GO

--REPORT 3 PRESCRIBING ----------------------------------------------------------------------------------------------------------------------------

-- drop all temp tables if exist
BEGIN -- drop all temp tables if exist

	DECLARE @DropGlobal bit=0 --Default dont drop global temp table

	DECLARE @DROP_STATEMENT nvarchar(1000)
	DECLARE cursorDEL CURSOR FOR
	SELECT 'DROP TABLE '
	   + case
			   when name like '##%' then name
			   when name like '#%' then SUBSTRING(name, 1, CHARINDEX( '____', name)-1)
		end as DropSQL
	from tempdb..sysobjects
	WHERE name LIKE '#%'
	   AND OBJECT_ID('tempdb..' + name) IS NOT NULL
	   AND name not like case
						   when @DropGlobal=0 then '##%' --//Exclude global temp
						   else '#######%'    --//some fack expression so we can
											   --//select global temp for delete
						end
	OPEN cursorDEL
	FETCH NEXT FROM cursorDEL INTO @DROP_STATEMENT
	WHILE @@FETCH_STATUS = 0
	BEGIN
	EXEC (@DROP_STATEMENT)
	--print @DROP_STATEMENT
	FETCH NEXT FROM cursorDEL INTO @DROP_STATEMENT
	END
	CLOSE cursorDEL
	DEALLOCATE cursorDEL

END

GO

--------Diabetes: ---------------------------------------------------------------------------------------------------------------------------

-------------- Diabetes treatment insulins short acting  ------------------------------------------------------------------------

---------------- Prescribed within previous 12m  ------------------------------------------------------------------------

DECLARE @LastRefreshDate AS DATE
SET @LastRefreshDate = '2018-10-01'

SELECT
	 SK_PatientID
	,[MedicationTerm]
	,IssueDate


INTO  #InsulinsShortPrev12m

FROM
(
	SELECT

		 SK_PatientID
		,[MedicationTerm]
		,IssueDate
		,row_number() over(partition by [SK_PatientID] order by IssueDate DESC) as rn

	FROM
		[08V].[GPMedication]

	WHERE
		(
			BNFChapter = '6.1.1.1'
			OR
			MedicationTerm IN
			(
				'Human Actrapid  Injection  100 units/ml',
				'Human Actrapid  Preloaded Pen  100 units/ml',
				'Human Actrapid Penfill  Cartridges (3 Ml)  100 units/ml',
				'Human Insulatard Ge  Injection  100 units/ml',
				'Human Insulatard Ge  Penfill Cartridges (3 Ml)  100 units/ml',
				'Human Insulatard Ge  Preloaded Pen  100 units/ml',
				'Human Mixtard 10  Penfill Cartridges (3 Ml)  ',
				'Human Mixtard 20  Preloaded Pen  ',
				'Human Mixtard 30  Injection  10 ml vial',
				'Human Mixtard 30  Injection (Cartridges)  100 u/ml',
				'Human Mixtard 30  Penfill Cartridges (3 Ml)  ',
				'Human Mixtard 30  Preloaded Pen  ',
				'Human Mixtard 30 Ge  Injection  ',
				'Human Mixtard 40  Preloaded Pen  ',
				'Human Mixtard 50  Penfill Cartridges (3 Ml)  ',
				'Human Mixtard 50  Preloaded Pen  ',
				'Human Ultratard  Injection  100 units/ml',
				'Humapen  Re-Usable Pen  ',
				'Humapen Luxura  Re-Usable Pen  3 ml, 1-60 units',
				'Humapen Luxura Re-usable pen 3 ml, 1-60 units',
				'Humegon  Injection 75 I.U.  ',
				'Humulin M1  Cartridges (1.5 Ml)  ',
				'Humulin M1  Cartridges (3 Ml)  ',
				'Humulin M2  Cartridges (1.5 Ml)  ',
				'Humulin M2  Injection  ',
				'Humulin M4  Cartridges (3 Ml)  ',
				'Insulin Aspart  Prefilled Syringes  100 units/ml',
				'Isophane Insulin (Human Pyr)  Penfill Cartridges (1.5 Ml)  100 units/ml',
				'Isophane Insulin (Human Pyr)  Penfill Cartridges (3 Ml)  100 units/ml',
				'Isophane Insulin (Human Pyr)  Preloaded Pen  100 units/ml',
				'Lantus 100units/ml solution for injection 3ml pre-filled ...',
				'Mixtard 50  Injection  100 units/ml, 10 ml vial'
			)
		)
		AND
		IssueDate BETWEEN DATEADD(month,-12, @LastRefreshDate) AND 	@LastRefreshDate
		AND
		SK_PatientID IN(SELECT SK_PatientID FROM ceg.GenomicsData WHERE CCG = '08V')

)InsulinsShortPrev12m


WHERE rn = 1

---------------- Prescribed earliest issue date ever --------------------------------------------------------------------

SELECT
	 SK_PatientID
	,[MedicationTerm]
	,IssueDate


INTO  #InsulinsShortEarliest

FROM
(
	SELECT

		 SK_PatientID
		,[MedicationTerm]
		,IssueDate
		,row_number() over(partition by [SK_PatientID] order by IssueDate ASC) as rn

	FROM
		[08V].[GPMedication]

	WHERE
		(
			BNFChapter = '6.1.1.1'
			OR
			MedicationTerm IN
			(
				'Human Actrapid  Injection  100 units/ml',
				'Human Actrapid  Preloaded Pen  100 units/ml',
				'Human Actrapid Penfill  Cartridges (3 Ml)  100 units/ml',
				'Human Insulatard Ge  Injection  100 units/ml',
				'Human Insulatard Ge  Penfill Cartridges (3 Ml)  100 units/ml',
				'Human Insulatard Ge  Preloaded Pen  100 units/ml',
				'Human Mixtard 10  Penfill Cartridges (3 Ml)  ',
				'Human Mixtard 20  Preloaded Pen  ',
				'Human Mixtard 30  Injection  10 ml vial',
				'Human Mixtard 30  Injection (Cartridges)  100 u/ml',
				'Human Mixtard 30  Penfill Cartridges (3 Ml)  ',
				'Human Mixtard 30  Preloaded Pen  ',
				'Human Mixtard 30 Ge  Injection  ',
				'Human Mixtard 40  Preloaded Pen  ',
				'Human Mixtard 50  Penfill Cartridges (3 Ml)  ',
				'Human Mixtard 50  Preloaded Pen  ',
				'Human Ultratard  Injection  100 units/ml',
				'Humapen  Re-Usable Pen  ',
				'Humapen Luxura  Re-Usable Pen  3 ml, 1-60 units',
				'Humapen Luxura Re-usable pen 3 ml, 1-60 units',
				'Humegon  Injection 75 I.U.  ',
				'Humulin M1  Cartridges (1.5 Ml)  ',
				'Humulin M1  Cartridges (3 Ml)  ',
				'Humulin M2  Cartridges (1.5 Ml)  ',
				'Humulin M2  Injection  ',
				'Humulin M4  Cartridges (3 Ml)  ',
				'Insulin Aspart  Prefilled Syringes  100 units/ml',
				'Isophane Insulin (Human Pyr)  Penfill Cartridges (1.5 Ml)  100 units/ml',
				'Isophane Insulin (Human Pyr)  Penfill Cartridges (3 Ml)  100 units/ml',
				'Isophane Insulin (Human Pyr)  Preloaded Pen  100 units/ml',
				'Lantus 100units/ml solution for injection 3ml pre-filled ...',
				'Mixtard 50  Injection  100 units/ml, 10 ml vial'
			)
		)
		AND
		SK_PatientID IN(SELECT SK_PatientID FROM ceg.GenomicsData WHERE CCG = '08V')

) AS InsulinsShortEarliest


WHERE rn = 1


-------------- Diabetes treatment insulins Intermediate and long acting  ------------------------------------------------------------------------

---------------- Prescribed within previous 12m  ------------------------------------------------------------------------


SELECT
	 SK_PatientID
	,[MedicationTerm]
	,IssueDate


INTO  #InsulinsLongPrev12m

FROM
(
	SELECT

		 SK_PatientID
		,[MedicationTerm]
		,IssueDate
		,row_number() over(partition by [SK_PatientID] order by IssueDate DESC) as rn

	FROM
		[08V].[GPMedication]

	WHERE
		(
			BNFChapter = '6.1.1.2'
		)
		AND
		IssueDate BETWEEN DATEADD(month,-12, @LastRefreshDate) AND 	@LastRefreshDate
		AND
		SK_PatientID IN(SELECT SK_PatientID FROM ceg.GenomicsData WHERE CCG = '08V')

)InsulinsLongPrev12m


WHERE rn = 1


---------------- Prescribed earliest issue date ever --------------------------------------------------------------------

SELECT
	 SK_PatientID
	,[MedicationTerm]
	,IssueDate


INTO  #InsulinsLongEarliest

FROM
(
	SELECT

		 SK_PatientID
		,[MedicationTerm]
		,IssueDate
		,row_number() over(partition by [SK_PatientID] order by IssueDate ASC) as rn

	FROM
		[08V].[GPMedication]

	WHERE
		(
			BNFChapter = '6.1.1.2'
		)
		AND
		SK_PatientID IN(SELECT SK_PatientID FROM ceg.GenomicsData WHERE CCG = '08V')

) AS InsulinsLongEarliest


WHERE rn = 1


-------------- Antidiabetic drugs sulphonylureas  --------------------------------------------------------------------------------

---------------- Prescribed within previous 12m  ------------------------------------------------------------------------

SELECT
	 SK_PatientID
	,[MedicationTerm]
	,IssueDate


INTO  #SulphonylureasPrev12m

FROM
(
	SELECT

		 SK_PatientID
		,[MedicationTerm]
		,IssueDate
		,row_number() over(partition by [SK_PatientID] order by IssueDate DESC) as rn

	FROM
		[08V].[GPMedication]

	WHERE
		(
			BNFChapter = '6.1.2.1'
		)
		AND
		IssueDate BETWEEN DATEADD(month,-12, @LastRefreshDate) AND 	@LastRefreshDate
		AND
		SK_PatientID IN(SELECT SK_PatientID FROM ceg.GenomicsData WHERE CCG = '08V')

)SulphonylureasPrev12m


WHERE rn = 1

---------------- Prescribed earliest issue date ever --------------------------------------------------------------------

SELECT
	 SK_PatientID
	,[MedicationTerm]
	,IssueDate


INTO  #SulphonylureasEarliest

FROM
(
	SELECT

		 SK_PatientID
		,[MedicationTerm]
		,IssueDate
		,row_number() over(partition by [SK_PatientID] order by IssueDate ASC) as rn

	FROM
		[08V].[GPMedication]

	WHERE
		(
			BNFChapter = '6.1.2.1'
		)
		AND
		SK_PatientID IN(SELECT SK_PatientID FROM ceg.GenomicsData WHERE CCG = '08V')

) AS SulphonylureasEarliest


WHERE rn = 1

-------------- Antidiabetic drugs biganuides  ------------------------------------------------------------------------------

---------------- Prescribed within previous 12m  ------------------------------------------------------------------------

SELECT
	 SK_PatientID
	,[MedicationTerm]
	,IssueDate


INTO  #MetforminPrev12m

FROM
(
	SELECT

		 SK_PatientID
		,[MedicationTerm]
		,IssueDate
		,row_number() over(partition by [SK_PatientID] order by IssueDate DESC) as rn

	FROM
		[08V].[GPMedication]

	WHERE
		(
			BNFChapter = '6.1.2.2'
			OR
			MedicationTerm LIKE '%Glucamet%'	-- 'Glucamet  Tablets  850 mg'
		)
		AND
		IssueDate BETWEEN DATEADD(month,-12, @LastRefreshDate) AND 	@LastRefreshDate
		AND
		SK_PatientID IN(SELECT SK_PatientID FROM ceg.GenomicsData WHERE CCG = '08V')

)MetforminPrev12m


WHERE rn = 1

---------------- Prescribed earliest issue date ever --------------------------------------------------------------------

SELECT
	 SK_PatientID
	,[MedicationTerm]
	,IssueDate


INTO  #MetforminEarliest

FROM
(
	SELECT

		 SK_PatientID
		,[MedicationTerm]
		,IssueDate
		,row_number() over(partition by [SK_PatientID] order by IssueDate ASC) as rn

	FROM
		[08V].[GPMedication]

	WHERE
		(
			BNFChapter = '6.1.2.2'
			OR
			MedicationTerm LIKE '%Glucamet%'	-- 'Glucamet  Tablets  850 mg'
		)
		AND
		SK_PatientID IN(SELECT SK_PatientID FROM ceg.GenomicsData WHERE CCG = '08V')

) AS MetforminEarliest


WHERE rn = 1


-------------- Antidiabetic drugs Other  -----------------------------------------------------------------------------------

----------------Alpha-glucosidase inhibitors - prescribed within previous 12m  ---------------------------------------------

SELECT
	 SK_PatientID
	,[MedicationTerm]
	,IssueDate


INTO  #AGIsPrev12m

FROM
(
	SELECT

		 SK_PatientID
		,[MedicationTerm]
		,IssueDate
		,row_number() over(partition by [SK_PatientID] order by IssueDate DESC) as rn

	FROM
		[08V].[GPMedication]

	WHERE
		(
			BNFChapter = '6.1.2.3'
			AND
			(
				MedicationTerm LIKE 'Acarbose%'
				OR
				MedicationTerm LIKE 'Glucobay%'
			)
		)
		AND
		IssueDate BETWEEN DATEADD(month,-12, @LastRefreshDate) AND 	@LastRefreshDate
		AND
		SK_PatientID IN(SELECT SK_PatientID FROM ceg.GenomicsData WHERE CCG = '08V')

)AGIsPrev12m


WHERE rn = 1

---------------- Alpha-glucosidase inhibitors - prescribed earliest issue date ever ----------------------------------------------

SELECT
	 SK_PatientID
	,[MedicationTerm]
	,IssueDate


INTO  #AGIsEarliest

FROM
(
	SELECT

		 SK_PatientID
		,[MedicationTerm]
		,IssueDate
		,row_number() over(partition by [SK_PatientID] order by IssueDate ASC) as rn

	FROM
		[08V].[GPMedication]

	WHERE
		(
			BNFChapter = '6.1.2.3'
			AND
			(
				MedicationTerm LIKE 'Acarbose%'
				OR
				MedicationTerm LIKE 'Glucobay%'
			)
		)
		AND
		SK_PatientID IN(SELECT SK_PatientID FROM ceg.GenomicsData WHERE CCG = '08V')

) AS AGIsEarliest

WHERE rn = 1


---------------- Dipeptidylpeptidase-4 inhibitors (gliptins) - prescribed within previous 12m  ---------------------------------------------


SELECT
	 SK_PatientID
	,[MedicationTerm]
	,IssueDate


INTO  #DPP4Prev12m

FROM
(
	SELECT

		 SK_PatientID
		,[MedicationTerm]
		,IssueDate
		,row_number() over(partition by [SK_PatientID] order by IssueDate DESC) as rn

	FROM
		[08V].[GPMedication]

	WHERE
		(
			BNFChapter = '6.1.2.3'
			AND
			(
				MedicationTerm LIKE 'Alogliptin%'
				OR
				MedicationTerm LIKE 'Vipidia%'
				OR
				MedicationTerm LIKE 'Linagliptin%'
				OR
				MedicationTerm LIKE 'Trajenta%'
				OR
				MedicationTerm LIKE 'Saxagliptin%'
				OR
				MedicationTerm LIKE 'Onglyza%'
				OR
				MedicationTerm LIKE 'Sitagliptin%'
				OR
				MedicationTerm LIKE 'Januvia%'
				OR
				MedicationTerm LIKE 'Vildagliptin%'
				OR
				MedicationTerm LIKE 'Galvus%'
			)
		)
		AND
		IssueDate BETWEEN DATEADD(month,-12, @LastRefreshDate) AND 	@LastRefreshDate
		AND
		SK_PatientID IN(SELECT SK_PatientID FROM ceg.GenomicsData WHERE CCG = '08V')

)DPP4Prev12m


WHERE rn = 1

---------------- Dipeptidylpeptidase-4 inhibitors (gliptins) - prescribed earliest issue date ever ----------------------------------------------

SELECT
	 SK_PatientID
	,[MedicationTerm]
	,IssueDate


INTO  #DPP4Earliest

FROM
(
	SELECT

		 SK_PatientID
		,[MedicationTerm]
		,IssueDate
		,row_number() over(partition by [SK_PatientID] order by IssueDate ASC) as rn

	FROM
		[08V].[GPMedication]

	WHERE
		(
			BNFChapter = '6.1.2.3'
			AND
			(
				MedicationTerm LIKE 'Alogliptin%'
				OR
				MedicationTerm LIKE 'Vipidia%'
				OR
				MedicationTerm LIKE 'Linagliptin%'
				OR
				MedicationTerm LIKE 'Trajenta%'
				OR
				MedicationTerm LIKE 'Saxagliptin%'
				OR
				MedicationTerm LIKE 'Onglyza%'
				OR
				MedicationTerm LIKE 'Sitagliptin%'
				OR
				MedicationTerm LIKE 'Januvia%'
				OR
				MedicationTerm LIKE 'Vildagliptin%'
				OR
				MedicationTerm LIKE 'Galvus%'
			)
		)
		AND
		SK_PatientID IN(SELECT SK_PatientID FROM ceg.GenomicsData WHERE CCG = '08V')

) AS DPP4Earliest

WHERE rn = 1


---------------- Glucagon-like peptide-1 receptor agonists - prescribed within previous 12m  ---------------------------------------------


SELECT
	 SK_PatientID
	,[MedicationTerm]
	,IssueDate


INTO  #GLP1Prev12m

FROM
(
	SELECT

		 SK_PatientID
		,[MedicationTerm]
		,IssueDate
		,row_number() over(partition by [SK_PatientID] order by IssueDate DESC) as rn

	FROM
		[08V].[GPMedication]

	WHERE
		(
			BNFChapter = '6.1.2.3'
			AND
			(
				-- Albiglutide
				MedicationTerm LIKE 'Albiglutide%'
				OR
				MedicationTerm LIKE 'Eperzan%'
				OR
				-- Dulaglutide
				MedicationTerm LIKE 'Dulaglutide%'
				OR
				MedicationTerm LIKE 'Trulicity%'
				OR

				-- Exenatide
				MedicationTerm LIKE 'Byetta%'
				OR
				MedicationTerm LIKE 'Bydureon%'
				OR
				MedicationTerm LIKE 'Exenatide%'
				OR

				-- Liraglutide,
				MedicationTerm LIKE 'Liraglutide%'
				OR
				MedicationTerm LIKE 'Victoza%'
				OR

				-- Lixisenatide
				MedicationTerm LIKE 'Lixisenatide%'
				OR
				MedicationTerm LIKE 'Lyxumia%'
			)
		)
		AND
		IssueDate BETWEEN DATEADD(month,-12, @LastRefreshDate) AND 	@LastRefreshDate
		AND
		SK_PatientID IN(SELECT SK_PatientID FROM ceg.GenomicsData WHERE CCG = '08V')

)GLP1Prev12m


WHERE rn = 1


---------------- Glucagon-like peptide-1 receptor agonists - prescribed earliest issue date ever ----------------------------------------------

SELECT
	 SK_PatientID
	,[MedicationTerm]
	,IssueDate


INTO  #GLP1Earliest

FROM
(
	SELECT

		 SK_PatientID
		,[MedicationTerm]
		,IssueDate
		,row_number() over(partition by [SK_PatientID] order by IssueDate ASC) as rn

	FROM
		[08V].[GPMedication]

	WHERE
		(
			BNFChapter = '6.1.2.3'
			AND
			(
				-- Albiglutide
				MedicationTerm LIKE 'Albiglutide%'
				OR
				MedicationTerm LIKE 'Eperzan%'
				OR
				-- Dulaglutide
				MedicationTerm LIKE 'Dulaglutide%'
				OR
				MedicationTerm LIKE 'Trulicity%'
				OR

				-- Exenatide
				MedicationTerm LIKE 'Byetta%'
				OR
				MedicationTerm LIKE 'Bydureon%'
				OR
				MedicationTerm LIKE 'Exenatide%'
				OR

				-- Liraglutide,
				MedicationTerm LIKE 'Liraglutide%'
				OR
				MedicationTerm LIKE 'Victoza%'
				OR

				-- Lixisenatide
				MedicationTerm LIKE 'Lixisenatide%'
				OR
				MedicationTerm LIKE 'Lyxumia%'
			)
		)
		AND
		SK_PatientID IN(SELECT SK_PatientID FROM ceg.GenomicsData WHERE CCG = '08V')

) AS GLP1Earliest

WHERE rn = 1


---------------- Meglitinides - prescribed within previous 12m  ---------------------------------------------

SELECT
	 SK_PatientID
	,[MedicationTerm]
	,IssueDate


INTO  #MeglitinidesPrev12m

FROM
(
	SELECT

		 SK_PatientID
		,[MedicationTerm]
		,IssueDate
		,row_number() over(partition by [SK_PatientID] order by IssueDate DESC) as rn

	FROM
		[08V].[GPMedication]

	WHERE
		(
			BNFChapter = '6.1.2.3'
			AND
			(
				-- Nateglinide
				MedicationTerm LIKE 'Nateglinide%'
				OR
				MedicationTerm LIKE 'Starlix%'
				OR

				-- Repaglinide
				MedicationTerm LIKE 'Prandin%'
				OR
				MedicationTerm LIKE 'Repaglinide%'
			)
		)
		AND
		IssueDate BETWEEN DATEADD(month,-12, @LastRefreshDate) AND 	@LastRefreshDate
		AND
		SK_PatientID IN(SELECT SK_PatientID FROM ceg.GenomicsData WHERE CCG = '08V')

)MeglitinidesPrev12m


WHERE rn = 1


---------------- Meglitinides - prescribed earliest issue date ever ----------------------------------------------

SELECT
	 SK_PatientID
	,[MedicationTerm]
	,IssueDate


INTO  #MeglitinidesEarliest

FROM
(
	SELECT

		 SK_PatientID
		,[MedicationTerm]
		,IssueDate
		,row_number() over(partition by [SK_PatientID] order by IssueDate ASC) as rn

	FROM
		[08V].[GPMedication]

	WHERE
		(
			BNFChapter = '6.1.2.3'
			AND
			(
				-- Nateglinide
				MedicationTerm LIKE 'Nateglinide%'
				OR
				MedicationTerm LIKE 'Starlix%'
				OR

				-- Repaglinide
				MedicationTerm LIKE 'Prandin%'
				OR
				MedicationTerm LIKE 'Repaglinide%'
			)
		)
		AND
		SK_PatientID IN(SELECT SK_PatientID FROM ceg.GenomicsData WHERE CCG = '08V')

) AS MeglitinidesEarliest

WHERE rn = 1

---------------- Sodium glucose co-transporter 2 inhibitors - prescribed within previous 12m  ---------------------------------------------

SELECT
	 SK_PatientID
	,[MedicationTerm]
	,IssueDate


INTO  #SGLT2Prev12m

FROM
(
	SELECT

		 SK_PatientID
		,[MedicationTerm]
		,IssueDate
		,row_number() over(partition by [SK_PatientID] order by IssueDate DESC) as rn

	FROM
		[08V].[GPMedication]

	WHERE
		(
			BNFChapter = '6.1.2.3'
			AND
			(
				-- Canagliflozin
				MedicationTerm LIKE 'Canagliflozin%'
				OR
				MedicationTerm LIKE 'Invokana%'
				OR

				-- Dapagliflozin
				MedicationTerm LIKE 'Dapagliflozin%'
				OR
				MedicationTerm LIKE 'Forxiga%'
				OR
				-- Empagliflozin
				MedicationTerm LIKE 'Empagliflozin%'
				OR
				MedicationTerm LIKE 'Jardiance%'
			)
		)
		AND
		IssueDate BETWEEN DATEADD(month,-12, @LastRefreshDate) AND 	@LastRefreshDate
		AND
		SK_PatientID IN(SELECT SK_PatientID FROM ceg.GenomicsData WHERE CCG = '08V')

)SGLT2Prev12m


WHERE rn = 1

---------------- Sodium glucose co-transporter 2 inhibitors - prescribed earliest issue date ever ----------------------------------------------

SELECT
	 SK_PatientID
	,[MedicationTerm]
	,IssueDate


INTO  #SGLT2Earliest

FROM
(
	SELECT

		 SK_PatientID
		,[MedicationTerm]
		,IssueDate
		,row_number() over(partition by [SK_PatientID] order by IssueDate ASC) as rn

	FROM
		[08V].[GPMedication]

	WHERE
		(
			BNFChapter = '6.1.2.3'
			AND
			(
				-- Canagliflozin
				MedicationTerm LIKE 'Canagliflozin%'
				OR
				MedicationTerm LIKE 'Invokana%'
				OR

				-- Dapagliflozin
				MedicationTerm LIKE 'Dapagliflozin%'
				OR
				MedicationTerm LIKE 'Forxiga%'
				OR
				-- Empagliflozin
				MedicationTerm LIKE 'Empagliflozin%'
				OR
				MedicationTerm LIKE 'Jardiance%'
			)
		)
		AND
		SK_PatientID IN(SELECT SK_PatientID FROM ceg.GenomicsData WHERE CCG = '08V')

) AS SGLT2Earliest

WHERE rn = 1

---------------- Thiazolidinediones - prescribed within previous 12m  ---------------------------------------------

SELECT
	 SK_PatientID
	,[MedicationTerm]
	,IssueDate


INTO  #ThiazolidinedionesPrev12m

FROM
(
	SELECT

		 SK_PatientID
		,[MedicationTerm]
		,IssueDate
		,row_number() over(partition by [SK_PatientID] order by IssueDate DESC) as rn

	FROM
		[08V].[GPMedication]

	WHERE
		(
			BNFChapter = '6.1.2.3'
			AND
			(
				-- Pioglitazone
				MedicationTerm LIKE 'Actos%'
				OR
				MedicationTerm LIKE 'Pioglitazone%'
			)
		)
		AND
		IssueDate BETWEEN DATEADD(month,-12, @LastRefreshDate) AND 	@LastRefreshDate
		AND
		SK_PatientID IN(SELECT SK_PatientID FROM ceg.GenomicsData WHERE CCG = '08V')

)ThiazolidinedionesPrev12m


WHERE rn = 1

---------------- Thiazolidinediones - prescribed earliest issue date ever ----------------------------------------------

SELECT
	 SK_PatientID
	,[MedicationTerm]
	,IssueDate


INTO  #ThiazolidinedionesEarliest

FROM
(
	SELECT

		 SK_PatientID
		,[MedicationTerm]
		,IssueDate
		,row_number() over(partition by [SK_PatientID] order by IssueDate ASC) as rn

	FROM
		[08V].[GPMedication]

	WHERE
		(
			BNFChapter = '6.1.2.3'
			AND
			(
				-- Pioglitazone
				MedicationTerm LIKE 'Actos%'
				OR
				MedicationTerm LIKE 'Pioglitazone%'
			)
		)
		AND
		SK_PatientID IN(SELECT SK_PatientID FROM ceg.GenomicsData WHERE CCG = '08V')

) AS ThiazolidinedionesEarliest

WHERE rn = 1

---------------- Combination drugs - prescribed within previous 12m  ---------------------------------------------

SELECT
	 SK_PatientID
	,[MedicationTerm]
	,IssueDate


INTO  #CombinationDrugsPrev12m

FROM
(
	SELECT

		 SK_PatientID
		,[MedicationTerm]
		,IssueDate
		,row_number() over(partition by [SK_PatientID] order by IssueDate DESC) as rn

	FROM
		[08V].[GPMedication]

	WHERE
		(
			BNFChapter = '6.1.2.3'
			AND
			(
				-- Alogliptin with metformin
					MedicationTerm LIKE 'Alogliptin%'
				OR	MedicationTerm LIKE 'Vipdomet%'

				-- Linagliptin with metformin
				OR	MedicationTerm LIKE 'Linagliptin%'
				OR	MedicationTerm LIKE 'Jentadueto%'

				-- Saxagliptin with metformin
				OR	MedicationTerm LIKE 'Saxagliptin%'
				OR	MedicationTerm LIKE 'Komboglyze%'

				-- Sitagliptin with metformin
				OR	MedicationTerm LIKE 'Metformin%'
				OR	MedicationTerm LIKE 'Janumet%'

				-- Vildaglitpin with metformin
				OR	MedicationTerm LIKE 'Vildagliptin%'
				OR	MedicationTerm LIKE 'Eucreas%'

				-- Canagliflozin with metformin
				OR	MedicationTerm LIKE 'Canagliflozin%'
				OR	MedicationTerm LIKE 'Vokanamet%'

				-- Dapagliflozin with metformin
				OR	MedicationTerm LIKE 'Dapagliflozin%'
				OR	MedicationTerm LIKE 'Xigduo%'

				-- Empagliflozin with metformin
				OR	MedicationTerm LIKE 'Empagliflozin%'
				OR	MedicationTerm LIKE 'Synjardy%'

				-- Pioglitazone with metformin
				OR	MedicationTerm LIKE 'Pioglitazone%'
				OR	MedicationTerm LIKE 'Competact%'

				-- INSULIN DEGLUDEC WITH LIRAGLUTIDE
				OR	MedicationTerm LIKE 'Insulin degludec%'
				OR	MedicationTerm LIKE 'Xultophy%'
			)
		)
		AND
		IssueDate BETWEEN DATEADD(month,-12, @LastRefreshDate) AND 	@LastRefreshDate
		AND
		SK_PatientID IN(SELECT SK_PatientID FROM ceg.GenomicsData WHERE CCG = '08V')

)CombinationDrugsPrev12m


WHERE rn = 1

---------------- Combination drugs - prescribed earliest issue date ever ----------------------------------------------

SELECT
	 SK_PatientID
	,[MedicationTerm]
	,IssueDate


INTO  #CombinationDrugsEarliest

FROM
(
	SELECT

		 SK_PatientID
		,[MedicationTerm]
		,IssueDate
		,row_number() over(partition by [SK_PatientID] order by IssueDate ASC) as rn

	FROM
		[08V].[GPMedication]

	WHERE
		(
			BNFChapter = '6.1.2.3'
			AND
			(
				-- Alogliptin with metformin
					MedicationTerm LIKE 'Alogliptin%'
				OR	MedicationTerm LIKE 'Vipdomet%'

				-- Linagliptin with metformin
				OR	MedicationTerm LIKE 'Linagliptin%'
				OR	MedicationTerm LIKE 'Jentadueto%'

				-- Saxagliptin with metformin
				OR	MedicationTerm LIKE 'Saxagliptin%'
				OR	MedicationTerm LIKE 'Komboglyze%'

				-- Sitagliptin with metformin
				OR	MedicationTerm LIKE 'Metformin%'
				OR	MedicationTerm LIKE 'Janumet%'

				-- Vildaglitpin with metformin
				OR	MedicationTerm LIKE 'Vildagliptin%'
				OR	MedicationTerm LIKE 'Eucreas%'

				-- Canagliflozin with metformin
				OR	MedicationTerm LIKE 'Canagliflozin%'
				OR	MedicationTerm LIKE 'Vokanamet%'

				-- Dapagliflozin with metformin
				OR	MedicationTerm LIKE 'Dapagliflozin%'
				OR	MedicationTerm LIKE 'Xigduo%'

				-- Empagliflozin with metformin
				OR	MedicationTerm LIKE 'Empagliflozin%'
				OR	MedicationTerm LIKE 'Synjardy%'

				-- Pioglitazone with metformin
				OR	MedicationTerm LIKE 'Pioglitazone%'
				OR	MedicationTerm LIKE 'Competact%'

				-- INSULIN DEGLUDEC WITH LIRAGLUTIDE
				OR	MedicationTerm LIKE 'Insulin degludec%'
				OR	MedicationTerm LIKE 'Xultophy%'
			)
		)
		AND
		SK_PatientID IN(SELECT SK_PatientID FROM ceg.GenomicsData WHERE CCG = '08V')

) AS CombinationDrugsEarliest

WHERE rn = 1


-------------- Blood glucose and ketone testing strips  -----------------------------------------------------------------------------------

---------------- Prescribed within previous 12m  ------------------------------------------------------------------------------------------

SELECT
	 SK_PatientID
	,[MedicationTerm]
	,IssueDate


INTO  #TestingStripsPrev12m

FROM
(
	SELECT

		 SK_PatientID
		,[MedicationTerm]
		,IssueDate
		,row_number() over(partition by [SK_PatientID] order by IssueDate DESC) as rn

	FROM
		[08V].[GPMedication]

	WHERE
		(
			BNFChapter = '6.1.6'
			OR
			MedicationTerm IN
			(
				-- BLOOD GLUCOSE STRIPS EMIS LIST CROSS CHECKED WITH BNF
				'Active testing strips (Roche Diabetes Care Ltd)',
				'Advocate Redi-Code+ testing strips (Diabetes Care Technology Ltd)',
				'AutoSense testing strips (Advance Diagnostic Products (NI) Ltd)',
				'Aviva testing strips (Roche Diabetes Care Ltd)',
				'BGStar testing strips (Sanofi)',
				'Betachek G5 testing strips (National Diagnostic Products)',
				'Betachek C50 cassette (National Diagnostic Products)',
				'Betachek Visual testing strips (National Diagnostic Products)',
				'Breeze 2 testing discs (Bayer Plc)',
				'CareSens N testing strips (Spirit Healthcare Ltd)',
				'Compact testing strips (Roche Diabetes Care Ltd)',
				'Contour Next testing strips (Bayer Diagnostics Manufacturing Ltd)',
				'Contour TS testing strips (Bayer Diagnostics Manufacturing Ltd)',
				'Contour testing strips (Bayer Diagnostics Manufacturing Ltd)',
				'CozyLab S7 testing strips (Health Integrated Technologies Ltd)',
				'Dario testing strips (Farla Medical Ltd)',
				'Dario Lite testing strips (LabStyle Innovations Ltd)',
				'Diastix testing strips (Bayer Diagnostics Manufacturing Ltd)',
				'eBchek testing strips (IRASCO Ltd)',
				'Element testing strips (Neon Diagnostics Ltd)',
				'FreeStyle testing strips (Abbott Laboratories Ltd)',
				'FreeStyle Optium testing strips (Abbott Laboratories Ltd)',
				'FreeStyle Lite testing strips (Abbott Laboratories Ltd)',
				'GlucoDock testing strips (Medisana Healthcare (UK) Ltd)',
				'Glucoflex-R testing strips (Bio-Diagnostics Ltd)',
				'GlucoLab testing strips (Neon Diagnostics Ltd)',
				'GlucoMen Visio testing strips (A Menarini Diagnostics Ltd)',
				'GlucoMen Sensor testing strips (A Menarini Diagnostics Ltd)',
				'GlucoMen LX Sensor testing strips (A Menarini Diagnostics Ltd)',
				'GlucoMen GM testing strips (A Menarini Diagnostics Ltd)',
				'GlucoMen areo Sensor testing strips (A Menarini Diagnostics Ltd)',
				'GlucoRx Nexus testing strips (GlucoRx Ltd)',
				'GlucoRx Original testing strips (GlucoRx Ltd)',
				'GlucoZen.auto testing strips (GlucoZen Ltd)',
				'GluNEO testing strips (Neon Diagnostics Ltd)',
				'Icare Advanced Testing strips',
				'Icare Advanced Solo Testing strips',
				'iHealth testing strips (Technomed Ltd)',
				'IME-DC testing strips (Arctic Medical Ltd)',
				'MediSense SoftSense testing strips (Abbott Laboratories Ltd)',
				'Medi-Test Glucose testing strips (BHR Pharmaceuticals Ltd)',
				'MediTouch testing strips (Medisana Healthcare (UK) Ltd)',
				'Mendor Discreet testing strips (SpringMed Solutions Ltd)',
				'Microdot testing strips (Cambridge Sensors Ltd)',
				'Mission Glucose testing strips 1G (Spirit Healthcare Ltd)',
				'Mobile cassette (Roche Diabetes Care Ltd)',
				'MODZ testing strips (Modz Oy)',
				'Myglucohealth testing strips (Entra Health Systems Ltd)',
				'Mylife Pura testing strips (Ypsomed Ltd)',
				'Mylife Unio testing strips (Ypsomed Ltd)',
				'Omnitest 3 testing strips (B.Braun Medical Ltd)',
				'On-Call Advanced testing strips (Point Of Care Testing Ltd)',
				'OneTouch testing strips (LifeScan)',
				'OneTouch Ultra testing strips (LifeScan)',
				'OneTouch Verio testing strips (LifeScan)',
				'OneTouch Vita testing strips (LifeScan)',
				'Performa testing strips (Roche Diabetes Care Ltd)',
				'SD CodeFree testing strips (SD Biosensor Inc)',
				'Sensocard testing strips (BBI Healthcare Ltd)',
				'SuperCheck 2 testing strips (Apollo Medical Technologies Ltd)',
				'SuperCheck Plus testing strips (Apollo Medical Technologies Ltd)',
				'SURESIGN Resure testing strips (Ciga Healthcare Ltd)',
				'TEE2 testing strips (Spirit Healthcare Ltd)',
				'TRUEone testing strips (Nipro Diagnostics (UK) Ltd)',
				'TRUEresult testing strips (Nipro Diagnostics (UK) Ltd)',
				'TrueTrack System testing strips (Nipro Diagnostics (UK) Ltd)',
				'TRUEyou testing strips (Nipro Diagnostics (UK) Ltd)',
				'WaveSense JAZZ testing strips (AgaMatrix Europe Ltd)',
				'WaveSense JAZZ Duo testing strips (AgaMatrix Europe Ltd)',
				'Accu-Chek Advantage  Meter  ',
				'Accu-Chek Aviva  Blood Glucose Testing System  ',
				'Accu-Chek Aviva Blood Glucose Testing System',
				'Accu-Chek Compact  Meter  ',
				'Accu-Chek Compact Meter',
				'Accu-Chek Compact Plus  Blood Glucose Testing System  ',
				'Accu-Chek Compact Plus Blood Glucose Testing System',
				'Ascensia Microfill biosensor strips [BAYER]',
				'Bm Test 5L  Reagent Strips  ',
				'Bm-Stix  Test Strips  ',
				'Glucorx  Strips  ',
				'GlucoRx Original testing strips (Disposable Medical Equipment Ltd)',
				'GlucoRx Original testing strips (GlucoRx Ltd)',
				'GlucoRx testing strips (Disposable Medical Equipment Ltd)',
				'Glucotrend  Test Strips  ',
				'Glucotrend color.strip(s) [ROCHE DIAG]',
				'Glucotrend Plus  Reagent Strips  ',
				'Hypoguard Ga  Reagent Strips  ',

				-- KETONE STRIPS
				'Ketostix testing strips (Bayer Diagnostics Manufacturing Ltd)',
				'Mission Ketone testing strips 1K (Spirit Healthcare Ltd'
			)
		)
		AND
		IssueDate BETWEEN DATEADD(month,-12, @LastRefreshDate) AND 	@LastRefreshDate
		AND
		SK_PatientID IN(SELECT SK_PatientID FROM ceg.GenomicsData WHERE CCG = '08V')

) AS TestingStripsPrev12m


WHERE rn = 1

---------------- Prescribed earliest issue date ever --------------------------------------------------------------------

SELECT
	 SK_PatientID
	,[MedicationTerm]
	,IssueDate


INTO  #TestingStripsEarliest

FROM
(
	SELECT

		 SK_PatientID
		,[MedicationTerm]
		,IssueDate
		,row_number() over(partition by [SK_PatientID] order by IssueDate ASC) as rn

	FROM
		[08V].[GPMedication]

	WHERE
		(
			BNFChapter = '6.1.6'
			OR
			MedicationTerm IN
			(
				-- BLOOD GLUCOSE STRIPS EMIS LIST CROSS CHECKED WITH BNF
				'Active testing strips (Roche Diabetes Care Ltd)',
				'Advocate Redi-Code+ testing strips (Diabetes Care Technology Ltd)',
				'AutoSense testing strips (Advance Diagnostic Products (NI) Ltd)',
				'Aviva testing strips (Roche Diabetes Care Ltd)',
				'BGStar testing strips (Sanofi)',
				'Betachek G5 testing strips (National Diagnostic Products)',
				'Betachek C50 cassette (National Diagnostic Products)',
				'Betachek Visual testing strips (National Diagnostic Products)',
				'Breeze 2 testing discs (Bayer Plc)',
				'CareSens N testing strips (Spirit Healthcare Ltd)',
				'Compact testing strips (Roche Diabetes Care Ltd)',
				'Contour Next testing strips (Bayer Diagnostics Manufacturing Ltd)',
				'Contour TS testing strips (Bayer Diagnostics Manufacturing Ltd)',
				'Contour testing strips (Bayer Diagnostics Manufacturing Ltd)',
				'CozyLab S7 testing strips (Health Integrated Technologies Ltd)',
				'Dario testing strips (Farla Medical Ltd)',
				'Dario Lite testing strips (LabStyle Innovations Ltd)',
				'Diastix testing strips (Bayer Diagnostics Manufacturing Ltd)',
				'eBchek testing strips (IRASCO Ltd)',
				'Element testing strips (Neon Diagnostics Ltd)',
				'FreeStyle testing strips (Abbott Laboratories Ltd)',
				'FreeStyle Optium testing strips (Abbott Laboratories Ltd)',
				'FreeStyle Lite testing strips (Abbott Laboratories Ltd)',
				'GlucoDock testing strips (Medisana Healthcare (UK) Ltd)',
				'Glucoflex-R testing strips (Bio-Diagnostics Ltd)',
				'GlucoLab testing strips (Neon Diagnostics Ltd)',
				'GlucoMen Visio testing strips (A Menarini Diagnostics Ltd)',
				'GlucoMen Sensor testing strips (A Menarini Diagnostics Ltd)',
				'GlucoMen LX Sensor testing strips (A Menarini Diagnostics Ltd)',
				'GlucoMen GM testing strips (A Menarini Diagnostics Ltd)',
				'GlucoMen areo Sensor testing strips (A Menarini Diagnostics Ltd)',
				'GlucoRx Nexus testing strips (GlucoRx Ltd)',
				'GlucoRx Original testing strips (GlucoRx Ltd)',
				'GlucoZen.auto testing strips (GlucoZen Ltd)',
				'GluNEO testing strips (Neon Diagnostics Ltd)',
				'Icare Advanced Testing strips',
				'Icare Advanced Solo Testing strips',
				'iHealth testing strips (Technomed Ltd)',
				'IME-DC testing strips (Arctic Medical Ltd)',
				'MediSense SoftSense testing strips (Abbott Laboratories Ltd)',
				'Medi-Test Glucose testing strips (BHR Pharmaceuticals Ltd)',
				'MediTouch testing strips (Medisana Healthcare (UK) Ltd)',
				'Mendor Discreet testing strips (SpringMed Solutions Ltd)',
				'Microdot testing strips (Cambridge Sensors Ltd)',
				'Mission Glucose testing strips 1G (Spirit Healthcare Ltd)',
				'Mobile cassette (Roche Diabetes Care Ltd)',
				'MODZ testing strips (Modz Oy)',
				'Myglucohealth testing strips (Entra Health Systems Ltd)',
				'Mylife Pura testing strips (Ypsomed Ltd)',
				'Mylife Unio testing strips (Ypsomed Ltd)',
				'Omnitest 3 testing strips (B.Braun Medical Ltd)',
				'On-Call Advanced testing strips (Point Of Care Testing Ltd)',
				'OneTouch testing strips (LifeScan)',
				'OneTouch Ultra testing strips (LifeScan)',
				'OneTouch Verio testing strips (LifeScan)',
				'OneTouch Vita testing strips (LifeScan)',
				'Performa testing strips (Roche Diabetes Care Ltd)',
				'SD CodeFree testing strips (SD Biosensor Inc)',
				'Sensocard testing strips (BBI Healthcare Ltd)',
				'SuperCheck 2 testing strips (Apollo Medical Technologies Ltd)',
				'SuperCheck Plus testing strips (Apollo Medical Technologies Ltd)',
				'SURESIGN Resure testing strips (Ciga Healthcare Ltd)',
				'TEE2 testing strips (Spirit Healthcare Ltd)',
				'TRUEone testing strips (Nipro Diagnostics (UK) Ltd)',
				'TRUEresult testing strips (Nipro Diagnostics (UK) Ltd)',
				'TrueTrack System testing strips (Nipro Diagnostics (UK) Ltd)',
				'TRUEyou testing strips (Nipro Diagnostics (UK) Ltd)',
				'WaveSense JAZZ testing strips (AgaMatrix Europe Ltd)',
				'WaveSense JAZZ Duo testing strips (AgaMatrix Europe Ltd)',
				'Accu-Chek Advantage  Meter  ',
				'Accu-Chek Aviva  Blood Glucose Testing System  ',
				'Accu-Chek Aviva Blood Glucose Testing System',
				'Accu-Chek Compact  Meter  ',
				'Accu-Chek Compact Meter',
				'Accu-Chek Compact Plus  Blood Glucose Testing System  ',
				'Accu-Chek Compact Plus Blood Glucose Testing System',
				'Ascensia Microfill biosensor strips [BAYER]',
				'Bm Test 5L  Reagent Strips  ',
				'Bm-Stix  Test Strips  ',
				'Glucorx  Strips  ',
				'GlucoRx Original testing strips (Disposable Medical Equipment Ltd)',
				'GlucoRx Original testing strips (GlucoRx Ltd)',
				'GlucoRx testing strips (Disposable Medical Equipment Ltd)',
				'Glucotrend  Test Strips  ',
				'Glucotrend color.strip(s) [ROCHE DIAG]',
				'Glucotrend Plus  Reagent Strips  ',
				'Hypoguard Ga  Reagent Strips  ',

				-- KETONE STRIPS
				'Ketostix testing strips (Bayer Diagnostics Manufacturing Ltd)',
				'Mission Ketone testing strips 1K (Spirit Healthcare Ltd'
			)
		)
		AND
		SK_PatientID IN(SELECT SK_PatientID FROM ceg.GenomicsData WHERE CCG = '08V')

) AS TestingStripsEarliest


WHERE rn = 1

------------------ Insulin pump earliest ever ----------------------------------------------------------------------------------------------

SELECT
	 [SK_PatientID]
	,[ClinicalCode] InsulinPumpEarliestCode
	,[EventDate] InsulinPumpDateRecorded
	--,[Value] eGFR_EarliestValue
	--,[Units] eGFR_EarliestUnits

INTO  #InsulinPumpEarliest

FROM
(
	SELECT

		 [SK_PatientID]
		,[ClinicalCode]
		,[EventDate]
		--,[Value]
		--,[Units]
		,row_number() over(partition by [SK_PatientID] order by [EventDate] ASC) as rn

	FROM  [08V].[GPEncounter]
	WHERE
		(
			[ClinicalCode] COLLATE Latin1_General_CS_AS LIKE '7L100%'	-- % to cover Synonym
		)
		AND
		SK_PatientID IN(SELECT SK_PatientID FROM ceg.GenomicsData WHERE CCG = '08V')

)as InsulinPumpEarliest

WHERE rn = 1

------------------ Metformin adverse reactions earliest ever ----------------------------------------------------------------------------------------------

SELECT
	 [SK_PatientID]
	,[ClinicalCode] MetforminAE_EarliestCode
	,[EventDate] MetforminAE_EarliestDateRecorded
	--,[Value] MetforminAE_EarliestValue
	--,[Units] MetforminAE_EarliestUnits

INTO  #MetforminAE_Earliest

FROM
(
	SELECT

		 [SK_PatientID]
		,[ClinicalCode]
		,[EventDate]
		--,[Value]
		--,[Units]
		,row_number() over(partition by [SK_PatientID] order by [EventDate] ASC) as rn

	FROM  [08V].[GPEncounter]
	WHERE
		(
			[ClinicalCode] COLLATE Latin1_General_CS_AS LIKE 'U6023%'	-- % to cover Synonym
			OR
			[ClinicalCode] COLLATE Latin1_General_CS_AS IN ('8I7B', 'TJ23A')
		)
		AND
		SK_PatientID IN(SELECT SK_PatientID FROM ceg.GenomicsData WHERE CCG = '08V')

)as MetforminAE_Earliest

WHERE rn = 1


-------------- Treatment of diabetic neuropthay  ------------------------------------------------------------------------------

---------------- Prescribed within previous 12m  ------------------------------------------------------------------------

SELECT
	 SK_PatientID
	,[MedicationTerm]
	,IssueDate


INTO  #NeuropathyPrev12m

FROM
(
	SELECT

		 SK_PatientID
		,[MedicationTerm]
		,IssueDate
		,row_number() over(partition by [SK_PatientID] order by IssueDate DESC) as rn

	FROM
		[08V].[GPMedication]

	WHERE
		(
			BNFChapter = '6.1.5'
			OR
			MedicationTerm IN
			(
				'Amitriptyline Hydrochloride  Capsules  25 mg',
				'Amitriptyline Hydrochloride  Capsules  50 mg',
				'Amitriptyline Hydrochloride  Capsules  75 mg',
				'Amitriptyline Hydrochloride  Mixture  Sugar Free 10 mg/5 ml',
				'Amitriptyline Sr  Capsules  75 mg'
			)
		)
		AND
		IssueDate BETWEEN DATEADD(month,-12, @LastRefreshDate) AND 	@LastRefreshDate
		AND
		SK_PatientID IN(SELECT SK_PatientID FROM ceg.GenomicsData WHERE CCG = '08V')

) AS NeuropathyPrev12m


WHERE rn = 1

---------------- Prescribed earliest issue date ever --------------------------------------------------------------------

SELECT
	 SK_PatientID
	,[MedicationTerm]
	,IssueDate


INTO  #NeuropathyEarliest

FROM
(
	SELECT

		 SK_PatientID
		,[MedicationTerm]
		,IssueDate
		,row_number() over(partition by [SK_PatientID] order by IssueDate ASC) as rn

	FROM
		[08V].[GPMedication]

	WHERE
		(
			BNFChapter = '6.1.5'
			OR
			MedicationTerm IN
			(
				'Amitriptyline Hydrochloride  Capsules  25 mg',
				'Amitriptyline Hydrochloride  Capsules  50 mg',
				'Amitriptyline Hydrochloride  Capsules  75 mg',
				'Amitriptyline Hydrochloride  Mixture  Sugar Free 10 mg/5 ml',
				'Amitriptyline Sr  Capsules  75 mg'
			)
		)
		AND
		SK_PatientID IN(SELECT SK_PatientID FROM ceg.GenomicsData WHERE CCG = '08V')

) AS NeuropathyEarliest


WHERE rn = 1


--------CVD -------------------------------------------------------------------------------------------------------------------------------------------

-------------- Cardiac glycosides  -----------------------------------------------------------------------------------------------------------

---------------- Prescribed within previous 12m  ------------------------------------------------------------------------

SELECT
	 SK_PatientID
	,[MedicationTerm]
	,IssueDate


INTO  #CardiacGlycosidePrev12m

FROM
(
	SELECT

		 SK_PatientID
		,[MedicationTerm]
		,IssueDate
		,row_number() over(partition by [SK_PatientID] order by IssueDate DESC) as rn

	FROM
		[08V].[GPMedication]

	WHERE
		(
			BNFChapter = '2.1.1'
		)
		AND
		IssueDate BETWEEN DATEADD(month,-12, @LastRefreshDate) AND 	@LastRefreshDate
		AND
		SK_PatientID IN(SELECT SK_PatientID FROM ceg.GenomicsData WHERE CCG = '08V')

) AS CardiacGlycosidePrev12m


WHERE rn = 1

---------------- Prescribed earliest issue date ever --------------------------------------------------------------------

SELECT
	 SK_PatientID
	,[MedicationTerm]
	,IssueDate


INTO  #CardiacGlycosideEarliest

FROM
(
	SELECT

		 SK_PatientID
		,[MedicationTerm]
		,IssueDate
		,row_number() over(partition by [SK_PatientID] order by IssueDate ASC) as rn

	FROM
		[08V].[GPMedication]

	WHERE
		(
			BNFChapter = '2.1.1'
		)
		AND
		SK_PatientID IN(SELECT SK_PatientID FROM ceg.GenomicsData WHERE CCG = '08V')

) AS CardiacGlycosideEarliest


WHERE rn = 1

-------------- Diruetics thiazides  -----------------------------------------------------------------------------------------------------------

---------------- Prescribed within previous 12m  ------------------------------------------------------------------------

SELECT
	 SK_PatientID
	,[MedicationTerm]
	,IssueDate


INTO  #ThiazidesPrev12m

FROM
(
	SELECT

		 SK_PatientID
		,[MedicationTerm]
		,IssueDate
		,row_number() over(partition by [SK_PatientID] order by IssueDate DESC) as rn

	FROM
		[08V].[GPMedication]

	WHERE
		(
			BNFChapter = '2.2.1'
		)
		AND
		IssueDate BETWEEN DATEADD(month,-12, @LastRefreshDate) AND 	@LastRefreshDate
		AND
		SK_PatientID IN(SELECT SK_PatientID FROM ceg.GenomicsData WHERE CCG = '08V')

) AS ThiazidesPrev12m


WHERE rn = 1

---------------- Prescribed earliest issue date ever --------------------------------------------------------------------

SELECT
	 SK_PatientID
	,[MedicationTerm]
	,IssueDate


INTO  #ThiazidesEarliest

FROM
(
	SELECT

		 SK_PatientID
		,[MedicationTerm]
		,IssueDate
		,row_number() over(partition by [SK_PatientID] order by IssueDate ASC) as rn

	FROM
		[08V].[GPMedication]

	WHERE
		(
			BNFChapter = '2.2.1'
		)
		AND
		SK_PatientID IN(SELECT SK_PatientID FROM ceg.GenomicsData WHERE CCG = '08V')

) AS ThiazidesEarliest


WHERE rn = 1

-------------- Loop diuretics  -----------------------------------------------------------------------------------------------------------

---------------- Prescribed within previous 12m  ------------------------------------------------------------------------

SELECT
	 SK_PatientID
	,[MedicationTerm]
	,IssueDate


INTO  #LoopDirueticsPrev12m

FROM
(
	SELECT

		 SK_PatientID
		,[MedicationTerm]
		,IssueDate
		,row_number() over(partition by [SK_PatientID] order by IssueDate DESC) as rn

	FROM
		[08V].[GPMedication]

	WHERE
		(
			BNFChapter = '2.2.2'
		)
		AND
		IssueDate BETWEEN DATEADD(month,-12, @LastRefreshDate) AND 	@LastRefreshDate
		AND
		SK_PatientID IN(SELECT SK_PatientID FROM ceg.GenomicsData WHERE CCG = '08V')

) AS LoopDirueticsPrev12m


WHERE rn = 1

---------------- Prescribed earliest issue date ever --------------------------------------------------------------------

SELECT
	 SK_PatientID
	,[MedicationTerm]
	,IssueDate


INTO  #LoopDirueticsEarliest

FROM
(
	SELECT

		 SK_PatientID
		,[MedicationTerm]
		,IssueDate
		,row_number() over(partition by [SK_PatientID] order by IssueDate ASC) as rn

	FROM
		[08V].[GPMedication]

	WHERE
		(
			BNFChapter = '2.2.2'
		)
		AND
		SK_PatientID IN(SELECT SK_PatientID FROM ceg.GenomicsData WHERE CCG = '08V')

) AS LoopDirueticsEarliest


WHERE rn = 1

-------------- Potassium sparing diuretics  -----------------------------------------------------------------------------------------------------------

---------------- Prescribed within previous 12m  ------------------------------------------------------------------------

SELECT
	 SK_PatientID
	,[MedicationTerm]
	,IssueDate


INTO  #PotassiumSparingPrev12m

FROM
(
	SELECT

		 SK_PatientID
		,[MedicationTerm]
		,IssueDate
		,row_number() over(partition by [SK_PatientID] order by IssueDate DESC) as rn

	FROM
		[08V].[GPMedication]

	WHERE
		(
			BNFChapter = '2.2.3'
		)
		AND
		IssueDate BETWEEN DATEADD(month,-12, @LastRefreshDate) AND 	@LastRefreshDate
		AND
		SK_PatientID IN(SELECT SK_PatientID FROM ceg.GenomicsData WHERE CCG = '08V')

) AS PotassiumSparingPrev12m


WHERE rn = 1

---------------- Prescribed earliest issue date ever --------------------------------------------------------------------

SELECT
	 SK_PatientID
	,[MedicationTerm]
	,IssueDate


INTO  #PotassiumSparingEarliest

FROM
(
	SELECT

		 SK_PatientID
		,[MedicationTerm]
		,IssueDate
		,row_number() over(partition by [SK_PatientID] order by IssueDate ASC) as rn

	FROM
		[08V].[GPMedication]

	WHERE
		(
			BNFChapter = '2.2.3'
		)
		AND
		SK_PatientID IN(SELECT SK_PatientID FROM ceg.GenomicsData WHERE CCG = '08V')

) AS PotassiumSparingEarliest


WHERE rn = 1

-------------- Potassium sparing diuretic combinations  -----------------------------------------------------------------------------------------------------------

---------------- Prescribed within previous 12m  ------------------------------------------------------------------------

SELECT
	 SK_PatientID
	,[MedicationTerm]
	,IssueDate


INTO  #DiureticCombiPrev12m

FROM
(
	SELECT

		 SK_PatientID
		,[MedicationTerm]
		,IssueDate
		,row_number() over(partition by [SK_PatientID] order by IssueDate DESC) as rn

	FROM
		[08V].[GPMedication]

	WHERE
		(
			BNFChapter = '2.2.4'
		)
		AND
		IssueDate BETWEEN DATEADD(month,-12, @LastRefreshDate) AND 	@LastRefreshDate
		AND
		SK_PatientID IN(SELECT SK_PatientID FROM ceg.GenomicsData WHERE CCG = '08V')

) AS DiureticCombiPrev12m


WHERE rn = 1

---------------- Prescribed earliest issue date ever --------------------------------------------------------------------

SELECT
	 SK_PatientID
	,[MedicationTerm]
	,IssueDate


INTO  #DiureticCombiEarliest

FROM
(
	SELECT

		 SK_PatientID
		,[MedicationTerm]
		,IssueDate
		,row_number() over(partition by [SK_PatientID] order by IssueDate ASC) as rn

	FROM
		[08V].[GPMedication]

	WHERE
		(
			BNFChapter = '2.2.4'
		)
		AND
		SK_PatientID IN(SELECT SK_PatientID FROM ceg.GenomicsData WHERE CCG = '08V')

) AS DiureticCombiEarliest


WHERE rn = 1

-------------- Diruetics with potassium  -----------------------------------------------------------------------------------------------------------

---------------- Prescribed within previous 12m  ------------------------------------------------------------------------

SELECT
	 SK_PatientID
	,[MedicationTerm]
	,IssueDate


INTO  #DiumidePrev12m

FROM
(
	SELECT

		 SK_PatientID
		,[MedicationTerm]
		,IssueDate
		,row_number() over(partition by [SK_PatientID] order by IssueDate DESC) as rn

	FROM
		[08V].[GPMedication]

	WHERE
		(
			BNFChapter = '2.2.8'
		)
		AND
		IssueDate BETWEEN DATEADD(month,-12, @LastRefreshDate) AND 	@LastRefreshDate
		AND
		SK_PatientID IN(SELECT SK_PatientID FROM ceg.GenomicsData WHERE CCG = '08V')

) AS DiumidePrev12m


WHERE rn = 1

---------------- Prescribed earliest issue date ever --------------------------------------------------------------------

SELECT
	 SK_PatientID
	,[MedicationTerm]
	,IssueDate


INTO  #DiumideEarliest

FROM
(
	SELECT

		 SK_PatientID
		,[MedicationTerm]
		,IssueDate
		,row_number() over(partition by [SK_PatientID] order by IssueDate ASC) as rn

	FROM
		[08V].[GPMedication]

	WHERE
		(
			BNFChapter = '2.2.8'
		)
		AND
		SK_PatientID IN(SELECT SK_PatientID FROM ceg.GenomicsData WHERE CCG = '08V')

) AS DiumideEarliest


WHERE rn = 1

-------------- Drugs for arrhythmias  -----------------------------------------------------------------------------------------------------------

---------------- Prescribed within previous 12m  ------------------------------------------------------------------------

SELECT
	 SK_PatientID
	,[MedicationTerm]
	,IssueDate


INTO  #AntiArrhythmicsPrev12m

FROM
(
	SELECT

		 SK_PatientID
		,[MedicationTerm]
		,IssueDate
		,row_number() over(partition by [SK_PatientID] order by IssueDate DESC) as rn

	FROM
		[08V].[GPMedication]

	WHERE
		(
			BNFChapter = '2.3'
		)
		AND
		IssueDate BETWEEN DATEADD(month,-12, @LastRefreshDate) AND 	@LastRefreshDate
		AND
		SK_PatientID IN(SELECT SK_PatientID FROM ceg.GenomicsData WHERE CCG = '08V')

) AS AntiArrhythmicsPrev12m


WHERE rn = 1

---------------- Prescribed earliest issue date ever --------------------------------------------------------------------

SELECT
	 SK_PatientID
	,[MedicationTerm]
	,IssueDate


INTO  #AntiArrhythmicsEarliest

FROM
(
	SELECT

		 SK_PatientID
		,[MedicationTerm]
		,IssueDate
		,row_number() over(partition by [SK_PatientID] order by IssueDate ASC) as rn

	FROM
		[08V].[GPMedication]

	WHERE
		(
			BNFChapter = '2.3'
		)
		AND
		SK_PatientID IN(SELECT SK_PatientID FROM ceg.GenomicsData WHERE CCG = '08V')

) AS AntiArrhythmicsEarliest


WHERE rn = 1

-------------- Beta-adrenoceptor blockers  -----------------------------------------------------------------------------------------------------------

---------------- Prescribed within previous 12m  ------------------------------------------------------------------------

SELECT
	 SK_PatientID
	,[MedicationTerm]
	,IssueDate


INTO  #BetaBlockersPrev12m

FROM
(
	SELECT

		 SK_PatientID
		,[MedicationTerm]
		,IssueDate
		,row_number() over(partition by [SK_PatientID] order by IssueDate DESC) as rn

	FROM
		[08V].[GPMedication]

	WHERE
		(
			BNFChapter = '2.4'
			OR
			MedicationTerm LIKE 'Propranolol Hydrochloride  Capsules%'
		)
		AND
		IssueDate BETWEEN DATEADD(month,-12, @LastRefreshDate) AND 	@LastRefreshDate
		AND
		SK_PatientID IN(SELECT SK_PatientID FROM ceg.GenomicsData WHERE CCG = '08V')

) AS BetaBlockersPrev12m


WHERE rn = 1

---------------- Prescribed earliest issue date ever --------------------------------------------------------------------

SELECT
	 SK_PatientID
	,[MedicationTerm]
	,IssueDate


INTO  #BetaBlockersEarliest

FROM
(
	SELECT

		 SK_PatientID
		,[MedicationTerm]
		,IssueDate
		,row_number() over(partition by [SK_PatientID] order by IssueDate ASC) as rn

	FROM
		[08V].[GPMedication]

	WHERE
		(
			BNFChapter = '2.4'
			OR
			MedicationTerm LIKE 'Propranolol Hydrochloride  Capsules%'
		)
		AND
		SK_PatientID IN(SELECT SK_PatientID FROM ceg.GenomicsData WHERE CCG = '08V')

) AS BetaBlockersEarliest


WHERE rn = 1

-------------- Anti-hypertensives vasodilators  -----------------------------------------------------------------------------------------------------------

---------------- Prescribed within previous 12m  ------------------------------------------------------------------------

SELECT
	 SK_PatientID
	,[MedicationTerm]
	,IssueDate


INTO  #VasodilatorsPrev12m

FROM
(
	SELECT

		 SK_PatientID
		,[MedicationTerm]
		,IssueDate
		,row_number() over(partition by [SK_PatientID] order by IssueDate DESC) as rn

	FROM
		[08V].[GPMedication]

	WHERE
		(
			BNFChapter = '2.5.1'
		)
		AND
		IssueDate BETWEEN DATEADD(month,-12, @LastRefreshDate) AND 	@LastRefreshDate
		AND
		SK_PatientID IN(SELECT SK_PatientID FROM ceg.GenomicsData WHERE CCG = '08V')

) AS VasodilatorsPrev12m


WHERE rn = 1

---------------- Prescribed earliest issue date ever --------------------------------------------------------------------

SELECT
	 SK_PatientID
	,[MedicationTerm]
	,IssueDate


INTO  #VasodilatorsEarliest

FROM
(
	SELECT

		 SK_PatientID
		,[MedicationTerm]
		,IssueDate
		,row_number() over(partition by [SK_PatientID] order by IssueDate ASC) as rn

	FROM
		[08V].[GPMedication]

	WHERE
		(
			BNFChapter = '2.5.1'
		)
		AND
		SK_PatientID IN(SELECT SK_PatientID FROM ceg.GenomicsData WHERE CCG = '08V')

) AS VasodilatorsEarliest


WHERE rn = 1

-------------- Centrally acting anti-hypertensives  -----------------------------------------------------------------------------------------------------------

---------------- Prescribed within previous 12m  ------------------------------------------------------------------------

SELECT
	 SK_PatientID
	,[MedicationTerm]
	,IssueDate


INTO  #CentralantihypertensivePrev12m

FROM
(
	SELECT

		 SK_PatientID
		,[MedicationTerm]
		,IssueDate
		,row_number() over(partition by [SK_PatientID] order by IssueDate DESC) as rn

	FROM
		[08V].[GPMedication]

	WHERE
		(
			BNFChapter = '2.5.2'
			OR
			MedicationTerm LIKE 'Methyldopa  Capsules%'
		)
		AND
		IssueDate BETWEEN DATEADD(month,-12, @LastRefreshDate) AND 	@LastRefreshDate
		AND
		SK_PatientID IN(SELECT SK_PatientID FROM ceg.GenomicsData WHERE CCG = '08V')

) AS CentralantihypertensivePrev12m

WHERE rn = 1

---------------- Prescribed earliest issue date ever --------------------------------------------------------------------

SELECT
	 SK_PatientID
	,[MedicationTerm]
	,IssueDate


INTO  #CentralantihypertensiveEarliest

FROM
(
	SELECT

		 SK_PatientID
		,[MedicationTerm]
		,IssueDate
		,row_number() over(partition by [SK_PatientID] order by IssueDate ASC) as rn

	FROM
		[08V].[GPMedication]

	WHERE
		(
			BNFChapter = '2.5.2'
			OR
			MedicationTerm LIKE 'Methyldopa  Capsules%'
		)
		AND
		SK_PatientID IN(SELECT SK_PatientID FROM ceg.GenomicsData WHERE CCG = '08V')

) AS CentralantihypertensiveEarliest

WHERE rn = 1

-------------- Alpha adrenoceptor blockers  -----------------------------------------------------------------------------------------------------------

---------------- Prescribed within previous 12m  ------------------------------------------------------------------------

SELECT
	 SK_PatientID
	,[MedicationTerm]
	,IssueDate


INTO  #AlphaBlockersPrev12m

FROM
(
	SELECT

		 SK_PatientID
		,[MedicationTerm]
		,IssueDate
		,row_number() over(partition by [SK_PatientID] order by IssueDate DESC) as rn

	FROM
		[08V].[GPMedication]

	WHERE
		(
			BNFChapter = '2.5.4'
			OR
			MedicationTerm IN
			(
				'Terazosin Hydrochloride  Starter Pack  7 x 1 mg, 14 x 2 mg, 7 x 5 mg'
			)
		)
		AND
		IssueDate BETWEEN DATEADD(month,-12, @LastRefreshDate) AND 	@LastRefreshDate
		AND
		SK_PatientID IN(SELECT SK_PatientID FROM ceg.GenomicsData WHERE CCG = '08V')

) AS AlphaBlockersPrev12m

WHERE rn = 1

---------------- Prescribed earliest issue date ever --------------------------------------------------------------------

SELECT
	 SK_PatientID
	,[MedicationTerm]
	,IssueDate


INTO  #AlphaBlockersEarliest

FROM
(
	SELECT

		 SK_PatientID
		,[MedicationTerm]
		,IssueDate
		,row_number() over(partition by [SK_PatientID] order by IssueDate ASC) as rn

	FROM
		[08V].[GPMedication]

	WHERE
		(
			BNFChapter = '2.5.4'
			OR
			MedicationTerm IN
			(
				'Terazosin Hydrochloride  Starter Pack  7 x 1 mg, 14 x 2 mg, 7 x 5 mg'
			)
		)
		AND
		SK_PatientID IN(SELECT SK_PatientID FROM ceg.GenomicsData WHERE CCG = '08V')

) AS AlphaBlockersEarliest

WHERE rn = 1

-------------- Angiotensin Converting Enzyme Inhibitors  -----------------------------------------------------------------------------------------------------------

---------------- Prescribed within previous 12m  ------------------------------------------------------------------------

SELECT
	 SK_PatientID
	,[MedicationTerm]
	,IssueDate

INTO  #ACE_InhibitorsPrev12m

FROM
(
	SELECT

		 SK_PatientID
		,[MedicationTerm]
		,IssueDate
		,row_number() over(partition by [SK_PatientID] order by IssueDate DESC) as rn
	FROM
		[08V].[GPMedication]

	WHERE
		(
			BNFChapter = '2.5.5.1'
			OR
			MedicationTerm IN
			(
				'Ramipril  Titration pack  7 x 2.5 mg, 21 x 5 mg, 7 x 10 mg',
				'Tritace  Tablet Titration Pack  7 x 2.5 mg, 21 x 5 mg, 7 x 10 mg',
				'Tritace titration pack tablets (Sanofi)'
			)
		)
		AND
		IssueDate BETWEEN DATEADD(month,-12, @LastRefreshDate) AND 	@LastRefreshDate
		AND
		SK_PatientID IN(SELECT SK_PatientID FROM ceg.GenomicsData WHERE CCG = '08V')

) AS ACE_InhibitorsPrev12m

WHERE rn = 1

---------------- Prescribed earliest issue date ever --------------------------------------------------------------------

SELECT
	 SK_PatientID
	,[MedicationTerm]
	,IssueDate

INTO  #ACE_InhibitorsEarliest

FROM
(
	SELECT
		 SK_PatientID
		,[MedicationTerm]
		,IssueDate
		,row_number() over(partition by [SK_PatientID] order by IssueDate ASC) as rn

	FROM
		[08V].[GPMedication]

	WHERE
		(
			BNFChapter = '2.5.5.1'
			OR
			MedicationTerm IN
			(
				'Ramipril  Titration pack  7 x 2.5 mg, 21 x 5 mg, 7 x 10 mg',
				'Tritace  Tablet Titration Pack  7 x 2.5 mg, 21 x 5 mg, 7 x 10 mg',
				'Tritace titration pack tablets (Sanofi)'
			)
		)
		AND
		SK_PatientID IN(SELECT SK_PatientID FROM ceg.GenomicsData WHERE CCG = '08V')

) AS ACE_InhibitorsEarliest

WHERE rn = 1

-------------- Angiotensin II antagonists  -----------------------------------------------------------------------------------------------------------

---------------- Prescribed within previous 12m  ------------------------------------------------------------------------

SELECT
	 SK_PatientID
	,[MedicationTerm]
	,IssueDate

INTO  #AngiotensinAntagonistsPrev12m

FROM
(
	SELECT

		 SK_PatientID
		,[MedicationTerm]
		,IssueDate
		,row_number() over(partition by [SK_PatientID] order by IssueDate DESC) as rn
	FROM
		[08V].[GPMedication]

	WHERE
		(
			BNFChapter = '2.5.5.2'
		)
		AND
		IssueDate BETWEEN DATEADD(month,-12, @LastRefreshDate) AND 	@LastRefreshDate
		AND
		SK_PatientID IN(SELECT SK_PatientID FROM ceg.GenomicsData WHERE CCG = '08V')

) AS AngiotensinAntagonistsPrev12m

WHERE rn = 1

---------------- Prescribed earliest issue date ever --------------------------------------------------------------------

SELECT
	 SK_PatientID
	,[MedicationTerm]
	,IssueDate

INTO  #AngiotensinAntagonistsEarliest

FROM
(
	SELECT
		 SK_PatientID
		,[MedicationTerm]
		,IssueDate
		,row_number() over(partition by [SK_PatientID] order by IssueDate ASC) as rn

	FROM
		[08V].[GPMedication]

	WHERE
		(
			BNFChapter = '2.5.5.2'
		)
		AND
		SK_PatientID IN(SELECT SK_PatientID FROM ceg.GenomicsData WHERE CCG = '08V')

) AS AngiotensinAntagonistsEarliest

WHERE rn = 1

-------------- Nitrates  -----------------------------------------------------------------------------------------------------------

---------------- Prescribed within previous 12m  ------------------------------------------------------------------------

SELECT
	 SK_PatientID
	,[MedicationTerm]
	,IssueDate

INTO  #NitratesPrev12m

FROM
(
	SELECT

		 SK_PatientID
		,[MedicationTerm]
		,IssueDate
		,row_number() over(partition by [SK_PatientID] order by IssueDate DESC) as rn
	FROM
		[08V].[GPMedication]

	WHERE
		(
			BNFChapter = '2.6.1'
		)
		AND
		IssueDate BETWEEN DATEADD(month,-12, @LastRefreshDate) AND 	@LastRefreshDate
		AND
		SK_PatientID IN(SELECT SK_PatientID FROM ceg.GenomicsData WHERE CCG = '08V')

) AS NitratesPrev12m

WHERE rn = 1

---------------- Prescribed earliest issue date ever --------------------------------------------------------------------

SELECT
	 SK_PatientID
	,[MedicationTerm]
	,IssueDate

INTO  #NitratesEarliest

FROM
(
	SELECT
		 SK_PatientID
		,[MedicationTerm]
		,IssueDate
		,row_number() over(partition by [SK_PatientID] order by IssueDate ASC) as rn

	FROM
		[08V].[GPMedication]

	WHERE
		(
			BNFChapter = '2.6.1'
		)
		AND
		SK_PatientID IN(SELECT SK_PatientID FROM ceg.GenomicsData WHERE CCG = '08V')

) AS NitratesEarliest

WHERE rn = 1

-------------- Calcium channel blockers  -----------------------------------------------------------------------------------------------------------

---------------- Prescribed within previous 12m  ------------------------------------------------------------------------

SELECT
	 SK_PatientID
	,[MedicationTerm]
	,IssueDate

INTO  #CalciumChannelPrev12m

FROM
(
	SELECT

		 SK_PatientID
		,[MedicationTerm]
		,IssueDate
		,row_number() over(partition by [SK_PatientID] order by IssueDate DESC) as rn
	FROM
		[08V].[GPMedication]

	WHERE
		(
			BNFChapter = '2.6.2'
			OR
			MedicationTerm IN
			(
				'Amlodipine Besilate  Tablets  5 mg',
				'Amlodipine Besilate  Tablets  10 mg',
				'Amlodipine Besylate  Tablets  5 mg',
				'Amlodipine Besylate  Tablets  10 mg',
				'Diltiazem Hydrochloride  Tablets  60 mg',
				'Nifedipine  Tablets  10 mg',
				'Nifedipine  Tablets  20 mg'
			)
		)
		AND
		IssueDate BETWEEN DATEADD(month,-12, @LastRefreshDate) AND 	@LastRefreshDate
		AND
		SK_PatientID IN(SELECT SK_PatientID FROM ceg.GenomicsData WHERE CCG = '08V')

) AS CalciumChannelPrev12m

WHERE rn = 1


---------------- Prescribed earliest issue date ever --------------------------------------------------------------------

SELECT
	 SK_PatientID
	,[MedicationTerm]
	,IssueDate

INTO  #CalciumChannelEarliest

FROM
(
	SELECT
		 SK_PatientID
		,[MedicationTerm]
		,IssueDate
		,row_number() over(partition by [SK_PatientID] order by IssueDate ASC) as rn

	FROM
		[08V].[GPMedication]

	WHERE
		(
			BNFChapter = '2.6.2'
			OR
			MedicationTerm IN
			(
				'Amlodipine Besilate  Tablets  5 mg',
				'Amlodipine Besilate  Tablets  10 mg',
				'Amlodipine Besylate  Tablets  5 mg',
				'Amlodipine Besylate  Tablets  10 mg',
				'Diltiazem Hydrochloride  Tablets  60 mg',
				'Nifedipine  Tablets  10 mg',
				'Nifedipine  Tablets  20 mg'
			)
		)
		AND
		SK_PatientID IN(SELECT SK_PatientID FROM ceg.GenomicsData WHERE CCG = '08V')

) AS CalciumChannelEarliest

WHERE rn = 1

-------------- Other Antianginal  -----------------------------------------------------------------------------------------------------------

---------------- Prescribed within previous 12m  ------------------------------------------------------------------------

SELECT
	 SK_PatientID
	,[MedicationTerm]
	,IssueDate

INTO  #AntianginalPrev12m

FROM
(
	SELECT

		 SK_PatientID
		,[MedicationTerm]
		,IssueDate
		,row_number() over(partition by [SK_PatientID] order by IssueDate DESC) as rn
	FROM
		[08V].[GPMedication]

	WHERE
		(
			BNFChapter = '2.6.3'
		)
		AND
		IssueDate BETWEEN DATEADD(month,-12, @LastRefreshDate) AND 	@LastRefreshDate
		AND
		SK_PatientID IN(SELECT SK_PatientID FROM ceg.GenomicsData WHERE CCG = '08V')

) AS AntianginalPrev12m

WHERE rn = 1

---------------- Prescribed earliest issue date ever --------------------------------------------------------------------

SELECT
	 SK_PatientID
	,[MedicationTerm]
	,IssueDate

INTO  #AntianginalEarliest

FROM
(
	SELECT
		 SK_PatientID
		,[MedicationTerm]
		,IssueDate
		,row_number() over(partition by [SK_PatientID] order by IssueDate ASC) as rn

	FROM
		[08V].[GPMedication]

	WHERE
		(
			BNFChapter = '2.6.3'
		)
		AND
		SK_PatientID IN(SELECT SK_PatientID FROM ceg.GenomicsData WHERE CCG = '08V')

) AS AntianginalEarliest

WHERE rn = 1

-------------- Peripheral vasodilators  -----------------------------------------------------------------------------------------------------------

---------------- Prescribed within previous 12m  ------------------------------------------------------------------------

SELECT
	 SK_PatientID
	,[MedicationTerm]
	,IssueDate

INTO  #PeripheralVasodilatorsPrev12m

FROM
(
	SELECT

		 SK_PatientID
		,[MedicationTerm]
		,IssueDate
		,row_number() over(partition by [SK_PatientID] order by IssueDate DESC) as rn
	FROM
		[08V].[GPMedication]

	WHERE
		(
			BNFChapter = '2.6.4'
		)
		AND
		IssueDate BETWEEN DATEADD(month,-12, @LastRefreshDate) AND 	@LastRefreshDate
		AND
		SK_PatientID IN(SELECT SK_PatientID FROM ceg.GenomicsData WHERE CCG = '08V')

) AS PeripheralVasodilatorsPrev12m

WHERE rn = 1

---------------- Prescribed earliest issue date ever --------------------------------------------------------------------

SELECT
	 SK_PatientID
	,[MedicationTerm]
	,IssueDate

INTO  #PeripheralVasodilatorsEarliest

FROM
(
	SELECT
		 SK_PatientID
		,[MedicationTerm]
		,IssueDate
		,row_number() over(partition by [SK_PatientID] order by IssueDate ASC) as rn

	FROM
		[08V].[GPMedication]

	WHERE
		(
			BNFChapter = '2.6.4'
		)
		AND
		SK_PatientID IN(SELECT SK_PatientID FROM ceg.GenomicsData WHERE CCG = '08V')

) AS PeripheralVasodilatorsEarliest

WHERE rn = 1

-------------- Oral Anticoagulants  -----------------------------------------------------------------------------------------------------------

---------------- Prescribed within previous 12m  ------------------------------------------------------------------------

SELECT
	 SK_PatientID
	,[MedicationTerm]
	,IssueDate

INTO  #OralAnticoagulantsPrev12m

FROM
(
	SELECT

		 SK_PatientID
		,[MedicationTerm]
		,IssueDate
		,row_number() over(partition by [SK_PatientID] order by IssueDate DESC) as rn
	FROM
		[08V].[GPMedication]

	WHERE
		(
			BNFChapter = '2.8.2'
		)
		AND
		IssueDate BETWEEN DATEADD(month,-12, @LastRefreshDate) AND 	@LastRefreshDate
		AND
		SK_PatientID IN(SELECT SK_PatientID FROM ceg.GenomicsData WHERE CCG = '08V')

) AS OralAnticoagulantsPrev12m

WHERE rn = 1

---------------- Prescribed earliest issue date ever --------------------------------------------------------------------

SELECT
	 SK_PatientID
	,[MedicationTerm]
	,IssueDate

INTO  #OralAnticoagulantsEarliest

FROM
(
	SELECT
		 SK_PatientID
		,[MedicationTerm]
		,IssueDate
		,row_number() over(partition by [SK_PatientID] order by IssueDate ASC) as rn

	FROM
		[08V].[GPMedication]

	WHERE
		(
			BNFChapter = '2.8.2'
		)
		AND
		SK_PatientID IN(SELECT SK_PatientID FROM ceg.GenomicsData WHERE CCG = '08V')

) AS OralAnticoagulantsEarliest

WHERE rn = 1

-------------- Antiplatelet drugs  -----------------------------------------------------------------------------------------------------------

---------------- Prescribed within previous 12m  ------------------------------------------------------------------------

SELECT
	 SK_PatientID
	,[MedicationTerm]
	,IssueDate

INTO  #AntiplateletPrev12m

FROM
(
	SELECT

		 SK_PatientID
		,[MedicationTerm]
		,IssueDate
		,row_number() over(partition by [SK_PatientID] order by IssueDate DESC) as rn
	FROM
		[08V].[GPMedication]

	WHERE
		(
			BNFChapter = '2.9'
			OR
			MedicationTerm IN
			(
				'Soluble Aspirin  Tablets  300 mg',
				'Soluble Aspirin Paediatric  Tablets  75 mg'
			)
		)
		AND
		IssueDate BETWEEN DATEADD(month,-12, @LastRefreshDate) AND 	@LastRefreshDate
		AND
		SK_PatientID IN(SELECT SK_PatientID FROM ceg.GenomicsData WHERE CCG = '08V')

) AS AntiplateletPrev12m

WHERE rn = 1

---------------- Prescribed earliest issue date ever --------------------------------------------------------------------

SELECT
	 SK_PatientID
	,[MedicationTerm]
	,IssueDate

INTO  #AntiplateletEarliest

FROM
(
	SELECT
		 SK_PatientID
		,[MedicationTerm]
		,IssueDate
		,row_number() over(partition by [SK_PatientID] order by IssueDate ASC) as rn

	FROM
		[08V].[GPMedication]

	WHERE
		(
			BNFChapter = '2.9'
			OR
			MedicationTerm IN
			(
				'Soluble Aspirin  Tablets  300 mg',
				'Soluble Aspirin Paediatric  Tablets  75 mg'
			)
		)
		AND
		SK_PatientID IN(SELECT SK_PatientID FROM ceg.GenomicsData WHERE CCG = '08V')

) AS AntiplateletEarliest

WHERE rn = 1

-------------- Lipid regulating drugs  -----------------------------------------------------------------------------------------------------------

---------------- Prescribed within previous 12m  ------------------------------------------------------------------------

SELECT
	 SK_PatientID
	,[MedicationTerm]
	,IssueDate

INTO  #LipidRegulationPrev12m

FROM
(
	SELECT

		 SK_PatientID
		,[MedicationTerm]
		,IssueDate
		,row_number() over(partition by [SK_PatientID] order by IssueDate DESC) as rn
	FROM
		[08V].[GPMedication]

	WHERE
		(
			BNFChapter = '2.12'
		)
		AND
		IssueDate BETWEEN DATEADD(month,-12, @LastRefreshDate) AND 	@LastRefreshDate
		AND
		SK_PatientID IN(SELECT SK_PatientID FROM ceg.GenomicsData WHERE CCG = '08V')

) AS LipidRegulationPrev12m

WHERE rn = 1

---------------- Prescribed earliest issue date ever --------------------------------------------------------------------

SELECT
	 SK_PatientID
	,[MedicationTerm]
	,IssueDate

INTO  #LipidRegulationEarliest

FROM
(
	SELECT
		 SK_PatientID
		,[MedicationTerm]
		,IssueDate
		,row_number() over(partition by [SK_PatientID] order by IssueDate ASC) as rn

	FROM
		[08V].[GPMedication]

	WHERE
		(
			BNFChapter = '2.12'
		)
		AND
		SK_PatientID IN(SELECT SK_PatientID FROM ceg.GenomicsData WHERE CCG = '08V')

) AS LipidRegulationEarliest

WHERE rn = 1


------------------ Statin adverse reactions earliest ever ----------------------------------------------------------------------------------------------

SELECT
	 [SK_PatientID]
	,[ClinicalCode] StatinAE_EarliestCode
	,[EventDate] StatinAE_DateRecorded
	--,[Value] eGFR_EarliestValue
	--,[Units] eGFR_EarliestUnits

INTO  #StatinAE_Earliest

FROM
(
	SELECT

		 [SK_PatientID]
		,[ClinicalCode]
		,[EventDate]
		--,[Value]
		--,[Units]
		,row_number() over(partition by [SK_PatientID] order by [EventDate] ASC) as rn

	FROM  [08V].[GPEncounter]
	WHERE

		[ClinicalCode] COLLATE Latin1_General_CS_AS IN ('U60CA', '8I76', 'TJ2C', 'U60C9')
		AND
		SK_PatientID IN(SELECT SK_PatientID FROM ceg.GenomicsData WHERE CCG = '08V')

)as StatinAE_Earliest

WHERE rn = 1

------------------ Statin muscle symptoms earliest ever ----------------------------------------------------------------------------------------------

SELECT
	 [SK_PatientID]
	,[ClinicalCode] StatinMuscleEarliestCode
	,[EventDate] StatinMuscleDateRecorded
	--,[Value] eGFR_EarliestValue
	--,[Units] eGFR_EarliestUnits

INTO  #StatinMuscleEarliest

FROM
(
	SELECT

		 [SK_PatientID]
		,[ClinicalCode]
		,[EventDate]
		--,[Value]
		--,[Units]
		,row_number() over(partition by [SK_PatientID] order by [EventDate] ASC) as rn

	FROM  [08V].[GPEncounter]
	WHERE

		[ClinicalCode] COLLATE Latin1_General_CS_AS IN ('N24', 'N241', 'N2411', 'N2333')
		AND
		SK_PatientID IN(SELECT SK_PatientID FROM ceg.GenomicsData WHERE CCG = '08V')

)as StatinMuscleEarliest

WHERE rn = 1

-------------- Treatment of erectile dysfunction  -----------------------------------------------------------------------------------------------------------

---------------- Prescribed within previous 12m  ------------------------------------------------------------------------

SELECT
	 SK_PatientID
	,[MedicationTerm]
	,IssueDate

INTO  #ErectileDysfunctionPrev12m

FROM
(
	SELECT

		 SK_PatientID
		,[MedicationTerm]
		,IssueDate
		,row_number() over(partition by [SK_PatientID] order by IssueDate DESC) as rn
	FROM
		[08V].[GPMedication]

	WHERE
		(
			BNFChapter = '7.4.5'
		)
		AND
		IssueDate BETWEEN DATEADD(month,-12, @LastRefreshDate) AND 	@LastRefreshDate
		AND
		SK_PatientID IN(SELECT SK_PatientID FROM ceg.GenomicsData WHERE CCG = '08V')

) AS ErectileDysfunctionPrev12m

WHERE rn = 1

---------------- Prescribed earliest issue date ever --------------------------------------------------------------------

SELECT
	 SK_PatientID
	,[MedicationTerm]
	,IssueDate

INTO  #ErectileDysfunctionEarliest

FROM
(
	SELECT
		 SK_PatientID
		,[MedicationTerm]
		,IssueDate
		,row_number() over(partition by [SK_PatientID] order by IssueDate ASC) as rn

	FROM
		[08V].[GPMedication]

	WHERE
		(
			BNFChapter = '7.4.5'
		)
		AND
		SK_PatientID IN(SELECT SK_PatientID FROM ceg.GenomicsData WHERE CCG = '08V')

) AS ErectileDysfunctionEarliest

WHERE rn = 1

-------------- Weight loss drugs   -----------------------------------------------------------------------------------------------------------

---------------- Prescribed within previous 12m  ------------------------------------------------------------------------

SELECT
	 SK_PatientID
	,[MedicationTerm]
	,IssueDate

INTO  #WeightLossPrev12m

FROM
(
	SELECT

		 SK_PatientID
		,[MedicationTerm]
		,IssueDate
		,row_number() over(partition by [SK_PatientID] order by IssueDate DESC) as rn
	FROM
		[08V].[GPMedication]

	WHERE
		(
			BNFChapter = '4.5.1'
			OR
			MedicationTerm IN
			(
				'Orlistat 120mg capsules  ',
				'Xenical 120mg capsules (Roche Products Ltd)'
			)
		)
		AND
		IssueDate BETWEEN DATEADD(month,-12, @LastRefreshDate) AND 	@LastRefreshDate
		AND
		SK_PatientID IN(SELECT SK_PatientID FROM ceg.GenomicsData WHERE CCG = '08V')

) AS WeightLossPrev12m

WHERE rn = 1

---------------- Prescribed earliest issue date ever --------------------------------------------------------------------

SELECT
	 SK_PatientID
	,[MedicationTerm]
	,IssueDate

INTO  #WeightLossEarliest

FROM
(
	SELECT
		 SK_PatientID
		,[MedicationTerm]
		,IssueDate
		,row_number() over(partition by [SK_PatientID] order by IssueDate ASC) as rn

	FROM
		[08V].[GPMedication]

	WHERE
		(
			BNFChapter = '4.5.1'
			OR
			MedicationTerm IN
			(
				'Orlistat 120mg capsules  ',
				'Xenical 120mg capsules (Roche Products Ltd)'
			)
		)
		AND
		SK_PatientID IN(SELECT SK_PatientID FROM ceg.GenomicsData WHERE CCG = '08V')

) AS WeightLossEarliest

WHERE rn = 1

GO


--------------------------------------------------------------------------------------------------------------------------
---------------------------------------- Joining Tables ------------------------------------------------------------------
--------------------------------------------------------------------------------------------------------------------------


SELECT

	(Select CommissionerName From Dictionary.[dbo].[Commissioner] Where CommissionerCode = '08V') AS Locality
	,P.EncryptedNHSNumber
	,P.SK_PatientID

	-- PRESCRIBING
	-----Diabetes
	,InsulinsShortPrev12m.MedicationTerm AS [Insulins Short Prev12m]
	,InsulinsShortPrev12m.IssueDate

	,InsulinsShortEarliest.MedicationTerm AS [Insulins Short Earliest]
	,InsulinsShortEarliest.IssueDate

	,InsulinsLongPrev12m.MedicationTerm AS  [Insulins Long Prev12m]
	,InsulinsLongPrev12m.IssueDate

	,InsulinsLongEarliest.MedicationTerm AS [Insulins Long Earliest]
	,InsulinsLongEarliest.IssueDate

	,SulphonylureasPrev12m.MedicationTerm AS [Sulphonylureas Prev12m]
	,SulphonylureasPrev12m.IssueDate

	,SulphonylureasEarliest.MedicationTerm AS [Sulphonylureas Earliest]
	,SulphonylureasEarliest.IssueDate

	,MetforminPrev12m.MedicationTerm AS [Metformin Prev12m]
	,MetforminPrev12m.IssueDate

	,MetforminEarliest.MedicationTerm AS [Metformin Earliest]
	,MetforminEarliest.IssueDate

	--,OtherAntidiabeticPrev12m.MedicationTerm AS [Other Antidiabetic Prev12m]
	--,OtherAntidiabeticPrev12m.IssueDate

	--,OtherAntidiabeticEarliest.MedicationTerm AS [Other Antidiabetic Earliest]
	--,OtherAntidiabeticEarliest.IssueDate

	,AGIsPrev12m.MedicationTerm AS [Alpha-glucosidase inhibitors Prev12m]
	,AGIsPrev12m.IssueDate

	,AGIsEarliest.MedicationTerm AS [Alpha-glucosidase inhibitors Earliest]
	,AGIsEarliest.IssueDate

	,DPP4Prev12m.MedicationTerm AS [Dipeptidylpeptidase-4 inhibitors Prev12m]
	,DPP4Prev12m.IssueDate

	,DPP4Earliest.MedicationTerm AS [Dipeptidylpeptidase-4 inhibitors Earliest]
	,DPP4Earliest.IssueDate

	,GLP1Prev12m.MedicationTerm AS [Glucagon-like peptide-1 receptor agonists  Prev12m]
	,GLP1Prev12m.IssueDate

	,GLP1Earliest.MedicationTerm AS [Glucagon-like peptide-1 receptor agonists  Earliest]
	,GLP1Earliest.IssueDate

	,MeglitinidesPrev12m.MedicationTerm AS [Meglitinides Prev12m]
	,MeglitinidesPrev12m.IssueDate

	,MeglitinidesEarliest.MedicationTerm AS [Meglitinides Earliest]
	,MeglitinidesEarliest.IssueDate

	,SGLT2Prev12m.MedicationTerm AS [Sodium glucose co-transporter 2 inhibitors Prev12m]
	,SGLT2Prev12m.IssueDate

	,SGLT2Earliest.MedicationTerm AS [Sodium glucose co-transporter 2 inhibitors Earliest]
	,SGLT2Earliest.IssueDate

	,ThiazolidinedionesPrev12m.MedicationTerm AS [Thiazolidinediones Prev12m]
	,ThiazolidinedionesPrev12m.IssueDate

	,ThiazolidinedionesEarliest.MedicationTerm AS [Thiazolidinediones Earliest]
	,ThiazolidinedionesEarliest.IssueDate

	,CombinationDrugsPrev12m.MedicationTerm AS [Combination drugs Prev12m]
	,CombinationDrugsPrev12m.IssueDate

	,CombinationDrugsEarliest.MedicationTerm AS [Combination drugs Earliest]
	,CombinationDrugsEarliest.IssueDate

	,TestingStripsPrev12m.MedicationTerm AS [Testing Strips Prev12m]
	,TestingStripsPrev12m.IssueDate

	,TestingStripsEarliest.MedicationTerm AS [Testing Strips Earliest]
	,TestingStripsEarliest.IssueDate

	,InsulinPumpEarliest.InsulinPumpEarliestCode AS [Insulin Pump Earliest]
	,InsulinPumpEarliest.InsulinPumpDateRecorded

	,MetforminAE_Earliest.MetforminAE_EarliestCode AS [Metformin AE Earliest]
	,MetforminAE_Earliest.MetforminAE_EarliestDateRecorded

	,NeuropathyPrev12m.MedicationTerm AS [Neuropathy Prev12m]
	,NeuropathyPrev12m.IssueDate

	,NeuropathyEarliest.MedicationTerm AS [Neuropathy Earliest]
	,NeuropathyEarliest.IssueDate


	-----CVD
	,CardiacGlycosidePrev12m.MedicationTerm AS [Cardiac Glycoside Prev12m]
	,CardiacGlycosidePrev12m.IssueDate

	,CardiacGlycosideEarliest.MedicationTerm AS [Cardiac Glycoside Earliest]
	,CardiacGlycosideEarliest.IssueDate

	,ThiazidesPrev12m.MedicationTerm AS [Thiazides Prev12m]
	,ThiazidesPrev12m.IssueDate

	,ThiazidesEarliest.MedicationTerm AS [Thiazides Earliest]
	,ThiazidesEarliest.IssueDate

	,LoopDirueticsPrev12m.MedicationTerm AS [Loop Diruetics Prev12m]
	,LoopDirueticsPrev12m.IssueDate

	,LoopDirueticsEarliest.MedicationTerm AS [Loop Diruetics Earliest]
	,LoopDirueticsEarliest.IssueDate

	,PotassiumSparingPrev12m.MedicationTerm AS [Potassium Sparing Prev12m]
	,PotassiumSparingPrev12m.IssueDate

	,PotassiumSparingEarliest.MedicationTerm AS [Potassium Sparing Earliest]
	,PotassiumSparingEarliest.IssueDate

	,DiureticCombiPrev12m.MedicationTerm AS [Diuretic Combi Prev12m]
	,DiureticCombiPrev12m.IssueDate

	,DiureticCombiEarliest.MedicationTerm AS [Diuretic Combi Earliest]
	,DiureticCombiEarliest.IssueDate

	,DiumidePrev12m.MedicationTerm AS [Diumide Prev12m]
	,DiumidePrev12m.IssueDate

	,DiumideEarliest.MedicationTerm AS [DiumideEarliest]
	,DiumideEarliest.IssueDate

	,AntiArrhythmicsPrev12m.MedicationTerm AS [Anti-Arrhythmics Prev12m]
	,AntiArrhythmicsPrev12m.IssueDate

	,AntiArrhythmicsEarliest.MedicationTerm AS [Anti-Arrhythmics Earliest]
	,AntiArrhythmicsEarliest.IssueDate

	,BetaBlockersPrev12m.MedicationTerm AS [Beta Blockers Prev12m]
	,BetaBlockersPrev12m.IssueDate

	,BetaBlockersEarliest.MedicationTerm AS [Beta Blockers Earliest]
	,BetaBlockersEarliest.IssueDate

	,VasodilatorsPrev12m.MedicationTerm AS [Vasodilators Prev12m]
	,VasodilatorsPrev12m.IssueDate

	,VasodilatorsEarliest.MedicationTerm AS [Vasodilators Earliest]
	,VasodilatorsEarliest.IssueDate

	,CentralantihypertensivePrev12m.MedicationTerm AS [Centralantihypertensive Prev12m]
	,CentralantihypertensivePrev12m.IssueDate

	,CentralantihypertensiveEarliest.MedicationTerm AS [Centralantihypertensive Earliest]
	,CentralantihypertensiveEarliest.IssueDate

	,AlphaBlockersPrev12m.MedicationTerm AS [Alpha Blockers Prev12m]
	,AlphaBlockersPrev12m.IssueDate

	,AlphaBlockersEarliest.MedicationTerm AS [Alpha Blockers Earliest]
	,AlphaBlockersEarliest.IssueDate

	,ACE_InhibitorsPrev12m.MedicationTerm AS [ACE Inhibitors Prev12m]
	,ACE_InhibitorsPrev12m.IssueDate

	,ACE_InhibitorsEarliest.MedicationTerm AS [ACE Inhibitors Earliest]
	,ACE_InhibitorsEarliest.IssueDate

	,AngiotensinAntagonistsPrev12m.MedicationTerm AS [Angiotensin Antagonists Prev12m]
	,AngiotensinAntagonistsPrev12m.IssueDate

	,AngiotensinAntagonistsEarliest.MedicationTerm AS [Angiotensin Antagonists Earliest]
	,AngiotensinAntagonistsEarliest.IssueDate

	,NitratesPrev12m.MedicationTerm AS [Nitrates Prev12m]
	,NitratesPrev12m.IssueDate

	,NitratesEarliest.MedicationTerm AS [Nitrates Earliest]
	,NitratesEarliest.IssueDate

	,CalciumChannelPrev12m.MedicationTerm AS [Calcium Channel Prev12m]
	,CalciumChannelPrev12m.IssueDate

	,CalciumChannelEarliest.MedicationTerm AS [Calcium Channel Earliest]
	,CalciumChannelEarliest.IssueDate

	,AntianginalPrev12m.MedicationTerm AS [Antianginal Prev12m]
	,AntianginalPrev12m.IssueDate

	,AntianginalEarliest.MedicationTerm AS [Antianginal Earliest]
	,AntianginalEarliest.IssueDate

	,PeripheralVasodilatorsPrev12m.MedicationTerm AS [Peripheral Vasodilators Prev12m]
	,PeripheralVasodilatorsPrev12m.IssueDate

	,PeripheralVasodilatorsEarliest.MedicationTerm AS [Peripheral Vasodilators Earliest]
	,PeripheralVasodilatorsEarliest.IssueDate

	,OralAnticoagulantsPrev12m.MedicationTerm AS [Oral Anticoagulants Prev12m]
	,OralAnticoagulantsPrev12m.IssueDate

	,OralAnticoagulantsEarliest.MedicationTerm AS [Oral Anticoagulants Earliest]
	,OralAnticoagulantsEarliest.IssueDate

	,AntiplateletPrev12m.MedicationTerm AS [Antiplatelet Prev12m]
	,AntiplateletPrev12m.IssueDate

	,AntiplateletEarliest.MedicationTerm [Antiplatelet Earliest]
	,AntiplateletEarliest.IssueDate

	,LipidRegulationPrev12m.MedicationTerm AS [Lipid Regulation Prev12m]
	,LipidRegulationPrev12m.IssueDate

	,LipidRegulationEarliest.MedicationTerm AS [Lipid Regulation Earliest]
	,LipidRegulationEarliest.IssueDate

	,StatinAE_Earliest.StatinAE_EarliestCode AS [Statin AE Earliest]
	,StatinAE_Earliest.StatinAE_DateRecorded

	,StatinMuscleEarliest.StatinMuscleEarliestCode AS [Statin Muscle Earliest]
	,StatinMuscleEarliest.StatinMuscleDateRecorded

	,ErectileDysfunctionPrev12m.MedicationTerm AS [Erectile Dysfunction Prev12m]
	,ErectileDysfunctionPrev12m.IssueDate

	,ErectileDysfunctionEarliest.MedicationTerm AS [Erectile Dysfunction Earliest]
	,ErectileDysfunctionEarliest.IssueDate

	,WeightLossPrev12m.MedicationTerm AS [Weight Loss Prev12m]
	,WeightLossPrev12m.IssueDate

	,WeightLossEarliest.MedicationTerm AS [Weight Loss Earliest]
	,WeightLossEarliest.IssueDate

FROM
	CEG.ceg.GenomicsData P

	-- PRESCRIBING
	-----Diabetes
	LEFT JOIN  #InsulinsShortPrev12m	AS	InsulinsShortPrev12m	ON 	InsulinsShortPrev12m.SK_PatientID = P.SK_PatientID
	LEFT JOIN  #InsulinsShortEarliest	AS	InsulinsShortEarliest	ON 	InsulinsShortEarliest.SK_PatientID = P.SK_PatientID

	LEFT JOIN  #InsulinsLongPrev12m	AS	InsulinsLongPrev12m	ON 	InsulinsLongPrev12m.SK_PatientID = P.SK_PatientID
	LEFT JOIN  #InsulinsLongEarliest	AS	InsulinsLongEarliest	ON 	InsulinsLongEarliest.SK_PatientID = P.SK_PatientID

	LEFT JOIN  #SulphonylureasPrev12m	AS	SulphonylureasPrev12m	ON 	SulphonylureasPrev12m.SK_PatientID = P.SK_PatientID
	LEFT JOIN  #SulphonylureasEarliest	AS	SulphonylureasEarliest	ON 	SulphonylureasEarliest.SK_PatientID = P.SK_PatientID

	LEFT JOIN  #MetforminPrev12m	AS	MetforminPrev12m	ON 	MetforminPrev12m.SK_PatientID = P.SK_PatientID
	LEFT JOIN  #MetforminEarliest	AS	MetforminEarliest	ON 	MetforminEarliest.SK_PatientID = P.SK_PatientID

	-------------- Antidiabetic drugs Other
		--LEFT JOIN  #OtherAntidiabeticPrev12m	AS	OtherAntidiabeticPrev12m	ON 	OtherAntidiabeticPrev12m.SK_PatientID = P.SK_PatientID
		--LEFT JOIN  #OtherAntidiabeticEarliest	AS	OtherAntidiabeticEarliest	ON 	OtherAntidiabeticEarliest.SK_PatientID = P.SK_PatientID

	LEFT JOIN  #AGIsPrev12m	AS	AGIsPrev12m	ON 	AGIsPrev12m.SK_PatientID = P.SK_PatientID
	LEFT JOIN  #AGIsEarliest AS	AGIsEarliest ON AGIsEarliest.SK_PatientID = P.SK_PatientID

	LEFT JOIN  #DPP4Prev12m	AS	DPP4Prev12m	ON DPP4Prev12m.SK_PatientID = P.SK_PatientID
	LEFT JOIN  #DPP4Earliest AS	DPP4Earliest ON DPP4Earliest.SK_PatientID = P.SK_PatientID

	LEFT JOIN  #GLP1Prev12m	AS	GLP1Prev12m	ON GLP1Prev12m.SK_PatientID = P.SK_PatientID
	LEFT JOIN  #GLP1Earliest AS	GLP1Earliest ON GLP1Earliest.SK_PatientID = P.SK_PatientID

	LEFT JOIN  #MeglitinidesPrev12m	AS	MeglitinidesPrev12m	ON MeglitinidesPrev12m.SK_PatientID = P.SK_PatientID
	LEFT JOIN  #MeglitinidesEarliest AS	MeglitinidesEarliest ON MeglitinidesEarliest.SK_PatientID = P.SK_PatientID

	LEFT JOIN  #SGLT2Prev12m AS	SGLT2Prev12m ON SGLT2Prev12m.SK_PatientID = P.SK_PatientID
	LEFT JOIN  #SGLT2Earliest AS SGLT2Earliest ON SGLT2Earliest.SK_PatientID = P.SK_PatientID

	LEFT JOIN  #ThiazolidinedionesPrev12m AS ThiazolidinedionesPrev12m ON ThiazolidinedionesPrev12m.SK_PatientID = P.SK_PatientID
	LEFT JOIN  #ThiazolidinedionesEarliest AS ThiazolidinedionesEarliest ON ThiazolidinedionesEarliest.SK_PatientID = P.SK_PatientID

	LEFT JOIN  #CombinationDrugsPrev12m	AS CombinationDrugsPrev12m ON CombinationDrugsPrev12m.SK_PatientID = P.SK_PatientID
	LEFT JOIN  #CombinationDrugsEarliest AS	CombinationDrugsEarliest ON CombinationDrugsEarliest.SK_PatientID = P.SK_PatientID

	LEFT JOIN  #TestingStripsPrev12m	AS	TestingStripsPrev12m	ON 	TestingStripsPrev12m.SK_PatientID = P.SK_PatientID
	LEFT JOIN  #TestingStripsEarliest	AS	TestingStripsEarliest	ON 	TestingStripsEarliest.SK_PatientID = P.SK_PatientID

	LEFT JOIN  #InsulinPumpEarliest	AS	InsulinPumpEarliest	ON 	InsulinPumpEarliest.SK_PatientID = P.SK_PatientID

	LEFT JOIN  #MetforminAE_Earliest	AS	MetforminAE_Earliest	ON 	MetforminAE_Earliest.SK_PatientID = P.SK_PatientID

	LEFT JOIN  #NeuropathyPrev12m	AS	NeuropathyPrev12m	ON 	NeuropathyPrev12m.SK_PatientID = P.SK_PatientID
	LEFT JOIN  #NeuropathyEarliest	AS	NeuropathyEarliest	ON 	NeuropathyEarliest.SK_PatientID = P.SK_PatientID

	--CVD
	LEFT JOIN  #CardiacGlycosidePrev12m	AS	CardiacGlycosidePrev12m	ON 	CardiacGlycosidePrev12m.SK_PatientID = P.SK_PatientID
	LEFT JOIN  #CardiacGlycosideEarliest	AS	CardiacGlycosideEarliest	ON 	CardiacGlycosideEarliest.SK_PatientID = P.SK_PatientID

	LEFT JOIN  #ThiazidesPrev12m	AS	ThiazidesPrev12m	ON 	ThiazidesPrev12m.SK_PatientID = P.SK_PatientID
	LEFT JOIN  #ThiazidesEarliest	AS	ThiazidesEarliest	ON 	ThiazidesEarliest.SK_PatientID = P.SK_PatientID

	LEFT JOIN  #LoopDirueticsPrev12m	AS	LoopDirueticsPrev12m	ON 	LoopDirueticsPrev12m.SK_PatientID = P.SK_PatientID
	LEFT JOIN  #LoopDirueticsEarliest	AS	LoopDirueticsEarliest	ON 	LoopDirueticsEarliest.SK_PatientID = P.SK_PatientID

	LEFT JOIN  #PotassiumSparingPrev12m	AS	PotassiumSparingPrev12m	ON 	PotassiumSparingPrev12m.SK_PatientID = P.SK_PatientID
	LEFT JOIN  #PotassiumSparingEarliest	AS	PotassiumSparingEarliest	ON 	PotassiumSparingEarliest.SK_PatientID = P.SK_PatientID

	LEFT JOIN  #DiureticCombiPrev12m	AS	DiureticCombiPrev12m	ON 	DiureticCombiPrev12m.SK_PatientID = P.SK_PatientID
	LEFT JOIN  #DiureticCombiEarliest	AS	DiureticCombiEarliest	ON 	DiureticCombiEarliest.SK_PatientID = P.SK_PatientID

	LEFT JOIN  #DiumidePrev12m	AS	DiumidePrev12m	ON 	DiumidePrev12m.SK_PatientID = P.SK_PatientID
	LEFT JOIN  #DiumideEarliest	AS	DiumideEarliest	ON 	DiumideEarliest.SK_PatientID = P.SK_PatientID

	LEFT JOIN  #AntiArrhythmicsPrev12m	AS	AntiArrhythmicsPrev12m	ON 	AntiArrhythmicsPrev12m.SK_PatientID = P.SK_PatientID
	LEFT JOIN  #AntiArrhythmicsEarliest	AS	AntiArrhythmicsEarliest	ON 	AntiArrhythmicsEarliest.SK_PatientID = P.SK_PatientID

	LEFT JOIN  #BetaBlockersPrev12m	AS	BetaBlockersPrev12m	ON 	BetaBlockersPrev12m.SK_PatientID = P.SK_PatientID
	LEFT JOIN  #BetaBlockersEarliest	AS	BetaBlockersEarliest	ON 	BetaBlockersEarliest.SK_PatientID = P.SK_PatientID

	LEFT JOIN  #VasodilatorsPrev12m	AS	VasodilatorsPrev12m	ON 	VasodilatorsPrev12m.SK_PatientID = P.SK_PatientID
	LEFT JOIN  #VasodilatorsEarliest	AS	VasodilatorsEarliest	ON 	VasodilatorsEarliest.SK_PatientID = P.SK_PatientID

	LEFT JOIN  #CentralantihypertensivePrev12m	AS	CentralantihypertensivePrev12m	ON 	CentralantihypertensivePrev12m.SK_PatientID = P.SK_PatientID
	LEFT JOIN  #CentralantihypertensiveEarliest	AS	CentralantihypertensiveEarliest	ON 	CentralantihypertensiveEarliest.SK_PatientID = P.SK_PatientID

	LEFT JOIN  #AlphaBlockersPrev12m	AS	AlphaBlockersPrev12m	ON 	AlphaBlockersPrev12m.SK_PatientID = P.SK_PatientID
	LEFT JOIN  #AlphaBlockersEarliest	AS	AlphaBlockersEarliest	ON 	AlphaBlockersEarliest.SK_PatientID = P.SK_PatientID

	LEFT JOIN  #ACE_InhibitorsPrev12m	AS	ACE_InhibitorsPrev12m	ON 	ACE_InhibitorsPrev12m.SK_PatientID = P.SK_PatientID
	LEFT JOIN  #ACE_InhibitorsEarliest	AS	ACE_InhibitorsEarliest	ON 	ACE_InhibitorsEarliest.SK_PatientID = P.SK_PatientID

	LEFT JOIN  #AngiotensinAntagonistsPrev12m	AS	AngiotensinAntagonistsPrev12m	ON 	AngiotensinAntagonistsPrev12m.SK_PatientID = P.SK_PatientID
	LEFT JOIN  #AngiotensinAntagonistsEarliest	AS	AngiotensinAntagonistsEarliest	ON 	AngiotensinAntagonistsEarliest.SK_PatientID = P.SK_PatientID

	LEFT JOIN  #NitratesPrev12m	AS	NitratesPrev12m	ON 	NitratesPrev12m.SK_PatientID = P.SK_PatientID
	LEFT JOIN  #NitratesEarliest	AS	NitratesEarliest	ON 	NitratesEarliest.SK_PatientID = P.SK_PatientID

	LEFT JOIN  #CalciumChannelPrev12m	AS	CalciumChannelPrev12m	ON 	CalciumChannelPrev12m.SK_PatientID = P.SK_PatientID
	LEFT JOIN  #CalciumChannelEarliest	AS	CalciumChannelEarliest	ON 	CalciumChannelEarliest.SK_PatientID = P.SK_PatientID

	LEFT JOIN  #AntianginalPrev12m	AS	AntianginalPrev12m	ON 	AntianginalPrev12m.SK_PatientID = P.SK_PatientID
	LEFT JOIN  #AntianginalEarliest	AS	AntianginalEarliest	ON 	AntianginalEarliest.SK_PatientID = P.SK_PatientID

	LEFT JOIN  #PeripheralVasodilatorsPrev12m	AS	PeripheralVasodilatorsPrev12m	ON 	PeripheralVasodilatorsPrev12m.SK_PatientID = P.SK_PatientID
	LEFT JOIN  #PeripheralVasodilatorsEarliest	AS	PeripheralVasodilatorsEarliest	ON 	PeripheralVasodilatorsEarliest.SK_PatientID = P.SK_PatientID

	LEFT JOIN  #OralAnticoagulantsPrev12m	AS	OralAnticoagulantsPrev12m	ON 	OralAnticoagulantsPrev12m.SK_PatientID = P.SK_PatientID
	LEFT JOIN  #OralAnticoagulantsEarliest	AS	OralAnticoagulantsEarliest	ON 	OralAnticoagulantsEarliest.SK_PatientID = P.SK_PatientID

	LEFT JOIN  #AntiplateletPrev12m	AS	AntiplateletPrev12m	ON 	AntiplateletPrev12m.SK_PatientID = P.SK_PatientID
	LEFT JOIN  #AntiplateletEarliest	AS	AntiplateletEarliest	ON 	AntiplateletEarliest.SK_PatientID = P.SK_PatientID

	LEFT JOIN  #LipidRegulationPrev12m	AS	LipidRegulationPrev12m	ON 	LipidRegulationPrev12m.SK_PatientID = P.SK_PatientID
	LEFT JOIN  #LipidRegulationEarliest	AS	LipidRegulationEarliest	ON 	LipidRegulationEarliest.SK_PatientID = P.SK_PatientID

	LEFT JOIN  #StatinAE_Earliest	AS	StatinAE_Earliest	ON 	StatinAE_Earliest.SK_PatientID = P.SK_PatientID
	LEFT JOIN  #StatinMuscleEarliest	AS	StatinMuscleEarliest	ON 	StatinMuscleEarliest.SK_PatientID = P.SK_PatientID

	LEFT JOIN  #ErectileDysfunctionPrev12m	AS	ErectileDysfunctionPrev12m	ON 	ErectileDysfunctionPrev12m.SK_PatientID = P.SK_PatientID
	LEFT JOIN  #ErectileDysfunctionEarliest	AS	ErectileDysfunctionEarliest	ON 	ErectileDysfunctionEarliest.SK_PatientID = P.SK_PatientID

	LEFT JOIN  #WeightLossPrev12m	AS	WeightLossPrev12m	ON 	WeightLossPrev12m.SK_PatientID = P.SK_PatientID
	LEFT JOIN  #WeightLossEarliest	AS	WeightLossEarliest	ON 	WeightLossEarliest.SK_PatientID = P.SK_PatientID

WHERE
	P.CCG = '08V'

GO


-- drop all used temp tables
BEGIN -- drop all temp tables if exist

	DECLARE @DropGlobal bit=0 --Default dont drop global temp table

	DECLARE @DROP_STATEMENT nvarchar(1000)
	DECLARE cursorDEL CURSOR FOR
	SELECT 'DROP TABLE '
	   + case
			   when name like '##%' then name
			   when name like '#%' then SUBSTRING(name, 1, CHARINDEX( '____', name)-1)
		end as DropSQL
	from tempdb..sysobjects
	WHERE name LIKE '#%'
	   AND OBJECT_ID('tempdb..' + name) IS NOT NULL
	   AND name not like case
						   when @DropGlobal=0 then '##%' --//Exclude global temp
						   else '#######%'    --//some fack expression so we can
											   --//select global temp for delete
						end
	OPEN cursorDEL
	FETCH NEXT FROM cursorDEL INTO @DROP_STATEMENT
	WHILE @@FETCH_STATUS = 0
	BEGIN
	EXEC (@DROP_STATEMENT)
	--print @DROP_STATEMENT
	FETCH NEXT FROM cursorDEL INTO @DROP_STATEMENT
	END
	CLOSE cursorDEL
	DEALLOCATE cursorDEL

END

GO
```
