## Supplementary File 1 for "Cohort Profile: East London Genes & Health (ELGH), a community based population genomics and health study of British-Bangladeshi and British-Pakistani people"

Barts Health NHS
NHS Trust

East London Genes & Health

Queen Mary
University of London

www.genesandhealth.org | 020 7882 2366 |

Study ID
ELGH

### Volunteer questionnaire | Stage 1

Thank you for taking part in this study. We would like some further information

**Please complete all the information below**

**Title:** ○ Mr ○ Mrs ○ Miss ○ Ms ○ Dr Other (please state) ________

**First names:**

**Last name:**

**Date of Birth:** DD / MM / YYYY

**How would you define your ethnic group?**
○ Bangladeshi or British-Bangladeshi ○ Pakistani or British-Pakistani
○ Other (please state) ________

**Are you** ○ Male ○ Female

**Your home address:**

**Postcode** -

**Your email address:**

**NHS Number (if known):** - -

**Your mobile phone number:**

**Your home phone number:**

**Your work phone number:**

**Name of GP or GP surgery:**

**Address of GP surgery:**

**Do you have diabetes?** ○ Yes ○ No ○ Don't Know

**Were your parents related by blood? (not just by marriage - for example, cousins)** ○ Yes ○ No ○ Don't Know

If Yes, how were your parents related? ○ First cousins ○ Don't Know
○ Other related by blood - please state if known ________

**Do you consider your health and wellbeing to be** ○ Poor ○ Average ○ Good

0656507256
